# Museum of Spatial Transcriptomics

**DOI:** 10.1101/2021.05.11.443152

**Authors:** Lambda Moses, Lior Pachter

**Affiliations:** Division of Biology and Biological Engineering, California Institute of Technology, Pasadena, CA 91125, USA; Department of Computing and Mathematical Sciences, California Institute of Technology, Pasadena, CA 91125, USA

## Abstract

The function of many biological systems, such as embryos, liver lobules, intestinal villi, and tumors depends on the spatial organization of their cells. In the past decade high-throughput technologies have been developed to quantify gene expression in space, and computational methods have been developed that leverage spatial gene expression data to identify genes with spatial patterns and to delineate neighborhoods within tissues. To assess the ability and potential of spatial gene expression technologies to drive biological discovery, we present a curated database of literature on spatial transcriptomics dating back to 1987, along with a thorough analysis of trends in the field such as usage of experimental techniques, species, tissues studied and computational approaches used. Our analysis places current methods in historical context, and we derive insights about the field that can guide current research strategies. A companion supplement offers a more detailed look at the technologies and methods analyzed: https://pachterlab.github.io/LP_2021/.

## 1 Introduction

It has long been recognized that in biological systems ranging from the *Drosophila* embryo to the hepatic lobule, many genes need to be properly regulated in space for the system to function. In order to study the spatial patterns of gene expression, many different spatial transcriptomics methods, which produce spatially localized quantification of mRNA transcripts as proxies for gene expression, have been developed. Thanks to growing interest in the field, several reviews have been written in the past 5 years, providing overviews of experimental techniques for data collection [1, 2], and describing how such techniques can be applied to specific biological systems, e.g. tumors [3], brain [4], and liver [5]. These reviews typically begin with either laser capture microdissection (LCM) or single molecular fluorescent *in situ* hybridization (smFISH) in the late 1990s, although the quest to profile the transcriptome in space is much older. The methods underlying such “prequel” technologies are important because many of them are used, in updated forms, in modern “current era” high-throughput methods.

Some important technologies enabling spatial transcriptomics date back to the 1970s (Supplementary Material; Chapter 2). Various forms of *in situ* hybridization (ISH) have been used for a long time to visualize gene expression in space. Radioactive ISH was first introduced in 1969, visualizing ribosomal RNA [6] and DNA [7] in *Xenopus laevis* oocytes, and was first used to visualize transcripts of specific genes (globin) in 1973 [8] (Figure 1A). Non-radioactive fluorescent or colorimetric ISH was developed in the 1970s and the early 1980s, improving spatial resolution, enabling 3D staining, and shortening required exposure times [9, 10] (Figure 1A). Early ISH was performed in tissue sections, making it challenging to apply to blastrulas and to reconstruct 3D tissue structures; whole mount ISH (WMISH) was first introduced in *Drosophila* in 1989 [11] and was soon adapted to other species such as mice in the early 1990s [12].

**Figure 1:**
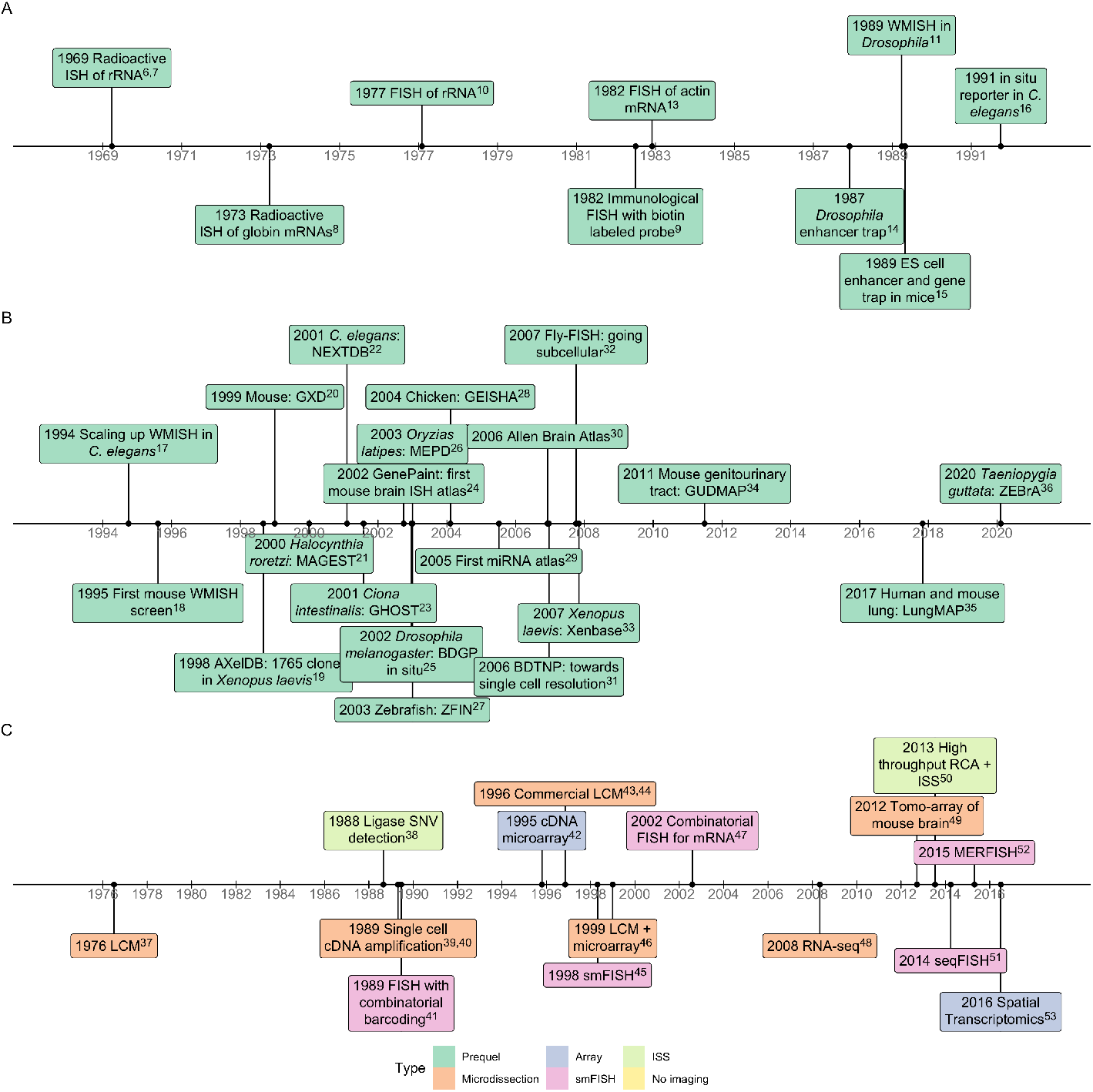
A) Timeline of development of prequel era technologies. References: 1969 radioactive ISH [6, 7], 1973 goblin [8], 1977 FISH [10], 1982 immunological [9], 1982 FISH [13], 1987 enhancer trap [14], 1989 WMISH [11], 1989 ES cell [15], 1991 *C. elegans* [16]. B) Timeline of major (WM)ISH atlases and gene expression pattern databases. References: 1994 WMISH [17], 1995 mouse WMISH [18], 1998 AXelDB [19], 1999 GXD [20], 2000 MAGEST [21], 2001 NEXTDB [22], 2001 GHOST [23], 2002 GenePaint [24], 2002 BDGP [25], 2003 MEPD [26], 2003 ZFIN [27], 2004 GEISHA [28], 2005 miRNA [29], 2006 Allen [30], 2006 BDTNP [31], 2007 Fly-FISH [32], 2007 Xenbase [33], 2011 GUDMAP [34], 2017 LungMAP [35], 2020 ZEBrA [36]. C) Timeline of development of current era technologies and their notable precursors, colored by type of technology. References: 1976 LCM [37], 1988 ligase mediated single nucleotide variant (SNV) detection [38], 1989 amplification [39, 40], 1989 FISH [41], 1995 microarray [42], 1996 LCM [43, 44], 1998 smFISH [45], 1999 LCM [46], 2002 combinatorial [47], 2008 RNA-seq [48], 2012 Tomo-array [49], 2013 ISS [50], 2014 seqFISH [51], 2015 MERFISH [52], 2016 ST [53].

Another strand of development in early spatial transcriptomics was the enhancer and gene trap screen which was developed in the 1980s when DNA sequencing throughput was increasing [54] and metazoan genomes were newly opened frontiers. The first screens in *Drosophila* [14] and mice [15] were performed in the late 1980s in order to visualize expression of untargeted, and often previously unknown, genes. These could be identified with 5’ rapid amplification of cDNA ends (RACE) polymerase chain reaction (PCR) followed by Sanger sequencing of the PCR products. A related technology, *in situ* reporter, in which a promoter of a known gene or a predefined genomic fragment drives the expression of a reporter, was first used in a gene expression screen in *C. elegans* in 1991 [16]. With increasing throughput, enhancer and gene traps became the technology of choice for spatial transcriptomics in the 1990s, until the rise of (WM)ISH in the late 1990s which leveraged automation. WM(ISH) also avoided the need for transgenic lines, and was facilitated by the widespread availability of reference genomes in the early 2000s, which could be used for computational probe design. Although now eclipsed by newer methods, enhancer trap, gene trap, and *in situ* reporter methods have been used to build reference databases of gene expression and enhancer usage patterns in transgenic lines throughout the 2000s and 2010s [55, 56].

The foundation for many current era technologies was built in the decades between the 1970s and the 2000s (Figure 1C). For example, UV laser was first used to cut tissue in 1976 [37]. Popular IR and UV LCM systems were first reported in 1996 [43, 44] and were soon commercialized. Microarray technology was first reported in 1995 [42], and was originally used to quantify transcripts hybridized to cDNAs printed on a slide, but it was soon adapted to quantify the transcriptome from LCM samples in 1999 [46]. Popular current era technologies such as Spatial Transcriptomics (ST) [53] and 10X Visium rely on such microarray technology, but to capture transcripts from the tissue mounted on the microarray slides rather than from solution. Some highly multiplexed sm-FISH technologies such as seqFISH [51] rely on combinatorial barcoding, i.e. encoding each gene with a combination of colors so transcripts of more genes than easily discernible colors (up to 5) can be quantified simultaneously. Combinatorial barcoding was first reported in immunological DNA FISH in 1989 [41] and was first used for transcripts in 2002 [47]. The first unequivocal demonstration of smFISH showing each mRNA molecule as a spot was reported in 1998 [45]. Highly multiplexed smFISH would not have been possible without the development of these technologies.

(WM)ISH was the technology of choice in the late 1990s and the 2000s before the rise of highly multiplexed, high resolution, and more quantitative technologies, and has been used to create gene expression atlases in embryos of several species such as *Drosophila melanogaster* [25], *Mus musculus*, and *Gallus gallus* [28], in various mouse organs such as the brain [30], genitourinary tract [34], and lung [35], and for specific types of genes such as miRNAs [29] (Figure 1B). For species other than mice and humans, organs other than the brain, and miRNAs, the only spatial transcriptomics resources currently available are for the most part (WM)ISH atlases. Model organism databases collecting the proliferating gene expression patterns from various sources were also established in this period, such as gene expression database (GXD) [57] and Zebrafish Information Network (ZFIN) [58] (Figure 1B). The golden age of (WM)ISH seems to have ended in the 2010s (Figure 1B), perhaps due to some of the disadvantages of (WM)ISH, such as requiring stereotypical tissue structure, the need for thousands of animals to generate an atlas, and the largely qualitative nature of results.

Early motivating applications for spatial transcriptomics included identification of genes with restricted patterns which indicated function in development, identification of novel cell type markers, and identification of novel cell types not evident from tissue morphology [14, 15]. In the 1980s and 1990s, analyses were typically done manually, although more recently automated methods have been developed (Supplementary Material Chapter 3). Recent developments in machine learning, coupled to more powerful computing infrastructure and the availability of more quantitative data, have opened up new possibilities. However, the legacy of the prequel era still lives on; current era studies still frequently reference prequel atlases [59–61].

## 2 Data collection

Current era technologies broadly fall into five categories: Microdissection (Supplementary Material Section 5.1), smFISH (Supplementary Material Section 5.2), *in situ* sequencing (ISS) (Supplementary Material Section 5.3), array (Supplementary Material Section 5.4), and no imaging (Supplementary Material Section 5.5). Developers of such technologies often seek to enable a trifecta of transcriptome wide profiling, single cell resolution, and high gene detection efficiency. While this achievement appears to be increasingly within reach, current era technologies are characterized by trade offs between these goals.

### 2.1 Microdissection

Current era microdissection constitutes the compilation of spatial locations of samples during the course of microdissection, and the assaying of transcriptomes of samples by cDNA microarray or RNA-seq. Since 1999, by far the most widely used microdissection technology is LCM, which has been applied to various biological fields such as oncology, neuroscience, immunology, developmental biology, and botany (see Supplementary Material Chapter 6 for topic modeling of PubMed LCM literature). In UV LCM, sometimes also called laser microbeam microdissection (LMM) or laser microdissection (LMD), a UV laser ablates a narrow strand of tissue around the desired area, which is then collected into tubes by gravity (Leica LMD) or laser pressure catapult (LPC, as in Zeiss PALM microbeam) (Figure 2B, C). In IR LCM (Arcturus PixCell II), the tissue section is mounted on a plastic membrane on tube caps. The IR laser briefly heats the desired area, melting the membrane so the tissue in the area is fused to the cap and captured (Figure 2A). IR and UV can be combined; for example UV can be used to cut the tissue and IR to remove the desired area with the plastic membrane (recent versions of Arcturus).

**Figure 2:**
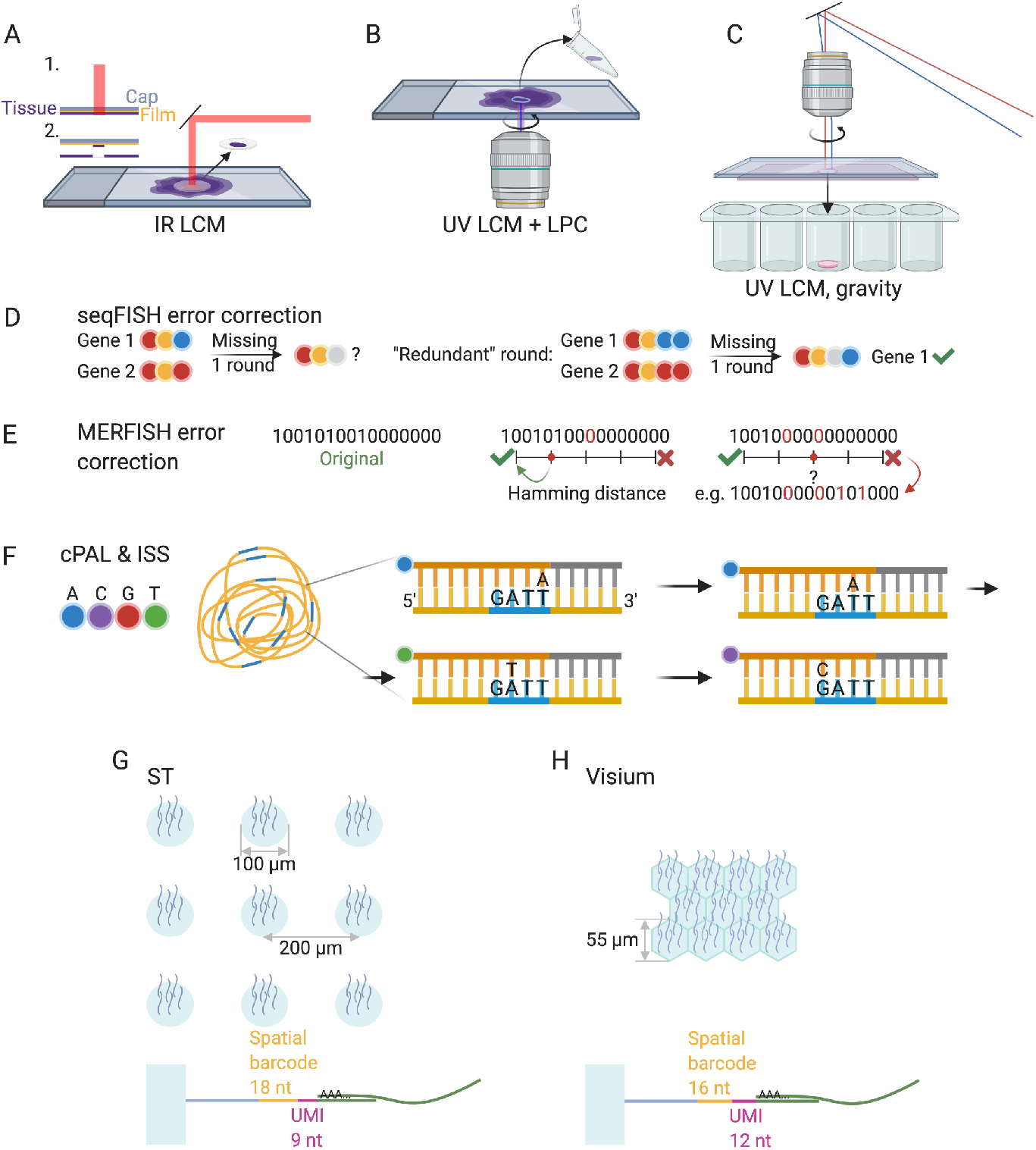
Schematics of common current era technologies. A) IR LCM. B) UV LCM with LPC. C) UV LCM removing cut era by gravity. D) seqFISH barcoding and error correction scheme. E) MERFISH Hamming distance 4 barcoding and error correction scheme. F) Cartana ISS with cPAL sequencing. Yellow line stands for the RCA amplicon. Short blue lines stand for the gene barcode. Brown stands for the probe; bases not labeled are degenerate. Gray stands for primer matching constant region. G) Top: ST spot diameter and spacing. Bottom: barcode and unique molecular identifier (UMI) lengths; the light blue block denotes the glass slide. H) Similar to G, but for Visium. Created with BioRender.com

Advantages of LCM include transcriptome wide profiling, precise cuts informed by histology, compatibility with formalin fixed paraffin embedded (FFPE) tissues [62], and the possibility of applying the method to single cells from both frozen [63] and FFPE sections [64]. LCM can also be applied to 3D tissues by microdissecting non-overlapping domains as in Geo-seq [65]. Disadvantages of LCM include difficulty to scale to larger number of samples, exclusion of whole mount samples, and potential RNA degradation [66]. While LCM is widely used to capture targeted areas according to histology, it can also cut tissue into an untargeted grid [67].

Other microdissection techniques generally fall into two categories: Mechanical (Supplementary Material 5.1.4) and optical (Supplementary Material 5.1.5). The former includes 2000s voxelation [68] (Figure 4), and Tomo-seq [69], which sections a tissue with a cryotome along an axis of interest, followed by RNA-seq on each section. Optical microdissection includes GeoMX DSP [70], which shines UV light on regions of interest (ROIs) to release photo-cleavable gene barcodes for quantification, and Niche-seq [71], which uses fluorescence activated cell sorting (FACS) to isolate cells expressing photoactivated GFP in transgenic mice for scRNA-seq.

### 2.2 Single Molecule FISH

Chronologically, the next technology developed in the current era is highly multiplexed single-molecule FISH (smFISH), which began with a 2012 prototype (seqFISH) that relied on super-resolution microscopy (SRM) to simultaneously profile 32 genes in yeast by hybridizing probes with different colors to transcripts, and then deducing relative locations of the colors present [72]. SRM is no longer needed; in 2014 seqFISH [51] was published, in which one color per gene is visualized per round of hybridization and the probes are stripped before the next round for the next color in the barcode. With 4 colors, 8 rounds of hybridization (4^8^ = 65536) are more than enough to encode all genes in the human or mouse genome. In practice, an error correcting round of hybridization is performed, so that genes can still be distinguished if signal from one round of hybridization is missing [73] (Figure 2D). More recently in a version of seqFISH based on RNA SPOTS [74], the “colors” themselves are one hot encoded by a sequence of hybridizations, expanding the palette to 20 “colors” per channel and enabling the profiling of 10,000 genes [75].

Another smFISH technique is multiplexed error-robust FISH MERFISH [52], which uses a different barcoding strategy, in which each gene is encoded by a binary code. The color codes in each experiment must be separated by a Hamming distance (HD) of 4 to allow for correction of missing signal in one round, and by 2 to identify error without the facility for correcting it (Figure 2D). The length of barcodes can be increased to encode 10,000 genes [76]. As only the fluorophores are removed but the probes are not stripped, numerous rounds of hybridization in MERFISH are less time consuming than those in seqFISH. Most other smFISH based techniques, such as HybISS [77] and split-FISH [78], use either seqFISH-like or MERFISH-like barcoding.

Advantages of smFISH based techniques include high gene detection efficiency (~95% for Hamming distance 4 MERFISH [79] compared to smFISH), single cell resolution, and subcellular transcript localization, which can be biologically relevant [80, 81]. Single round smFISH has nearly 100% detection efficiency [72], and multiple rounds of hybridization tend to decrease the efficiency in part because barcodes with incorrigible errors are discarded. Disadvantages include requirement of pre-defined gene panel and probes, difficulty in probing shorter transcripts, lengthy imaging time, limited scalability to large area of tissues, possible challenges in cell segmentation, and the need to process terabytes of images.

Challenges with smFISH have been addressed by various methods: Signal to noise ratio can be improved with rolling circle amplification (RCA) [77], branched DNA (bDNA) [82], hybridization chain reaction (HCR) [73], and tissue clearing [83]. With increasing number of genes profiled, the transcript spots are increasing likely to overlap, causing optical crowding. This can be mitigated by expansion microscopy (ExM) [84], only imaging a subset of probes at a time and using computational super-resolution [75], imaging highly expressed genes without combinatorial barcoding [52], and computationally resolving overlapping spots [85]. While smFISH based techniques are typically designed for frozen sections, SCRINSHOT is designed for FFPE sections [86].

### 2.3 *In Situ* Sequencing

ISS methods yield spatial transcriptome information by sequencing, typically by ligation (SBL), gene barcodes (targeted) or short fragments of cDNAs (untargeted) *in situ*. Such methods rely on ligase only joining two pieces of DNA – a primer with known sequence and a probe – if they match the template, with non-matching probes washed away. The probes used are degenerate except for one or two query bases encoded by a color. The 2013 ISS [50], later commercialized by Cartana, uses one query base per probe as in cPAL [87] to sequence gene barcodes (Figure 2E). FISSEQ [88] and a later adaptation with ExM called ExSeq [89] use SOLiD, which uses two query bases per probe to sequence circularized and RCA amplified cDNAs. In STARmap [90], gene barcodes are sequenced by SEDAL, in which SOLiD-like two query bases are also used to reject error, but one base encoding can also be used [90]. BARseq also RCA amplifies probes with gene barcodes, but uses sequencing by synthesis (SBS) instead of SBL to sequence the barcodes [91].

Advantages of ISS include single cell resolution and subcellular transcript localization, as ISS displays each mRNA molecule as a spot. The ISS technologies mentioned above use RCA, which greatly amplifies signal, but misses many transcripts. This can be an advantage, as brighter and less crowded spots allow ISS to be applied to larger tissue areas such as whole mouse brain sections, with lower magnification (20x, while MERFISH uses 60x) [92, 93]. However, ISS techniques tend to have low detection efficiency. Whereas detection efficiency of scRNA-seq techniques is between 3% - 25% [94–98], the detection efficiencies of Cartana ISS and FISSEQ [99] are ~5% and ~1% respectively, with STARmap only marginally better. However, ExSeq claims up to 62% efficiency compared to smFISH, i.e. for genes of interest in the same cell type, ExSeq detects around 62% as many transcripts as smFISH. Untargeted ExSeq can be transcriptome wide, but the targeted ISS techniques have only been used for up to 1020 genes [90] though more typically fewer than 300 genes [89], perhaps due to limited read length of SBL and challenges of imaging and image processing as in smFISH.

### 2.4 Arrays

Spatial locations of transcripts can also be preserved by capturing the transcripts from tissue sections on *in situ* arrays. Such arrays can be manufactured by printing spot barcodes, UMIs, and poly-T oligos on commercial microarray slides to capture polyadenylated transcripts, as in the ST and Visium technologies (Figure 2G, H). They can also be Drop-seq-like beads [94] with split pool barcodes, UMIs, and poly-T oligos spread on slides in a single layer (e.g., Slide-Seq [100]) or confined in wells etched on the slides (e.g., HDST [101]), with bead barcodes subsequently located using *in situ* SBL. Alternatively, in DBiT-seq [102], an array is generated by microfluidic channels, which are used to deposit one type of barcode in one direction, and then another in a perpendicular direction, with the orthogonal barcodes ligated so each spot can be identified with a unique pairwise combination. While array based techniques are typically designed for frozen sections and 3’ end Illumina sequencing, Visium has recently been adapted to FFPE sections [103] and Nanopore long read sequencing [104].

Array based techniques have been applied to large areas of tissue [61], and their use is increasing (Figure 3A). Nevertheless, they do not have single cell resolution, and the spot diameter of ST is 100 μm with spots 200 μm apart (Figure 2G). Visium, which is an improved version of ST released after 10X Genomics acquired ST, has spots in a hexagonal array with diameter 55 μm (Figure 2H). Bead diameter is 10 μm in Slide-seq, and 2 μm in HDST. Slide-seq and HDST use bead size smaller than single cells, however they still don’t provide single cell resolution because one bead can span two or more cells. Resolution of DBiT-seq is determined by channel width (either 50, 25, or 10 μm). More recently, the spot size can be reduced to below 1 μm, with RCA amplified DNA nanoballs as small as 0.22 μm across with spot barcodes deposited in wells 0.5 or 0.715 μm apart in Stereo-seq [105], and in Seq-Scope polonies with spatial barcodes 0.5 μm in diameter on an Illumina flow cell re-purposed to capture transcripts from tissue sections [106]. Another polony based method PIXEL-seq achieves spot diameter of about 1.22 μm but unlike in the flow cell, PIXEL-seq does not have much spacing around each polony [107].

**Figure 3:**
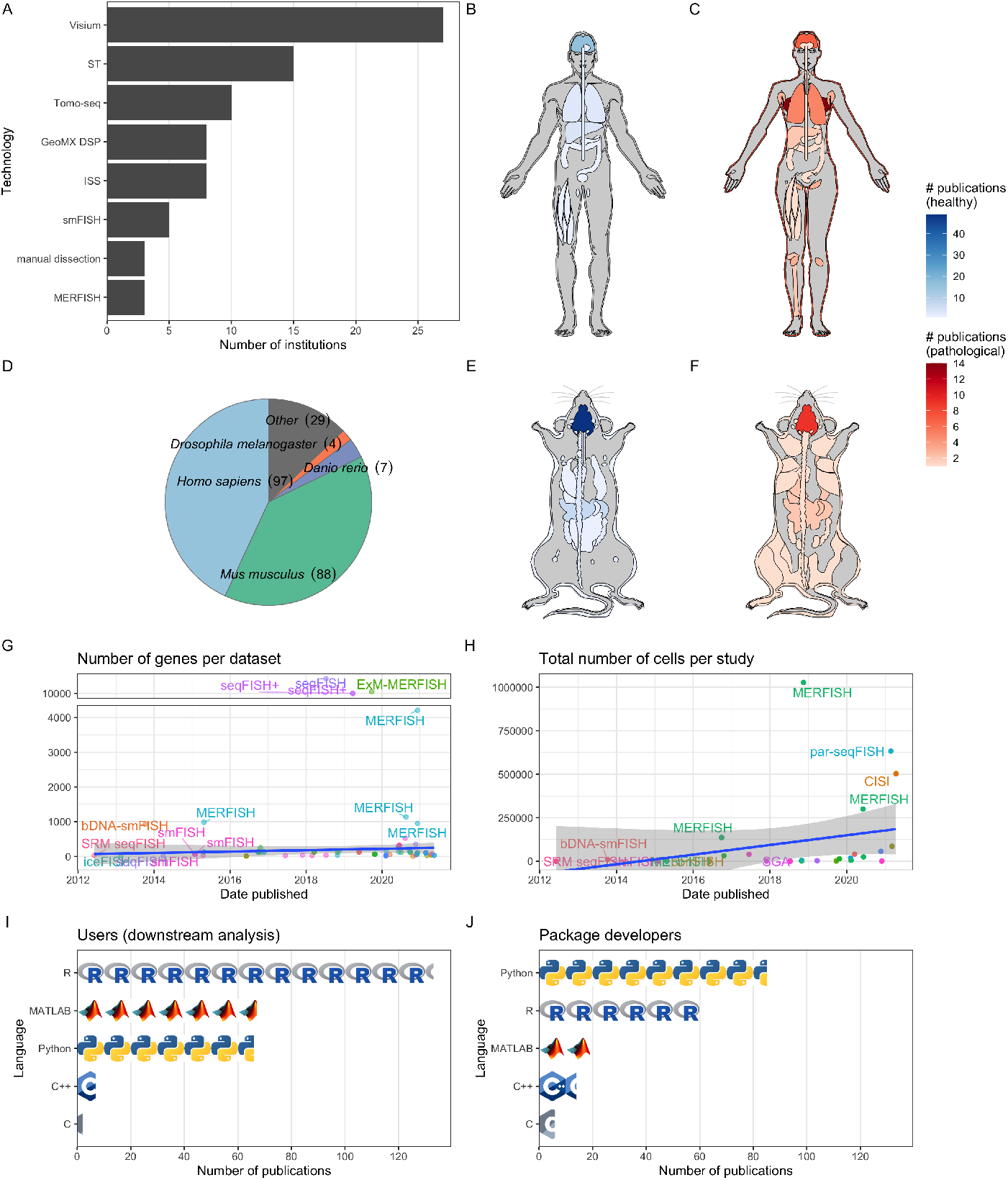
A) Number of institutions that have published papers or preprints with each technique, excluding LCM. Only techniques used by at least 3 institutions are shown. B) Number of publications for each healthy organ in human (male shown here, as there is no study on healthy female specific organs in humans at present). C) Number of publications for pathological organs in human (female shown here, but there are two studies on prostate cancer [146, 147]). D) Number of publication per species. E) Number of publications per healthy organ in the mouse. F) Number of publications for pathological organs in mouse. G) Number of genes per dataset over time. The studies profiling 10,000 genes are shown with broken y-axis to better show the trend among more ordinary studies. Gray ribbon in G and H stands for 95% confidence interval and the trend line is fitted without the 10,000 gene datasets. H) Total number of cells per study profiled by smFISH based techniques over time among studies that reported the number of cells. All studies that reported number of cells are shown. In C and D, the slopes of the fitted line do not significantly differ from 0 (t-test). I) Number of publications for data collection using each of the 5 most popular programming languages for downstream data analysis. J) Number of publication for data analysis using each of the 5 most popular programming languages for package development. In both A and B, each icon stands for 10 publications. Note that multiple programming languages can be used in one publication.

**Figure 4:**
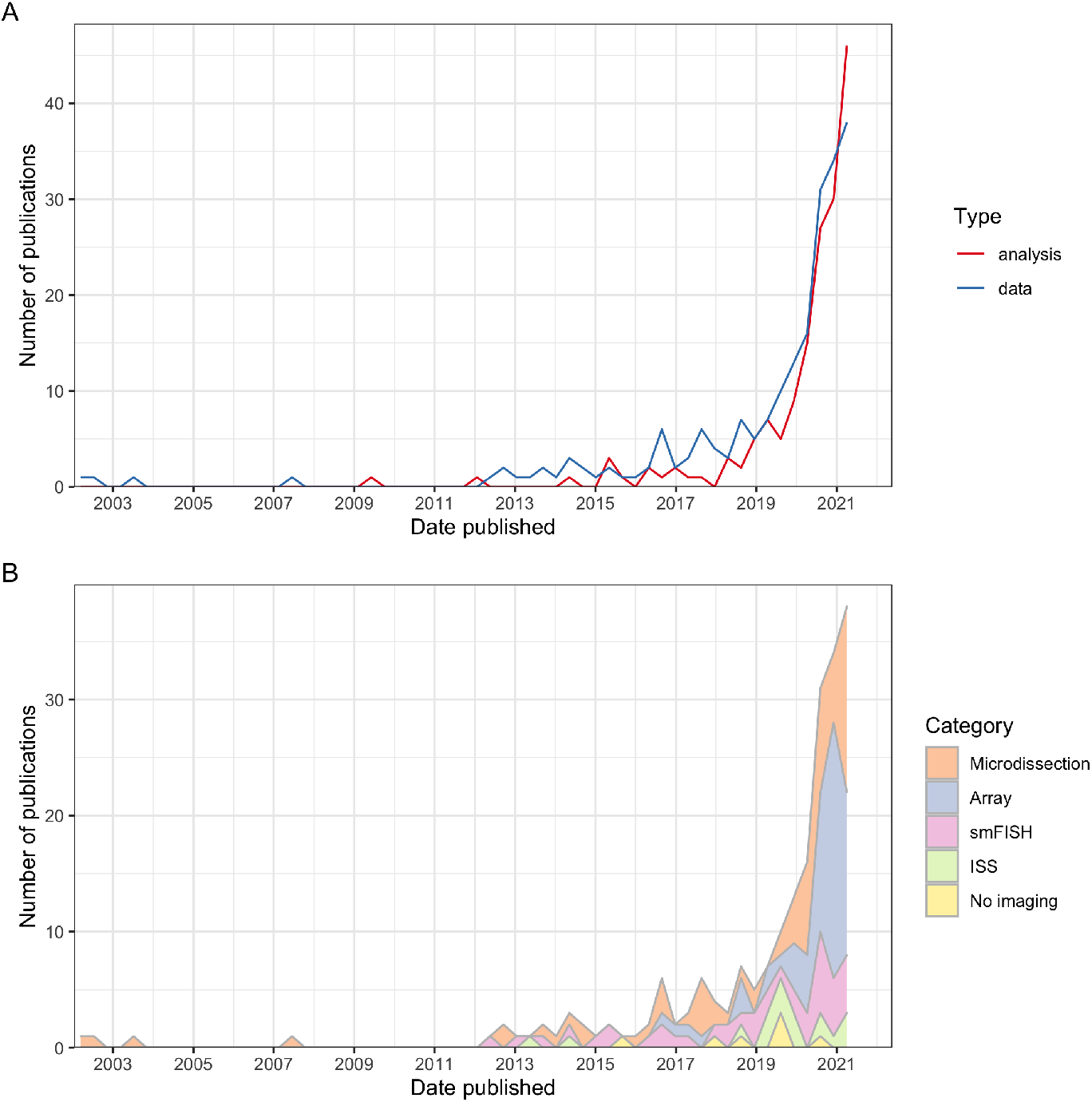
A) Number of publication over time for current era data collection and data analysis. Bin width is 120 days. Non-curated LCM literature is excluded. B) The data collection curve in A, broken down by category of techniques. The colors are stacked and sorted in descending order of total number of publications using techniques in that category.

Array based techniques tend to have low detection efficiency. The efficiency of ST is estimated to be 6.9% compared to smFISH for select genes in the same tissue type, comparable to that of scRNA-seq. Visium’s efficiency seems to be moderately higher than that of ST, and DBiT-seq’s is even higher, at ~15.5% compared to smFISH. Slide-seq and HDST are much less efficient. Efficiency of Slide-seq1 is only ~2.7% of that of Drop-seq, while Slide-seq2 is on par with Drop-seq [108]. Efficiency of HDST is ~1.3% per bead compared to smFISH. Efficiencies of the submicron techniques, in number of UMIs per unit area in the same tissue, might be comparable to that of Visium [107].

### 2.5 Methods that don’t rely on imaging

Some technologies have been developed to preserve information necessary to computationally reconstruct spatial gene expression patterns without imaging. One such technology is DNA microscopy [109], which records proximity between cDNAs. This information can be used to reconstruct relative locations of transcripts. At the cellular level, gene expression in rare cell types can be reconstructed by deliberately assaying multiplets, and then mapping them to locations in a spatial reference based on gene expression of cells from common cell types attached to cells from the rare cell types [110, 111]. Variants of the term “spatial transcriptomics” have also been used to describe techniques localizing transcripts to organelles (e.g., APEX-seq [112]), although no spatial coordinates are recorded.

### 2.6 Multi-omics

Another direction of development is multi-omics (Supplementary Material 4.6). Oligo-tagged antibodies are used to detect proteins of interest, and the oligonucleotide signifying the protein species can be detected with smFISH based methods. Such antibody panels have been combined with a variant of ST as SM-Omics [113], GeoMX DSP [70], and MERFISH [90]. A disadvantage of antibody panels is that the number of proteins profiled is limited to at most a few dozens. MERFISH [114] and seqFISH+ [115] have been adapted to visualize chromatin structure, by targeting DNA genomic loci [114] or introns of nascent transcripts [114, 115]. Multiplexed transcript quantification can also be combined with neuron projection tracing. For instance, cholera toxin subunit b (CTb) retrograde tracing has been used in conjunction with MERFISH to visualize axons [116]. Also, BARseq was originally designed to use ISS for axon tracing by sequencing neuron specific barcodes introduced by a virus injected into the brain, but was later adapted to sequence gene barcodes as well.

## 3 Data analysis

The processing and analysis of high-throughput spatial transcriptomics data requires novel methods and tools, especially for problems such as image preprocessing, spatial reconstruction of scRNA-seq data, cell type deconvolution of array-based data, identification of spatially variable genes, and inference of cell-cell interactions.

For smFISH and ISS based data, the raw data consists of images of fluorescent spots, which must be processed to identify transcript spots, match spots to genes, and assign spots to cells (Section 5.1). SmFISH and ISS studies often use classical image processing tools such as top-hat filtering to remove background, translation to align images from different rounds of hybridization, and watershed for cell segmentation [73, 79, 90]. Machine learning in Ilastik, deep learning packages like DeepCell [117], and alternative tools incorporating scRNA-seq data [118], can also be used for cell segmentation. However, without visualizing the plasma membrane, accuracy of cell segmentation is limited. Some analyses, such as identification of tissue regions, can be performed without cell segmentation [118]. Until 2019, image processing was typically performed with poorly documented and technique specific code written in the proprietary language MATLAB, but more recently such code is increasingly written in the open source language Python. The package `starfish` [119] was developed as an attempt to provide a unified and well-documented user interface to process images from different techniques such as seqFISH, MERFISH, and ISS, but it has not yet been widely adopted.

Recently, improvements in scRNA-seq technology have inspired new methods for leveraging the complementary nature of high-resolution transcriptome quantification with spatial transcriptomics data. For smFISH and ISS data that is not transcriptome wide, expression patterns of genes not profiled in the spatial data can be imputed with scRNA-seq data, either by mapping dissociated scRNA-seq cells to the spatial reference or by directly imputing gene expression in space using expression profiles from scRNA-seq (Supplementary Material 7.3). Cells can be mapped to spatial locations on an existing spatial dataset with genes shared by the two datasets, with an *ad hoc* score favoring similarity between cell and location [60] or via optimal transport modeling [120]. While *ad hoc* scoring is simple to implement, the results tend to be qualitative. Alternatively, spatial locations of scRNA-seq cells can be reconstructed *de novo*, by optimal transport modeling, or projection into 2 or 3 dimensions corresponding to the spatial dimensions [121]. Optimal transport exploits spatial autocorrelation of gene expression to map cells to locations. However, spatial autocorrelation in tissues breaks down when different cell types are co-localized. When a spatial reference is not available, the *de novo* approach often does not yield results resembling the original tissue [121].

Gene expression in space can also be imputed from scRNA-seq without explicitly mapping scRNA-seq cells to locations. A common approach is to project the spatial and scRNA-seq data into a shared low dimensional and batch-free latent space, and to subsequently estimate gene expression by projecting the spatial cells into the latent space. Examples of this approach include Seurat3 [122] and gimVI [123]. These methods may also be used to add spatial context to single cell multi-omics data when spatial techniques for some of the multi-omics data are not available.

In spatial data without single cell resolution, such as those derived from ST and Visium, scRNA-seq data can inform cell type composition of the spots or voxels (Supplementary Material 7.4). A common approach is to explicitly model observed gene expression at the spots as a weighted sum of mean gene expression of each cell type from scRNA-seq. Gene counts can then be modeled with negative binomial (NB) or Poisson distributions, and cell type proportions in each spot can then be estimated from the parameters of the model. Examples of such methods include stereoscope [124], and RCTD [124].

Given the relevance of scRNA-seq to spatial data, popular scRNA-seq exploratory data analysis (EDA) ecosystems such as Seurat [122], SCANPY (Squidpy) [125], and SingleCellExperiment (SpatialExperiment) [126] have added functionalities for spatial data, such as updates to data containers and functions to facilitate visualization of gene expression and cell/spot metadata at spatial locations (Supplementary Material 7.2). EDA packages dedicated to spatial data with beautiful graphics and good documentation have also been written, such as Giotto [127], STUtility [128], and SPATA [129]. Seurat, Giotto, and SPATA also implement basic methods to identify spatially variable genes. In addition, Giotto implements methods to identify cell type enrichment in ST and Visium spots, identify gene coexpression and association between gene expression and cell type colocalization, and to identify spatial regions [130].

Spatially variable genes are genes whose expression is associated with spatial location (Supplementary Material 7.5). Two approaches are commonly used: Gaussian process regression (GPR) [131] and its generalization to Poisson [132] and NB [133], and Laplacian score [134]. The former models normalized gene expression or the rate parameter of Poisson or NB gene expression as a GPR and finds whether the model better describes the data with the spatial term than without. To speed up computation, the cells or spots can be aggregated with self-organizing map before the GPR approach is applied [135]. The latter approach identifies genes whose expression better reflects the structure of a spatial neighborhood graph. The locations of cells can also be modeled as a spatial point process with gene expression as marks; spatially variable genes can be identified as marks associated with locations [136]. GPR based methods approximate the p-values of genes from theory, while other methods use permutation testing, which makes them less scalable. However, the Gaussian kernel commonly used for GPR based methods doesn’t account for anisotropy observed in tissues.

Spatial information also enables identification of potential cell-cell interaction (Supplementary Material 7.8). This is commonly done with knowledge of ligand-receptor (L-R) pairs and testing of whether L-R pairs are more likely to be expressed in neighboring cells or spots [137]. Expression of genes of interest can also be modeled, including a term for cell-cell co-localization; the gene is considered associated with cell-cell co-localization if the model better describes the data with this term than without [138, 139].

There are many other types of analyses that are useful for spatial transcriptomics analysis, including identification of archetypal gene patterns (Supplementary Material 7.6), spatial regions defined by the transcriptome (Supplementary Material 7.7), inferring gene-gene interactions (Supplementary Material 7.9), subcellular transcript localization (Supplementary Material 7.10), and gene expression imputation from H&E (Supplementary Material 7.11).

## 4 Trends in the spatial transcriptomics field

The quality vs. quantity trade off inherent in existing technologies means that there is no single “best” solution currently available, and the difficulty in implementing methods has resulted in many technologies never spreading beyond their institutions of origin. LCM, ST, Visium, ISS, and Tomo-seq have been the most widely adopted (Figure 3A), and in almost all cases in the US and western Europe (Supplementary Figures 4.9, 5.25, 5.31). In terms of tissues analyzed, multiplexed current era techniques have been used widely to characterize human tissues [140], tumors [53] (especially breast tumors and squamous cell carcinoma), and pathological tissues that don’t necessarily have a stereotypical structure [141] (Figure 3B, C). In the SARS-CoV-2 pandemic, GeoMX DSP has been used for spatial transcriptomic profiling in lung autopsy of COVID victims [142, 143].

Some of the processed data, and associated spatially variable genes, can be downloaded and visualized from SpatialDB [144]. Excluding LCM, the vast majority of current era studies were performed on either humans or mice (Figure 3D), and the brain is the most studied organ (Figure 3B, E, F). In particular, the international project Brain Research through Advancing Innovative Neurotechnologies (BRAIN) Initiative - Cell Census Network (BICCN) is constructing a multi-modal atlas for the human, mouse, and non-human primate brain, including spatial data such as MERFISH and seqFISH [145]. While the smFISH techniques being utilized for this project can in principle scale to many genes, in practice they have for the most part been used for limited numbers of genes (Figure 3G) and cells (Figure 3H).

All packages mentioned in the Data analysis section are open source and written in languages such as R, Python, and Julia. Downstream analyses in studies primarily concerning new data and data analysis packages predominantly use open source programming languages such as R, Python, and C++ (Figure 3I, J). While MATLAB is still popular, its use appears to be declining with R and Python gaining in popularity (Supplementary Figure 5.9). While R is more popular for downstream analyses and EDA, Python and C++ are more popular for package development (Figure 3I, J), reflecting relative strengths of these languages and their surrounding cultures. Most of the packages are not hosted on standard repositories such as the Comprehensive R Archive Network (CRAN), Bioconductor, pip, and conda (Supplementary Figure 7.9). While most packages using R, Python, and C++ are well-documented, most MATLAB packages are not, perhaps reflecting the central role of documentation in open source culture (Supplementary Figure 7.9).

## 5 Future prospective

While technologies of the prequel are rapidly being deprecated, the ideas and methods that underlie them are fundamental to current era spatial transcriptomics. The field has dramatically expanded over the past 5 years (Figure 4A), with a plethora of new techniques and popularization of Visium driving growth (Figure 4B, Supplementary Figures 4.8, 5.34). The popularity of these techniques may be attributable to applicability to diverse tissues and the availability of commercial kits and core facilities translating to less work and cost to set up instruments and train personnel.

What lies ahead of the rising curves? First, more can be done to improve data collection techniques. For example, most current era techniques require tissue sections. Highly multiplexed whole mount smFISH and tissue clearing protocols, and more efficient computational tools for aligning multiple sections that may come from multiple individuals or even developmental stages, should be developed to extend current era techniques to 3D and to spatiotemporal analysis. Furthermore, smFISH and ISS techniques, with signal amplification to reduce the number of probes per transcript, can be adapted to target isoform specific exons or untranslated regions rather than all transcripts of a gene.

Second, current era data has not yet been integrated into comprehensive databases. Prequel databases such as GXD, e-Mouse Atlas and Gene Expression (EMAGE) [148], and FlyExpress [149] include data from multiple sources and can be queried by gene symbol and developmental and spatial ontologies. In addition, ABA [30], EMAGE, and FlyExpress aligned ISH images to common coordinates and can be queried with expression patterns. While some current era authors provide online interactive visualization of datasets from their studies [61], comprehensive databases integrating data from multiple sources as in the prequel era have not yet been developed. Furthermore, while prequel ontologies are still used in current era studies, such ontologies may be improved with the transcriptome wide quantitative data from the current era.

Third, outside of LCM, the current era is highly focused on the brain in human and mice, with potential spatial transcriptomics investigations of other organs such as the liver and the leaf lagging behind. Technological modernization of prequel consortia for organisms other than human and mice, and for organs other than the brain, holds much promise for the development of useful spatial transcriptomics atlases.

Fourth, an open source, well-documented, interoperable, and scalable workflow with an integrated easy-to-use interface would greatly simplify spatial transcriptomics data collection and analysis. At present, for tasks beyond EDA, users still often need to learn new syntax, convert object types, and even learn new languages to use some data analysis tools. Finally, our survey of methods shows that spatial transcriptomics methods need to be more open and accessible so that they become adopted around the world, and are not restricted to Western elite institutions.

## 6 Methods

### 6.1 Database

The criterion for including studies in the database was that the study used a method for quantification of transcripts while recording spatial context of samples within a tissue or cell. Methods were required to be able to quantify more transcripts and genes than possible with one round of FISH or immunofluorescence. Reviews and protocols were excluded. For publications on data analysis methods, the criterion for inclusion was that the proposed method went beyond the use of existing packages, and demonstrated a novel methodological approach or technique on a spatial transcriptomic dataset. In addition to collating publications through extensive reading and citation searching, keywords such as “spatial transcriptomics”, “digital spatial profiling”, “in situ sequencing”, as well as technology specific terms such as “visium”, “seqfish”, and “merfish”, were searched on PubMed and bioRxiv. The search results, as well as related papers and other papers citing the publication of interest on PubMed were manually screened and entered into the database. The sharing of the database is intended to foster crowd sourcing of literature in the future.

### 6.2 Analysis

All analyses were performed with R version 4.0.4. The timelines were generated with the R package `ggtext` [150]. The tissue maps in Figure 3B-C, E-F were generated with the R package `gganatogram` [151]. All trend lines were estimated by linear regression, and significance of the slope for each was computed by a t-test. The pictograms in Figure 3I-J were generated with the R package `ggtextures` [152].

In the supplement, the medium resolution world map was generated with the R package `rnaturalearth` using the Robinson projection option. Cities and institutions were geocoded with the Google geocoding API.

For LCM text mining, PubMed abstracts and metadata were downloaded by searching for the expression “((laser capture microdissection) OR (laser microdissection)) AND ((microarray) OR (transcriptome) OR (RNA-seq))” with the PubMed API, with R package `easyPubMed` [153], BioRxiv abstracts were downloaded by web scraping with Python package `biorxiv-retriever` [154] and metadata was downloaded with the bioRxiv API. The abstracts were tokenized into unigrams while preserving common phrases. Stopwords and punctuations were removed and the words were stemmed. With token counts in each abstract, the R package `stm` [155] was used for topic modeling of LCM abstracts from PubMed and bioRxiv. The date of publication, city of first authors (PubMed) or corresponding authors (bioRxiv), and journal (including bioRxiv) were used as covariates to model topic prevalence. Due to the large number of cities and journals, cities and journals with fewer than 5 publications were grouped into “Other” prior to modeling. The number of topics was chosen based on a trade off between hold out likelihood and residual, and between topic exclusivity and semantic coherence; 50 topics were used. Global vector (GloVe) embedding [156] of words from the abstracts was performed with the R package `text2vec` [157], using 125 dimensions. The word embeddings were then Louvain clustered [158] with the R package `igraph` [159] and projected to lower dimensions for visualization with principal component analysis (PCA) using the R function prcomp, and uniform manifold approximation and projection (UMAP) [160] with the R package `uwot` [161].

The paper and supplement can be continuously updated to reflect the contents of the database.

## 6.3 Code and data availability

All code for generating the figures is available at https://github.com/pachterlab/LP_2021, and the version as of submission is archived at https://doi.org/10.5281/zenodo.4795375. The curated database can be accessed at https://docs.google.com/spreadsheets/d/1sJDb9B7AtYmfKv4-m8XR7uc3XXw_k4kGSout8cqZ8bY/.

## References

1. Liao, J., Lu, X., Shao, X., Zhu, L. & Fan, X. Uncovering an Organ’s Molecular Architecture at Single-Cell Resolution by Spatially Resolved Transcriptomics. Trends in Biotechnology. ISSN: 01677799. https://linkinghub.elsevier.com/retrieve/pii/S0167779920301402 (June 2020).

2. Asp, M., Bergenstråhle, J. & Lundeberg, J. Spatially Resolved Transcriptomes—Next Generation Tools for Tissue Exploration. BioEssays 42, 1900221. ISSN: 0265-9247. https://doi.org/10.1002/bies.201900221 (Oct. 2020).

3. Smith, E. A. & Hodges, H. C. The Spatial and Genomic Hierarchy of Tumor Ecosystems Revealed by Single-Cell Technologies. Trends in Cancer 5, 411–425. ISSN: 24058033. https://linkinghub.elsevier.com/retrieve/pii/S2405803319301013 (July 2019).

4. Lein, E., Borm, L. E. & Linnarsson, S. The promise of spatial transcriptomics for neuroscience in the era of molecular cell typing 2017.

5. Saviano, A., Henderson, N. C. & Baumert, T. F. Single-cell genomics and spatial transcriptomics: Discovery of novel cell states and cellular interactions in liver physiology and disease biology. Journal of Hepatology 73, 1219–1230. ISSN: 01688278. https://linkinghub.elsevier.com/retrieve/pii/S016882782030372X (Nov. 2020).

6. Gall, J. G., Lou, M., Kline, P. & Giles, N. H. Formation and Detecction of RNA-DNA Hybrid Molecules in Cytological Preparations. PNAS 63, 378–383. https://www.pnas.org/content/63/2/378 (1969).

7. John, H. A., Birnstiel, M. L. & Jones, K. W. RNA-DNA hybrids at the cytological level. Nature 223, 582–587. ISSN: 00280836. https://www.nature.com/articles/223582a0 (1969).

8. Harrison, P., Conkie, D., Paul, J. & Jones, K. Localisation of cellular globin messenger RNA by in situ hybridisation to complementary DNA. FEBS Letters 32, 109–112. ISSN: 00145793. http://doi.wiley.com/10.1016/0014-5793%2873%2980749-5 (May 1973).

9. Langer-Safer, P. R., Levine, M. & Ward, D. C. Immunological method for mapping genes on Drosophila polytene chromosomes. Proceedings of the National Academy of Sciences 79, 4381–4385. ISSN: 0027-8424. https://www.pnas.org/content/79/14/4381 (July 1982).

10. Rudkin, G. & Stollar, B. D. High resolution detection of DNA–RNA hybrids in situ by indirect immunofluorescence. Nature 265, 472–473. ISSN: 1476-4687. https://doi.org/10.1038/265472a0 (1977).

11. Tautz, D. & Pfeifle, C. A non-radioactive in situ hybridization method for the localization of specific RNAs in Drosophila embryos reveals translational control of the segmentation gene hunchback. Chromosoma 98, 81–85. ISSN: 0009-5915. https://link.springer.com/article/10.1007/BF00291041 (Aug. 1989).

12. Rosen, B. & Beddington, R. S. Whole-mount in situ hybridization in the mouse embryo: gene expression in three dimensions. Trends in Genetics 9, 162–167. ISSN: 01689525. https://linkinghub.elsevier.com/retrieve/pii/016895259390162B (May 1993).

13. Singer, R. H. & Ward, D. C. Actin gene expression visualized in chicken muscle tissue culture by using in situ hybridization with a biotinated nucleotide analog. Proceedings of the National Academy of Sciences of the United States of America 79, 7331–7335. ISSN: 00278424. http://www.pnas.org/content/79/23/7331.abstract (Dec. 1982).

14. O’Kane, C. J. & Gehring, W. J. Detection in situ of genomic regulatory elements in Drosophila. Proceedings of the National Academy of Sciences of the United States of America 84, 9123–9127. ISSN: 00278424. https://www.pnas.org/content/84/24/9123 (Dec. 1987).

15. Gossler, A., Joyner, A., Rossant, J. & Skarnes, W. Mouse embryonic stem cells and reporter constructs to detect developmentally regulated genes. Science 244, 463–465. ISSN: 0036-8075. https://www.sciencemag.org/lookup/doi/10.1126/science.2497519 (Apr. 1989).

16. Hope, I. A. ‘Promoter trapping’ in Caenorhabditis elegans tech. rep. (1991), 399–408. https://dev.biologists.org/content/develop/113/2/399.full.pdf.

17. Seydoux, G. & Fire, A. Soma-germline asymmetry in the distributions of embryonic RNAs in Caenorhabditis elegans. Development 120, 2823 LP–2834. http://dev.biologists.org/content/120/10/2823.abstract (Oct. 1994).

18. Bettenhausen, B. & Gossler, A. Efficient Isolation of Novel Mouse Genes Differentially Expressed in Early Postimplantation Embryos. Genomics 28, 436–441. ISSN: 08887543. https://linkinghub.elsevier.com/retrieve/pii/S088875438571172X (Aug. 1995).

19. Gawantka, V. et al. Gene expression screening in Xenopus identifies molecular pathways, predicts gene function and provides a global view of embryonic patterning. Mechanisms of Development 77, 95–141. ISSN: 09254773. https://linkinghub.elsevier.com/retrieve/pii/S0925477398001154 (Sept. 1998).

20. Ringwald, M., Mangan, M. E., Eppig, J. T., Kadin, J. A. & Richardson, J. E. GXD: a Gene Expression Database for the laboratory mouse. Nucleic Acids Research 27, 106–112. ISSN: 0305-1048. https://doi.org/10.1093/nar/27.1.106 (Jan. 1999).

21. Kawashima, T. MAGEST: MAboya Gene Expression patterns and Sequence Tags. Nucleic Acids Research 28, 133–135. ISSN: 13624962. http://www.genome.ad.jp/magest/%20https://academic.oup.com/nar/article-lookup/doi/10.1093/nar/28.1.133 (Jan. 2000).

22. Maeda, I., Kohara, Y., Yamamoto, M. & Sugimoto, A. Large-scale analysis of gene function in Caenorhabditis elegans by high-throughput RNAi. Current Biology 11, 171–176. ISSN: 0960-9822. https://www.sciencedirect.com/science/article/pii/S0960982201000525 (2001).

23. Satou, Y. et al. Gene expression profiles in &lt;em&gt;Ciona intesti-nalis&lt;/em&gt; tailbud embryos. Development 128, 2893 LP–2904. http://dev.biologists.org/content/128/15/2893.abstract (Aug. 2001).

24. Carson, J. P., Thaller, C. & Eichele, G. A transcriptome atlas of the mouse brain at cellular resolution. Current Opinion in Neurobiology 12, 562–565. ISSN: 09594388. https://linkinghub.elsevier.com/retrieve/pii/S0959438802003562 (Oct. 2002).

25. Tomancak, P. et al. Systematic determination of patterns of gene expression during Drosophila embryogenesis. Genome biology 3, research0088.1. ISSN: 14656914. http://genomebiology.biomedcentral.com/articles/10.1186/gb-2002-3-12-research0088 (Dec. 2002).

26. Henrich, T. MEPD: a Medaka gene expression pattern database. Nucleic Acids Research 31, 72–74. ISSN: 13624962. https://academic.oup.com/nar/article-lookup/doi/10.1093/nar/gkg017 (Jan. 2003).

27. Sprague, J. The Zebrafish Information Network (ZFIN): the zebrafish model organism database. Nucleic Acids Research 31, 241–243. ISSN: 13624962. http://zfin.org/zf_info/%20https://academic.oup.com/nar/article-lookup/doi/10.1093/nar/gkg027 (Jan. 2003).

28. Bell, G. W., Yatskievych, T. A. & Antin, P. B. GEISHA, a whole-mount in situ hybridization gene expression screen in chicken embryos. Developmental Dynamics 229, 677–687. ISSN: 1058-8388. http://doi.wiley.com/10.1002/dvdy.10503 (Mar. 2004).

29. Wienholds, E. MicroRNA Expression in Zebrafish Embryonic Development. Science 309, 310–311. ISSN: 0036-8075. https://www.sciencemag.org/lookup/doi/10.1126/science.1114519 (July 2005).

30. Lein, E. S. et al. Genome-wide atlas of gene expression in the adult mouse brain. Nature 445, 168–176. ISSN: 0028-0836. http://www.nature.com/articles/nature05453 (Jan. 2007).

31. Luengo Hendriks, C. L. et al. Three-dimensional morphology and gene expression in the Drosophila blastoderm at cellular resolution I: Data acquisition pipeline. Genome Biology 7, R123. ISSN: 14747596. http://genomebiology.biomedcentral.com/articles/10.1186/gb-2006-7-12-r123 (Dec. 2006).

32. Lécuyer, E. et al. Global Analysis of mRNA Localization Reveals a Prominent Role in Organizing Cellular Architecture and Function. Cell 131, 174–187. ISSN: 00928674. https://linkinghub.elsevier.com/retrieve/pii/S0092867407010227 (Oct. 2007).

33. Bowes, J. B. et al. Xenbase: a Xenopus biology and genomics resource. Nucleic Acids Research 36, D761–D767. ISSN: 0305-1048. https://doi.org/10.1093/nar/gkm826 (Jan. 2008).

34. Harding, S. D. et al. The GUDMAP database - an online resource for genitourinary research. Development 138, 2845–2853. ISSN: 0950-1991. http://golgi.ana.%20http://dev.biologists.org/cgi/doi/10.1242/dev.063594 (July 2011).

35. Ardini-Poleske, M. E. et al. LungMAP: The Molecular Atlas of Lung Development Program. American Journal of Physiology-Lung Cellular and Molecular Physiology 313, L733–L740. ISSN: 1040-0605. https://www.physiology.org/doi/10.1152/ajplung.00139.2017 (Nov. 2017).

36. Lovell, P. V. et al. ZEBrA: Zebra finch Expression Brain Atlas—A resource for comparative molecular neuroanatomy and brain evolution studies. Journal of Comparative Neurology 528, 2099–2131. ISSN: 0021-9967. https://onlinelibrary.wiley.com/doi/abs/10.1002/cne.24879 (Aug. 2020).

37. Meier-Ruge, W. et al. The laser in the Lowry technique for microdissection of freeze-dried tissue slices. The Histochemical Journal 8, 387–401. ISSN: 0018-2214. http://link.springer.com/10.1007/BF01003828 (July 1976).

38. Landegren, U., Kaiser, R., Sanders, J. & Hood, L. A ligase-mediated gene detection technique. Science 241, 1077 LP–1080. http://science.sciencemag.org/content/241/4869/1077.abstract (Aug. 1988).

39. Belyavsky, A., Vinogradova, T. & Rajewsky, K. PCR-based cDNA library construction: general cDNA libraries at the level of a few cells. eng. Nucleic acids research 17, 2919–2932. ISSN: 0305-1048. https://pubmed.ncbi.nlm.nih.gov/2471144%20https://www.ncbi.nlm.nih.gov/pmc/articles/PMC317702/ (Apr. 1989).

40. Van Gelder, R. N. et al. Amplified RNA synthesized from limited quantities of heterogeneous cDNA. Proceedings of the National Academy of Sciences 87, 1663 LP–1667. http://www.pnas.org/content/87/5/1663.abstract (Mar. 1990).

41. Nederlof, P. M. et al. Multiple fluorescence in situ hybridization. Cytometry 11, 126–131. ISSN: 0196-4763. http://doi.wiley.com/10.1002/cyto.990110115 (1990).

42. Schena, M., Shalon, D., Davis, R. W. & Brown, P. O. Quantitative Monitoring of Gene Expression Patterns with a Complementary DNA Microarray. Science 270, 467 LP–470. http://science.sciencemag.org/content/270/5235/467.abstract (Oct. 1995).

43. Emmert-Buck, M. R. et al. Laser Capture Microdissection. Science 274, 998–1001. ISSN: 0036-8075. https://www.sciencemag.org/lookup/doi/10.1126/science.274.5289.998 (Nov. 1996).

44. Becker, I. et al. Single-cell mutation analysis of tumors from stained histologic slides. Laboratory Investigation 75, 801–807. ISSN: 00236837. https://pubmed.ncbi.nlm.nih.gov/8973475/ (Dec. 1996).

45. Femino, A. M., Fay, F. S., Fogarty, K. & Singer, R. H. Visualization of Single RNA Transcripts in Situ. Science 280, 585 LP–590. http://science.sciencemag.org/content/280/5363/585.abstract (Apr. 1998).

46. Luo, L. et al. Gene expression profiles of laser-captured adjacent neuronal subtypes. Nature Medicine 5, 117–122. ISSN: 1078-8956. http://www.nature.com/articles/nm0199_117 (Jan. 1999).

47. Levsky, J. M. Single-Cell Gene Expression Profiling. Science 297, 836–840. ISSN: 00368075. https://www.sciencemag.org/lookup/doi/10.1126/science.1072241 (Aug. 2002).

48. Lister, R. et al. Highly Integrated Single-Base Resolution Maps of the Epigenome in Arabidopsis. Cell 133, 523–536. ISSN: 0092-8674. https://www.sciencedirect.com/science/article/pii/S0092867408004480 (2008).

49. Okamura-Oho, Y. et al. Transcriptome Tomography for Brain Analysis in the Web-Accessible Anatomical Space. PLoS ONE 7 (ed Hayasaka, S.) e45373. ISSN: 1932-6203. https://dx.plos.org/10.1371/journal.pone.0045373 (Sept. 2012).

50. Ke, R. et al. In situ sequencing for RNA analysis in preserved tissue and cells. Nature Methods 10, 857–860. ISSN: 15487091. https://www.nature.com/articles/nmeth.2563 (Sept. 2013).

51. Lubeck, E., Coskun, A. F., Zhiyentayev, T., Ahmad, M. & Cai, L. Single-cell in situ RNA profiling by sequential hybridization Mar. 2014. https://www.nature.com/articles/nmeth.2892.

52. Chen, K. H., Boettiger, A. N., Moffitt, J. R., Wang, S. & Zhuang, X. Spatially resolved, highly multiplexed RNA profiling in single cells. Science. ISSN: 10959203 (2015).

53. Ståhl, P. L. et al. Visualization and analysis of gene expression in tissue sections by spatial transcriptomics. Science 353, 78–82. ISSN: 00368075. https://www.sciencemag.org/lookup/doi/10.1126/science.aaf2403 (July 2016).

54. Giani, A. M., Gallo, G. R., Gianfranceschi, L. & Formenti, G. Long walk to genomics: History and current approaches to genome sequencing and assembly. Computational and Structural Biotechnology Journal 18, 9–19. ISSN: 20010370. https://linkinghub.elsevier.com/retrieve/pii/S2001037019303277 (Jan. 2020).

55. Jenett, A. et al. A GAL4-Driver Line Resource for Drosophila Neurobiology. Cell Reports 2, 991–1001. ISSN: 22111247. http://dx.doi.org/10.1016/j.celrep.2012.09.011 (Oct. 2012).

56. Gong, S. et al. A gene expression atlas of the central nervous system based on bacterial artificial chromosomes. Nature 425, 917–925. ISSN: 0028-0836. http://www.nature.com/articles/nature02033 (Oct. 2003).

57. Ringwald, M. et al. A database for mouse development. S’cience 265, 2033–2034. ISSN: 00368075. https://science.sciencemag.org/content/265/5181/2033.abstract (Sept. 1994).

58. Howe, D. G. et al. The Zebrafish Model Organism Database: New support for human disease models, mutation details, gene expression phenotypes and searching. Nucleic Acids Research 45, D758–D768. ISSN: 13624962. https://academic.oup.com/nar/article/45/D1/D758/2605740 (Jan. 2017).

59. Satija, R., Farrell, J. A., Gennert, D., Schier, A. F. & Regev, A. Spatial reconstruction of single-cell gene expression data. Nature Biotechnology 33, 495–502. ISSN: 15461696. https://doi.org/10.1038/nbt.3192 (May 2015).

60. Karaiskos, N. et al. The Drosophila embryo at single-cell transcriptome resolution. Science 358, 194–199. ISSN: 10959203. https://science.sciencemag.org/content/358/6360/194 (Oct. 2017).

61. Ortiz, C. et al. Molecular atlas of the adult mouse brain. Science Advances 6, eabb3446. ISSN: 2375-2548. www.brain-map.org%20https://advances.sciencemag.org/lookup/doi/10.1126/sciadv.abb3446 (June 2020).

62. Morton, M. L. et al. Identification of mRNAs and lincRNAs associated with lung cancer progression using next-generation RNA sequencing from laser micro-dissected archival FFPE tissue specimens. Lung Cancer 85, 31–39. ISSN: 01695002. https://linkinghub.elsevier.com/retrieve/pii/S016950021400141X (July 2014).

63. Nichterwitz, S. et al. Laser capture microscopy coupled with Smart-seq2 for precise spatial transcriptomic profiling. Nature Communications 7, 12139. ISSN: 2041-1723. http://www.nature.com/articles/ncomms12139 (Nov. 2016).

64. Foley, J. W. et al. Gene expression profiling of single cells from archival tissue with laser-capture microdissection and Smart-3SEQ. Genome Research 29, 1816–1825. ISSN: 1088-9051. http://genome.cshlp.org/lookup/doi/10.1101/gr.234807.118 (Nov. 2019).

65. Peng, G. et al. Spatial Transcriptome for the Molecular Annotation of Lineage Fates and Cell Identity in Mid-gastrula Mouse Embryo. Developmental Cell 36, 681–697. ISSN: 18781551. http://dx.doi.org/10.1016/j.devcel.2016.02.020 (Mar. 2016).

66. Kerman, I. A., Buck, B. J., Evans, S. J., Akil, H. & Watson, S. J. Combining laser capture microdissection with quantitative real-time PCR: Effects of tissue manipulation on RNA quality and gene expression. Journal of Neuroscience Methods 153, 71–85. ISSN: 01650270. https://linkinghub.elsevier.com/retrieve/pii/S016502700500364X (May 2006).

67. Zechel, S., Zajac, P., Lönnerberg, P., Ibáñez, C. F. & Linnarsson, S. Topographical transcriptome mapping of the mouse medial ganglionic eminence by spatially resolved RNA-seq. Genome biology 15, 486. ISSN: 1474760X. http://genomebiology.biomedcentral.com/articles/10.1186/s13059-014-0486-z (Oct. 2014).

68. Brown, V. M. et al. Multiplex Three-Dimensional Brain Gene Expression Mapping in a Mouse Model of Parkinson’s Disease. Genome Research 12, 868–884. ISSN: 1088-9051. http://www.genome.org/cgi/doi/10.1101/gr.229002 (May 2002).

69. Junker, J. P. et al. Genome-wide RNA Tomography in the Zebrafish Embryo. Cell. ISSN: 10974172 (2014).

70. Merritt, C. R. et al. High multiplex, digital spatial profiling of proteins and RNA in fixed tissue using genomic detection methods. bioRxiv 38, 559021. ISSN: 1546-1696. https://www.biorxiv.org/content/10.1101/559021v2 (Feb. 2019).

71. Medaglia, C. et al. Spatial reconstruction of immune niches by combining photoactivatable reporters and scRNA-seq. Science 358, 1622–1626. ISSN: 0036-8075. https://www.sciencemag.org/lookup/doi/10.1126/science.aao4277 (Dec. 2017).

72. Lubeck, E. & Cai, L. Single-cell systems biology by super-resolution imaging and combinatorial labeling. Nature Methods 2012 9:7 9, 743–748. ISSN: 1548-7105. https://www.nature.com/articles/nmeth.2069 (June 2012).

73. Shah, S., Lubeck, E., Zhou, W. & Cai, L. In Situ Transcription Profiling of Single Cells Reveals Spatial Organization of Cells in the Mouse Hippocampus. Neuron. ISSN: 10974199 (2016).

74. Eng, C.-H. L., Shah, S., Thomassie, J. & Cai, L. Profiling the transcriptome with RNA SPOTs. Nature Methods 14, 1153–1155. ISSN: 1548-7105. https://doi.org/10.1038/nmeth.4500 (2017).

75. Eng, C. H. L. et al. Transcriptome-scale super-resolved imaging in tissues by RNA seqFISH÷. Nature 568, 235–239. ISSN: 14764687. https://www.nature.com/articles/s41586-019-1049-y (Apr. 2019).

76. Xia, C., Fan, J., Emanuel, G., Hao, J. & Zhuang, X. Spatial transcriptome profiling by MERFISH reveals subcellular RNA compartmentalization and cell cycle-dependent gene expression. Proceedings of the National Academy of Sciences of the United States of America. ISSN: 10916490 (2019).

77. Gyllborg, D. et al. Hybridization-based In Situ Sequencing (HybISS): spatial transcriptomic detection in human and mouse brain tissue. bioRxiv, 2020.02.03.931618. https://doi.org/10.1101/2020.02.03.931618 (Feb. 2020).

78. Goh, J. J. L. et al. Highly specific multiplexed RNA imaging in tissues with split-FISH. Nature Methods 17, 689–693. ISSN: 15487105. https://doi.org/10.1038/s41592-020-0858-0 (July 2020).

79. Moffitt, J. R. & Zhuang, X. in Methods in Enzymology (2016).

80. Samacoits, A. et al. A computational framework to study sub-cellular RNA localization. Nature Communications 9, 4584. ISSN: 2041-1723. http://www.nature.com/articles/s41467-018-06868-w (Dec. 2018).

81. Cabili, M. N. et al. Localization and abundance analysis of human lncRNAs at single-cell and single-molecule resolution. Genome Biology 16, 20. ISSN: 1474-760X. https://genomebiology.biomedcentral.com/articles/10.1186/s13059-015-0586-4 (Dec. 2015).

82. Battich, N., Stoeger, T. & Pelkmans, L. Image-based transcriptomics in thousands of single human cells at single-molecule resolution. Nature Methods 10, 1127–1136. ISSN: 15487091. https://www.nature.com/articles/nmeth.2657 (Nov. 2013).

83. Wang, X. et al. Three-dimensional intact-tissue sequencing of single-cell transcriptional states. Science 361, eaat5691. ISSN: 0036-8075. https://www.sciencemag.org/lookup/doi/10.1126/science.aat5691 (July 2018).

84. Chen, F., Tillberg, P. W. & Boyden, E. S. Expansion microscopy. Science 347, 543–548. ISSN: 0036-8075. https://www.sciencemag.org/lookup/doi/10.1126/science.1260088 (Jan. 2015).

85. Coskun, A. F. & Cai, L. Dense transcript profiling in single cells by image correlation decoding. Nature Methods 13, 657–660. ISSN: 15487105. https://www.nature.com/articles/nmeth.3895 (July 2016).

86. Sountoulidis, A. et al. SCRINSHOT, a spatial method for single-cell resolution mapping of cell states in tissue sections. bioRxiv, 2020.02.07.938571. https://www.biorxiv.org/content/biorxiv/early/2020/02/07/2020.02.07.938571.full.pdf (Feb. 2020).

87. Shendure, J. Accurate Multiplex Polony Sequencing of an Evolved Bacterial Genome. Science 309, 1728–1732. ISSN: 0036-8075. https://www.sciencemag.org/lookup/doi/10.1126/science.1117389 (Sept. 2005).

88. Lee, J. H. et al. Highly Multiplexed Subcellular RNA Sequencing in Situ. Science 343, 1360 LP–1363. http://science.sciencemag.org/content/343/6177/1360.abstract (Mar. 2014).

89. Alon, S. et al. Expansion Sequencing: Spatially Precise &lt;em&gt;In Situ&lt;/em&gt; Transcriptomics in Intact Biological Systems. bioRxiv, 2020.05.13.094268. http://biorxiv.org/content/early/2020/05/15/2020.05.13.094268.abstract (Jan. 2020).

90. Wang, G., Moffitt, J. R. & Zhuang, X. Multiplexed imaging of high-density libraries of RNAs with MERFISH and expansion microscopy. Scientific Reports. ISSN: 20452322 (2018).

91. Sun, Y.-C. et al. Integrating barcoded neuroanatomy with spatial transcriptional profiling reveals cadherin correlates of projections shared across the cortex. bioRxiv, 2020.08.25.266460. http://biorxiv.org/content/early/2020/08/26/2020.08.25.266460.abstract (Jan. 2020).

92. Partel, G. et al. Identification of spatial compartments in tissue from in situ sequencing data. bioRxiv, 765842. https://doi.org/10.1101/765842 (Sept. 2019).

93. Qian, X. et al. Probabilistic cell typing enables fine mapping of closely related cell types in situ. Nature Methods 17, 101–106. ISSN: 15487105. https://doi.org/10.1038/s41592-019-0631-4 (Jan. 2020).

94. Macosko, E. Z. et al. Highly parallel genome-wide expression profiling of individual cells using nanoliter droplets. Cell 161, 1202–1214. ISSN: 10974172. http://dx.doi.org/10.1016/j.cell.2015.05.002 (May 2015).

95. Zheng, G. X. Y. et al. Massively parallel digital transcriptional profiling of single cells. Nature Communications 8, 14049. ISSN: 2041-1723. http://www.nature.com/articles/ncomms14049 (Apr. 2017).

96. Klein, A. M. et al. Droplet Barcoding for Single-Cell Transcriptomics Applied to Embryonic Stem Cells. Cell 161, 1187–1201. ISSN: 00928674. https://linkinghub.elsevier.com/retrieve/pii/S0092867415005000 (May 2015).

97. Hashimshony, T. et al. CEL-Seq2: sensitive highly-multiplexed single-cell RNA-Seq. Genome Biology 17, 77. ISSN: 1474-760X. http://genomebiology.biomedcentral.com/articles/10.1186/s13059-016-0938-8 (Dec. 2016).

98. Grun, D., Kester, L. & van Oudenaarden, A. Validation of noise models for single-cell transcriptomics. Nature Methods 11, 637–640. ISSN: 15487091. https://www.nature.com/articles/nmeth.2930 (June 2014).

99. Lee, J. H. et al. Fluorescent in situ sequencing (FISSEQ) of RNA for gene expression profiling in intact cells and tissues. Nature Protocols 10, 442–458. ISSN: 1754-2189. http://www.nature.com/articles/nprot.2014.191 (Mar. 2015).

100. Rodriques, S. G. et al. Slide-seq: A scalable technology for measuring genome-wide expression at high spatial resolution. Science. ISSN: 10959203 (2019).

101. Vickovic, S. et al. High-definition spatial transcriptomics for in situ tissue profiling. Nature Methods. ISSN: 15487105 (2019).

102. Liu, Y. et al. High-Spatial-Resolution Multi-Omics Sequencing via Deterministic Barcoding in Tissue. Cell 183, 1665–1681. ISSN: 0092-8674. https://www.sciencedirect.com/science/article/pii/S0092867420313908 (2020).

103. Villacampa, E. G. et al. Genome-wide Spatial Expression Profiling in FFPE Tissues. bioRxiv, 2020.07.24.219758. https://doi.org/10.1101/2020.07.24.219758 (July 2020).

104. Lebrigand, K. et al. The spatial landscape of gene expression isoforms in tissue sections. bioRxiv, 2020.08.24.252296. https://doi.org/10.1101/2020.08.24.252296 (Aug. 2020).

105. Chen, A. et al. Large field of view-spatially resolved transcriptomics at nanoscale resolution. bioRxiv, 2021.01.17.427004. http://biorxiv.org/content/early/2021/01/24/2021.01.17.427004.abstract (Jan. 2021).

106. Cho, C.-S. et al. Seq-Scope: Submicrometer-resolution spatial transcriptomics for single cell and subcellular studies. bioRxiv, 2021.01.25.427807. http://biorxiv.org/content/early/2021/01/27/2021.01.25.427807.abstract (Jan. 2021).

107. Fu, X. et al. Continuous Polony Gels for Tissue Mapping with High Resolution and RNA Capture Efficiency. bioRxiv, 2021.03.17.435795. http://biorxiv.org/content/early/2021/03/17/2021.03.17.435795.abstract (Jan. 2021).

108. Stickels, R. et al. Sensitive spatial genome wide expression profiling at cellular resolution. bioRxiv, 2020.03.12.989806. https://doi.org/10.1101/2020.03.12.989806 (Mar. 2020).

109. Weinstein, J. A., Regev, A. & Zhang, F. DNA Microscopy: Optics-free Spatio-genetic Imaging by a Stand-Alone Chemical Reaction. Cell 178, 229–241. ISSN: 10974172. https://doi.org/10.1016/j.cell.2019.05.019 (June 2019).

110. Halpern, K. B. et al. Paired-cell sequencing enables spatial gene expression mapping of liver endothelial cells. Nature Biotechnology 36, 962. ISSN: 15461696. https://www.nature.com/articles/nbt.4231 (Nov. 2018).

111. Manco, R. et al. Clump sequencing exposes the spatial expression programs of intestinal secretory cells. bioRxiv, 2020.08.05.237917. https://www.biorxiv.org/content/10.1101/2020.08.05.237917v1 (Aug. 2020).

112. Fazal, F. M. et al. Atlas of Subcellular RNA Localization Revealed by APEX-Seq. Cell 178, 473–490. ISSN: 00928674. https://linkinghub.elsevier.com/retrieve/pii/S0092867419305550 (July 2019).

113. Vickovic, S. et al. SM-Omics: An automated platform for high-throughput spatial multi-omics. bioRxiv, 2020.10.14.338418. http://biorxiv.org/content/early/2020/10/15/2020.10.14.338418.abstract (Jan. 2020).

114. Su, J.-H., Zheng, P., Kinrot, S. S., Bintu, B. & Zhuang, X. GenomeScale Imaging of the 3D Organization and Transcriptional Activity of Chromatin. Cell 182, 1641–1659. ISSN: 00928674. https://linkinghub.elsevier.com/retrieve/pii/S0092867420309405 (Sept. 2020).

115. Shah, S. et al. Dynamics and Spatial Genomics of the Nascent Transcriptome by Intron seqFISH. Cell. ISSN: 10974172 (2018).

116. Zhang, M. et al. Molecular, spatial and projection diversity of neurons in primary motor cortex revealed by in situ single-cell transcriptomics. bioRxiv, 2020.06.04.105700. http://biorxiv.org/content/early/2020/06/05/2020.06.04.105700.abstract (Jan. 2020).

117. Van Valen, D. A. et al. Deep Learning Automates the Quantitative Analysis of Individual Cells in Live-Cell Imaging Experiments. PLOS Computational Biology 12, e1005177. https://doi.org/10.1371/journal.pcbi.1005177 (Nov. 2016).

118. Petukhov, V., Soldatov, R. A., Khodosevich, K. & Kharchenko, P. V. Bayesian segmentation of spatially resolved transcriptomics data. bioRxiv, 2020.10.05.326777. http://biorxiv.org/content/early/2020/10/06/2020.10.05.326777.abstract (Jan. 2020).

119. Axelrod, S. et al. {Starfish}: Open Source Image Based Transcriptomics and Proteomics Tools http://github.com/spacetx/starfish.

120. Nitzan, M., Karaiskos, N., Friedman, N. & Rajewsky, N. Gene expression cartography. Nature 576, 132–137. ISSN: 14764687. https://doi.org/10.1038/s41586-019-1773-3 (Dec. 2019).

121. Zhu, J. & Sabatti, C. Integrative Spatial Single-cell Analysis with Graphbased Feature Learning. bioRx?v, 2020.08.12.248971. http://biorxiv.org/content/early/2020/08/13/2020.08.12.248971.abstract (Jan. 2020).

122. Stuart, T. et al. Comprehensive Integration of Single-Cell Data. Cell 177, 1888–1902. ISSN: 00928674. https://linkinghub.elsevier.com/retrieve/pii/S0092867419305598 (June 2019).

123. Lopez, R. et al. A joint model of unpaired data from scRNA-seq and spatial transcriptomics for imputing missing gene expression measurements. http://arxiv.org/abs/1905.02269 (May 2019).

124. Andersson, A. et al. Spatial mapping of cell types by integration of transcriptomics data. bioRxiv, 2019.12.13.874495. https://www.biorxiv.org/content/10.1101/2019.12.13.874495v1.full#ref-20 (Dec. 2019).

125. Palla, G. et al. Squidpy: a scalable framework for spatial single cell analysis. bioRxiv, 2021.02.19.431994. http://biorxiv.org/content/early/2021/02/20/2021.02.19.431994.abstract (Jan. 2021).

126. Righelli, D. et al. SpatialExperiment: infrastructure for spatially resolved transcriptomics data in R using Bioconductor. bioRxiv, 2021.01.27.428431. http://biorxiv.org/content/early/2021/01/27/2021.01.27.428431.abstract (Jan. 2021).

127. Dries, R. et al. Giotto, a pipeline for integrative analysis and visualization of single-cell spatial transcriptomic data. bioRxiv, 701680. https://www.biorxiv.org/content/10.1101/701680v1.full (May 2019).

128. Bergenstråhle, J., Larsson, L. & Lundeberg, J. Seamless integration of image and molecular analysis for spatial transcriptomics workflows. BMC Genomics 21, 482. ISSN: 1471-2164. https://bmcgenomics.biomedcentral.com/articles/10.1186/s12864-020-06832-3 (Dec. 2020).

129. Kueckelhaus, J. et al. Inferring spatially transient gene expression pattern from spatial transcriptomic studies. bioRxiv, 2020.10.20.346544. http://biorxiv.org/content/early/2020/10/21/2020.10.20.346544.abstract (Jan. 2020).

130. Zhu, Q., Shah, S., Dries, R., Cai, L. & Yuan, G. C. Identification of spatially associated subpopulations by combining scRNAseq and sequential fluorescence in situ hybridization data. Nature Biotechnology 36, 1183–1190. ISSN: 15461696. https://www.nature.com/articles/nbt.4260 (Dec. 2018).

131. Svensson, V., Teichmann, S. A. & Stegle, O. SpatialDE: Identification of spatially variable genes. Nature Methods 15, 343–346. ISSN: 15487105. https://www.nature.com/articles/nmeth.4636 (Apr. 2018).

132. Sun, S., Zhu, J. & Zhou, X. Statistical analysis of spatial expression patterns for spatially resolved transcriptomic studies. Nature Methods 17, 193–200. ISSN: 15487105. https://doi.org/10.1038/s41592-019-0701-7 (Feb. 2020).

133. BinTayyash, N. et al. Non-parametric modelling of temporal and spatial counts data from RNA-seq experiments. bioRxiv, 2020.07.29.227207. http://biorxiv.org/content/early/2020/07/30/2020.07.29.227207.abstract (Jan. 2020).

134. Govek, K. W., Yamajala, V. S. & Camara, P. G. Clustering-independent analysis of genomic data using spectral simplicial theory. PLOS Computational Biology 15, e1007509. https://doi.org/10.1371/journal.pcbi.1007509 (Nov. 2019).

135. Hao, M., Hua, K. & Zhang, X. SOMDE: A scalable method for identifying spatially variable genes with self-organizing map. bioRxiv, 2020.12.10.419549. http://biorxiv.org/content/early/2021/03/24/2020.12.10.419549.abstract (Jan. 2021).

136. Edsgärd, D., Johnsson, P. & Sandberg, R. Identification of spatial expression trends in single-cell gene expression data. Nature Methods 15, 339–342. ISSN: 1548-7091. http://www.nature.com/articles/nmeth.4634 (May 2018).

137. Pham, D. et al. stLearn: integrating spatial location, tissue morphology and gene expression to find cell types, cell-cell interactions and spatial trajectories within undissociated tissues. bioRxiv, 2020.05.31.125658. http://biorxiv.org/content/early/2020/05/31/2020.05.31.125658.abstract (Jan. 2020).

138. Arnol, D., Schapiro, D., Bodenmiller, B., Saez-Rodriguez, J. & Stegle, O. Modeling Cell-Cell Interactions from Spatial Molecular Data with Spatial Variance Component Analysis. Cell Reports 29, 202–211. ISSN: 22111247. https://doi.org/10.1016/j.celrep.2019.08.077 (Oct. 2019).

139. Tanevski, J. et al. Gene selection for optimal prediction of cell position in tissues from single-cell transcriptomics data. Life Science Alliance 3, e202000867. http://www.life-science-alliance.org/content/3/11/e202000867.abstract (Nov. 2020).

140. Maynard, K. R. et al. Transcriptome-scale spatial gene expression in the human dorsolateral prefrontal cortex. bioRxiv, 788992. https://doi.org/10.1101/2020.02.28.969931 (Oct. 2019).

141. Lundmark, A. et al. Gene expression profiling of periodontitis-affected gingival tissue by spatial transcriptomics. Scientific Reports 8, 9370. ISSN: 2045-2322. https://doi.org/10.1038/s41598-018-27627-3 (2018).

142. Butler, D. et al. Shotgun transcriptome, spatial omics, and isothermal profiling of SARS-CoV-2 infection reveals unique host responses, viral diversification, and drug interactions. Nature Communications 12, 1660. ISSN: 2041-1723. https://doi.org/10.1038/s41467-021-21361-7 (2021).

143. Margaroli, C. et al. Spatial mapping of SARS-CoV-2 and H1N1 Lung Injury Identifies Differential Transcriptional Signatures. eng. Cell reports. Medicine, 100242. ISSN: 2666-3791. https://pubmed.ncbi.nlm.nih.gov/33778787%20https://www.ncbi.nlm.nih.gov/pmc/articles/PMC7985929/ (Mar. 2021).

144. Fan, Z., Chen, R. & Chen, X. SpatialDB: a database for spatially resolved transcriptomes. Nucleic Acids Research 48, D233–D237. ISSN: 0305-1048. https://doi.org/10.1093/nar/gkz934 (Jan. 2020).

145. Adkins, R. S. et al. A multimodal cell census and atlas of the mammalian primary motor cortex. bioRxiv, 2020.10.19.343129. http://biorxiv.org/content/early/2020/10/21/2020.10.19.343129.abstract (Jan. 2020).

146. Burgess, D. J. Spatial transcriptomics coming of age. Nature Reviews Genetics 20, 317–317. ISSN: 1471-0056. https://www.nature.com/articles/s41576-019-0129-z (June 2019).

147. Brady, L. et al. Inter- and intra-tumor heterogeneity of metastatic prostate cancer determined by digital spatial gene expression profiling. Nature Communications 12, 1426. ISSN: 2041-1723. https://doi.org/10.1038/s41467-021-21615-4 (2021).

148. Armit, C. et al. eMouseAtlas: An atlas-based resource for understanding mammalian embryogenesis. Developmental Biology 423, 1–11. ISSN: 00121606. https://linkinghub.elsevier.com/retrieve/pii/S0012160616308582 (Mar. 2017).

149. Kumar, S. et al. FlyExpress 7: An Integrated Discovery Platform To Study Coexpressed Genes Using in situ Hybridization Images in Drosophila. G3; Genes—Genomes—Genetics 7, 2791–2797. ISSN: 2160-1836. http://g3journal.org/lookup/doi/10.1534/g3.117.040345 (Aug. 2017).

150. Wilke, C. O. ggtext: Improved Text Rendering Support for ‘ggplot2’ 2020. https://cran.r-project.org/package=ggtext.

151. Maag, J. L. V. gganatogram: An R package for modular visualisation of anatograms and tissues based on ggplot2 [version 1; peer review: 2 approved]. F1000Research 7 (2018).

152. Wilke, C. ggtextures: Drawing Textured Rectangles and Bars with grid and ggplot2 2020.

153. Fantini, D. easyPubMed: Search and Retrieve Scientific Publication Records from PubMed 2019. https://cran.r-project.org/package=easyPubMed.

154. Schuster, T. BiorxivRetriever 2020. https://github.com/TalSchuster/BiorxivRetriever%20https://pypi.org/project/biorxiv-retriever/.

155. Roberts, M. E., Stewart, B. M. & Tingley, D. Stm: An R package for structural topic models. Journal of Statistical Software 91. ISSN: 15487660. https://www.jstatsoft.org/v091/i02 (2019).

156. Pennington, J., Socher, R. & Manning, C. D. GloVe: Global Vectors for Word Representation tech. rep. ().

157. Selivanov, D., Bickel, M. & Wang, Q. text2vec: Modern Text Mining Framework for R 2020. https://cran.r-project.org/package=text2vec.

158. Blondel, V. D., Guillaume, J.-L., Lambiotte, R. & Lefebvre, E. Fast unfolding of communities in large networks. Journal of Statistical Mechanics: Theory and Experiment 2008, P10008. ISSN: 1742-5468. http://dx.doi.org/10.1088/1742-5468/2008/10/P10008 (2008).

159. Csardi, G. & Nepusz, T. The igraph software package for complex network research. InterJournal Complex Sy, 1695. https://igraph.org (2006).

160. McInnes, L., Healy, J. & Melville, J. UMAP: Uniform Manifold Approximation and Projection for Dimension Reduction 2020.

161. Melville, J. uwot: The Uniform Manifold Approximation and Projection (UMAP) Method for Dimensionality Reduction 2020. https://cran.r-project.org/package=uwot.

