## Supplementary Material for "Museum of Spatial Transcriptomics"

2021-05-25

[1] mailto:
[2] mailto:

### Contents

**Preface** 5

**1 Introduction** 7

#### I Prequel era 11

**2 Prequel era** 13

#### II Current era 43

**4 From the past to the present** 45

**5 Current era technologies 57**

**III Future perspectives 177**

**8 From the past to the present to the future 179**

**References 183**

### Preface

This supplement to the paper Museum of Spatial Transcriptomics[1] and the associated database of spatial transcriptomics literature[2] is inspired by museum catalogs that provide insight and detail to further understanding of the exhibits. The results presented are based on code that can be run interactively on RStudio Cloud[3]. We present key analyses of metadata curated for the database, and provide further analyses and results beyond what could be included here in the `more_analyses` directory of this repository. The markdown that generates this text is on GitHub, and is version controlled so that its development can be tracked now and in the future. Please notify us of errors, omissions, or other suggestions via submission of issues on GitHub: `https://github.com/pachterlab/LP_2021`

This document is built with the `bookdown` package from a collection of R Markdown files. How some of figures look depends on parameters that can be changed, such as size of bins when binning number of publications in time to show a trend. The source code is on RStudio Cloud[4]. The dependencies are pre-installed in the RStudio Cloud project. By default, when the database is queried by code, the most up to date version is used, which can be newer than the rendered static version on github.io. To build the document in RStudio Cloud (i.e., both the web page `gitbook` version and a PDF), run this in the R console:

```
bookdown::render_book("index.Rmd", output_format = c("bookdown::gitbook", "bookdown::pdf_book"))
```

If you are cloning this repo into a fresh RStudio Cloud project or a fresh machine, install the packages required to build the book as follows:

First install `remotes` with `install.packages("remotes")`. Then use

[1] `https://www.biorxiv.org/content/10.1101/2021.05.11.443152v1`

[2] `https://docs.google.com/spreadsheets/d/1sJDb9B7AtYmfKv4-m8XR7uc3XXw_k4kGSout8cqZ8bY/edit?usp=sharing`

[3] `https://rstudio.cloud/project/2492054`

[4] `https://rstudio.cloud/project/2492054`

`remotes:install_deps(dependencies = TRUE)` to install all required packages from CRAN, Bioconductor, and GitHub. So in short,

```
install.packages("remotes")
remotes::install_deps(dependencies = TRUE)
```

Because many packages are installed, the installation can be sped up with the argument `Ncpus` in `install_deps()` to specify the number of CPU cores to use to install packages in parallel, such as `Ncpus = 2L` for 2 cores. The free plan of RStudio Cloud only has 1 core, but this argument can be used when multiple cores are available.

By default, the most up to date version of the database is downloaded for analyses in this book. However, as the `museumst` R package written for these analyses contains a cached version of the database, historical versions of the database can be viewed by installing older versions of `museumst` and setting `update = FALSE` when calling `museumst::read_metadata()` when running code from this book on RStudio Cloud or your computer. Older versions of `museumst` can be installed with

```
remotes::install_github("pachterlab/museumst", ref = "v0.0.0.9016")
```

where `ref` refers to a release. Release history of `museumst` can be seen here[5]. Documentation of `museumst` can be seen here[6].

[5] `https://github.com/pachterlab/museumst/releases`
[6] `https://pachterlab.github.io/museumst/`

### Chapter 1

### Introduction

The spatial organization of the components of biological systems is crucial for their proper function. For instance, morphogen gradients in embryos are tightly regulated to ensure that the right cell types differentiate at the right place. In adults, spatial organization of cells in tissues is important to proper functions of organs. For instance, the liver lobule is divided in labor according to distance from the portal triad as such distance affects suitability of different tasks. Both oxygen level and morphogen gradient regulate zonation of metabolism (Gebhardt 2014); there is more oxidative phosphorylation and gluconeogenesis in the more oxygenated periportal region and more glycolysis in the more deoxygenated pericentral region. How cell types and cellular functions vary in space is measured by quantifying gene expression in space. Conversely, the expression of an unknown gene in space can give clues to its function. Gene expression is usually quantified by quantifying proteins or transcripts encoded by the gene, and high throughput spatial methods exist for both protein and transcripts. In other words, cellular function exemplifies the maxim that “the whole is greater than the sum of its parts”, and in large part this follows from “location, location, location”.

Here we focus on spatial transcriptomics (the field of spatial proteomics is covered elsewhere (Lundberg and Borner 2019; Baharlou et al. 2019; Buchberger et al. 2018)). Even spatial transcriptomics is a vast field, and it is useful to begin by considering the scope of what it contains. Naïvely, one may say, spatial transcriptomics means quantifying the complete set of RNAs encoded by the genome in space. Usually the “in space” is at some microscopic resolution rather than geospatial as often assumed in the term “spatial statistics”; the resolution is usually cellular, though sometimes subcellular. The “spatial” is in contrast to other transcriptomics methods that by virtue of the nature of their assays, lose information of tissue structure in space. That is the case with microarray technology for bulk tissue analysis, for bulk RNA-seq, and single cell RNA-seq (scRNA-seq) that is based on dissociation of tissue – the “spatial” usually means tissue structure in space. More broadly, the "spatial" can mean knowing spatial context of samples although the spatial context is only a label and the coordinates are not collected or not used, such as in some laser capture microdissection (LCM) literature (Baccin et al. 2020; Nichterwitz et al. 2016), Niche-seq (Medaglia et al. 2017), and APEX-seq (Fazal et al. 2019). The "spatial" can also mean preserving spatial coordinates of samples within tissue, though the coordinates may or may not be explicitly used in data analysis, such as in the various single molecular fluorescent in situ hybridization (smFISH) based technologies such as seqFISH (Lubeck et al. 2014) and MERFISH (K. H. Chen et al. 2015) and array based technologies such as Spatial Transciptomics (ST) (Ståhl et al. 2016).

There is more complexity in defining "transcriptomics". While some technologies usually called "spatial transcriptomics" are indeed transcriptome-wide, such as ST, Visium, and LCM followed by RNA-seq, many technologies that only profile a panel of usually a few hundred genes are nevertheless considered part of "spatial transcriptomics". Here "transcriptomics" actually means high-throughput quantification of gene expression, preferably highly multiplexed, quantifying numerous genes within the same piece of tissue at the same time. However, what counts as "high-throughput"? Is there a minimum number of genes required? Should 50 genes be enough? Or a hundred genes? The threshold number of genes required to be considered "high-throughput" is difficult to define; here, by "high-throughput", we mean the intent to quantify expression of more genes than normally done with fluorescent in situ hybridization (FISH) or immunofluorescence when only color distinguishes between genes, which can mean more than about 5 genes. There is also some complication regarding whether "highly multiplexed" should be required. Some fairly recent studies that intended to perform high-throughput gene expression profiling in space did not profile most genes at the same time (e.g. multiple rounds of smFISH hybridization, each round for a different set of genes) (Lignell et al. 2017), or even profiled different genes in different tissue sections (Bayraktar et al. 2020; Battich, Stoeger, and Pelkmans 2013); these papers nevertheless claimed to be spatial transcriptomic or something similar.

When terms are to be defined by how they are used, then we rely on a generic and inclusive definition of "spatial transcriptomics", which can be summarized as: Quantifying transcripts while keeping spatial context of samples within tissue or cell, with intent to quantify transcripts of more genes than normally done with one round of FISH or immunofluorescence when color is the only way to distinguish between genes. This is the criterion we used in considering what methods to include in our review.

The field of spatial transcriptomics has grown drastically in the past 5 years, during which several reviews have already been written. These survey existing technologies (Crosetto, Bienko, and Van Oudenaarden 2015; Moor and Itzkovitz 2017; Strell et al. 2019; Liao et al. 2020; Waylen et al. 2020) or discuss how the technologies apply to specific biological systems such as tumors (Smith et al.

2019), brain (Lein, Borm, and Linnarsson 2017), and liver (Saviano, Henderson, and Baumert 2020). Unlike the review papers, we aim to be more systematic and detailed in our review of spatial transcriptomics technology. In addition, we review existing data analysis methods in this field, a crucial aspect of spatial transcriptomics which has not yet been reviewed in depth. Moreover, we present a curated database of spatial transcriptomics literature and analyses of the literature metadata to show trends in different aspects of spatial transcriptomics. This database is publicly available here[1]. A similar database has been curated for scRNA-seq literature, which has been analyzed to show trends in the field (Svensson, da Veiga Beltrame, and Pachter 2020) although the metadata in our database and the analyses are much more extensive.

There are some caveats to our review and database. First, while we narrate a history of evolution of techniques and in some cases explain how one technique influenced another, we do not present aspects of the history that are not apparent from the publications. Studying those aspects of the history of the field may require interviewing the people who developed the techniques, as well as exploration of additional unpublished material. Second, our database was originally only meant for papers, so relevant materials that are not in presented in that format are underrepresented. Examples of such materials include databases and software not presented as papers (e.g. the XDB3 database ("XDB3" 2004) and the `spicyR` R package (Canete and Patrick 2020)). This means that the metadata analyses in this book might not be representative of all material that exists in spatial transcriptomics. Third, as the curation was done manually, the database might not include some relevant literature unknown to us.

Our database starts with articles published in the 1980s. This provides historical context of what is now commonly known as spatial transcriptomics; this literature is summarized in Chapter 2, and historical methods of data analysis are reviewed in Chapter 3. The literature is broken down into the following categories, to be defined and elaborated on in the subsequent chapters. Technologies to collect data (Chapter 4): Microdissection (Section 5.1), array based techniques (Section 5.4), single molecular FISH (smFISH) (Section 5.2), in situ sequencing (ISS) (Section 5.3), and no imaging (Section 5.5). Data analysis methods (Chapter 7): Preprocessing (Section 7.1), exploratory data analysis (EDA) (Section 7.2), spatial reconstruction of single cell RNA-seq (scRNA-seq) data (Section 7.3), spatially variable genes (Section 7.5), archetypal gene expression patterns (Section 7.6), using transcriptome to identify spatially coherent regions in tissue (Section 7.7), cell type deconvolution of non-single-cell resolution spatial data (Section 7.4), cell-cell interaction (Section 7.8), and other types of analyses. Also, the literature metadata is analyzed to show relevant sociological trends such as who is using each technology, usage trends of technologies, and the programming languages used. The metadata analyses can be run interactively in RStudio Cloud[2].

[1] https://docs.google.com/spreadsheets/d/1sJDb9B7AtYmfKv4-m8XR7uc3XXw_k4kGSout8cqZ8bY/edit#gid=1693202466

[2] https://rstudio.cloud/project/1981124

### Part I

### Prequel era

### Chapter 2

### Prequel era

Some previous reviews on spatial transcriptomics start the history of spatial transcriptomics with laser capture microdissection (LCM) followed by microarray or RNA-seq and single molecular fluorescent *in situ* hybridization (smFISH) in the late 1990s (Lein, Borm, and Linnarsson 2017; Liao et al. 2020; Crosetto, Bienko, and Van Oudenaarden 2015). We will discuss these later, but note that by 1999 and the early 2000s, when the earliest LCM microarray studies were published (Luo et al. 1999; Sgroi et al. 1999; Ohyama et al. 2000; Kitahara et al. 2001), the quest to profile the transcriptome in space had already begun, with enhancer and gene trap screens, *in situ* reporter screens, and (whole mount) *in situ* hybridization ((WM)ISH) atlases. Although this early literature, dating from the late 1980s, generally does not refer to itself as "spatial transcriptomics", it fits into the definition of spatial transcriptomics as stated in Chapter 1.

We call this body of literature "prequel", because first, its origin predates LCM microarray. Second, unlike most technologies covered by existing spatial transcriptomics reviews, the techniques used were not multiplexed and were less quantitative, and as a result, they have fallen out of favor. In contrast, what comes after "prequel" will be called "current", although the prequel and current eras chronologically overlap. Given what current era spatial transcriptomics is commonly perceived to be, here "prequel" is broadly defined as methods that fulfill the more relaxed definition of "spatial transcriptomics" in this book, but do not involve cDNA microarray, next generation sequencing (NGS), or single molecular imaging.

There are 204 prequel papers in our database. Prequel literature is included in the database and covered here for the following reasons. First, the legacy of the prequel era has influenced more recent spatial transcriptomic research; the present and future are shaped by the past. For example, spatial reconstruction of scRNA-seq data in Seurat v1 (Satija et al. 2015), the Achim et al. *Platynereis* study (Achim et al. 2015), `DistMap` (Karaiskos et al. 2017), and the Zeisel et al. Mouse Brain Atlas (Zeisel et al. 2018) used (WM)ISH atlases as spatial references. Recent Spatial Transcriptomics$^{TM}$ (ST) mouse brain data are still compared to the ISH atlas of Allen Brain Atlas (ABA) (Ortiz et al. 2020; Chen et al. 2020). A study on spatial reconstruction of scATAC-seq data compared the *in silico* reconstruction to the FlyLight *Drosophila* enhancer atlas (Jenett et al. 2012; Bravo González-Blas et al. 2020). Hence prequel resources can still be useful in the current era. Second, some features of the prequel era may benefit future spatial transcriptomics studies; this will be discussed after more recent technologies are reviewed. Third, the various quests in the current era have already begun in the prequel era, and this history can show where we are in these quests.

Fourth, as shown later in this book, existing current era spatial transcriptomics data are by and large from humans and mice, and especially the brain (Figure 4.4, Figure 2.7). For other model and non-model organisms (e.g. *Xenopus laevis* (Bowes et al. 2009; "XDB3" 2004), *Ciona intestinalis* (Satou et al. 2001), *Danio rerio* (Sprague 2003; Belmamoune and Verbeek 2008), *Oryzias latipes* (Henrich 2003), *Gallus gallus* (Bell, Yatskievych, and Antin 2004), *Taeniopygia guttata* (Lovell et al. 2020), and to some extent, even *Drosophila melanogaster* (Tomancak et al. 2002; Luengo Hendriks et al. 2006)), some tissues other than the brain (e.g. lung (Ardini-Poleske et al. 2017), retina (Blackshaw et al. 2004), genitourinary tract (Harding et al. 2011)), and miRNAs (Ahmed et al. 2015; Karali et al. 2010; Diez-Roux et al. 2011; Aboobaker et al. 2005; Wienholds 2005; Darnell et al. 2006), the most comprehensive spatial transcriptomic resources, if any are are available at all, are still (WM)ISH atlases. For plants, the most comprehensive resources can still be enhancer and gene trap screens (Johnson et al. 2005; Nakayama et al. 2005). Hence, while current era technologies may produce more quantitative and highly multiplexed data, they have not completely superseded (WM)ISH atlases. Finally, the historical literature is curated for the same reason why museums and libraries keep historical maps and scientific works that have been superseded by more recent work; it is part of our heritage.

An overall timeline for prequel techniques is shown in Figure 2.1, which will be discussed in more details in the rest of this chapter.

#### 2.1 Enhancer and gene traps

Long before the advent of reference genomes for common model organisms, the quest to characterize genes based on expression pattern in space had already begun. The earliest high-throughput efforts to identify and characterize such genes were enhancer traps. To the best of our knowledge, the first use of a reporter to visualize gene expression in space was reported in 1983. It used lacZ fused to sequences upstream to the hsp70 gene encoding a heat shock protein in *Drosophila melanogaster* and inserted into the genome with P element to char-

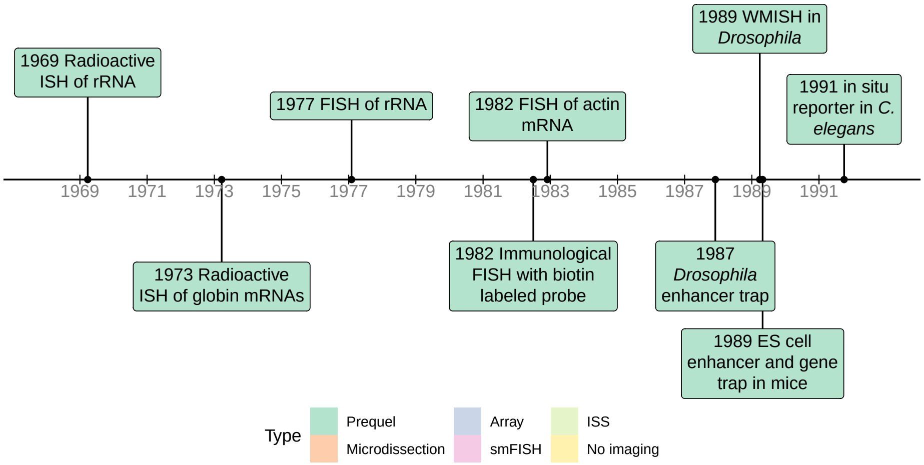

**Figure 2.1:** Timeline of prequel techniques.

acterize the puffs formed in polytene chromosomes and the tissue distribution of hsp70 in response to heat shock (Lis, Simon, and Sutton 1983).

The first enhancer trap screen in *Drosophila melanogaster* was published in 1987 (O'Kane and Gehring 1987). The P element is a transposable element found in *Drosophila*. In an enhancer trap vector, a reporter gene, such as lacZ, here with the polyadenylation site of the hsp70 gene, and a marker gene with its own promoter that can be used to identify individuals and their offspring with the vector integrated into the germline, such as rosy which can be used in *Drosophila* to identify the individuals with eye color, are flanked by the 5' and 3' ends of the P element necessary for transposition (Figure 2.2). The vector is injected into *Drosophila* embryos before the formation of pole cells (Spradling and Rubin 1982). As a transposon, the construct is randomly inserted into the genome, and since the P element promoter is so weak that an enhancer is required for the promoter to drive transcription of the reporter gene, the location of the reporter gene expression marks where the enhancer is active. As the transposon is inserted into different locations of the genome in different individuals, each individual that has the vector integrated into the germline forms a transformant line. In *Drosophila*, in many cases, expression patterns of $\beta$-galactosidase do reflect expression pattern of a nearby gene (Bellen et al. 1989; Wilson et al. 1989).

Since then, different vectors have been developed for better efficiency and flexibility (Stanford, Cohn, and Cordes 2001), and enhancer traps have been applied at increasing scale. The 1987 study recovered 39 lines (O'Kane and Gehring 1987), possibly characterizing 39 genes, but already in 1989, over 3000 lines were possible in one study (Bier et al. 1989). Enhancer trapping was also adapted to other species, such as mouse (Gossler et al. 1989; Allen et al. 1988) and *Arabidopsis thaliana* (Sundaresan et al. 1995).

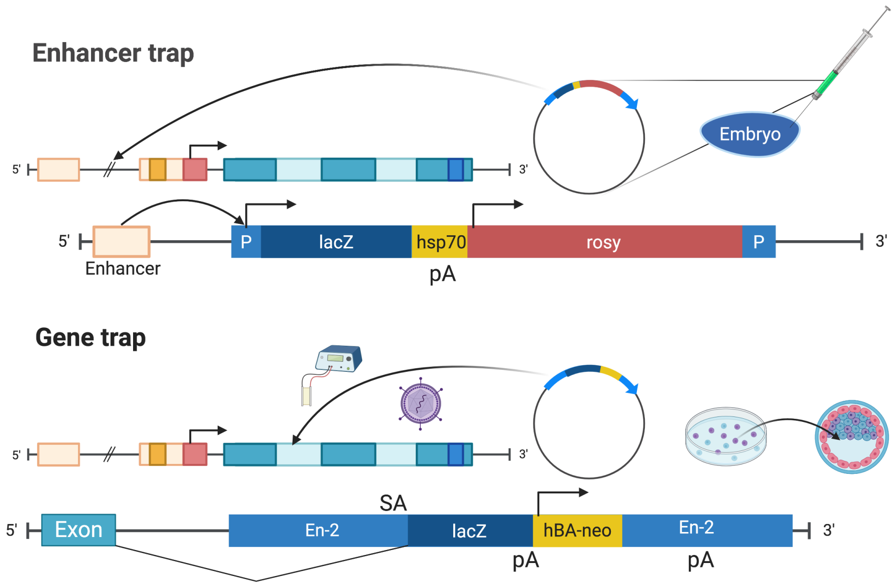

**Figure 2.2:** Illustrations of enhancer trap as described in (O'Kane and Gehring 1987) and gene trap as described in (Gossler et al. 1989) (Created with BioRender.com).

Enhancer traps were not intended to be mutagenic (O'Kane and Gehring 1987), nor is it highly mutagenic (Stanford, Cohn, and Cordes 2001). Gene trap and promoter traps were introduced to not only screen for genes with restricted expression patterns, but also to enable functional analysis of the gene from homozygote mutant phenotypes (Friedrich and Soriano 1991). Like the typical enhancer trap vector, gene trap and promoter trap vectors contain a reporter gene, such as lacZ ($\beta$-gal), to visualize gene expression, and sometimes also a marker to screen for integration, such as the neomycin resistance gene (neo). Though often, lacZ itself, or in a fusion with neo ($\beta$-geo), was also used as the marker when screening mouse embryonic stem (ES) cells (Figure 2.2).

Unlike the enhancer trap vector, gene trap and promoter trap vectors do not have a promoter for the reporter, though the marker, if present, can have its own promoter. In a promoter trap, the construct needs to be inserted in frame and in the correct orientation into an exon of a gene to be expressed, making it very inefficient (Friedrich and Soriano 1991; Stanford, Cohn, and Cordes 2001).

In contrast, in gene traps, a splice acceptance site is added to the 5' end of the reporter, so the construct can be expressed when inserted into an intron at the right orientation; this is over 50 times more efficient than a promoter trap because introns tend to be much longer than exons and the construct does not have to be in frame to an exon (Friedrich and Soriano 1991; Stanford, Cohn, and Cordes 2001). Gene traps and promoter traps are mutagenic as the reporter has a stop codon, thus truncating the endogenous protein.

While enhancer traps are more commonly used in *Drosophila*, gene traps are more commonly used in mice. In mice, in 1988, the enhancer trap vector was initially introduced by injection into the male pronucleus in the fertilized egg (Allen et al. 1988). The throughput of the screen is increased by inserting the construct into genomes of ES cells by electroporation or retroviral infection (Stanford, Cohn, and Cordes 2001), screening for ES cells expressing lacZ or the marker, injecting these ES cells into blastocysts to generate chimeric mice to characterize gene expression patterns; chimera are especially useful for characterizing dominant and lethal mutations (Friedrich and Soriano 1991; Gossler et al. 1989).

The first gene trap screen in mouse ES cells was reported in 1989 (Gossler et al. 1989), recovering 14 lines. Again, variants of the vector emerged and gene trap screens increased in scale. In 1995, nearly 300 mouse gene trap lines were recovered from one study (Wurst et al. 1995). Later, smaller gene trap studies specific to particular types of genes made possible by additional steps to screen ES cell colonies were performed, such as genes encoding membrane and secreted proteins (Skarnes et al. 1995), genes responding to retinoic acid (Forrester et al. 1996), and genes expressed in hematopoitic and endothelial lineages (Stanford et al. 1998). In 2001, gene trapping was used to examine not only expression pattern of genes in cell bodies of neurons in the mouse brain, but also axon guidance (Leighton et al. 2001). By 2001, a number of gene trap consortia have been established as resources of gene trap vectors and transformant mouse ES cell lines, hoping to create at least one line for each gene in the mouse genome (Stanford, Cohn, and Cordes 2001).

In the 1980s and 1990s, with increasing throughput of Sanger sequencing and the advent of shotgun sequencing, the amount of sequencing data in GenBank exploded (Giani et al. 2020). With 5' or 3' rapid amplification of cDNA ends (RACE) PCR, the fusion transcript of the reporter and an endogenous gene could be cloned (Frohman, Dush, and Martin 1988), sequenced, and potentially aligned to the existing sequences to identify the gene of interest (Stanford et al. 1998). However, the golden age of gene trapping was soon to pass, with the rise of ISH atlases in the late 1990s and the advent of reference genomes of *Drosophila melanogaster* (Myers 2000), mouse (Waterston et al. 2002), and human (Lander et al. 2001; Venter et al. 2001) in the early 2000s that would make it easier to design ISH probes from the reference genome to target annotated genes, as is done today. Nevertheless, enhancer and gene traps were not rendered obsolete by these developments. They have been used in plants and zebrafish through the 2000s and 2010s, as resources of gene expression patterns (Johnson et al. 2005; Nakayama et al. 2005; Pérez-Martín et al. 2017; Hiwatashi et al. 2001; Kawakami et al. 2010; Marquart et al. 2015) (Figure 2.3).

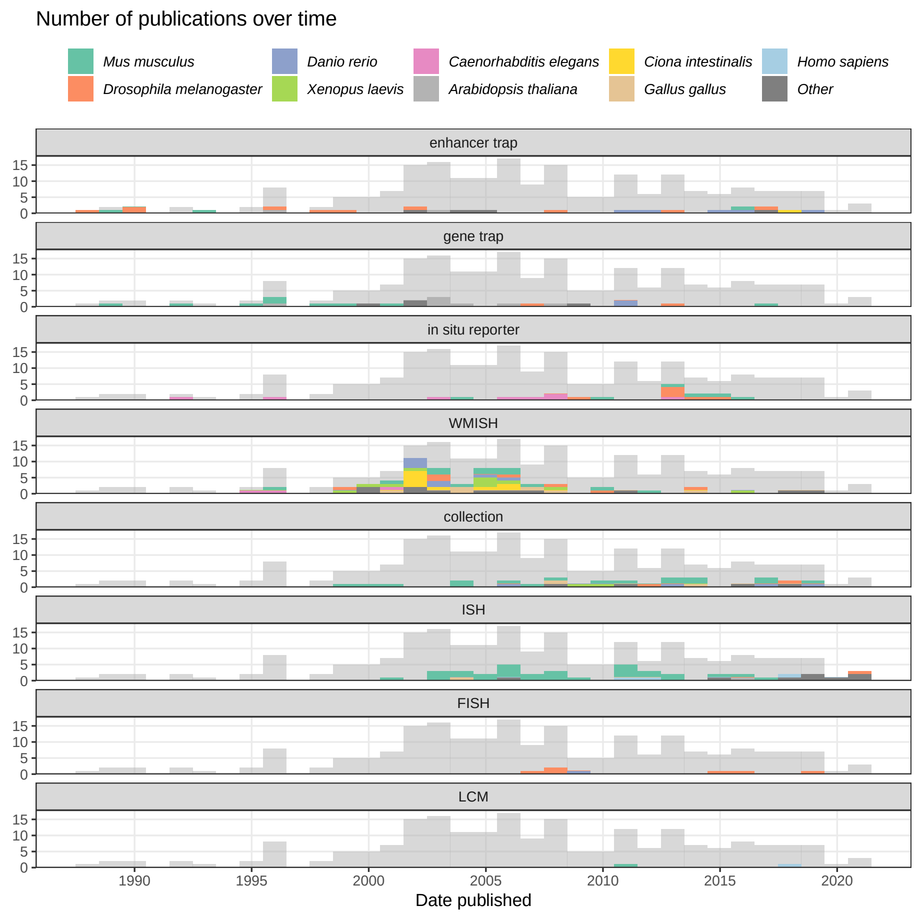

**Figure 2.3:** Number of publications over time in the prequel era, broken down by technique and colored by species. The gray histogram in the background is the histogram for all prequel publications over time. The bin width of this histogram is 365 days. Here WMISH and ISH exclude fluorescent ISH (FISH).

#### 2.2 In situ reporter

In enhancer, gene, or promoter trap screens, the reporter is randomly inserted into the genome, not targeting predetermined genes. In contrast, in what we call *in situ* reporter screens, the reporter is fused to predefined regulatory sequences of a gene of interest, with the hope that expression pattern of the reporter would recapitulate that of the gene of interest. Chronologically, this is the second type of high throughput method to profile gene expression patterns (Figure 2.3).

A precursor to this type of method was used in 1991, where random genomic fragments were fused to a lacZ reporter lacking a transcription start signal and injected as plasmids, screening for fragments driving lacZ expression and characterizing the expression patterns in *C. elegans* (Hope 1991). To the best of our knowledge, the first time *in situ* reporter with predefined regulatory sequences was used to screen for gene expression patterns in a multicellular organism, was in 1995, in *C. elegans* (Lynch, Briggs, and Hope 1995). At that time, the *C. elegans* genome sequencing project was already in progress (Lynch, Briggs, and Hope 1995; Sulston et al. 1992), and the genome sequence was declared "essentially complete" in 1998 (The C. elegans Sequencing Consortium 1998). Computationally predicted upstream regulatory sequences of 35 putative genes were fused to a promoterless lacZ as a reporter, cloned into plasmid vectors, and microinjected into *C. elegans* gonads to create transformed lines then stained with X-gal (Lynch, Briggs, and Hope 1995).

A reliable *in situ* reporter was first reported in mice in 1997. It used a recombinant bacterial artificial chromosome (BAC) with part of the full RU49 gene in the BAC replaced by a lacZ construct and showed that the construct is heritable (Yang, Model, and Heintz 1997). In 2003, a similar strategy, replacing coding sequences of genes in BACs with EGFP reporter gene, was used to create a mouse brain gene expression atlas GENSAT[1] with BAC transgenic mouse lines (Gong et al. 2003). The GENSAT lines were used again in 2009 to create a gene expression atlas for retina (Siegert et al. 2009). Again, GENSAT benefited from the reference genome, which greatly helped with identifying BACs that include sequences flanking a gene that may contain regulatory elements that make the reporter better recapitulate expression pattern of the endogenous gene (Gong et al. 2003).

Through the 2000s and 2010s, *in situ* reporters have been used as a targeted alternative to enhancer and gene trap screens informed by the reference genomes. To address limitations of gene traps, such as inability to precisely define the allele and favoring genes expressed in ES cells when screening for transformant colonies, high-throughput mouse knock out resources with knock out alleles computationally designed according to a reference genome and annotations have been established (Skarnes et al. 2011; "A Mouse for All Reasons" 2007). As these alleles contain a lacZ reporter, these resources have been used to characterize gene expression in over 40 tissues in mutant mice with lacZ staining (White et al. 2013; West et al. 2015; Tuck et al. 2015). However, for some tissues, only low resolution whole mount staining was performed. Similarly, in both mouse (Visel et al. 2013) and *Drosophila* (Jenett et al. 2012; Kvon et al. 2014), transgenic lines with genomic fragments containing putative enhancers driving expression of reporter genes were established as alternatives to enhancer traps. The enhancer candidates can be selected from sequence homology and ChIP-seq predictions (Visel et al. 2013), or from tiles of sequences flanking genes thought to have restricted expression patterns or within introns of such genes (Jenett et al. 2012).

*In situ* reporter atlases exceeded the scale of enhancer and gene trap screens.

[1]`http://www.gensat.org/index.html`

The largest such atlas in *C. elegans*, WormAtlas, profiled 1886 genes (Hunt-Newbury et al. 2007); we are unaware of enhancer and gene trap screens in *C. elegans* because *C. elegans* genome sequencing was already underway by 1992 (Sulston et al. 1992), making *in situ* reporter screening feasible before it was so in mice and *Drosophila*. The largest such study in *Drosophila* profiled 7705 enhancer candidates (Kvon et al. 2014), which far exceeded the 3768 enhancer trap lines in 1989 (Bier et al. 1989). *In situ* reporters were used in mice to profile up to 536 genes(Siegert et al. 2009) and 329 enhancer candidates (Visel et al. 2013), while the large scale gene trap screen in 1995 only reached 279 lines (Wurst et al. 1995) and later mouse gene trap screens did not typically exceed 100 lines. However, where comparable, *in situ* reporter atlases never reached the scale of (WM)ISH atlases, perhaps because of the large number of transgenic lines required. Allen Brain Atlas (ABA[2]) profiled over 20,000 genes in the mouse brain, and as of April 2021, the Berkeley Drosophila Genome Project (BDGP[3]) WMISH atlas already has 8533 genes. However, *in situ* reporters might still be a good way to profile enhancer usage in space.

#### 2.3 ISH and WMISH atlases

*In situ* hybridization was first used in 1969 to visualize ribosomal RNA (rRNA) (Gall et al. 1969) and ribosomal DNA (rDNA) (John, Birnstiel, and Jones 1969) in *Xenopus laevis* oocytes with probes labeled with radioisotope $^3$H (Figure 2.1). To the best of our knowledge, the earliest use of ISH to visualize what was thought to be a specific transcript was done in 1973, to visualize globin mRNAs in various cultured erythroid and non-erythoid cell types by hybridization of radiolabeled cDNA to the mRNA (Harrison et al. 1973). As radioactive ISH requires long exposure time (several weeks), has low spatial resolution and high background, and requires handling hazardous radioactive material, alternatives emerged in the mid 1970s and early 1980s. Among the alternatives were variants of FISH and labeled probes detected by primary and enzyme or fluorophore labeled secondary antibodies (Huber, Voith von Voithenberg, and Kaigala 2018; Langer-Safer, Levine, and Ward 1982); the latter, immunological method is commonly used in ISH and WMISH atlases. To the best of our knowledge, the first report of using immunological fluorescent and peroxidase ISH to visualize expression of a specific gene was published in 1982, the same year such technique was published (Langer-Safer, Levine, and Ward 1982), visualizing actin transcripts in chicken muscle tissue culture; the authors reported puncta of cytoplasmic fluorescence which might be clumps of mRNAs or artefact, but could possibly be individual transcripts (Singer and Ward 1982).

Non-radioactive ISH not only has shorter exposure time and higher resolution than radioactive ISH, but also made WMISH possible. WMISH was first reported in Drosophila embryos in 1989 (Tautz and Pfeifle 1989), and was adapted

[2] `https://portal.brain-map.org`

[3] `https://insitu.fruitfly.org/cgi-bin/ex/insitu.pl`

to other model organisms such as mice, *Xenopus laevis*, and *Paracentrotus lividus* (purple sea urchin) in the early 1990s (Rosen and Beddington 1993). Advantages of WMISH compared to section ISH is preservation of 3D structure of the tissue, ease of interpretation in blastoderm stage embryos, and ease of performing ISH on larger number of embryos (Rosen and Beddington 1993; Tautz and Pfeifle 1989).

Just like genome sequencing in multi-cellular organisms and *in situ* reporter screens, WMISH atlases got a head start in *C. elegans*. The first WMISH screen with higher throughput than typically used on select marker genes was reported in 1994, of 21 genes in *C. elegans* (Seydoux and Fire 1994). Early (WM)ISH atlases in the late 1990s typically made probes from cDNA clones from poly-A selected RNAs in tissue or developmental stage of interest without pre-selecting genes to stain for (Tomancak et al. 2002; Stapleton 2002; Gawantka et al. 1998; Bettenhausen and Gossler 1995). Some early atlases were intended to be improvements to enhancer and gene trapping and *in situ* reporter screens, as a simpler and more direct alternative (Bettenhausen and Gossler 1995) or as a way that can better capture endogenous and dynamic spatial distribution of transcripts (Gawantka et al. 1998). Since 1998, (WM)ISH has been automated, enabling staining for thousands of probes (Gawantka et al. 1998; Carson, Thaller, and Eichele 2002).

The genes from which the clones come from were often unknown, so early (WM)ISH atlases referred to the entities stained for as “clones” (Figure 2.4), though the genes, homology, and putative functions of the genes can be identified by aligning sequences of the cDNA clones to existing sequences in databases (Bettenhausen and Gossler 1995; Gawantka et al. 1998; Kopczynski et al. 1998). However, again, the first WMISH screen with probes made from cloning PCR amplified pre-defined genomic sequences was performed in *C. elegans* in 1995 (Birchall, Fishpool, and Albertson 1995). By the turn of the century, the entities stained for were sometimes referred to as “clusters”, especially in the GHOST atlas for *Ciona intestinalis* (Satou et al. 2001) (Figure 2.4); the sequences of the probes were clustered by alignment and these probes might have come from the same gene.

The rise of (WM)ISH atlases started before the completion of genome projects in humans and common model organisms, although their later growth was transformed by the reference genome. In the 2000s, with the availability of sequenced cDNA collections covering increasing proportion of predicted genes and the consequent rise of transcriptome-wide microarray (Stapleton 2002; Carter 2003), genes to be stained for in (WM)ISH atlases could be pre-screened based on microarray data of the tissue of interest, with probes made from cDNA clones readily available from such collections (Yoshikawa et al. 2006; Lein 2004). In addition, probes could be computationally designed based on reference genome sequences (Lein et al. 2007). Perhaps because of these developments, since the turn of the century, entities stained for have been predominantly referred to as “genes” (Figure 2.4). Notably, while radioactive ISH has been mostly replaced

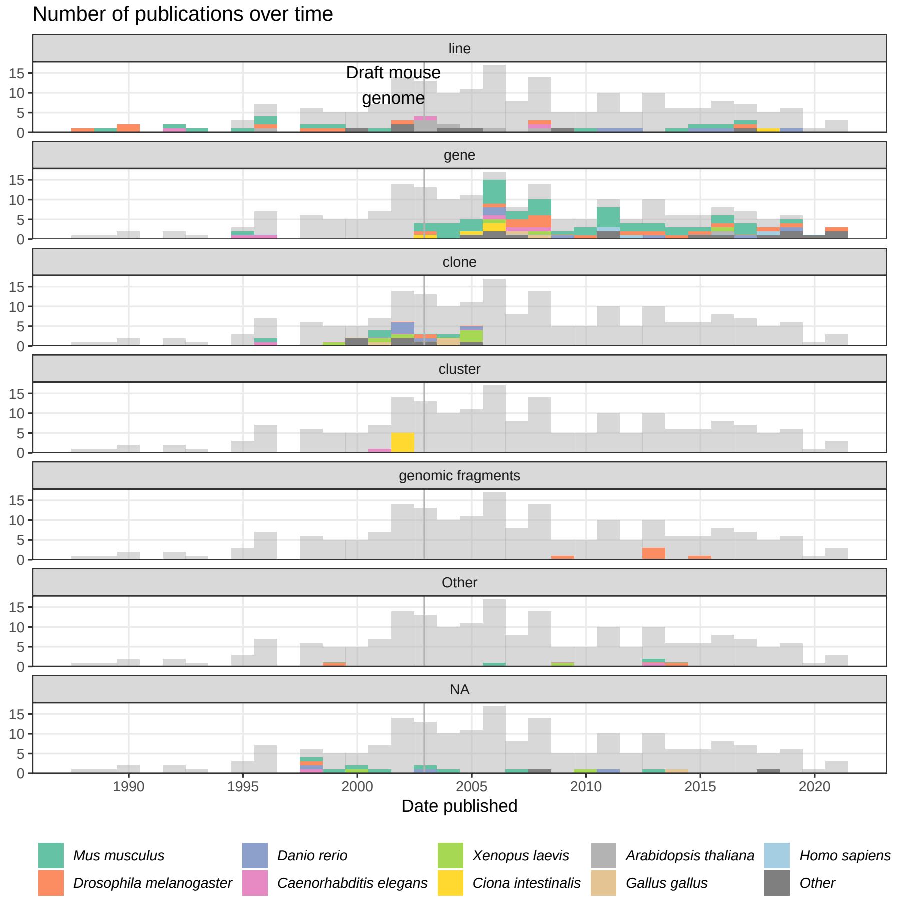

**Figure 2.4:** Number of prequel publications over time, broken down by what the entities stained for were called and colored by species. Bin width is 365 days. Vertical line marks the date when the draft mouse reference genome was published (Waterston et al. 2002), as context of transition from “clone” and “line” to “gene”.

by non-radioactive ISH by the 2000s, there is a mouse hippocampus ISH atlas published in 2004 that used radioactive ISH to profile all of its 104 genes (Lein 2004).

Also with the rise of cDNA microarray in the late 1990s and early 2000s, some (WM)ISH atlases were made as an improvement to microarray with bulk tissue to profile the transcriptome, not only at cellular resolution, but also preserving spatial and sometimes temporal context (Lein et al. 2007; Bell, Yatskievych, and Antin 2004), analogous to how scRNA-seq and various later forms of spatial transcriptomics were developed in response to bulk RNA-seq.

Since the 2000s, (WM)ISH atlases have been made for specific types of genes and a number of mouse tissues. In 2004, locked nucleic acid (LNA) modified oligonucleotide probes were introduced, greatly improving sensitivity of miRNA northern blot (Valoczi 2004) and made (WM)ISH atlases for miRNAs possible. The first miRNA WMISH atlas was published in 2005, which profiled 115 miRNAs in zebrafish embryos (Wienholds 2005). Since then, miRNA atlases were created for mice (Karali et al. 2010; Diez-Roux et al. 2011; Kloosterman et al. 2006), Drosophila (Aboobaker et al. 2005), chicken (Darnell et al. 2006), and Xenopus laevis (Ahmed et al. 2015).

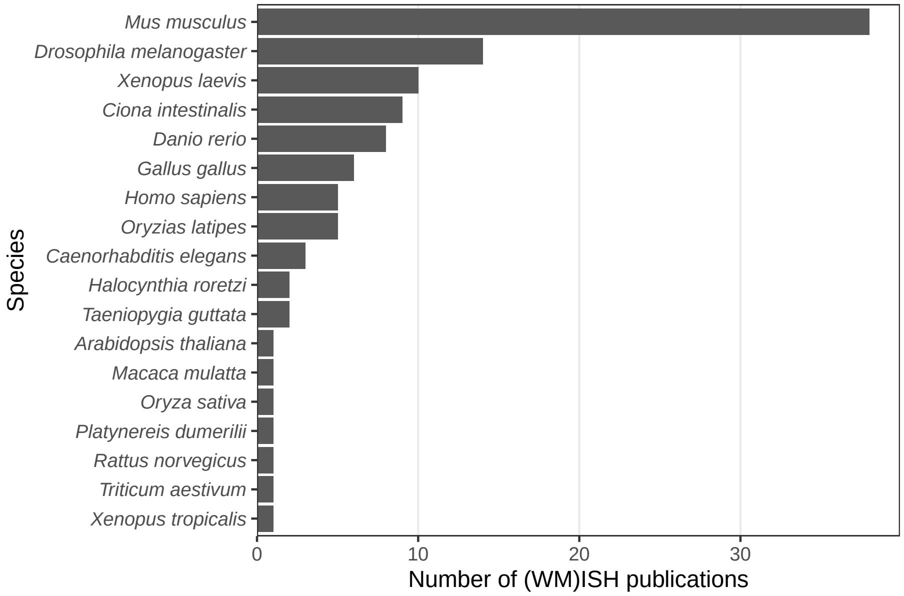

**Figure 2.5:** Number of (WM)ISH publications per species.

While (WM)ISH atlases are available for several species, the mouse is by far the favored model organism (Figure 2.5). A timeline of the first (WM)ISH atlas for each of the species and some notable atlases are shown in Figure 2.6. Especially for mice, atlases for other specific types of genes were published in the late 2000s and the 2010s, such as genes coding for RNA binding proteins (McKee et al. 2005), fibroblast growth factors and their receptors (Yaylaoglu et al. 2005), proteins with catalytic activities (Cankaya et al. 2007), transcription factors and cofactors (Yokoyama et al. 2009), metabolic enzymes and soluble carriers (Geffers et al. 2012), cholesterol biosynthetic enzymes (Şişecioğlu et al. 2015), and ion channels (in rats) (Shcherbatyy et al. 2015). Among the mouse atlases, while the brain gets disproportionately strong interests, with the influential ABA[4] (Lein et al. 2007) and GenePaint[5] (Carson, Thaller, and Eichele 2002), ISH atlases exist for the eye (Thut et al. 2001; Blackshaw et al. 2004), genitourinary tract (GenitoUrinary Development Molecular Anatomy Project (GUDMAP)[6]) (Harding et al. 2011), and lung (LungMAP[7]) (Ardini-Poleske et al. 2017) (Figure 2.6, Figure 2.7).

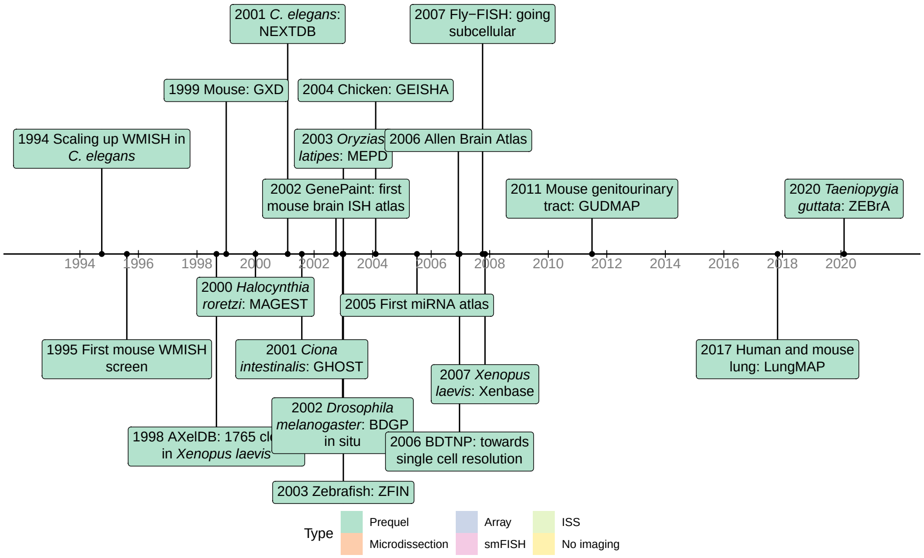

**Figure 2.6:** Timeline of the first (WM)ISH databases for each species for which such databases are available, as well as some notable databases.

While the vast majority of (WM)ISH atlases used bright field imaging, a few used FISH (Figure 2.3), for advantages conferred by FISH discussed below. A notable FISH atlas is the Berkeley *Drosophila* Transcription Network Project (BDTNP[8]) from 2006 to 2008, which profiled expression patterns of 95 genes in the *Drosophila* embryo across 6 developmental stages up to the beginning of gastrulation (Fowlkes et al. 2008; Luengo Hendriks et al. 2006). Two genes are imaged in each embryo, and the images of 1822 embryos were registered

[4] `https://portal.brain-map.org`
[5] `https://gp3.mpg.de`
[6] `https://www.gudmap.org`
[7] `https://lungmap.net`
[8] `http://www.cb.uu.se/~cris/BDTNP_Imaging.html`

across both space and time to construct 3D virtual embryos on which patterns of different genes can be quantitatively compared (Fowlkes et al. 2008); the 3D imaging and penetration into the opaque yolk is made possible by two photon microscopy, in which only the fluorophores in the region of focus are excited (Luengo Hendriks et al. 2006). Another notable FISH atlas is Fly-FISH[9] from 2007, which profiled subcellular localization of transcripts of 3370 genes in *Drosophila* embryos (Lécuyer et al. 2007). While subcellular localization of transcripts can sometimes be discerned in bright field WMISH (Tomancak et al. 2002), Fly-FISH shows higher subcellular resolution thanks to a FISH protocol using tyramide signal amplification. To our best knowledge, this is the first transcriptomic atlas of a multi-cellular organism to profile subcellular transcript localization. While more recent smFISH based methods record subcellular information, such information is typically not used in downstream analyses.

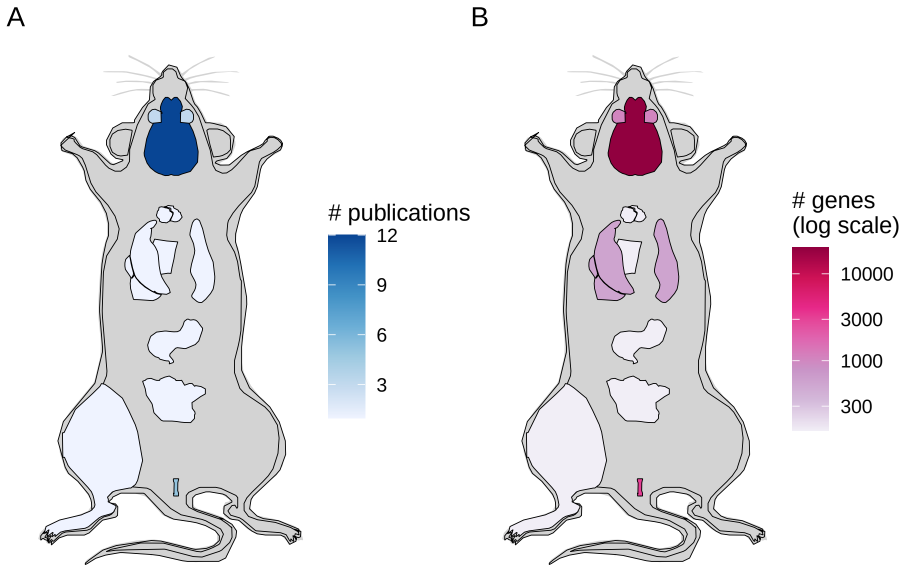

**Figure 2.7:** A) Number of mouse publications per organ for (WM)ISH atlases (including FISH). B) Maximum number of genes in atlases for each organ, as of publication of the paper about the atlases. The color is in log scale to improve dynamic range.

WMISH was the most commonly used technique in the prequel era, followed by ISH (Figure 2.8). In summary, advances of non-radioactive ISH and WMISH from radioactive ISH, limitations of enhancer and gene trap and *in situ* reporter screens, cDNA collections that cover most of predicted genes, limitations of

[9] http://fly-fish.ccbr.utoronto.ca

bulk microarray, reference genomes that allow for computational probe design, and ISH robots may have been responsible for the rise of (WM)ISH atlases. Another important factor may be the rise of digital photography and the internet in the 1990s, as developing thousands of analogue photos is an arduous task. Moreover, online digital atlases have been much more accessible to the wider community. Assuming that the number of publications in a field reflects interest in that field during a period of time, and if our collection is representative of the actual body of literature, then the golden age of the prequel era was the 2000s and WMISH was responsible for that peak, while section ISH and "collection", i.e. databases of gene expression patterns curated from publications and some (WM)ISH atlases, account for much of the interest after 2010 (Figure 2.3). The websites of many of the older (WM)ISH atlases are no longer accessible. However, some of the atlases from that period of time still live on in extant curated databases, which will be discussed in the next section.

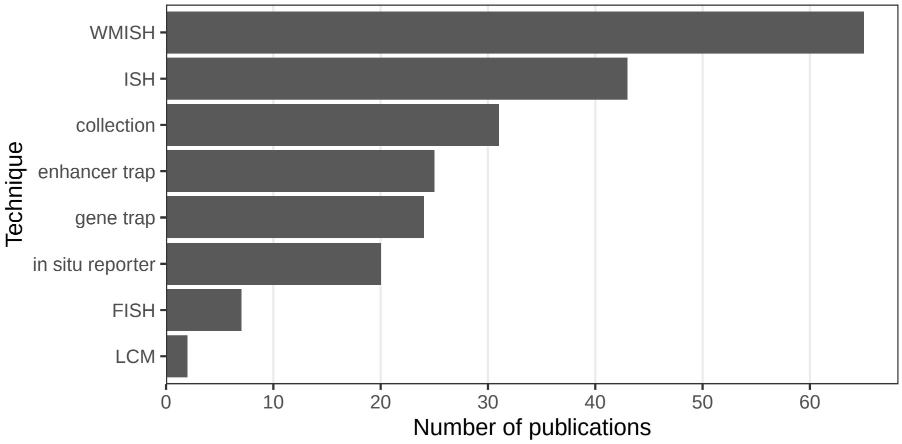

**Figure 2.8:** Number of prequel publications per technique.

The golden age declined before the rise of current era spatial transcriptomics, which started around 2014 4.2. What contributed to the decline of the golden age? Perhaps with proliferation of such atlases, curated databases exceeding 10,000 genes, and especially with over 20,000 genes in ABA mapped to a high quality 3D mouse brain model, there are already enough gene expression pattern resources for the most commonly studied genes, tissues (especially the brain), and developmental stages in the most common model organisms, thus making new atlases in those systems unnecessary. Moreover, in the last decade, the under-utilization of gene expression atlases (Boer et al. 2009) may have reduced motivation to build new atlases. Or perhaps, more importantly, inherent limitations of non-multiplexed (WM)ISH contributed to the decline in interest in such methods. In these atlases, typically only one gene is stained for in each individual embryo or tissue section. Gene expression patterns of different genes can only be meaningfully compared and classified in tissues with a stereotypical structure, such as wild type embryos and the brain, but not tumors and pathological tissues, even though there is intense interest in spatial transcriptomics in tumors as evidenced by the LCM and ST literature 6.3. A large number of embryos or sections are required for such atlases, thus increasing cost and making human atlases extremely difficult and costly, if ethical at all. Furthermore, since the chromogenic reaction in bright field ISH can be prolonged to increase staining intensity, the patterns are not quantitative and consequently, analyses of such patterns typically involve binarization and quantitative expression levels of genes cannot be compared. Even with a stereotypical structure, image registration can be challenging because of biological differences between individuals (Fowlkes et al. 2008).

#### 2.4 Databases of the prequel era

Many of the (WM)ISH atlases discussed above, such as BDGP (Tomancak et al. 2002), Gallus *In Situ* Hybridization Atlas (GEISHA)[10] (Bell, Yatskievych, and Antin 2004), ABA (Lein et al. 2007), BDTNP (Fowlkes et al. 2008), GUDMAP (Harding et al. 2011), and LungMAP (Ardini-Poleske et al. 2017) are stored in databases that can be queried online, typically by gene symbol or by controlled anatomical or developmental vocabulary (i.e. ontology, reviewed in depth in (Clarkson 2016)). There are additional gene expression databases for images curated from publications, some containing non-spatial data as well and some specifically for spatial data.

The rise of the curated databases started in the 1990s. Already in 1992, the challenges of managing the increasing amount of gene expression data in developmental biology emerged and a spatiotemporal database of mouse gene expression that would later become the Edinburgh Mouse Atlas of Gene Expression (EMAGE[11]) was discussed (Baldock et al. 1992). In 1994, Jackson Laboratory proposed the Gene Expression Database (GXD[12]) (Ringwald et al. 1994), in collaboration with EMAGE to build the most comprehensive mouse gene expression database. In 1997, work was already in progress to produce (WM)ISH atlases and construct the database infrastructure for mouse (Ringwald et al. 1997) (GXD and EMAGE), Drosophila melanogaster (Janning 1997), *C. elegans* (Martinelli, Brown, and Durbin 1997), and zebrafish (Westerfield et al. 1997). Curated databases of mice (GXD and EMAGE), zebrafish (Zebrafish Information Network (ZFIN)[13] (Howe et al. 2017)), and *Xenopus laevis* (Xenbase[14] (Bowes et al. 2009)) were released in the 2000s, within a tide of (WM)ISH atlases for new species (Figure 2.6). Some of these databases are regularly updated and the updates are responsible for many of the "collection" publications

[10] `http://geisha.arizona.edu/geisha/`
[11] `http://www.emouseatlas.org/emage/home.php`
[12] `http://www.informatics.jax.org/menus/expression_menu.shtml`
[13] `http://zfin.org`
[14] `http://www.xenbase.org/entry/`

after 2010 (Figure 2.3, Figure 2.8); our historical literature collection has not only the original publications for the databases, but also publications for later updates that involve new spatial gene expression images. Examples of other extant curated databases: for *Drosophila melanogaster* FlyExpress[15] (Kumar et al. 2017), for *Xenopus laevis* XenMARK[16] (Gilchrist et al. 2009), and for ascidians Ascidian Network for *In Situ* Expression and Embryological Data (ANISEED)[17] (Tassy et al. 2010). Databases, curated or not, are available for several species; mice, *Drosophila*, and zebrafish have the most extant databases (Figure 2.9).

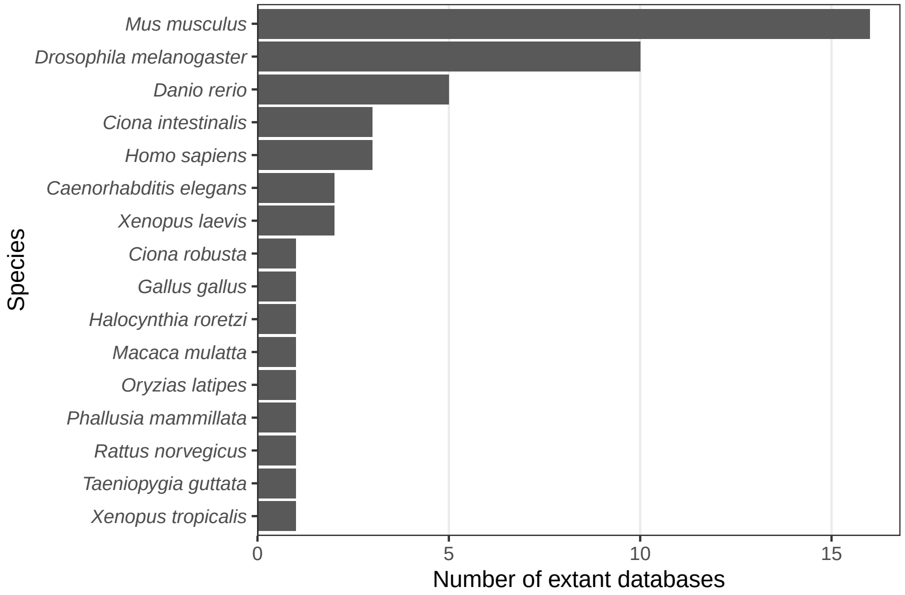

**Figure 2.9:** Number of extant spatial gene expression databases per species.

Data can be exchanged between databases. For example, among mouse databases GenePaint (Carson, Thaller, and Eichele 2002) and EMAGE now contain data from Eurexpress[18] (Diez-Roux et al. 2011; Boer et al. 2009), and EMAGE uses data from GXD for the 3D gene expression models (Ringwald et al. 1999). ANISEED contains data from WMISH atlases GHOST[19] for Ciona intestinalis (Satou et al. 2001) and MAboya Gene Expression patterns and Sequence Tags (MAGEST) for *Halocynthia roretzi* (Kawashima 2000). FlyExpress contains data from *Drosophila* atlases such as BDGP and Fly-FISH. Data in databases that ceased to operate may still be available in extant

[15] `http://www.flyexpress.net`
[16] `http://genomics.crick.ac.uk/cgi-bin/search.exe`
[17] `https://www.aniseed.cnrs.fr`
[18] `http://www.eurexpress.org/ee/`
[19] `http://ghost.zool.kyoto-u.ac.jp`

databases. For instance, AXelDb WMISH atlas and database for *Xenopus laevis* (Gawantka et al. 1998) has been subsumed in Xenbase while AXelDb's own website has long been defunct. Likewise, as of April 2021, the MAGEST website is defunct but the data lives on in ANISEED.

Some of the databases go beyond collecting data from other databases. Databases such as EMAGE, ANISEED, and ABA registered multiple 2D section images to map gene expression patterns onto 3D anatomical models for better comparison between different genes. FlyExpress also standardized the images from the atlases and enables search for coexpressed genes by expression pattern (Kumar et al. 2017). There have also been efforts to integrate databases from multiple model organisms. In 2007, COMPARE (Salgado et al. 2008) and 4DXpress (Haudry et al. 2007) were developed to make gene expression patterns and developmental stages in zebrafish, mouse, and *Drosophila* (also medaka in 4DXpress) comparable. While COMPARE and 4DXpress are no longer available, interest in integrating the databases continues, so in 2016, the Alliance of Genome Resource was founded, producing a unified user interface to genome and gene expression databases for *Saccharomyces cerevisiae*, *C. elegans*, *Drosophila melanogaster*, mouse, rat, and zebrafish (Agapite et al. 2020), although spatial patterns are not its focus.

#### 2.5 Geography of the prequel era

Where were prequel era research conducted? Our database includes affiliation of the first author as of publication for all papers, and the affiliations have been geocoded to plot on maps. Around the world, most of prequel studies were performed in coastal US and Western Europe, but a some studies were performed in Asia and Oceania, but especially Japan (Figure 2.10). Not all of the top contributing institutions are readily recognizable "elite" institutions. Institutions include BDGP from UC Berkeley, ZFIN from University of Oregon (UO), ABA from Allen Brain Institute (Allen), GEISHA from University of Arizona (UofA), GXD from Jackson Laboratory (JAX), EMAGE from Western General Hospital (WGH), MEPD[20] (for *Oryzias latipes*) from European Molecular Biology Laboratory (EMBL), and GHOST from Kyoto University (Kyodai), and mouse gene trap lines from Mount Sinai.

This can be better visualized by breaking the map down by species. Here we see locations of some model organism consortia, and that GHOST is a result of collaboration of multiple Japanese institutions (Figure 2.11).

That some institutions have disproportional contribution of one technique can also be shown. Here it's clear that prequel techniques are used by many different institutions (Figure 2.12). In contrast, as will be shown in Chapter 4, most current era techniques never spread beyond their institutions of origin.

[20] `https://www.embl-heidelberg.de/mepd/`

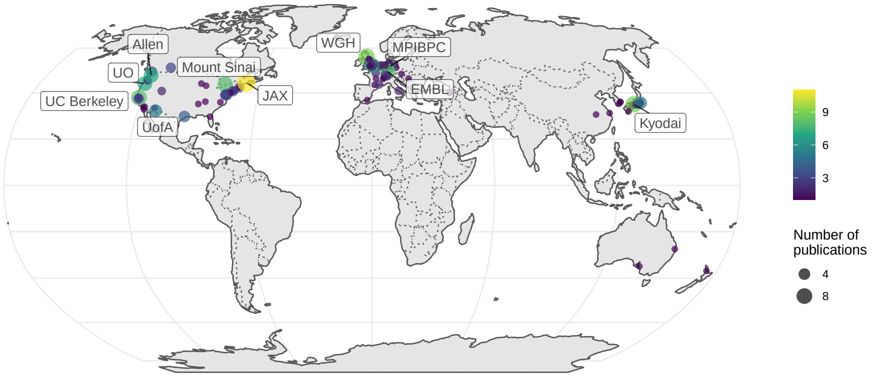

**Figure 2.10:** Number of prequel publications per city around the world, with top contributing institutions labeled.

The LCM study comes from Allen Brain Institute's atlases for Allen's mouse sleep deprivation atlas (Thompson et al. 2010) and human glioblastoma atlas (Puchalski et al. 2018); although LCM is a current era technique, those two studies are in the prequel sheet because they also have ISH atlases.

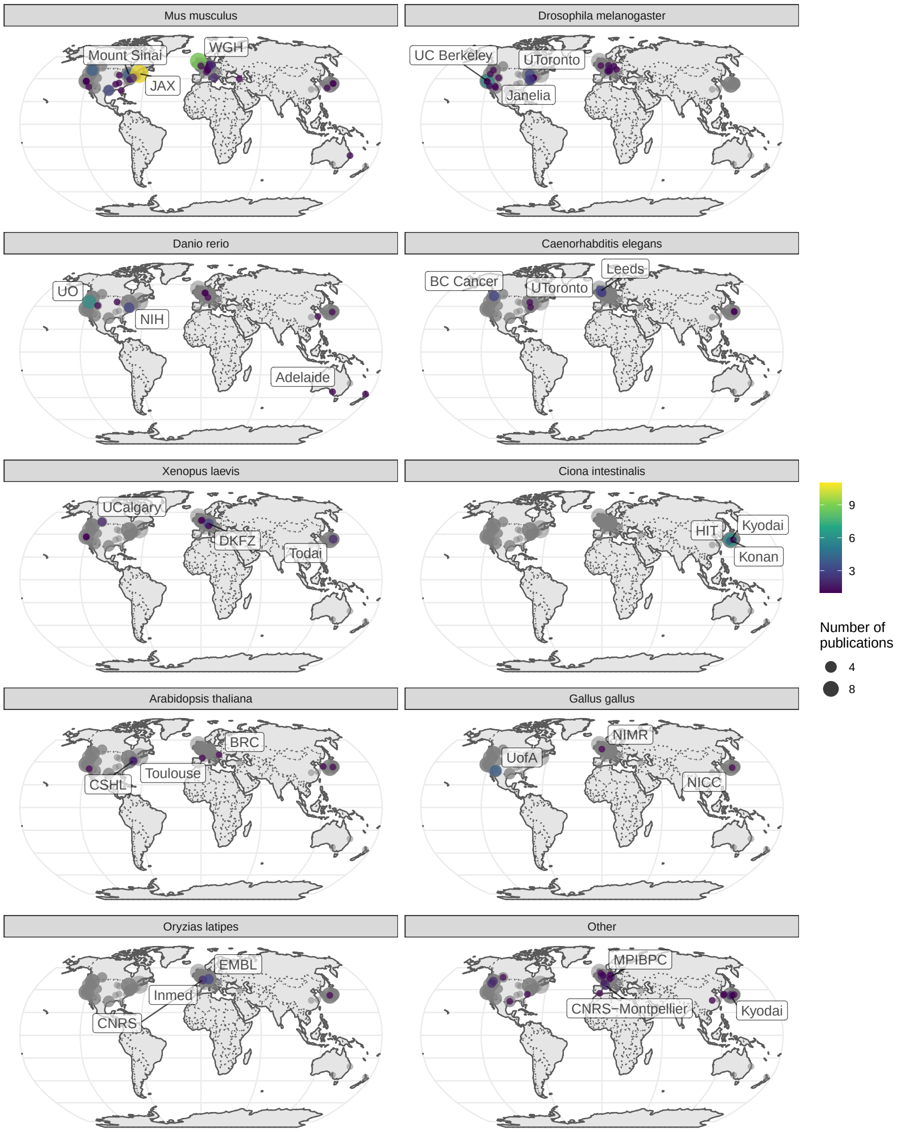

**Figure 2.11:** Number of prequel publications per city broken down by species. Gray points are the overall number as a reference of contributions from each city and region.

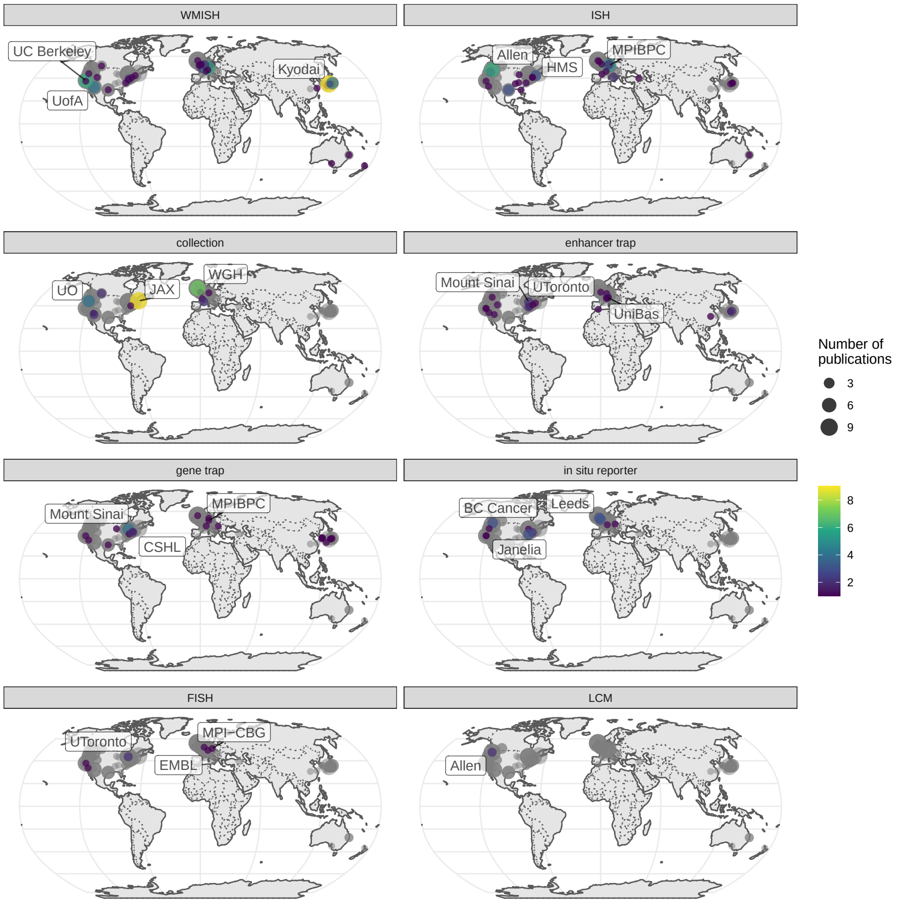

**Figure 2.12:** Number of prequel publications per city broken down by technique. Gray points are the overall number as a reference of contributions from each city and region.

### Chapter 3

### Data analysis in the prequel era

Many machine learning and statistics methods are mentioned in this chapter. The names of these methods are linked to articles explaining them for those who are unfamiliar. Some of them are math heavy.

From the earliest days of enhancer and gene traps to the (WM)ISH atlases, identifying genes with spatially and temporally variable expression patterns, comparing and classifying the patterns, identifying new marker genes of cell types and developmental stages, and using gene expression to redefine cell types have been among the goals of the studies (O'Kane and Gehring 1987; Gossler et al. 1989; Wurst et al. 1995; Sundaresan et al. 1995; Gawantka et al. 1998; Tomancak et al. 2002; Lein et al. 2007). In the prequel era, these were typically done manually, which, with the growing size of atlases in the 2000s, was time consuming and potentially inconsistent between curators. Thus, computational methods were developed to analyze images from the (WM)ISH atlases. This chapter reviews data analysis methods designed for (WM)ISH atlases and does not involve scRNA-seq data; methods involving both (WM)ISH and scRNA-seq are reviewed in Chapter 7 for the current era because scRNA-seq is at present a popular and rapidly growing field, too in vogue to be considered "prequel". If our collection is representative, then the rise of prequel data analysis methods arrived much later than that of data collection (Figure 3.1).

Except for one study on *Platynereis dumereilii* in 2014 (Pettit et al. 2014), on *Xenopus tropicalis* in 2018 (Patrushev et al. 2018), one on post mortem human brain in 2021 (Abed-Esfahani et al. 2021), all data analysis methods in our collection were designed for either *Drosophila melanogaster* or *Mus musculus*

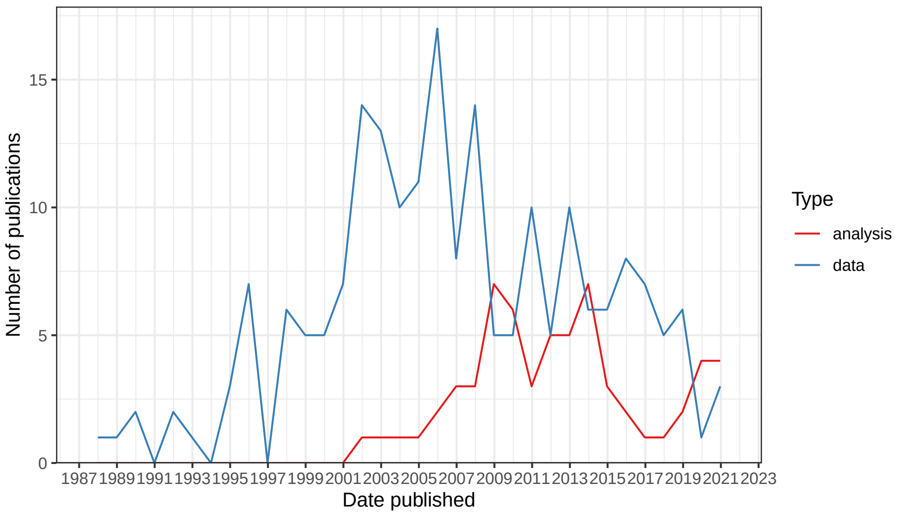

**Figure 3.1:** Comparing trends in data collection and data analysis in the prequel era. Bin width is 365 days.

(Figure 3.2). There seem to have been two waves; the first for *Drosophila*, peaking in the late 2000s, mostly concerning the BDGP in situ atlas, and the second for mice, peaking in early 2010s, mostly concerning ABA (Figure 3.2). The apparent rise since 2019 is in part driven by deep learning methods to annotate gene expression patterns or infer gene interactions. Given the small number of publications in this category and potential incompleteness of the curation, the trends should be taken with a grain of salt.

#### 3.1 Gene patterns

The most common goal of these data analysis methods was to annotate and compare gene expression patterns, especially to automate annotation of the BDGP atlas (Figure 3.3). It seems reasonable to focus on 4 phases in this category: first, in early to mid 2000s, after image registration, the images were binarized into "expressed" and "not expressed" regions, and the shapes of the expressed regions were summarized and compared. Metrics to summarize the shapes included moment invariant[1] (Jayaraman, Panchanathan, and Kumar 2001; Gurunathan et al. 2004), Hamming distance (Kumar et al. 2002), and a weighted score involving L1 distance[2] between column or row histograms of two images (Liu et al. 2007). These unsupervised methods enabled clustering

[1] `https://towardsdatascience.com/introduction-to-the-invariant-moment-and-its-application-to-the-feature-extraction-ee991f39ec`

[2] `https://iq.opengenus.org/manhattan-distance/`

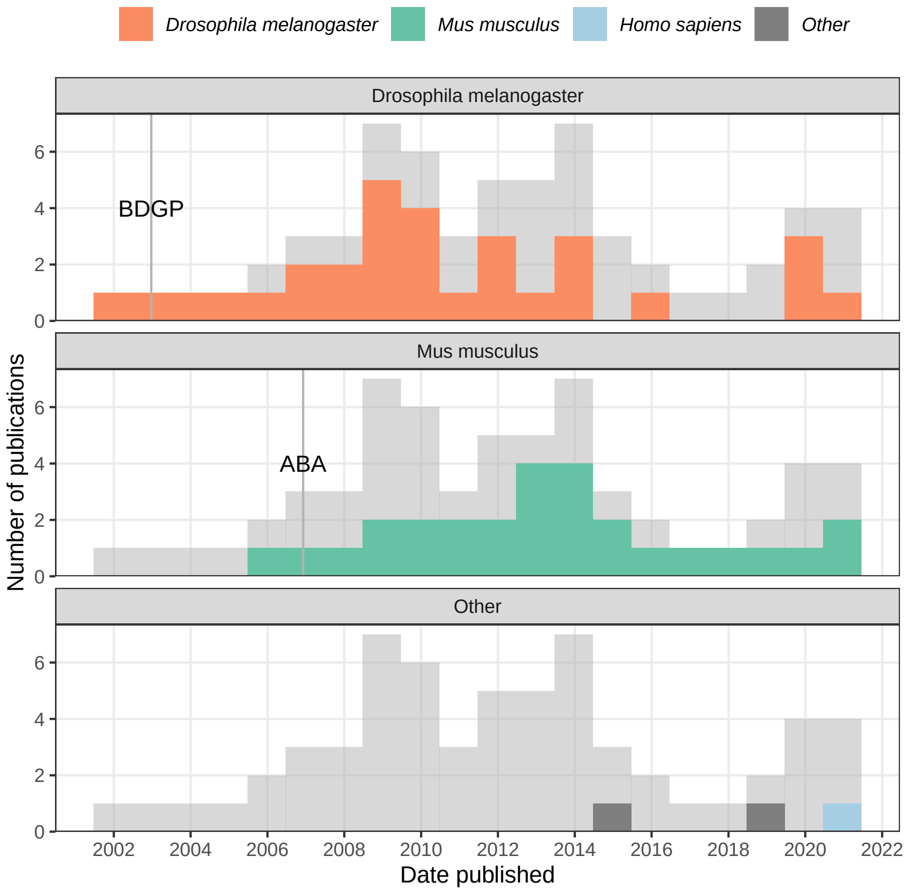

**Figure 3.2:** Gray histogram in the background is overall histogram of prequel data analysis literature. Number of publications in each time bin for each species is highlighted in the facets.

of patterns and querying genes with similar patterns to a given gene.

Second, from the mid 2000s to mid 2010s, many supervised and unsupervised methods for gene expression pattern annotation or comparison were developed. In supervised methods, extensive feature engineering more sophisticated than binarization was performed on registered images for image annotation with machine learning classification. These methods were trained with existing BDGP annotations and developed to automatically annotate the BDGP expression patterns with controlled vocabulary (CV) of anatomical regions where genes were expressed. In BDGP, a gene gets annotated with a CV if the gene was deemed expressed in the anatomical region and developmental stage denoted by the CV, so the annotation typically contained a list of CVs.

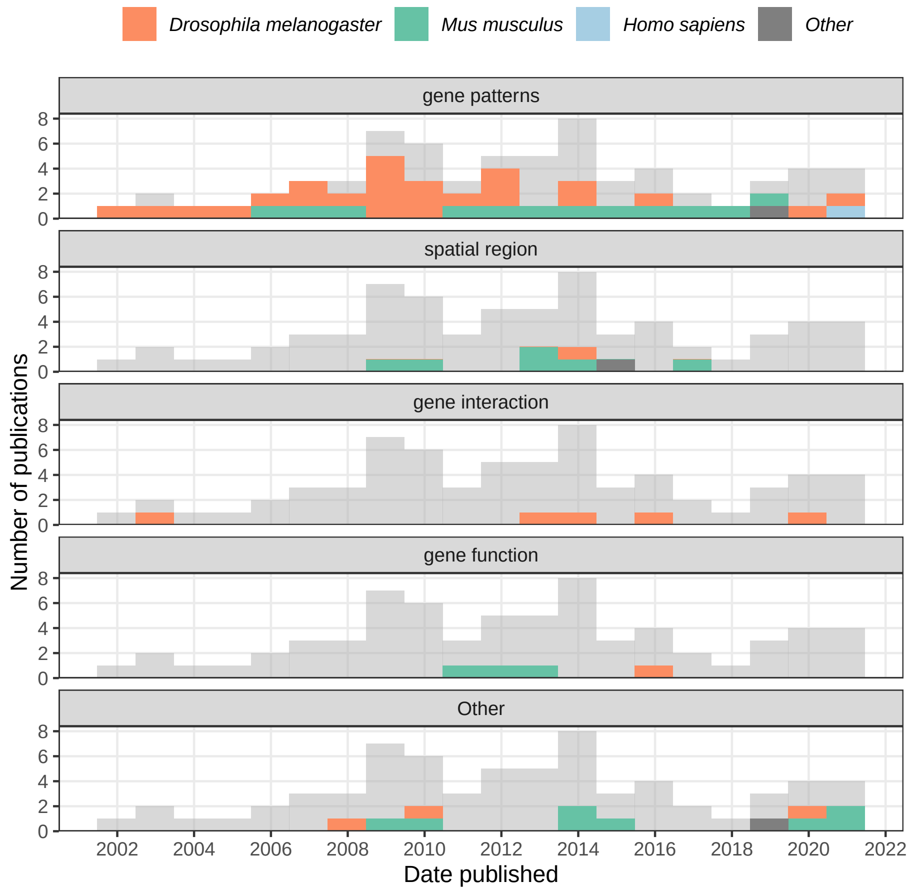

**Figure 3.3:** Number of publications in each time bin for each category of data analysis is highlighted in the facets.

The feature engineering can be based on the wavelet transform (Zhou and Peng 2007) and Fourier coefficients (Heffel et al. 2008), but a particularly popular feature engineering method was scale-invariant feature transform (SIFT)[3] (Lowe 2004; Ji et al. 2008; Li et al. 2009; S. Ji, Li, et al. 2009). A method published in 2009 that used SIFT followed by bag of words[4] where "word" is a k means[5] cluster (code book) was quite influential (S. Ji, Li, et al. 2009); several later methods were inspired by this method, with improved code books (Sun et al. 2013; S. Ji, Yuan, et al. 2009; Yuan et al. 2012; Liscovitch, Shalit, and Chechik 2013). The most common classifier that take in the features to predict annota-

[3] `https://docs.opencv.org/master/da/df5/tutorial_py_sift_intro.html`
[4] `https://machinelearningmastery.com/gentle-introduction-bag-words-model/`
[5] `https://medium.com/analytics-vidhya/k-means-clustering-explained-419ee66d095e`

tions is support vector machine (SVM)[6] (Sun et al. 2013; Yuan et al. 2012) or multi-label variants of it (Ji et al. 2008; S. Ji, Li, et al. 2009).

Unsupervised methods rely on clustering algorithms after images are registered on a common mesh, such as affinity propagation clustering[7] (Frise, Hammonds, and Celniker 2010) and co-clustering (rows and columns of matrix are clustered simultaneously) (Jagalur et al. 2007; Zhang et al. 2013).

Third, another notable type of the feature engineering is dimension reduction. In 2006, some methods applied dimension reduction methods such as principal component analysis (PCA)[8] and independent component analysis (ICA)[9] to the registered images to find "eigen" patterns (Pan et al. 2006; Hanchuan Peng et al. 2006). Instead of PCA or ICA, the dimension reduction can also be sparse Bayesian factor analysis (Pruteanu-Malinici, Mace, and Ohler 2011), sparse dictionary learning (Li et al. 2017), and non-negative matrix factorization (NMF)[10] (Noto, Barnagian, and Castro 2017; Wu et al. 2016). The dimension reduction can be used for unsupervised clustering of genes (Pan et al. 2006; Hanchuan Peng et al. 2006; Pruteanu-Malinici, Mace, and Ohler 2011), as well as supervised classification methods such as SVM and logistic regression[11] to annotate gene expression patterns with controlled vocabulary (Pruteanu-Malinici, Mace, and Ohler 2011; Wu et al. 2016). Notably, in NMF, both the matrix for basis patterns and the coefficient matrix for the genes tend to exhibit block structures; the blocks in the gene coefficient matrix has been used to cluster genes (Noto, Barnagian, and Castro 2017).

Fourth, since 2015, convolutional neural networks (CNNs)[12] have been adopted to analyze gene expression patterns. Typically, a pre-trained model, such as ResNet50, OverFeat, or Alexnet is used. With some modifications or retraining of the original model, the model can be used to extract features for gene pattern annotation with logistic regression (Zeng et al. 2015), classifying new patterns (Andonian et al. 2019, @Long2021), and predicting interactions between genes (Yang, Fang, and Shen 2019).

---

[6] `https://towardsdatascience.com/support-vector-machine-introduction-to-machine-learning-algorithms-934a444fca47`

[7] `https://towardsdatascience.com/unsupervised-machine-learning-affinity-propagation-algorithm-explained-d1fef85f22c8`

[8] `https://liorpachter.wordpress.com/2014/05/26/what-is-principal-component-analysis/`

[9] `http://wwwf.imperial.ac.uk/~nsjones/TalkSlides/HyvarinenSlides.pdf`

[10] `http://www.cs.cmu.edu/~11755/lectures/Lee_Seung_NMF.pdf`

[11] `https://towardsdatascience.com/logistic-regression-detailed-overview-46c4da4303bc`

[12] `https://towardsdatascience.com/a-comprehensive-guide-to-convolutional-neural-networks-the-eli5-way-3bd2b1164a53`

#### 3.2 Spatial regions

Closely related to classifying gene expression patterns are these questions: What are the implications of gene expression patterns to traditional anatomical regions as in the CV? Can we discover novel anatomical regions from gene expression? How well do expression-based regions correspond to the traditional regions? A few studies, which we call "spatial region", tried to answer these questions in the ABA (Figure 3.3). Clusters of expression patterns of cell type specific genes (Ko et al. 2013), or the most localized genes (Grange et al. 2014), principal components of the patterns (Bohland et al. 2010), or patterns of coexpression modules were compared to traditional anatomy (Grange et al. 2014). At least in the mouse brain, while with the principal components, these clusters may correspond to traditional anatomy quite well (Bohland et al. 2010), when cell types are taken into account in clustering, gene expression seems to be able to refine traditional anatomy (Ko et al. 2013; Grange et al. 2014).

A clustering strategy for identifying spatial regions that takes the spatial neighborhood into account is Markov random field (MRF)[13]. In MRFs, nearby voxels can be made to be more likely to share a label, which can be cell type or histological region, and the probability of a voxel taking each of the labels only depends on labels of neighboring voxels. MRFs were used to delineate spatial regions in a 3D FISH atlas of the developing *Platynereis dumereilii*[14] brain (Pettit et al. 2014), with 86 high quality genes. The images in the atlas were aligned into a 3D model and broken into voxels 3 $\mu$m per side, which is smaller than a typical single cell; the spatial neighborhood graph is the 3D square grid of the voxels. As FISH is not very quantitative, the gene expression was manually binarized. Expression of each gene at each voxel is modeled with a Bernoulli distribution[15], and the 86 genes are assumed to be independent. Cluster label assignment is modeled with Potts model[16], a type of MRF in which only neighboring voxels with the same label contribute to the probability distribution of the labels, thus favoring neighbors with the same label. The parameters, such as interaction strength between neighboring voxels for the Potts model and the probability parameter of the Bernoulli distributions are estimated with expectation maximization (EM)[17].

#### 3.3 Gene interactions

While not single cell resolution, (WM)ISH atlases provide transcriptomes within the tissue at a resolution far higher than that of typical bulk RNA-seq and bulk

[13] `https://ocw.mit.edu/courses/electrical-engineering-and-computer-science/6-867-machine-learning-fall-2006/lecture-notes/lec23.pdf`
[14] `https://platynereis.github.io/`
[15] `https://mathworld.wolfram.com/BernoulliDistribution.html`
[16] `https://en.wikipedia.org/wiki/Potts_model`
[17] `http://ai.stanford.edu/~chuongdo/papers/em_tutorial.pdf`

microarray, thus opening the way to studying coexperssion and interaction between genes within the tissue. There are a few methods that aim to decide whether two genes interact according to (WM)ISH images, some dating published long before the popularization of scRNA-seq. Already in 2002, an early method that compares binarized gene expression patterns was used to identify interactions among genes by comparing patterns from wild type and mutant backgrounds (Kumar et al. 2002).

However, as mutant lines are harder to obtain than wild type images, the simplest method is to set a threshold in Pearson correlation[18] coefficient between two genes to decide an edge should be drawn on the gene coexpression graph (Wu et al. 2016; Campiteli et al. 2013).

Alternatively, a sparse Markov network[19] whose nodes are genes and edges are presence of interaction can be learnt from expression profiles in each voxel (Puniyani and Xing 2013), or a CNN can be trained on known interactions and predict new interactions based on gene expression patterns (Yang, Fang, and Shen 2019). There are other types of analyses, such as inferring gene function from expression pattern, identifying spatially variable genes, and gene expression imputation at locations. The latter two are still important topics in current era data analysis.

#### 3.4 Decline

What contributed to the decline of the golden age of prequel data analysis? Partly a lack of usage of the methods developed, which was exacerbated by the decline of the golden age of (WM)ISH atlases in the 2010s (Figure 2.3). While many methods to automate gene expression pattern annotation for BDGP were developed before 2013, for the 2013 BDGP update that added images of 708 transcription factors, the BDGP annotated the new images with human curators instead of the automated methods (Hammonds et al. 2013). Nor did BDGP use the new methods to compare and classify the new gene expression patterns; instead, the curator assigned CV annotations were used for analysis (Hammonds et al. 2013; Tomancak et al. 2007). BDGP did not have a major update after 2013; as existing images have already been annotated, there is no need to automate annotations.

There are additional possible reasons why these methods were not used: First, it is unclear from the publications of the methods where the software implementation can be obtained. Second, a reason why most prequel analysis methods were developed for either BDGP or ABA is that since one gene is stained for in one embryo/section at a time, the images must be registered and standardized for different genes to be comparable; BDGP, through FlyExpress (Kumar et al.

[18] https://www.questionpro.com/blog/pearson-correlation-coefficient/

[19] http://ml.informatik.uni-freiburg.de/former/_media/teaching/ws1314/gm/11-markov_logic_networks.handout.pdf

2017), and ABA, provide images that have already been registered and standardized, while many other atlases, such as GEISHA, do not. Due to challenges in image registration in other organisms, the automated gene expression pattern analysis methods can't be applied. Third, lack of usage of these methods can also be due to insufficient accuracy; from 2009 to 2013, the area under the curve (AUC)[20] of the automated annotations is typically around 0.8 and rarely exceeded 0.9 (S. Ji, Li, et al. 2009; Pruteanu-Malinici, Mace, and Ohler 2011; Yuan et al. 2012; Sun et al. 2013), which means when using such tools to annotate new images, extensive human review would still be required.

#### 3.5 Geography of prequel data analysis

If our collection is representative, then contribution to prequel data analysis concentrates in a few institutions (Figure 3.4), not all of which are elite.

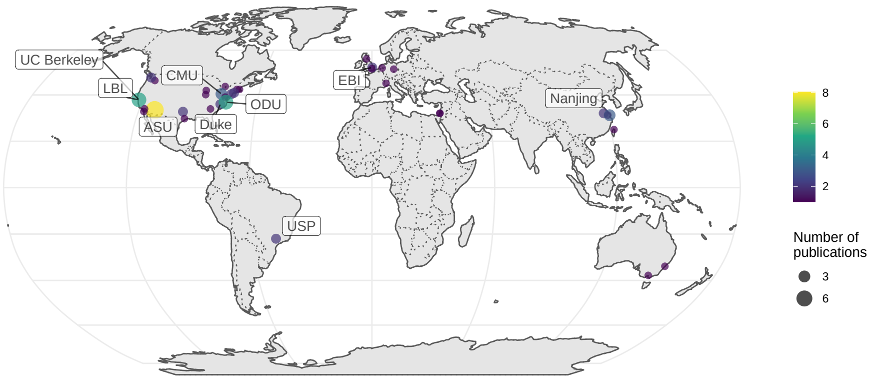

**Figure 3.4:** Number of publications per city for prequel data analysis.

When broken down by species, it seems that distinct institutions contributed to data analysis of *Drosophila* and mouse data. UC Berkeley and Lawrence Berkeley National Laboratory (LBL) are responsible for BDGP, and Allen is responsible for ABA. However, among the top contributors are other institutions such as Arizona State University (ASU) and Old Dominion University (ODU) (Figure 3.5).

[20]https://towardsdatascience.com/understanding-auc-roc-curve-68b2303cc9c5

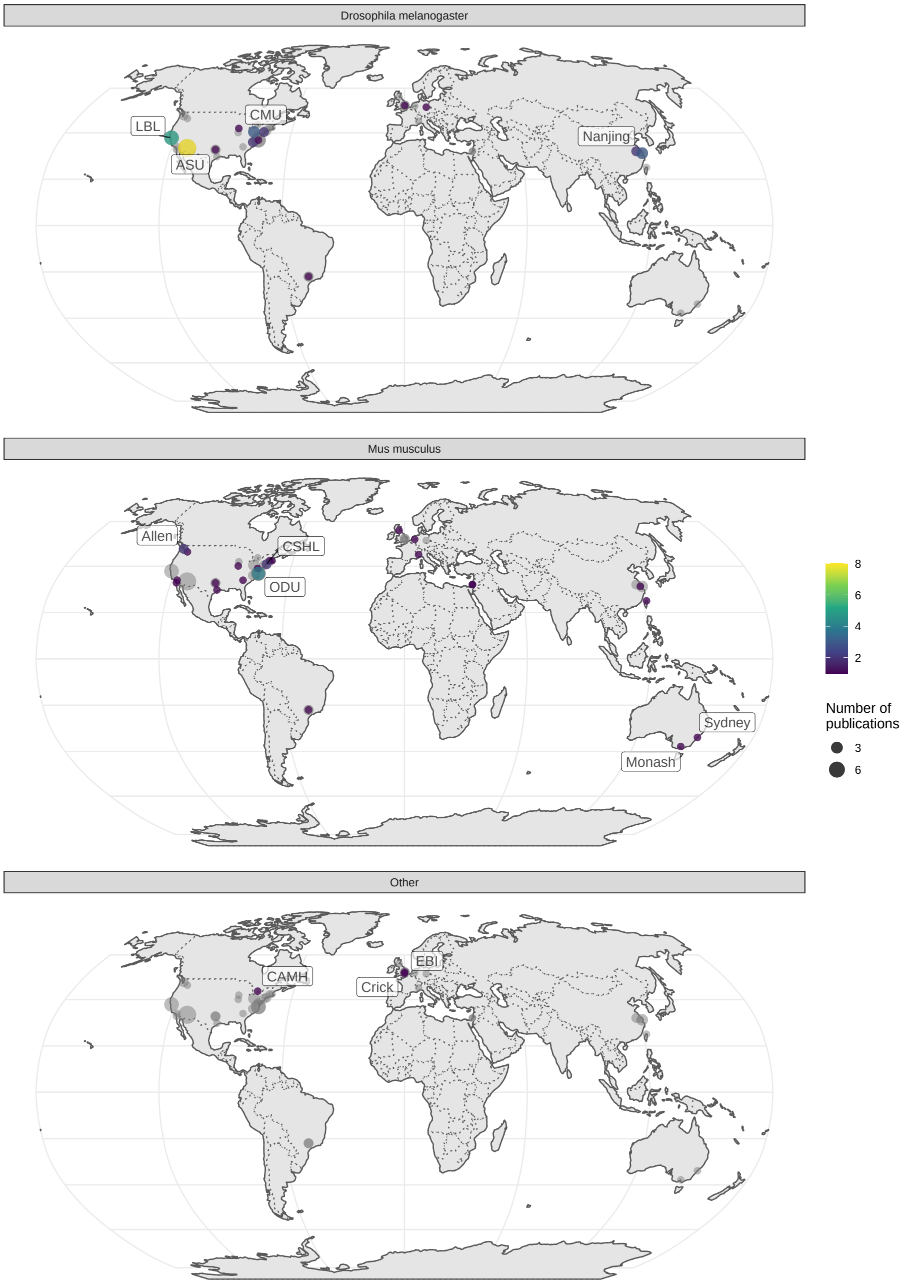

**Figure 3.5:** Number of publications per city for prequel data analysis broken down by species of interest.

### Part II

### Current era

### Chapter 4

### From the past to the present

The current era continues many of the quests of the prequel era, such as to profile the transcriptome in space, to identify genes with restricted expression, to classify gene expression patterns, to build reference gene expression atlases for model systems, and to infer anatomical regions based on gene expression. While the prequel era also sought to identify cell type markers, this has been taken over by non-spatial transcriptomics, which has been used to identify marker genes to stain for with spatial transcriptomics methods not easily scalable to the whole transcriptome. As already mentioned, (WM)ISH atlases can be understood as an improved alternative to microarray and *in situ* reporter screens, and the latter can be in turn understood as an improved alternative to enhancer and gene traps. To some extent, current era spatial transcriptomics started as an improved alternative to (WM)ISH atlases, to profile the whole transcriptome in the same cells (Junker et al. 2014; Lee et al. 2014). On the other hand, part of current era of spatial transcriptomics can be seen as an improvement to bulk microarray or RNA-seq (V. M. Brown et al. 2002; Junker et al. 2014; Ståhl et al. 2016; Luo et al. 1999), and lower throughput single cell biology (Lubeck and Cai 2012; K. H. Chen et al. 2015).

**Table 4.1:** Summary of spatial transcriptomics techniques in the current era

| Method | First published | Category | Max # genes | Min spot diameter ($\mu$m) |
|---|---|---|---|---|
| voxelation | 2002-02-01 | Microdissection | Tx wide | NA |
| SRM seqFISH | 2012-06-03 | smFISH | 32 | single cell |
| Tomo-array | 2012-09-19 | Microdissection | Tx wide | NA |

**Table 4.1:** Summary of spatial transcriptomics techniques in the current era *(continued)*

| Method | First published | Category | Max # genes | Min spot diameter ($\mu$m) |
|---|---|---|---|---|
| iceFISH | 2013-02-17 | smFISH | 20 | single cell |
| ISS | 2013-07-14 | ISS | 222 | single cell |
| Tomo-seq | 2013-08-12 | Microdissection | Tx wide | NA |
| bDNA-smFISH | 2013-10-06 | smFISH | 928 | single cell |
| TIVA | 2014-01-12 | Microdissection | Tx wide | NA |
| FISSEQ | 2014-03-21 | ISS | 8102 | single cell |
| seqFISH | 2014-03-28 | smFISH | 10421 | single cell |
| MERFISH | 2015-04-24 | smFISH | 4209 | single cell |
| Puzzle Imaging | 2015-07-20 | No imaging | NA | NA |
| Geo-seq | 2016-03-21 | Microdissection | Tx wide | NA |
| corrFISH | 2016-06-06 | smFISH | 10 | single cell |
| ST | 2016-07-01 | Array | Tx wide | 100 |
| HCR-seqFISH | 2016-10-19 | smFISH | 249 | single cell |
| punch | 2017-06-28 | Microdissection | Tx wide | NA |
| SGA | 2017-11-28 | smFISH | 35 | single cell |
| APEX-RIP | 2017-12-14 | No imaging | NA | NA |
| Niche-seq | 2017-12-22 | Microdissection | Tx wide | NA |
| ExM-MERFISH | 2018-03-19 | smFISH | 10050 | single cell |
| STARmap | 2018-07-27 | ISS | 1020 | single cell |
| Paired-cell sequencing | 2018-09-17 | No imaging | NA | NA |
| osmFISH | 2018-10-30 | smFISH | 33 | single cell |
| seqFISH+ | 2019-03-25 | smFISH | 10000 | single cell |
| slide-seq | 2019-03-29 | Array | Tx wide | 10 |
| bDNA-MERFISH | 2019-05-25 | smFISH | 130 | single cell |
| DNA microscopy | 2019-06-27 | No imaging | NA | NA |
| APEX-seq | 2019-07-11 | No imaging | NA | NA |
| INSTA-seq | 2019-08-06 | ISS | NA | single cell |
| PARSIFT | 2019-09-04 | No imaging | NA | NA |
| HDST | 2019-09-09 | Array | Tx wide | 2 |
| GaST-seq | 2019-10-10 | Microdissection | Tx wide | NA |

**Table 4.1:** Summary of spatial transcriptomics techniques in the current era *(continued)*

| Method | First published | Category | Max # genes | Min spot diameter ($\mu$m) |
|---|---|---|---|---|
| BARseq | 2019-10-17 | ISS | 79 | single cell |
| SCRINSHOT | 2020-02-07 | smFISH | 29 | single cell |
| Visium | 2020-02-28 | Array | Tx wide | 55 |
| PIC | 2020-03-23 | Microdissection | Tx wide | NA |
| miRNA nanowell | 2020-05-09 | Array | 9 | 300 |
| GeoMX DSP | 2020-05-11 | Microdissection | 2093 | NA |
| split-FISH | 2020-06-15 | smFISH | 317 | single cell |
| HybISS | 2020-07-03 | smFISH | 120 | single cell |
| ZipSeq | 2020-07-06 | Microdissection | Tx wide | NA |
| ClumpSeq | 2020-08-06 | No imaging | NA | NA |
| BARseq2 | 2020-08-26 | ISS | 65 | single cell |
| SM-Omics | 2020-10-15 | Array | Tx wide | 100 |
| slide-seq2 | 2020-10-17 | Array | Tx wide | 10 |
| DBiT-seq | 2020-10-19 | Array | Tx wide | 10 |
| C-FISH | 2020-10-23 | smFISH | 2 | single cell |
| HybRISS | 2020-12-02 | smFISH | 50 | single cell |
| Stereo-seq | 2021-01-19 | Array | Tx wide | 0.22 |
| Seq-Scope | 2021-01-27 | Array | Tx wide | 0.5 |
| ExSeq | 2021-01-29 | ISS | 297 | single cell |
| par-seqFISH | 2021-02-25 | smFISH | 105 | single cell |
| EASI-FISH | 2021-03-08 | smFISH | 26 | single cell |
| Pick-Seq | 2021-03-09 | Microdissection | Tx wide | NA |
| nanoneedles | 2021-03-10 | Microdissection | 9 | NA |
| PIXEL-seq | 2021-03-17 | Array | Tx wide | 1.22 |
| GeoMX WTA | 2021-03-20 | Microdissection | 18190 | NA |
| CISI | 2021-04-15 | smFISH | 37 | single cell |
| STRP-seq | 2021-04-19 | Microdissection | Tx wide | NA |
| XYZeq | 2021-04-21 | Array | Tx wide | 500 |
| electro-seq | 2021-04-23 | ISS | 201 | single cell |
| centrifugation on 384 well plate | 2021-04-30 | Microdissection | Tx wide | NA |

The current era started with LCM followed by microarray in 1999 (Luo et al. 1999). Due to the immense popularity of LCM followed by microarray or RNA-seq, the body of LCM literature is too vast for unbiased and comprehensive manual curation, so the curated database does not include most LCM literature, which was instead collected from a PubMed search and text mined (Figure 6.3, Chapter 6). Because the search results—without manual inspection and curation—may contain irrelevant entries and miss relevant ones, they are separated from the curated database in our analyses. Current era literature in the curated database is classified into Microdissection, smFISH, ISS, Array, and No Imaging, to be defined in detail in their corresponding sections below.

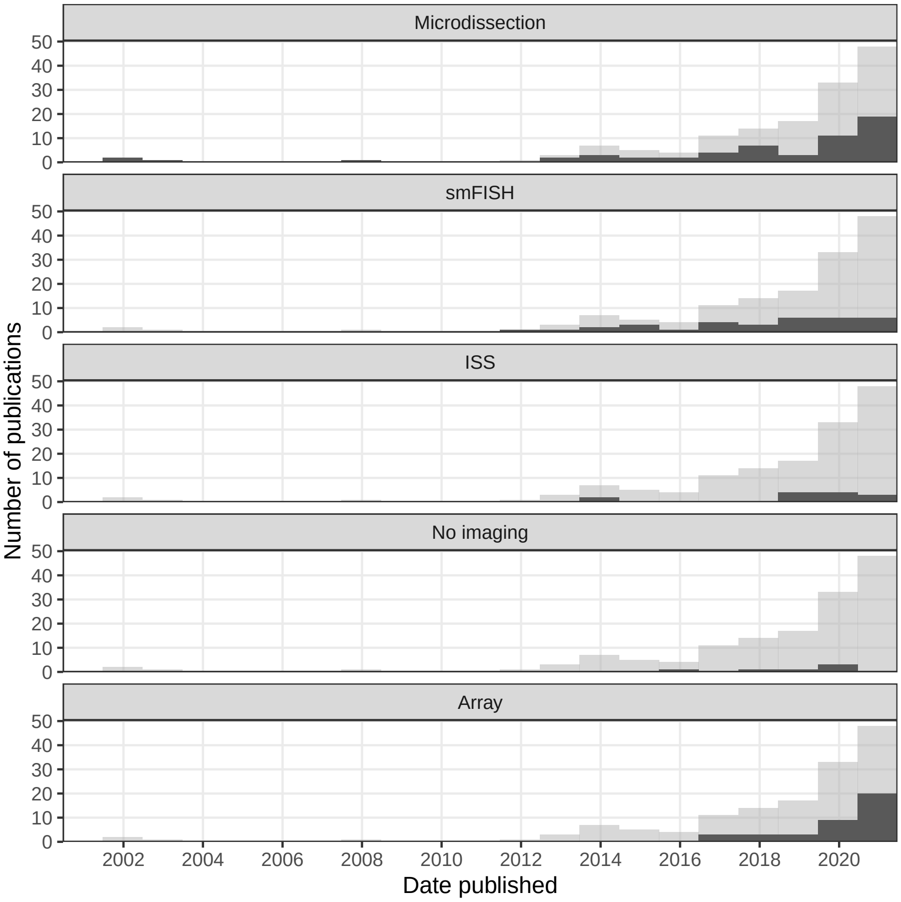

**Figure 4.1:** Number of publications over time in the current era. The gray histogram in the background is the overall trend of all current era literature. Each facet highlights a category, ordered chronologically in terms of first report. Bin width is 365 days. Plots in this figure include curated LCM literature, but not the non-curated literature.

Chronologically, in the curated database, microdissection came first, with voxelation in 2002 (V. M. Brown et al. 2002), followed by smFISH, ISS, no imaging, and array (Figure 4.1). Despite an early start in the midst of the (WM)ISH golden age, if not including non-curated LCM literature, the current era did not really take off until around 2014 (Figure 4.2). Ever since, its has seen drastic growth, far exceeding that of the prequel era in the 1990s and 2000s (Figure 4.2). Growth in microdissection and array seemed to have contributed the most to this overall drastic growth (Figure 4.1). All techniques in the curated database, along with their classification, maximum number of genes, spatial resolution, and references are listed in Table 4.1. A timeline of major techniques in the current era is shown in Figure 4.3.

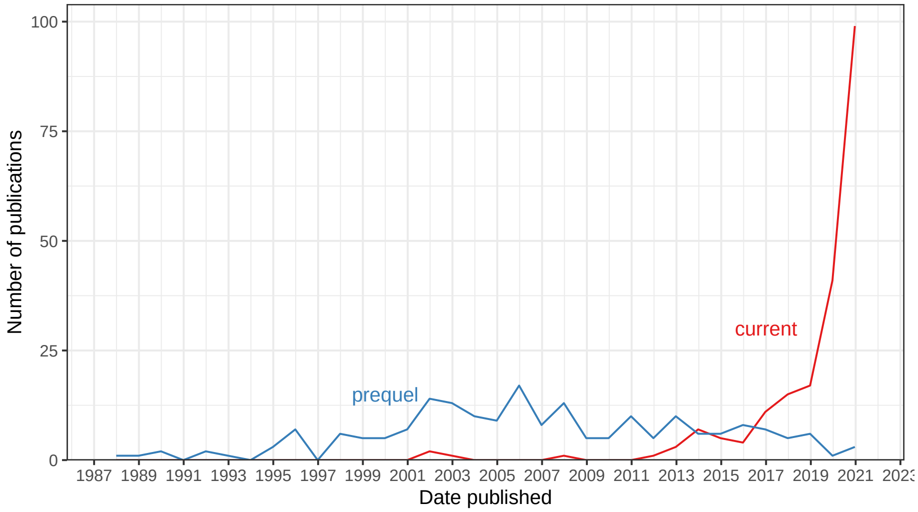

**Figure 4.2:** Comparing number of publications over time in the prequel and the current eras. Bin width is 365 days.

The prequel era started with untargeted screens and grew into atlases and databases striving to be comprehensive. Screens are still a theme in the current era and spatial transcriptomics is still used in untargeted searches for genes involved in development of model organisms, but with highly multiplexed technology, this can also be done for pathological and human tissues (Figure 4.4, Figure 4.5). Thanks to multiplexing, while mouse was the most popular species in the prequel era, in the current era, there are almost as many studies on human tissues as those on mice and the vast majority of studies are on either humans or mice (Figure 4.4). *Drosophila* is no longer as commonly used in the current era (Figure 4.4), perhaps because any current era technique requiring tissue sectioning is less amenable to *Drosophila* embryos, making Tomo-seq along one body axis the only technique that has been demonstrated to be amenable (Combs and Eisen 2013; Combs and Fraser 2018). Whole mount smFISH protocols exist for *Drosophila* brains (Long et al. 2017), zebrafish embryos (Oka and Sato 2015),

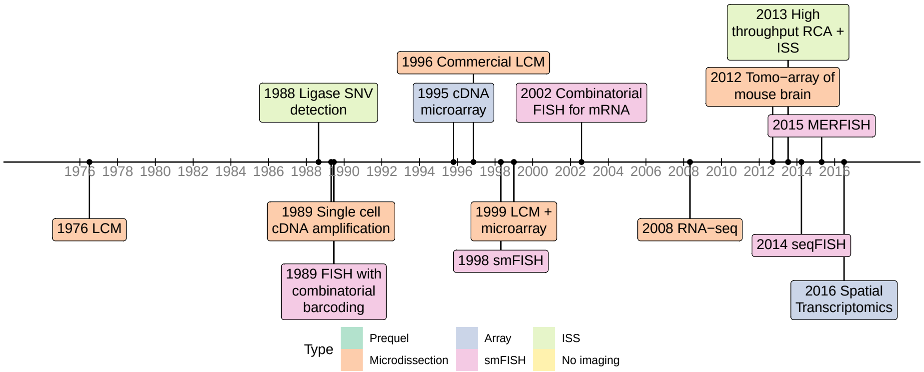

**Figure 4.3:** Timeline of major techniques related to the current era.

and embryonic mouse organs (Wang et al. 2019), but to our best knowledge, highly multiplexed smFISH and ISS have not been adapted to whole mount samples, although they have been adapted to thick slices of cleared tissue (X. Wang et al. 2018; Wang et al. 2021). For *Drosophila* tissue sections, while microdissection, smFISH, and ISS may be applied, the resolution of ST and Visium may be too low to discern sufficiently fine patterns in such a small model organism.

While atlases have so far not been in the center of the stage, some atlases have been made with current era technology, such as MERFISH (Zhang et al. 2020), HybISS (Manno et al. 2020), and ST (Ortiz et al. 2020), described with similar language to that of (WM)ISH atlases. Also as in the prequel era, the brain is still the most favored organ (Figure 4.5, Figure 4.6). Among pathological tissues, breast tumors are the most used (Figure 4.5). More recently, in the wake of the SARS-CoV-2 pandemic, a number of studies using GeoMX Digital Spatial Profiler (DSP) to profile spatial transcriptomes of lungs of COVID victims have been published (Park et al. 2021; Butler et al. 2021; Delorey et al. 2021; Margaroli et al. 2021).

However, unlike in the prequel era, in which older technologies were adapted to larger scale to produce the screens and atlases, the current era has another major theme – new techniques, due to the challenges to be discussed in the following sections; the number of new techniques published each year has grown steadily in the past few years (Figure 4.7). However, this difference might be due to bias in curation and change in culture. In the prequel era, very different enhancer and gene trap vectors were lumped together into enhancer or gene trap in our database, and there might have been many different early non-radioactive ISH protocols not included in our database because they were not used to profile a sufficiently large number of genes. Furthermore, in the current era, authors like to give techniques new names, making related techniques seem distinct rather than lumped together in a wider category like enhancer or gene trap. While a few techniques other than LCM have become somewhat popular,

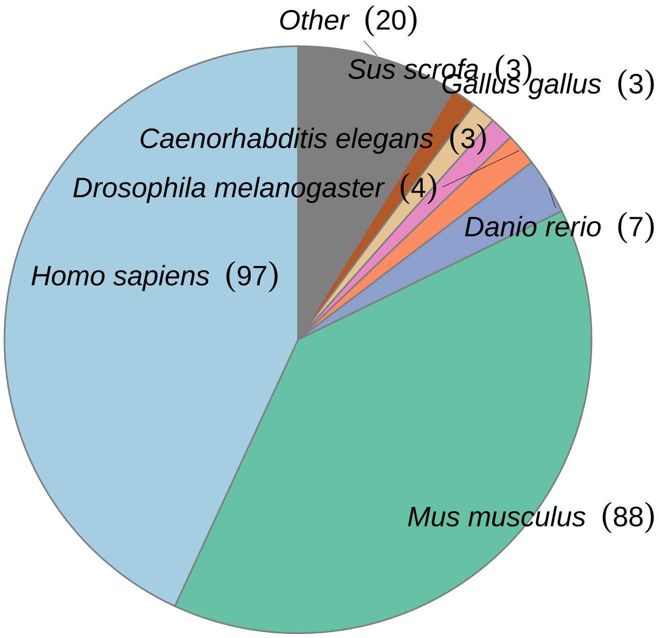

**Figure 4.4:** Number of publication per species.

such as ISS (2013), Tomo-seq (2013), MERFISH (2015), ST (late 2016), GeoMX DSP (2019), and Visium (first preprint in 2020), most techniques never or rarely spread beyond their institutions of origin (Figure 4.7).

Furthermore, especially in the US, research in the current era tends to be more concentrated in a few elite institutions, while research in the prequel era tends to be more spread out to some less well-known institutions (Figure 4.9). Among the top contributing institutions in the prequel era are those hosting databases, such as Allen Institute for ABA, University of Oregon (UO) for ZFIN, UC Berkeley and Lawrence Berkeley National Laboratory (LBL) for BDGP, University of Arizona (UofA) for GEISHA, Jackson Laboratory (JAX) for GXD, Western General Hospital (WGH) for EMAGE, and Kyoto University (Kyodai) for GHOST (Figure 4.9).

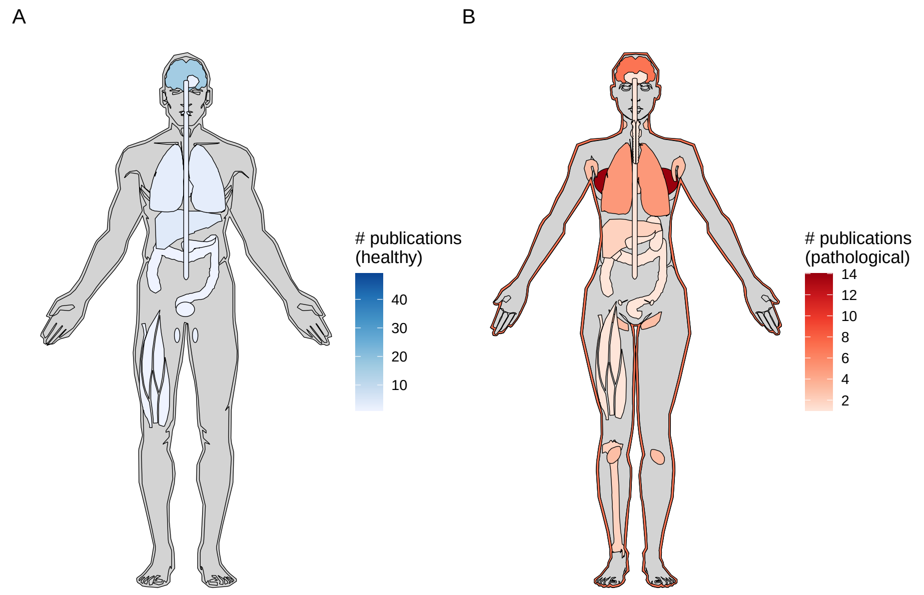

**Figure 4.5:** A) Number of publications for each healthy organ in human (male shown here, as there is no study on healthy female specific organs in humans at present). B) Number of publications for pathological organs in human (female shown here, but there are two studies on prostate cancer (Burgess 2019; Brady et al. 2021)).

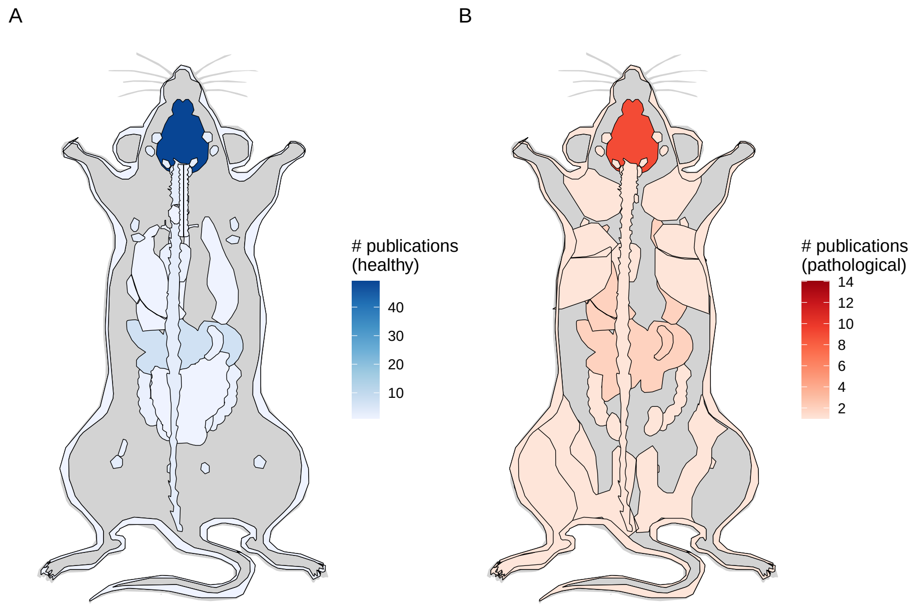

**Figure 4.6:** A) Number of publications per healthy organ in the mouse. B) Number of publications for pathological organs in mouse.

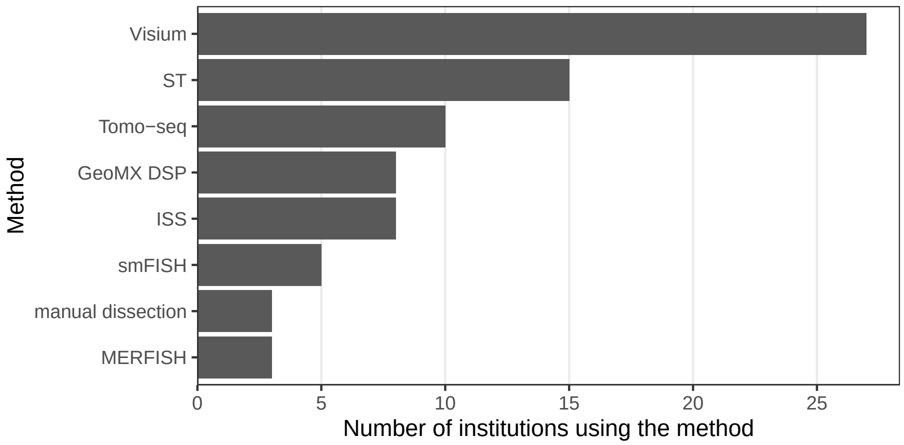

**Figure 4.7:** Techniques used by at least 3 institutions and the number of institutions that have used them.

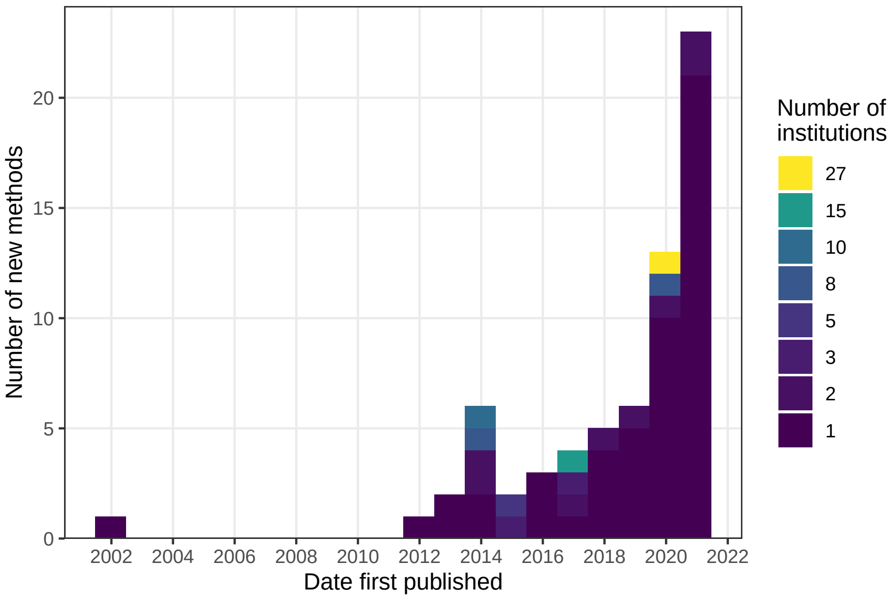

**Figure 4.8:** Number of new methods per year, colored by the number of institutions that have used the method.

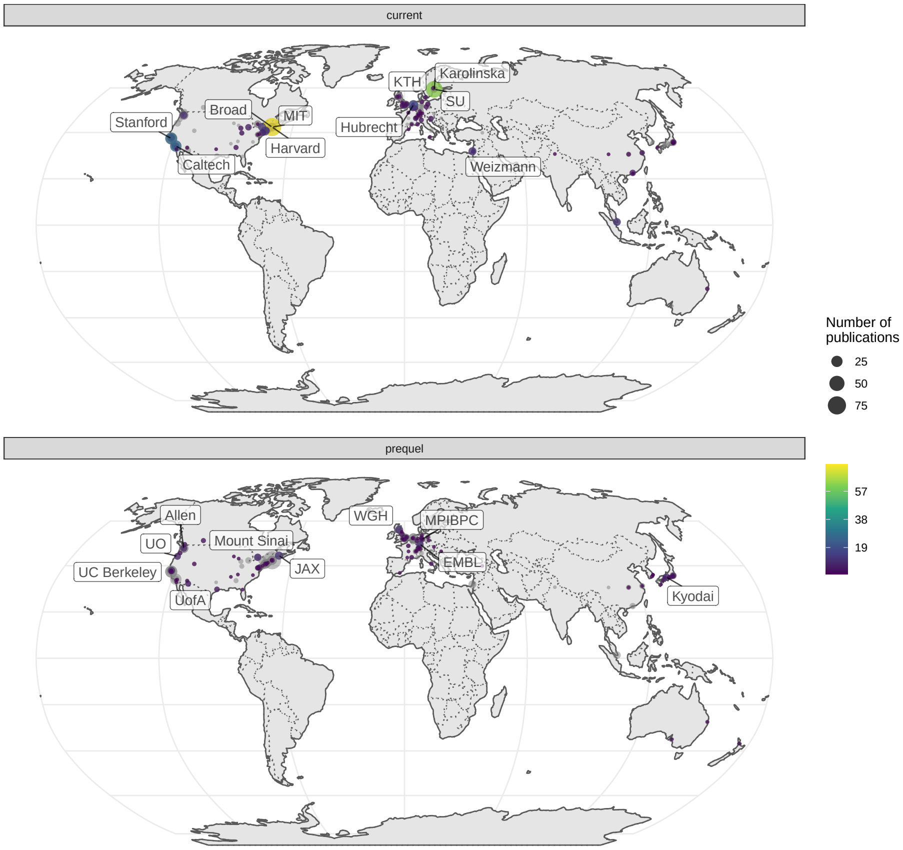

**Figure 4.9:** World map of institutions. Area of the point is proportional to the number of publications from that city. Gray points are sum of both prequel and current eras for each city. Top 10 institutions in each era are labeled.

### Chapter 5

### Current era technologies

#### 5.1 Microdissection

A simple way to preserve spatial information is to isolate the samples from known locations in the tissue. The samples can be isolated physically or by molecular techniques. The known locations can be targeted, for cells with certain histological characteristics, or untargeted, on a grid over the tissue.

##### 5.1.1 History of LCM

###### 5.1.1.1 Microdissection

LCM, also known as laser microdissection (LMD), is by far the most commonly used method of microdissection. Before LCM, manual microdissection could isolate small pieces of tissue, but the process was laborious (Bidarimath, Edwards, and Tayade 2015). Laser microdissection predates ISH, though it was not used for spatial transcriptomics until it was possible to profile the transcriptome from small quantity of tissue.

A precursor to laser microdissection is the 1912 "Strahlenstich", which focused a conventional light source to a spot a few micrometers in size to cut tissues (Greulich 1999). Soon after the invention of the laser in 1960, ruby laser was used to manipulate mitochondria, and a ruby laser microdissection system was commercialized by Zeiss in 1965 (Greulich 1999). UV laser was used to create chromosomal lesions in 1969 (Berns, Olson, and Rounds 1969). The first use of UV laser to cut tissue was in 1976 (Meier-Ruge et al. 1976) (Figure 4.3).

At present, there are two main types of LCM: IR and UV. IR LCM was introduced in 1996 (Emmert-Buck et al. 1996). It utilizes a cap with thermoplastic film which is placed over area of interest, and an IR laser to briefly heat select

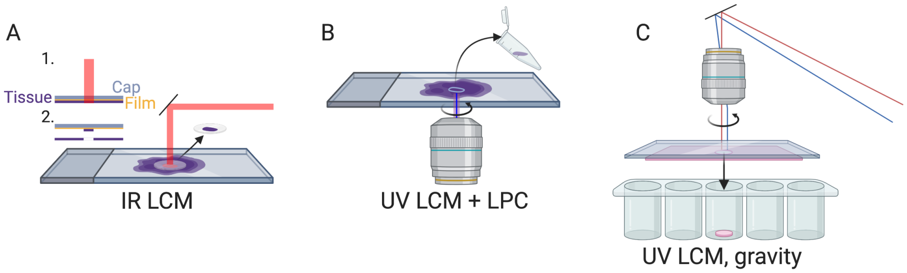

**Figure 5.1:** A) IR LCM schematic. B) UV LCM and LPC schematic, like in Zeiss PALM Microbeam. C) UV LCM, letting microdissected region fall by gravity, like in Leica LMD. All schematics in this book, i.e. anything not made with `ggplot2`, were created with BioRender.com

areas of tissues to 90 °C so the film melts in the area and fuses to the area of tissue of interest (Emmert-Buck et al. 1996) (Figure 5.1 A). This was commercialized as the Arcturus PixCell II LCM System in 1997, which was used in several early LCM studies including the first one in 1999 (Luo et al. 1999; Ohyama et al. 2000; Sgroi et al. 1999; Kitahara et al. 2001) (Figure 4.3).

UV LCM is also known as laser microbeam microdissection (LMM) due to the microbeam of UV laser used. A popular commercial UV LCM system is the Robot-Microbeam (P.A.L.M. Wolfratshausen, Germany), now Zeiss PALM Microbeam. In this method, a narrow UV laser beam ablates a narrow strip of tissue surrounding the area of interest, isolating the area of interest from the rest of the section, so the area of interest is minimally heated. Then, the area of interest is removed from the slide into the collection vial with laser pressure catapult (LPC), avoiding physical contact so as to prevent cross contamination (Figure 5.1 B). An early version of this system was first used in 1996 to isolate single cells from gastric tumors, followed by PCR to analyze E-cadherin mutations, but the cells were removed with a needle rather than LPC (Becker et al. 1996). Another popular commercial UV LCM system is the Leica LMD; unlike the PALM system, the Leica system lets the isolated tissue fall into collection vials by gravity, still avoiding physical contact (Figure 5.1 C). UV LCM was used in some early LCM spatial transcriptomics studies as well (Nakamura et al. 2004), and remains popular in recent years while IR LCM seems to have fallen out of favor (Moor et al. 2017; Zechel et al. 2014; Baccin et al. 2020).

Recent versions of the Arcturus LCM system have both IR and UV, which can be used in conjunction. UV can be used to cut the region of interest (ROI) and IR can then be used to fuse the region to the film at a few points for removal ("Arcturus XT Laser Capture Microdissection (LCM) Instrument - US," n.d.).

###### 5.1.1.2 Amplification

The minuscule amount of transcripts from microdissected tissues, which can be single cells, needs to be amplified to be detected by microarray or RNA-seq. Indeed, RNA amplification is a part of one of the most prevalent topics in LCM related search results (Figures 6.3, 6.4). To this day, there are two main strategies of amplification of minuscule amount of mRNA or cDNA – *in vitro* transcription (IVT) of cDNA (linear amplification) and PCR (exponential amplification), or a combination of both (Tang, Lao, and Surani 2011). These two strategies have coexisted since their beginnings in 1989 and 1990 (Figure 4.3).

Heterogeneous cDNAs can be amplified with PCR by appending known sequences to one or both ends of the cDNA so primers with known sequences can be used to amplify the heterogeneous cDNAs. Early approaches meant for single cells or small number of cells include tailing the cDNA with poly-dA (Belyavsky, Vinogradova, and Rajewsky 1989) or poly-dG (Brady, Barbara, and Iscove 1990) after reverse transcription, and use as PCR primers sequences containing poly-dT (both poly-dA tail and reverse transcription (RT) primer of poly-A mRNAs) or poly-dC (poly-dG tail) and poly-dT (RT primer). Alternatively, lone linkers ("lone" because they are designed to prevent linker polymerization) could be ligated to both ends of the DNA fragments of interest to anneal to PCR primers (Ko et al. 1990). Some of the early single cell (or small number of cells) transcriptomic studies used PCR amplification, prior to quantification or differential expression analayses with Southern blot with radiolabeled cDNA probes hybridizing to cDNA clones of interest screened from plaque lift hybridization of a phage cDNA library (Dulac and Axel 1995), or with cDNA microarray (Klein et al. 2002; Tietjen et al. 2003). LCM was used to isolate the single cells in (Tietjen et al. 2003). Before the advent of CEL-seq, early scRNA-seq methods also used PCR amplification (Tang et al. 2009; Islam et al. 2011). An influential method is switching mechanism at the 5 end of the RNA transcript (SMART) (Zhu et al. 2001), for construction of cDNA (clone) libraries covering the full length of mRNAs, though not originally for single cells. The full length scRNA-seq method Smart-seq(2) (Ramsköld et al. 2012) is based on SMART but adapted to the minuscule amount of transcripts from single cells, with PCR amplification of the cDNA. Smart-seq(2) is one of the most commonly used library preparation methods for LCM since the 2010s, and was used for RNA-seq of LCM isolated single cells (Nichterwitz et al. 2016).

Alternatively, transcripts can be amplified by IVT, with a T7 RNA polymerase promoter attached to the 5' end of the poly-dT primer, so the RNA polymerase transcribes the cDNAs into many copies of antisense RNAs (aRNA) (Van Gelder et al. 1990; Eberwine et al. 1992). Some of the early single cell (or small number of cells) transcriptomic studies used IVT amplification. Quantification and differential expression analyses of the aRNA can be performed with differential display (Liang and Pardee 1992; Kacharmina, Crino, and Eberwine 1999), cDNA microarray (Hemby et al. 2002; Kamme et al. 2003), or with "expression profiling" (Eberwine et al. 1992; Kacharmina, Crino, and Eberwine 1999), i.e. reverse northern blot with radiolabeled aRNAs hybridizing to cDNA clones of interest, where the cDNA clones can be blotted onto a Southern blot membrane in a macroarray, which may have inspired the development of the cDNA microarray printed on glass (Kacharmina, Crino, and Eberwine 1999). LCM was used to isolate the single cells in (Kamme et al. 2003). Since the 2010s, Cell Expression by Linear amplification and Sequencing (CEL-seq) (Hashimshony et al. 2012) and derivatives (e.g. CEL-seq2, MARS-seq, and SORT-seq), which use IVT amplification, have been commonly used for library preparation for microdissected or *de facto* microdissected samples such as from LCM (Tzur et al. 2018), Tomo-seq (Junker et al. 2014), and Niche-seq (Medaglia et al. 2017).

##### 5.1.2 Usage of LCM

Usage trends of LCM as reflected in PubMed and bioRxiv search results are analyzed in Chapter 6. LCM can be used to isolate targeted ROIs based on histology, or to create a grid for untargeted search of gene expression patterns in space, and examples of both are highlighted here. Moreover, the three themes of screening, atlas curation, and new technique development, are all represented in LCM literature. In the "screening" theme, LCM is used to isolate cell populations of interest based on histology (targeted) to discover genes associated with pathological conditions such as cancer metastasis (Nakamura et al. 2004) and cell types (Aguila et al. 2018), or to discover cell type localization in healthy tissue difficult to other spatial transcriptomics techniques such as the bone marrow (Baccin et al. 2020).

LCM can also be used to dissect the tissue in a grid, not targeting very specific histological regions (untargeted), to identify genes associated with locations on the grid (Moor et al. 2018; Peng et al. 2016) or transcriptomically defined regions (Zechel et al. 2014; Peng et al. 2016), or to map cells from scRNA-seq to spatial locations (Zechel et al. 2014; Peng et al. 2016). The untargeted studies can also touch upon the "atlas" theme, providing an online interface to query and explore the spatial transcriptomes (Peng et al. 2016).

However, targeted approaches can also be used for the "atlas" theme, such as in the human (Hawrylycz et al. 2012; Miller et al. 2014) and macaque (Bakken et al. 2016) atlases of the ABA, isolating histologically annotated regions for microarray profiling to build systematic resources for exploration. This addresses the limitation of bright field ISH that only one gene can be stained per section thus requiring large number of brains, which is too costly for primates; in LCM, while often not single cell resolution, the same brain can be used to profile the whole transcriptome. The "technique development" theme is evident in the text mining results (Figure 6.3), and contributes to some of the advantages of LCM as discussed below.

As shown in Chapter 6, LCM transcriptomics has spread far and wide, and has been used on many research topics rarely featured in (WM)ISH atlases. These include cancer and botany (Figure 6.2, Figure 6.3). The following advantages of LCM might have contributed to its popularization: first, as already mentioned, both IR and UV LCM systems have been commercialized prior to their use for transcriptomics, making setup convenient. Second, while LCM equipment can be expensive and require specialized training to use, many institutions have core facilities that can perform LCM ("Translational Pathology Core Laboratory (TPCL)" n.d.; "Veritas Laser Capture Microdissection (LCM) and Laser Cutting System from Applied Biosystems" n.d.; "Dana-Farber Core Facilities" n.d.; "Johns Hopkins Cell Imaging Core Facility" n.d.), reducing cost and personnel training time in individual laboratories.

Third, in some cases, especially in the clinical setting, only archival formalin fixed, paraffin embedded (FFPE) tissues are available. While in 2020, newer current era technologies such as Visium (Villacampa et al. 2020) and GeoMX DSP (Hwang et al. 2020) have been demonstrated on FFPE tissues, LCM followed by microarray was already demonstrated on FFPE tissues in 2007 (Coudry et al. 2007) and with RNA-seq by 2014 (Morton et al. 2014). As a result, for several years, LCM may have been the only option to perform spatial transcriptomics on FFPE samples. In addition, LCM might still be the only way to profile transcriptomes of single cells in FFPE samples. With scRNA-seq library preparation methods such as Smart-seq2 (Nichterwitz et al. 2016), and CEL-seq (Tzur et al. 2018) it is possible to profile the transcriptome in minuscule amount of LCM isolated tissue, and even single cells (Nichterwitz et al. 2016). More recently, with Smart-3SEQ, LCM single cell transcriptomics has been made possible for FFPE tissues as well, even for samples that are several years old (Foley et al. 2019).

Finally, despite its long history, LCM cannot yet be replaced by newer spatial transcriptomics technologies. Unlike smFISH or ISS based techniques, LCM followed by RNA-seq is not restricted to known genes and allows for transcriptome wide profiling and other omics. Unlike ST and Visium, LCM can have single cell resolution, and unlike array based techniques with resolution of the size of a cell or higher, such as Slide-seq(2) and HDST, LCM can more unequivocally isolate individual cells or nuclei based on histology.

LCM has a number of disadvantages, some of which are addressed by other current era spatial transcriptomics technologies. First, compared to droplet based scRNA-seq and highly multiplexed barcoding, using LCM to isolate single cells is still too laborious, limiting its throughput. Second, LCM requires tissue sections, while preparation of many slides to cover a 3D volume can be laborious and it can be challenging to reconstruct 3D structures from tissue sections. To reiterate, sections of blastoderm stage embryos are hard to interpret, which motivated WMISH. Third, because it can be challenging to segment cells based on hematoxylin and eosin (H&E) or immunohistochemistry (IHC) staining and parts of different cells can be stacked within the thickness of the section even in thin sections, single cells isolated by LCM can have contents of other cells. Fourth, LCM can reduce RNA quality in cells (Kerman et al. 2006), perhaps due to heat and UV.

##### 5.1.3 Physical microdissection

###### 5.1.3.1 Voxelation

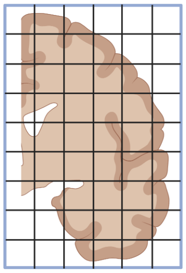

**Figure 5.2:** Voxelation of human brain, as in (Vanessa M. Brown 2002).

LCM did not completely replace microdissection with a physical blade. Voxelation was one of the alternatives to LCM developed to profile spatial transcriptomes in 3D and address the limitation of throughput of ISH. In voxelation, a grid of steel blades is used to cut tissue into cubes for microarray profiling, but the resolution is low. Human brains were first cut into 8 mm thick slabs and then a grid of 1 cm per side (Vanessa M. Brown 2002; Singh et al. 2003), and mouse brains were first cut into 1 mm thick slabs and then a grid of 1 mm per side (V. M. Brown et al. 2002; Singh et al. 2003; Chin et al. 2007) (Figure 5.2). With low resolution, it's easier to use voxelation to profile large 3D tissues of multiple slabs that would be much more laborious with LCM's thinner sections and higher resolution (V. M. Brown et al. 2002). As the human voxels were quite large (almost 1 ml) and corresponding voxels of 20 to 30 mice were pooled (V. M. Brown et al. 2002; Chin et al. 2007) to get enough transcripts, the voxelation studies did not mention T7-based PCR amplification of transcripts, unlike for LCM samples (Nakamura et al. 2004). To the best of our knowledge, voxelation never spread beyond its institution of origin, UCLA School of Medicine, and has not been used in a publication to generate new data since 2007 (Chin et al. 2007) and for data analysis since 2009 (An et al. 2009).

###### 5.1.3.2 Tomo-seq

Another alternative to LCM is Tomo-seq/array, which has continued to be utilized in recent years. In this approach, the tissue is sectioned with a cryotome like tomography (hence the "Tomo"), and the transcripts in each section are extracted for microarray (Tomo-array) or RNA-seq (Tomo-seq) profiling; the

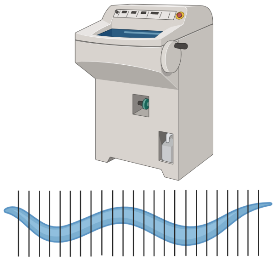

**Figure 5.3:** Tomo-seq, here showing C. elegans.

resolution is limited by section thickness, which has gone down to 8 $\mu$m (Brink et al. 2020). Three-D expression maps can be reconstructed from sections along the anterior-posterior (AP), dorsal-ventral (DV), and left-right (LR) axes. All three themes, namely screening, atlas cutation, and new technique development, are present in Tomo-seq/array literature.

Tomo-array was first used in 2012 to build a 3D mouse brain transcriptome atlas, attempting to address difficulties in image registration in ISH atlases, low resolution of voxelation, and limitation of LCM to specific regions (Okamura-Oho et al. 2012) (Figure 4.3). Mouse brains were sectioned along all three axes and 200 adjacent 5 $\mu$m sections were pooled as "fractions" for microarray; again, PCR amplification was not mentioned. Fractions from the three axes were then used to reconstruct a 3D atlas.

Tomo-seq was first demonstrated in 2013, on *Drosophila melanogaster* embryos, with 60 and 25 $\mu$m sections, again in response to the difficulty to scale ISH atlases to the whole transcriptome (Combs and Eisen 2013). Genes patterned along the AP axis were identified, and the data is stored in an online database[1]. However, Tomo-seq is more commonly credited to a 2014 method first demonstrated on zebrafish embryos, with 18 $\mu$m sections (Junker et al. 2014). Gene expression patterns along the AP axis of straightened embryos were identified, and sections along all three axes were used for 3D reconstruction of embryos that were not straightened. The data and the 3D reconstruction are also stored in an online database[2], though the 3D reconstruction algorithm produced many artefacts.

Since then, Tomo-seq has been used in several different biological systems, typically when one axis is of primary interest. Tomo-seq has been used in *C. elegans* (Ebbing et al. 2018), developing zebrafish hearts (Burkhard and Bakkers 2018), *Drosophila* embryos (Combs and Fraser 2018), ischemic mouse hearts (Lacraz et al. 2017), and *Pristionchus pacificus*[3] (Rödelsperger et al. 2020) to identify genes associated with that axis of interest. Tomo-seq was also used on mouse

[1] http://eisenlab.org/sliceseq/
[2] http://zebrafish.genomes.nl/tomoseq/
[3] http://wormbook.org/chapters/www_genomesPristionchus/genomesPristionchus.html

(Brink et al. 2020) and human (Moris et al. 2020) gastruloids to demonstrate the viability of this in vitro and potentially high-throughput model for developmental biology. Again, due to the minuscule amount of tissue in each section, library preparation methods designed for scRNA-seq, such as CEL-seq(2) (Junker et al. 2014; Rödelsperger et al. 2020; Ebbing et al. 2018) have been adapted to Tomo-seq.

###### 5.1.3.3 Other methods of physical microdissection

More recently, Spatial Transcriptomics by Reoriented Projections and sequencing (STRP-seq) was developed in response to the limited number of genes of smFISH and ISS based techniques, degradation of RNA and technical complexity of LCM, and number of specimens required by and inadequacy of the 2014 Tomo-seq 3D reconstruction (Schede et al. 2020). In STRP-seq, adjacent sections of the tissue are sectioned in different orientations, and are then used for 3D construction with an algorithm inspired by reconstruction of ray-based computerized tomography. This has been shown to perform better than the 2014 Tomo-seq 3D reconstruction method, and was demonstrated on the brain of a non-model organism, the lizard *Pogona vitticeps*[4].

Because of the specialized equipment and technical complexity of LCM and degradation of RNA, other methods of physical microdissection have been developed. Examples of such techniques are Cell and Tissue Acquisition System (CTAS), which uses a disposable capillary unit connect to the vacuum to aspirate tissue (Kudo et al. 2012), and an automated micropuncch system that collects samples of tissue with diameter of 110 $\mu$m at 300 $\mu$m intervals (Yoda et al. 2017). In addition, for similar reasons, manual microdissection is still used (Figure 4.7), such as to dissect leaves on a grid of distances from a lesion to characterize response to infection (Giolai et al. 2019; Lukan et al. 2020). Manual microdissection of pre-defined anatomical regions was also used to create low resolution gene expression atlases of *Xenopus laevis* (Plouhinec et al. 2017) and *Xenopus tropicalis* (Blitz et al. 2017) embryos, to avoid sectioning as required for LCM and artefacts in Tomo-seq 3D reconstruction.

##### 5.1.4 *De facto* microdissection

Some methods have been developed that do not directly cut tissues. Instead, cells, or ROIs judged from histology, are optically and molecularly marked so that only transcripts or cells from the marked regions are captured. Because these methods involve selection of pre-defined ROIs within the section, we call them *de facto* microdissection.

Transcriptome *in vivo* analysis (TIVA) from 2014 can be viewed as the first of these methods (Lovatt et al. 2014). Live cell culture is incubated with the

[4] `https://en.wikipedia.org/wiki/Pogona_vitticeps`

photoactivable cage with a poly-U sequence that captures poly-A transcripts. Select cells are photoactivated by 405 nm laser and the captured transcripts are sequenced. TIVA is widely cited, perhaps because it is one of the earliest single cell resolution and transcriptome wide methods, predating RNA-seq from LCM isolated single cells. However, because TIVA has only been demonstrated on fewer than a dozen cells per sample, to the best of our knowledge it has not been used in any other publication to collect new data.

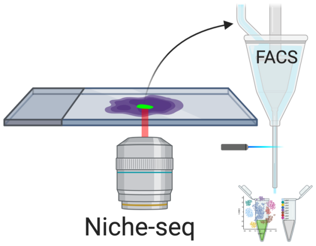

**Figure 5.4:** Niche-seq schematics. Green: cells with photoactivated PA-GFP.

Two *de facto* microdissection methods have spread beyond their institutions of origin. One of them is Niche-seq, which was developed as LCM is still usually used to isolate groups of cells rather than single cells and involves tissue fixation (Medaglia et al. 2017). Select regions of *ex vivo* tissues from transgenic mice expressing photoactivable GFP (PA-GFP), here lymph node and spleen B cell and T cell niches, are photoactivated at 820 nm with two photon irradiation. Then the tissue is dissociated and cells with photoactivated PA-GFP are collected from flow cytometry–based fluorescence-activated cell sorting (FACS) for scRNA-seq with MARS-seq (Figure 5.4). After its inception, Niche-seq has been used once more in lymph node niches (De Giovanni et al. 2020). However, as Niche-seq requires transgenic mice expressing PA-GFP and living tissue, it cannot be applied to human tissues, to fixed tissues, or when a PA-GFP line is unavailable. This might limit further growth of Niche-seq. Moreover, the spatial context of cells within the photoactivated region is lost, limiting spatial resolution.

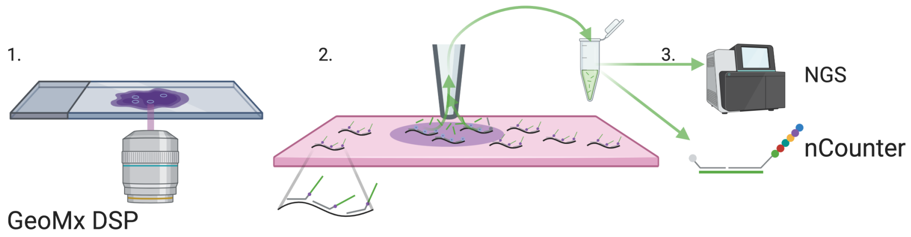

**Figure 5.5:** GeoMX DSP schematics, inspired by figures in (Merritt et al. 2019). Black: transcripts in tissue. Gray: probes. Green: indexing oligo.

Another method that spread beyond its institution of origin is the commercial GeoMX DSP from NanoString (Merritt et al. 2019), which can be used for both high throughput immunofluorescence and transcript quantification in FFPE tissue sections. For transcript quantification, probes are attached to indexing oligos with a UV cleavable linker (Figure 5.5). The selected ROI is illuminated by UV to remove the index oligos from the probes. Then the released index oligos are aspirated and quantified with either NGS or NanoString nCounter. This can be repeated for multiple ROIs, which can be a grid for unbiased profiling (Merritt et al. 2019). The probes tile the transcripts, and each probe has a distinct index oligo, so in NGS, each tile is counted separately, enabling isoform quantification (Merritt et al. 2019). GeoMx DSP is not transcriptome wide; up to 1860 target transcripts have been quantified (Margaroli et al. 2021). Also, as pre-defined probes are required, unlike in RNA-seq, novel transcripts cannot be quantified. Ready made probe sets for oncology, immunology, and neuroscience are sold by NanoString ("Gene Expression Panels | NanoString Technologies" n.d.). Although GeoMx DSP was published in 2019, it has spread to several different institutionss, and has been used on pancreatic ductal adenocarcinoma (PDAC) (Hwang et al. 2020), hepatocellular carcinoma (HCC) (Sharma et al. 2020), reactive lymph nodes (Tripodo et al. 2020), and COVID infected lungs from autopsy (Park et al. 2021; Butler et al. 2021; Delorey et al. 2021; Margaroli et al. 2021). A variant of GeoMX DSP, called GeoMX Whole Transcriptome Atlas (WTA), has been used to profile transcripts of 18190 genes, nearly covering the whole transcriptome (Roberts et al. 2021). In GeoMX WTA, the UV cleaved index oligo must be sequenced with NGS to identify the gene each transcript is from.

##### 5.1.5 Summary

Overall, in the current era, microdissection is the most widely used type of technique (Figure 5.6). Excluding LCM, Tomo-seq is the most popular technique after ST and Visium (Figure 4.7). Microdissection has not been replaced by other seemingly more sophisticated techniques such as ST and MERFISH, and is still popular in 2020 and 2021 (Figure 4.1, Figure 6.1). Microdissection techniques generally do not have single cell resolution, but combined with scRNA-seq or snRNA-seq data, cell type compositions of ROIs can be computationally deconvoluted (Baccin et al. 2020; Hwang et al. 2020). The popularity may be due to availability of commercial platforms (LCM and GeoMX DSP), core facilities (LCM), Nanostring's commercial data collection and analysis service for GeoMX DSP ("DSP Technology Access Program (TAP)" n.d.), not requiring specialized equipment (Tomo-seq, manual microdissection), or disadvantages of other techniques discussed later in this chapter.

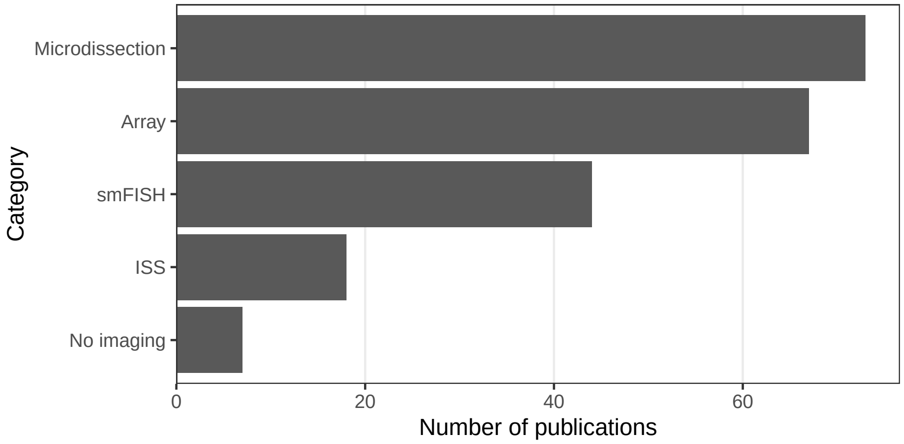

**Figure 5.6:** Number of publications per category of techniques in the current era. Non-curated LCM literature is excluded.

#### 5.2 Single molecular FISH

One quantitative approach to transcripts abundance estimation is to display individual transcripts as distinct puncta with FISH and count them. Prior to smFISH, transmission electron microscopy was used to visualize individual mRNA molecules in fibroblasts by labeling the poly-A tail with a single large colloidal gold particle and the *in situ* reverse transcribed cDNA with small gold particles (Bassell et al. 1994). That FISH can be used to visualize single mRNA molecules was first demonstrated in 1998 (Femino et al. 1998) (Figure 4.3). Five or more probes targeting adjacent parts of the transcript, each about 50 nt long and labeled with 5 fluorophores were hybridized to the transcripts. The puncta seen were shown to be likely individual mRNA molecules, as the fluorescence intensity of each punctum was consistent with the number of fluorophores, and the number of puncta for $\beta$-actin was consistent with the number of $\beta$-actin transcripts measured by other means, and the colors of puncta seen from probes with different colored fluorophores targeting different parts of the transcript were consistent with organization of the fluorophores on the transcript (Figure 5.7).

The 1998 approach had a number of disadvantages, leading to development of an alternative approach in 2008 (Raj et al. 2008). First, probes labeled with multiple fluorophore moieties are difficult to synthesize and purify. Second, the multiple fluorophores on the same probe can interact with each other and self-quench. Third, out of the 5 probes per transcript, only 1 or 2 may have actually hybridize to the transcript in most cases, making it difficult to distinguish between true signal and non-specific binding. In the 2008 method, each 17-22 nt probe is labeled with one fluorophore at the 3' end, and a larger number of probes (48 or more) targeting tandem sequences of the transcript were used

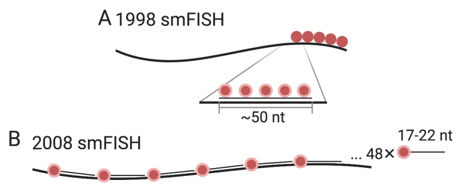

**Figure 5.7:** A) Schematic of smFISH from (Femino et al. 1998). The long thick line stands for the mRNA, and short think line stands for DNA oligo probe. B) smFISH with singly labeled probes from (Raj et al. 2008).

to improve signal to noise ratio (Figure 5.7). The probes were computationally designed and ordered from Biosearch Technologies. This method influenced later highly multiplexed smFISH techniques; computational probe design and commercial synthesis would remain crucial.

##### 5.2.1 Barcoding strategies

To use smFISH to quantify transcripts transcriptome wide, there is an obvious challenge – how to distinguish among over 20,000 genes with only about 5 easily distinguishable colors? Various strategies using multiple colors and/or rounds of hybridization or imaging have been devised to drastically expand the palette. The first attempt to do so was in 1989, using 3 colors to visualize 4 chromosomes in immunological DNA FISH (Nederlof et al. 1990) (Figure 4.3). Each probe can be labeled with one or two of the 3 haptens: biotin, 2-acetyl aminofluorene (AAF), and Chemiprobe. Red fluorophore was attached to avidin to target biotin label, and blue and green to different secondary antibodies targeting, respectively, mouse anti-Chemiprobe and rabbit anti-AAF primary antibodies (Figure 5.8). Then with one doubly labeled and 3 singly labeled probes, imaged with different excitation wavelengths or channels, 3 colors can distinguish 4 chromosomes. However, with this method, the palette size is limited by the number of haptens available and the number of their combinations.

For transcript detection, the first attempt was in 2002 (Levsky 2002); fluorophore labeled probes were synthesized as in the 1998 smFISH method, and either probes of one color or a mixture of probes of 2 colors were hybridized to the transcript, and imaged with different channels, to visualize transcription foci in the nucleus (Figure 5.8). This way, combinations of 2 of the 4 available colors plus blank were used to encode 10 different transcripts.

The above mentioned historical works in smFISH and combinatorial barcoding laid foundation to smFISH based spatial transcriptomics. The first attempt to quantify transcripts with combinatorial barcoding at single molecular resolution was in 2012 by Long Cai's group, which later developed seqFISH and its variants (Lubeck and Cai 2012). Like in the 2008 smFISH study, singly labeled probes

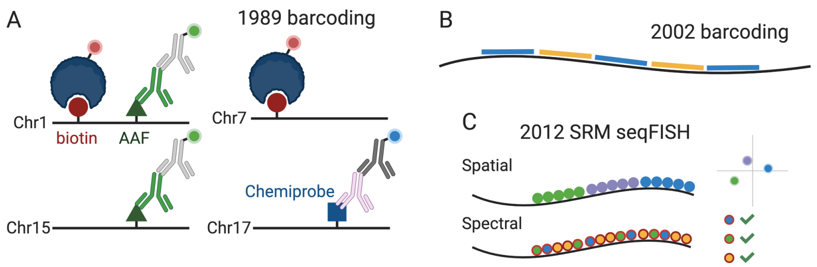

**Figure 5.8:** A) Combinatorial barcoding in immunological DNA FISH, as described in (Nederlof et al. 1990). The line stands for the probe and the circle, triangle, and square stand for haptens. Not to scale, and only one hapten of each kind is shown on one probe. B) Combinatorial barcoding in (Levsky 2002). Short colored lines stand for probes with fluorophores of the color. C) Schematic of SRM seqFISH as described in (Lubeck and Cai 2012).

purchased from Biosearch were used, but forming blocks of different colors as in the 1998 smFISH $\beta$-actin experiment. Then the transcripts were imaged with super-resolution microscopy (SRM), in particular stochastic optical reconstruction microscopy (STORM). In the spatial barcoding strategy, the ordering of the colors in space would distinguish between transcripts, but would require linearization of the transcripts and high resolution (20 nm) (Figure 5.8). To improve signal to noise ratio, cyanine dye–based photoswitchable dye pairs (Bates et al. 2012) was used so both the activator and the emitter fluorophores must be present and adjacent for the fluorophores to be reactivated. In the spectral barcoding approach, the pairs of fluorophores are spread across the transcript, so the transcripts are recognized by the pairs of fluorophores detected (Figure 5.8). The spectral approach requires lower resolution (100 nm) and does not require linearization, but because the ordering of the colors is not used, the number of possible barcode from the same number of colors is smaller than in the spatial approach. With spectral barcoding, transcripts of 32 genes were quantified in yeast, with 3 color barcodes chosen from 7 available colors. To the best of our knowledge, after its inception, this SRM method has not been used to generate new data, perhaps because it requires specialized equipment for SRM. None of the later methods in our curated database used SRM.

Thus far, probes with fluorophores of different colors were hybridized to mRNAs at the same time, without multiple rounds of hybridization. To obtain single molecular resolution but without SRM, there is a challenge of needing to use multiple probes of the same color to strengthen signal, which requires transcripts that are long enough to accommodate probes of different colors. The more colors that are used to encode more genes, the longer the transcripts must be.

This changed in 2014, with the advent of seqFISH (Lubeck et al. 2014). Twenty four singly labeled probes were designed for each gene, and 12 genes were encoded with 4 colors and 2 rounds of hybridization (Figure 5.9). After imaging the first round of hybridization and DAPI staining for DNA, the probes are removed with DNase I, and then probes for the second round are hybridized. Let $F$ denote the number of fluorophores or colors, and $N$ denote the number of rounds of hybridization, then the number of genes that can be barcoded is $F^N$. However, with longer barcodes to encode more genes, error can build up.

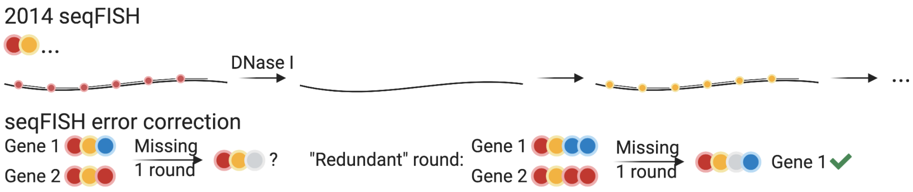

**Figure 5.9:** Probe structures of 2014 seqFISH (Lubeck et al. 2014) and seqFISH error correction.

The most common error in multi-round smFISH is missing signal, most likely in one round (Shah et al. 2016; K. H. Chen et al. 2015). If all $F^N$ barcodes are used and one round is missing for a mRNA molecule, then the existing signal of this molecule is consistent to $F$ genes, so it cannot be uniquely identified. If a small proportion of barcodes are intentionally left out to control for false positives, as done in this first version of seqFISH (4 out of 16), then error correction is still not guaranteed. A further defense against errors in 2014 seqFISH was to repeat the 2 rounds of hybridization 3 times, so 6 rounds were performed. This filtered out false positives where repeated rounds didn't match, and barring false positives, this can recover the original 2 barcoding rounds if up to 2 of the 6 total rounds have missing signal.

Another error correction scheme was introduced in 2016, with hybridization chain reaction (HCR) seqFISH (Shah et al. 2016), and was used in seqFISH+ (Eng et al. 2019) as well. One more round of hybridization than necessary to encode the number of genes of interest was used, and the barcodes are designed so that if one of the rounds is missing, the remaining rounds still uniquely identify the gene (Figure 5.9). For example, with 5 colors, 3 rounds are enough to encode 100 genes, as 125 barcodes are possible. However, a fourth round is used, so missing one round can still result in 3 remaining rounds that uniquely identify the gene.

An alternative to seqFISH was developed with error correction in mind – multiplexed error-robust FISH (MERFISH) (K. H. Chen et al. 2015). In MERFISH each encoding probe has a 30 nt long region that targets the transcript, and 2 or 3 20 nt (Moffitt et al. 2016) readout sequences to bind to readout probes (Figure 5.10). First, the encoding probes are hybridized to the transcripts. For each round of hybridization, readout probes, singly labeled, are hybridized to the readout sequences on the encoding probes and imaged. Then the fluorescence of the previous round is either photobleached (version 1) (K. H. Chen et al. 2015) or when the fluorophore is bound to the readout probe with a disulfide

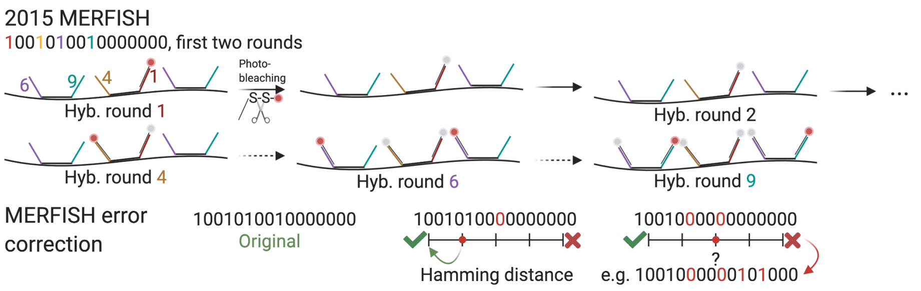

**Figure 5.10:** Schematic of MERFISH (K. H. Chen et al. 2015; Moffitt et al. 2016) and MERFISH error correction.

bond, cleaved off with a reducing agent such as Tris(2-carboxyethyl)phosphine (TCEP) (version 2) (Moffitt et al. 2016). The readout probes are not stripped, and in the next round, new readout probes are hybridized to new readout sequences and imaged.

The MERFISH barcodes are binary, with "1" for a round with fluorescence, and "0" without, and must differ from other barcodes at at least 4 places, i.e. with Hamming distance[5] of at least 4 (HD4). As missing signal is the most common error, each barcode has 4 1's, or Hamming weight 4. This way, when one round is missing, the gene can still be uniquely identified, but when 2 rounds are missing, the remaining barcode is equally distant to 2 genes, so the error cannot be corrected (Figure 5.10). Sixteen rounds of imaging, or 16 bits, would result in 140 barcodes. In this case, there are 16 different readout sequences, and each gene is assigned 4 of them, for the 4 1's in the barcode. If the code is expanded to 69 bits, then about 10,000 genes can be encoded, and imaging time can be cut to a third by using 3 colors to image 3 bits per round (C. Xia, Fan, et al. 2019). An HD2 code, i.e. barcodes are at least hamming distance 2 away from each other, can also be used, but errors can only be recognized but not corrected. All variants of MERFISH use this type of binary barcoding.

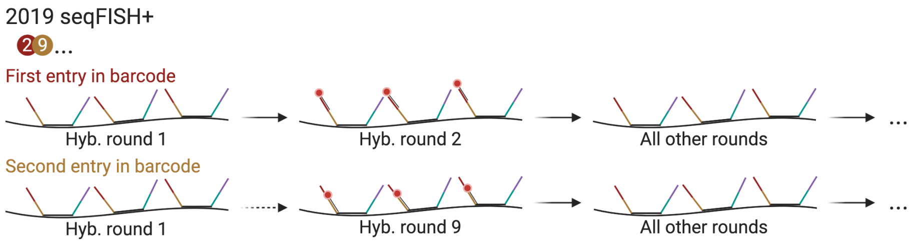

**Figure 5.11:** Schematic of seqFISH with pseudocolors.

More recently, a new variant of seqFISH was devised to scale up to 10,000 genes

[5] https://en.wikipedia.org/wiki/Hamming_distance

(Shah et al. 2018). The barcoding and hybridization scheme enabling such scale was first introduced in vitro in 2017 as RNA SPOTs (Eng et al. 2017), and was then adapted to cultured cells in 2018, targeting introns of nascent transcripts of over 10,000 genes (Shah et al. 2018). In 2019, this scheme was used to profile mature transcripts of 10,000 genes in both cell culture and the mouse brain, and with super-resolution (Eng et al. 2019). Super-resolution beyond the diffraction limit can be achieved by computationally super-resolving the transcript spots with a radial center algorithm (Parthasarathy 2012) when spot density is very high to help with decoding barcodes; the super-resolution version is known as seqFISH+. While this new version of seqFISH can reduce optical crowding and greatly expand the palette, the super-resolution algorithm that can further reduce crowding does not have to be used to locate the transcript spots when density is low. This version of seqFISH was again used to visualize genomic loci (super-resolution) (Takei et al. 2020) and mature transcripts of a smaller number of genes (not super-resolution) (Lohoff et al. 2020).

This method is quite different from previous seqFISH variants, and is in some ways reminiscent of MERFISH. Like previous versions of seqFISH, each barcode is a series of colors, but a large number of "pseudocolors", specifically 20 per channel in the seqFISH+ study, are used rather than the 5 fluorophores, so 3 rounds of hybridization can encode $20^3$ or 8000 genes per channel. Any number of pseudocolors and rounds can be used depending on the number of genes profiled. Each primary probe has a 28 nt region targeting the transcript and 4 readout sites of 15 nt. Each readout site has as many different sequences as there are pseudocolors, and the 4 sites correspond to the series of 4 pseudocolors in the barcode. First, 24 primary probes are hybridized to the transcripts. Then for each place of the barcode, 20 (or whatever number of pseudocolors) rounds of hybridization with readout probes are performed, stripping with formamide between rounds. In these 20 rounds, each gene should light up only once, and its place in the 20 rounds is its pseudocolor (Figure 5.11). This way, in each image, only 1 out of 20 molecules of interest imaged in the channel fluoresce, reducing optical crowding. For the entire barcode of length 4, there would be 80 rounds of hybridization. In contrast, in MERFISH, with the 16 bit barcode, this would be 1 out of 4. Like in MERFISH, a larger number of real colors, or channels, can be used to increase throughput, to image multiple pseudocolors simultaneously. So with 3 channels, 24,000 genes can be encoded. The same error correction method as in HCR seqFISH was used, so while a barcode of length 3 is sufficient, length 4 was used.

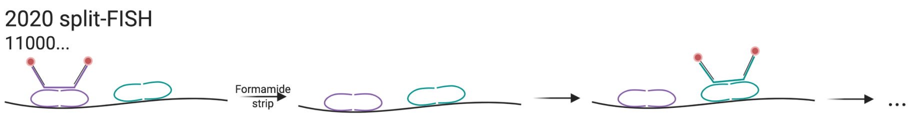

**Figure 5.12:** Schematic of split-FISH.

Another new method, called split-FISH (Goh et al. 2020) was devised to reduce off target hybridization, and thus background noise and some barcoding errors. For each encoding probe or bridge probe like in MERFISH, a pair of split probes hybridize to the transcript itself (Figure 5.12). Half of the split probes would bind to the transcript, and the other half bind to the bridge probe. Then as in MERFISH, the bridge probe has 2 readout sequences and singly labeled readout probes bind to the bridge probe for imaging. This method reduces off target hybridization because the bridge probe can only indirectly bind to the transcript if both of the split probes hybridize to the transcript. To encode 317 genes, 2 places out of 26 in binary barcodes are chosen to be "1", resulting into 325 possible barcodes; 8 of them are left blank to control for false positives. Error correction is not mentioned.

Despite the availability of the above barcoding schemes, when the number of genes stained for is not too large, each gene can still be encoded by only one round of hybridization and one color. When the number of genes is larger than the number of colors, each round of hybridization stains for as many genes as there as colors, and the probes are stripped so the next round stains for a different set of genes. This has been done in osmFISH (Codeluppi et al. 2018) staining for 33 genes, in a non-barcoded adaptation of HCR-seqFISH called Spatial Genomic Analysis (SGA) (Lignell et al. 2017) staining for 35 genes, and in Expansion-Assisted Iterative Fluorescence *In Situ* Hybridization (EASI-FISH) 26 genes (Wang et al. 2021).

##### 5.2.2 Signal amplification

As already mentioned, in smFISH, a large number of singly labeled probes can be used to boost signal, but not all transcripts are long enough to accommodate this number of probes. Furthermore, isoform specific exons are often not long enough to accommodate these probes for isoform specific staining. Without increasing the number of probes, background reduction such as by tissue clearing, split probes (e.g. in split-FISH), and using fluorophores with colors very different from the color of autofluorescence (Moffitt et al. 2016) can increase signal to noise ratio. There are also ways to boost signals without increasing the number of probes, the most common of which are branched DNA (bDNA), rolling circle amplification (RCA), and HCR. All of these methods non-covalently attach numerous fluorophores to the probe to amplify signal. Background reduction and signal amplification can be used in conjunction.

###### 5.2.2.1 Branched DNA

Dating back at least as far back as to 1993 (Urdea 1993), early use of bDNA in ISH was to detect low copy number of viral genomes, eventually down to single copies (Player et al. 2001). bDNA signal amplification involves several steps of hybridization (Figure 5.13). First, usually some sort of bridge probe binds to the transcript itself. Then the primary amplifier binds to the bridge probe, leaving a long overhang. Then multiple secondary amplifiers bind to the primary amplifier on the overhang of the primary amplifier, and each secondary amplifier also leaves an overhang. Finally, multiple labeled readout probes bind to each secondary amplifier. This way, space available for hybridization of the readout probes is drastically expanded, allowing for more fluorophores per unit transcript length.

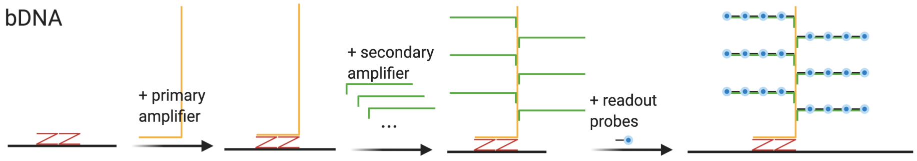

**Figure 5.13:** Schematic of bDNA. The Z probes are specific to RNAscope, but the other parts are generic to bDNA.

For FISH, a particularly influential bDNA method is RNAscope, introduced in 2012 for FFPE tissues, and is now commercially available from ACD (Wang et al. 2012). In addition to bDNA amplification, RNAscope reduces background noise from non-specific hybridization by using 2 bridge Z probes in between the transcript and the primary amplifier, so the primary amplifier will only bind when both Z probes are present. An smFISH RNAscope method has been used to profile around 1000 genes in cell culture (Battich, Stoeger, and Pelkmans 2013) and 49 genes in the mouse somatosensory cortex (Bayraktar et al. 2020), although these experiments were not highly multiplexed and only one or a handful of genes distinguishable by fluorophore color were stained for in the same cells or sections; numerous cells and sections were stained to cover all genes in the gene panels. However, bDNA has also made its way into highly multiplexed smFISH, as a variant of MERFISH (C. Xia, Babcock, et al. 2019). Here, the primary amplifier binds to the readout regions of the MERFISH encoding probe. Like in regular MERFISH (v2), the fluorophores are attached to the readout probes by a disulfide bond and removed by TCEP after each round of hybridization; the bDNA moiety is not removed. With bDNA amplification, only 16 probes per gene can detect about as many transcripts as with 92 unamplified probes (C. Xia, Babcock, et al. 2019).

###### 5.2.2.2 Rolling circle amplification

Chronologically, the next of the popular signal amplification method is padlock probe RCA. Padlock probe was introduced in 1994 by Mats Nilsson as a way to reduce background in ISH and to detect single nucleotide variants (SNVs) (Nilsson et al. 1994). Both ends of of the padlock probe must hybridize to the target without terminal mismatches for the ligase to connect the ends of the probe to form a circle (Figure 5.14); thus padlock probe and RCA can detect

SNPs and point mutations (Larsson et al. 2010; Lizardi et al. 1998). The circle encloses the target like a padlock on a string, hence the name “padlock probe”. Then probes that are not circularized are digested by an exonuclease. RCA was introduced in 1995 as a way to create tandem repeats and potentially point to origins of tandem repeats in genomes, not seeming to have signal amplification in mind (Fire and Xu 1995). A primer anneals to circularized DNA and is then elongated by Φ29 DNA polymerase, and as the polymerase goes around the circle many times, many copies of the complimentary sequences of the circle are made (Figure 5.14). In 1998, padlock probes and RCA were united to create a method of signal amplification (Baner et al. 1998; Lizardi et al. 1998).

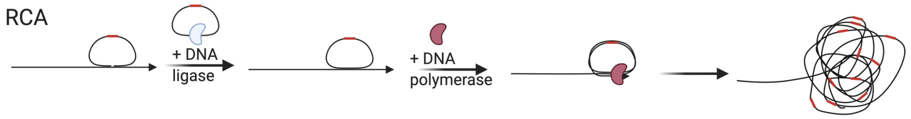

**Figure 5.14:** Schematic of RCA, here shown with target priming though a separate primer can also be used. Red segment is the gene barcode.

In spatial transcriptomics, padlock probe and RCA were initially used for *in situ* sequencing (ISS) (Ke et al. 2013), but more recently adapted to smFISH. The padlock probe with the gene barcode is hybridized to *in situ* reverse transcribed cDNA as in ISS and hybridization-based ISS (HybISS) (Gyllborg et al. 2020), or the mRNA itself as in SCRINSHOT (Sountoulidis et al. 2020) and hybridization-based RNA ISS (HybRISS) (Lee et al. 2020). RCA can be initiated with the target cDNA itself as a primer or with a separate primer when the target is mRNA. Then readout probes are hybridized to the RCA amplified gene barcode, with (Gyllborg et al. 2020) or without (Sountoulidis et al. 2020) a bridge probe. In Hyb(R)ISS and SCRINSHOT, multiple rounds of readout hybridization encode each gene with a sequence of colors as in seqFISH; although error correction is not discussed, the seqFISH error correction scheme can be easily adapted. Perhaps because of larger number of copies of the gene barcode sequence produced by RCA, Hyb(R)ISS and SCRINSHOT use 5 probes per gene, each with a 30 nt (HybISS, target sequences are proprietary information of CARTANA for HybRISS) or 40 nt (SCRINSHOT) region to target the transcript. While we are unaware of isoform specific studies conducted with Hyb(R)ISS or SCRINSHOT, isoform specific exons may more realistically accommodate the 5 probes. While smFISH based techniques were typically designed for frozen sections, SCRINSHOT was designed for FFPE sections.

###### 5.2.2.3 Hybridization chain reaction

A third signal amplification method is HCR, introduced in 2004 (Dirks and Pierce 2004), which has been adapted to seqFISH, giving rise to HCR-seqFISH. EASI-FISH also uses HCR for signal amplification. In singly labeled hairpins,

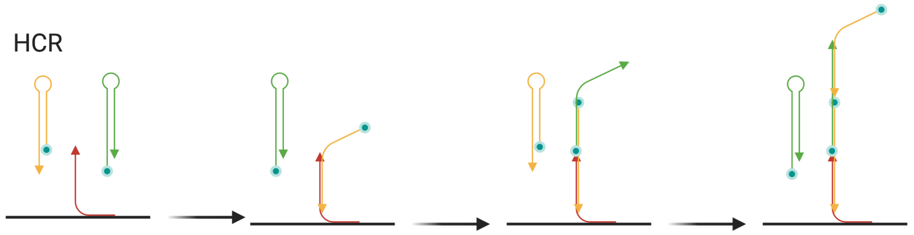

**Figure 5.15:** Schematic of HCR, showing 3 cycles, but this can continue indefinitely until H1 and H2 are exhausted. Arrow shows 5' to 3' direction.

the long stem is protected by the short stem, but can also hybridize with short stems of other hairpins (Figure 5.15). The long stem of H1 can hybridize to the short stem of H2, and vice versa (Figure 5.15). First, an initiator probe is hybridized to the transcript (24 per gene in the 2016 HCR-seqFISH study). Then the long stem of H1 hybridizes to the part of initiator not hybridized to the transcribe, now leaving the short stem vacant. Then the long stem of H2 hybridizes to the vacant short stem of H1, and now the short stem of H2 is vacant for another H1. This cycle can continue indefinitely until H1 and H2 are depleted. This way, many fluorophores are tethered to the target transcript without increasing the number of probes bound to the transcript, thus amplifying signal.

Similarly, RCA can continue indefinitely until DNA polymerase is inhibited or removed or when deoxynucleotides are depleted. In contrast, the bDNA moiety has a controlled size and does not grow indefinitely until stopped. In both bDNA and HCR, the amplified moiety is still anchored on the target transcript. In contrast, since when the padlock probe encloses the target, the DNA polymerase is inhibited (Baner et al. 1998), the padlock must be dissociated from the target before RCA, or in the case of target priming, the target cDNA itself grows into the RCA hairball. As the hairball is not anchored to the original target, it can drift away and obscure the original location of the target.

##### 5.2.3 Optical crowding

As we have seen, smFISH based spatial transcriptomics has been scaled to around 10,000 genes and can potentially be scaled to the whole transcriptome. With increasing number of mRNA molecules visualized, it's also increasingly likely for different target molecules to be so close to each other that their fluorescent spots overlap or are even within the diffraction limit of the optical microscope and appear as one point. This is the problem of optical crowding, and some existing ways to mitigate this problem are summarized below.

As already mentioned, SRM is not susceptible to this problem (Lubeck and Cai 2012), though access to SRM is not as common as access to regular confocal or epifluorescent microscopes. Another simple strategy is to select the most highly expressed genes from RNA-seq. These genes are imaged separately with smFISH, with one color and one round of hybridization per gene instead of combinatorial barcoding, as was done in the first MERFISH study (K. H. Chen et al. 2015). However, with increasing number of highly expressed genes, this method becomes increasingly laborious. Also as already mentioned, in seqFISH+, only 1 in 60 mRNA molecules of interest light up in each channel and round of hybridization (20 pseudocolors per channel and 3 channels), and the transcript spots can be computationally super-resolved, thus reducing optical crowding (Eng et al. 2019).

Another strategy is to allow transcript spots to overlap but computationally resolve them, as in corrFISH (Coskun and Cai 2016), BarDensr (S. Chen et al. 2021), ISTDECO (Andersson et al. 2021), and Composite In Situ Imaging (CISI) (Cleary et al. 2021). In corrFISH, Transcripts of highly expressed genes encoding ribosomal proteins were visualized with sequential hybridization and 2 colors but not every gene lights up in each round of hybridization; each gene is encoded by one color and a sequence of 0's (absence of fluorescence) and 1's (presence) of that color. Then images from different rounds of hybridization in the same FOV are correlated to identify transcripts that are 1's in both rounds amidst transcripts that are not 1's in both rounds. To the best of our knowledge, after its conception, corrFISH has not been applied to generate any new high throughput dataset.

A more recent method, BarDensr, models the observed brightness of potentially mixed spots in terms of the point spread function (PSF), codebook, unknown spot density, probe washing, background, and per round per channel gain. Then the unknown spot density and deconvolution of barcodes at mixed spots are inferred by maximizing sparsity of the spots in space (most voxels don't have spots) while keeping reconstruction loss of the observed brightness sufficiently low. BarDensr is very recently published, and, as of writing, we are unaware of studies that used the method. ISTDECO is similar but only uses a Gaussian PSF, codebook, and background.

CISI uses seqFISH-like barcoding, but does not even require spot detection. Gene abundance is computationally inferred with compressed sensing[6]. First, an autoencoder is trained on composite images with different channels. Then in the latent space inferred by the autoencoder, the channels are decompressed with compressed sensing principles and decoded into genes with the decoder branch of the trained autoencoder. The barcodes and genes must be carefully chosen from an existing dataset. The genes must be described by a small number of coexpression modules so module activity is sparse. Inferring the sparse module activity before inferring individual gene levels at the decompression step is more tractable than directly inferring individual gene abundances.

A strategy that has been reused is expansion microscopy (ExM). When a poly-

[6] https://en.wikipedia.org/wiki/Compressed_sensing

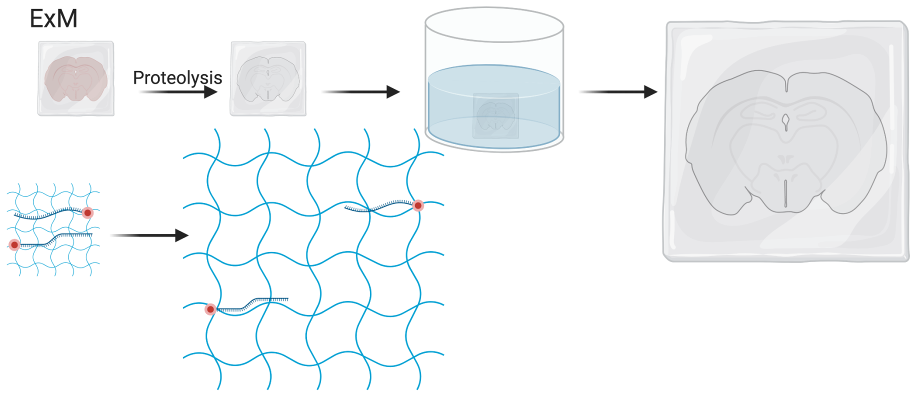

**Figure 5.16:** Schematic of expansion microscopy.

electrolyte gel is dialyzed in water, it expands as its polymer network changes into extended conformations (Chen, Tillberg, and Boyden 2015). First, the tissue is infused with monomers of the gel. Then with small molecule linkers, molecules of interest such as fluorophores and RNAs can be covalently incorporated to the polymer network over the course of free radical polymerization. After the gel forms, proteins in the tissue are digested to homogenize mechanical properties of the gel and to clear the tissue to reduce autofluorescent background. Then the gel is soaked in water to expand, linearly expanding 3 to 4.5 times on each side (Chen, Tillberg, and Boyden 2015; Chen et al. 2016) (Figure 5.16). This way, transcripts attached to the gel are physically separated, avoiding optical crowding. ExM has thus been adapted to MERFISH for this purpose (Wang, Moffitt, and Zhuang 2018), as well as EASI-FISH. In addition, EASI-FISH was used to quantify transcripts in 300 $\mu$m thick brain slices and imaging was accelerated with light sheet microscopy. However, a disadvantage of ExM is that each FOV now covers less of the original tissue, thus increasing imaging time. Furthermore, the expanded gel would continue to expand during the rounds of hybridization. As the expansion is non-linear and non-isotropic, barcode decoding is challenging as it's difficult to match transcript spots across rounds of hybridizations.

##### 5.2.4 Usage of smFISH based techniques

As already noted, the number of genes whose transcripts can be possibly quantified simultaneously in the same piece of tissue with highly multiplexed smFISH based technology has increased over time (Figure 5.17). The number of cells that can be imaged in one study has also increased (Figure 5.18). However, in practice, the actual number of genes and cells profiled has not significantly increased (Figure 5.19, Figure 5.21). These plots only show papers that reported the number of cells and genes in the main text; if we download and

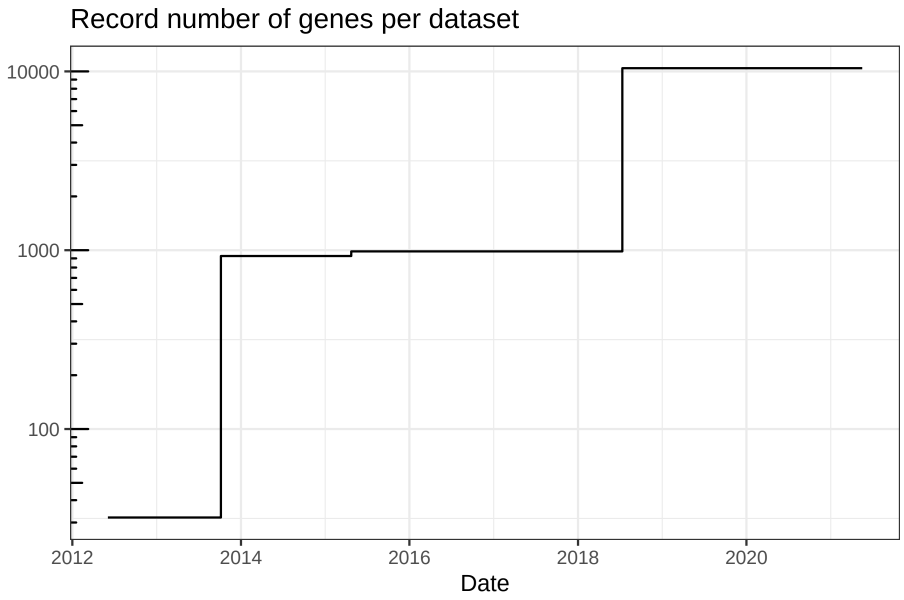

**Figure 5.17:** Record number of genes per dataset quantified by smFISH based techniques over time.

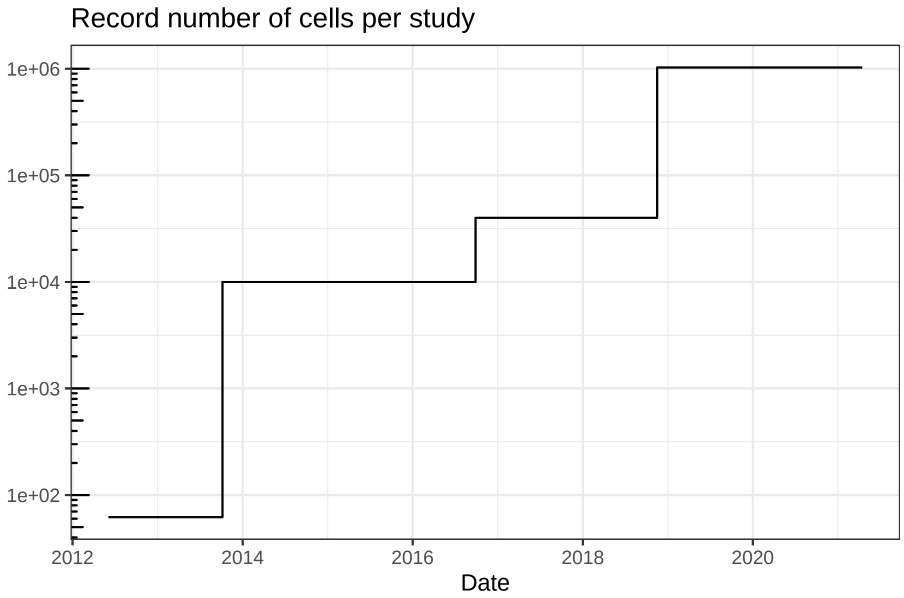

**Figure 5.18:** Record total number of cells per study profiled by smFISH based techniques over time.

process all publicly available datasets associated with such papers, the trends might change, although figures of papers that do not report the number of cells (number of genes is usually reported in smFISH and ISS studies) don't seem to indicate that the trend would change significantly. Moreover, as discussed in Section 5.6, some of the studies used smFISH based methods to visualize DNA loci and 3D chromatin structure alongside transcripts. The number of genes here is for the transcripts, including when only introns are targeted.

An earlier version of the plot of number of genes over time plotted the mean number of genes for each study, due to difficulty in defining what constitutes a dataset. However, since that version caused confusion as sometimes one study profiled very different number of genes in different experiments, we decided to give some definition of "dataset" and not to plot the mean. Here a "dataset" means either a different tissue, cell type, experimental or clinical condition, or a separate experiment profiling a different number or set of genes in the same study. One dataset can involve multiple sections and individuals.

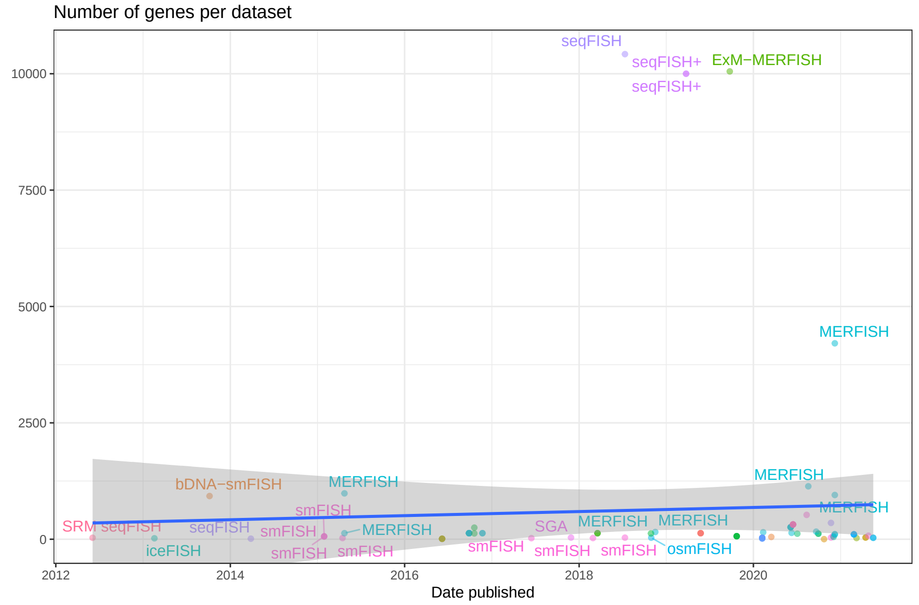

**Figure 5.19:** Number of genes per datasets in each study, over time. Gray ribbon is 95% confidence interval (CI). The points are translucent; more opaque points are multiple datasets from the same study.

The trend line looks pretty flat. Although studies quantifying a very large number of genes tend to be recent, many other studies profiling fewer genes pulled the line down. The slope (with all data, outliers and all) is not significantly different from 0 (t-test). The number of genes was not log transformed precisely to show that even the outliers can't give a significantly rising trend.

```
##
## Call:
## lm(formula = n_genes ~ date_published, data = smfish)
##
## Residuals:
##     Min      1Q  Median      3Q     Max
## -716.1 -618.5 -515.5 -407.1 9803.3
##
## Coefficients:
##                  Estimate Std. Error t value Pr(>|t|)
## (Intercept)    -1515.9241  4955.9415  -0.306    0.760
## date_published     0.1204     0.2772   0.434    0.665
##
## Residual standard error: 2108 on 89 degrees of freedom
##   (5 observations deleted due to missingness)
## Multiple R-squared:  0.002115,  Adjusted R-squared:  -0.009097
## F-statistic: 0.1886 on 1 and 89 DF,  p-value: 0.6651
```

To better show the trend of the vast majority of datasets that profiled far fewer than 10,000 genes, a broken y-axis is introduced to zoom into the majority of datasets (5.20).

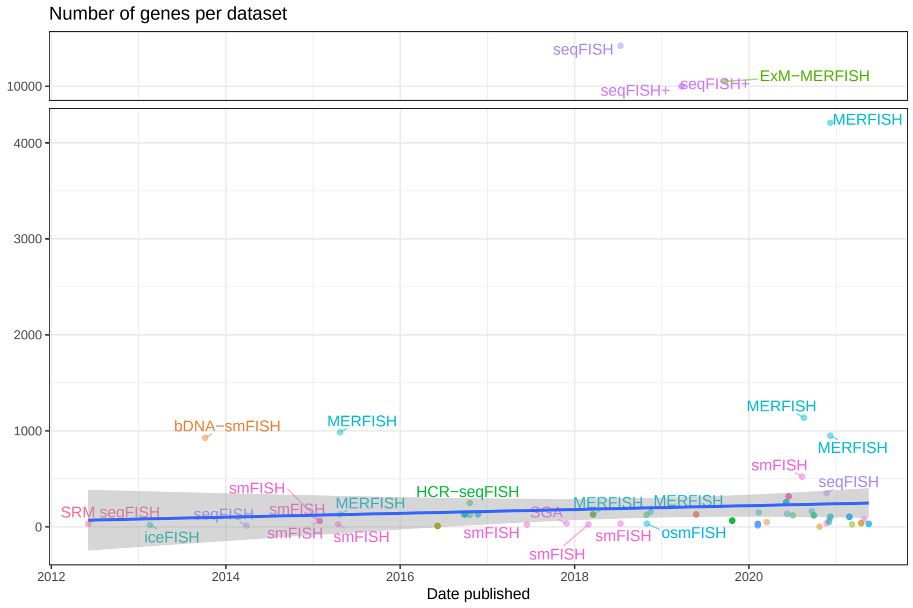

**Figure 5.20:** Like the previous figure, but more zoomed in to datasets with fewer than 10,000 genes.

How total number of cells profiled in each study that reported the number of cells in the main text is shown here. The total number across datasets is used because sometimes number of cells per dataset is not reported.

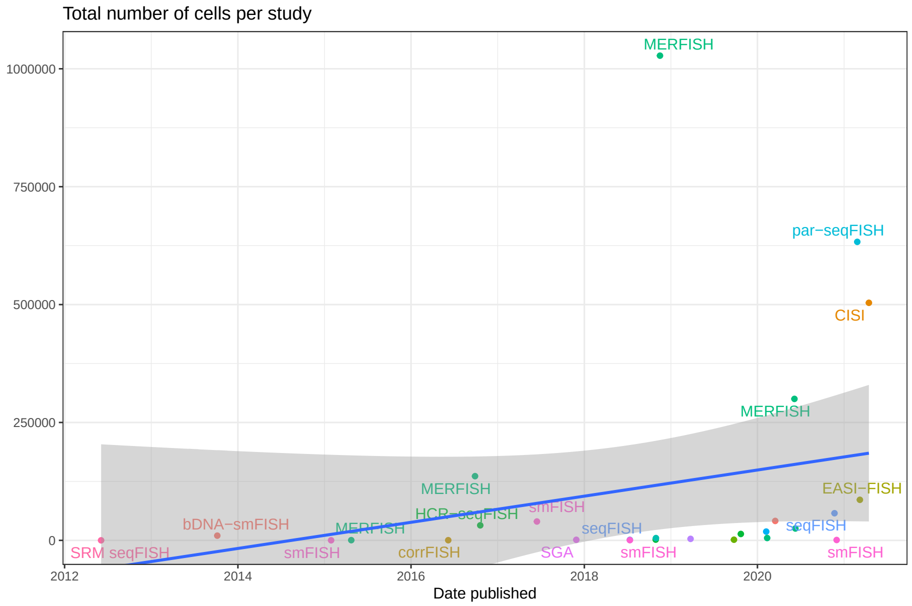

**Figure 5.21:** Total number of cells per study profiled by smFISH based techniques over time.

Again, although studies that profiled large numbers of cells tend to be more recent, as there are many recent studies with smaller numbers of cells, the slope (with outliers and all) is not significantly different from 0 (t-test).

```
##
## Call:
## lm(formula = n_cells ~ date_published, data = sum_cells)
##
## Residuals:
##     Min      1Q  Median      3Q     Max
## -173754 -120441  -95966   10604  909877
##
## Coefficients:
##                  Estimate Std. Error t value Pr(>|t|)
## (Intercept)    -1.237e+06  9.524e+05  -1.299    0.206
## date_published  7.591e+01  5.364e+01   1.415    0.169
##
## Residual standard error: 236800 on 25 degrees of freedom
## Multiple R-squared:  0.07416,    Adjusted R-squared:  0.03713
## F-statistic: 2.003 on 1 and 25 DF,  p-value: 0.1694
```

MERFISH is the technique used in the most studies (Figure 5.22), although most of the smFISH based techniques barely spread beyond their institutions of origin, if at all (Figure 5.23). The following advantages and disadvantages of smFISH based techniques may explain these trends in usage. Advantages and disadvantages of individual smFISH based techniques reviewed so far are summarized in Table 5.1.

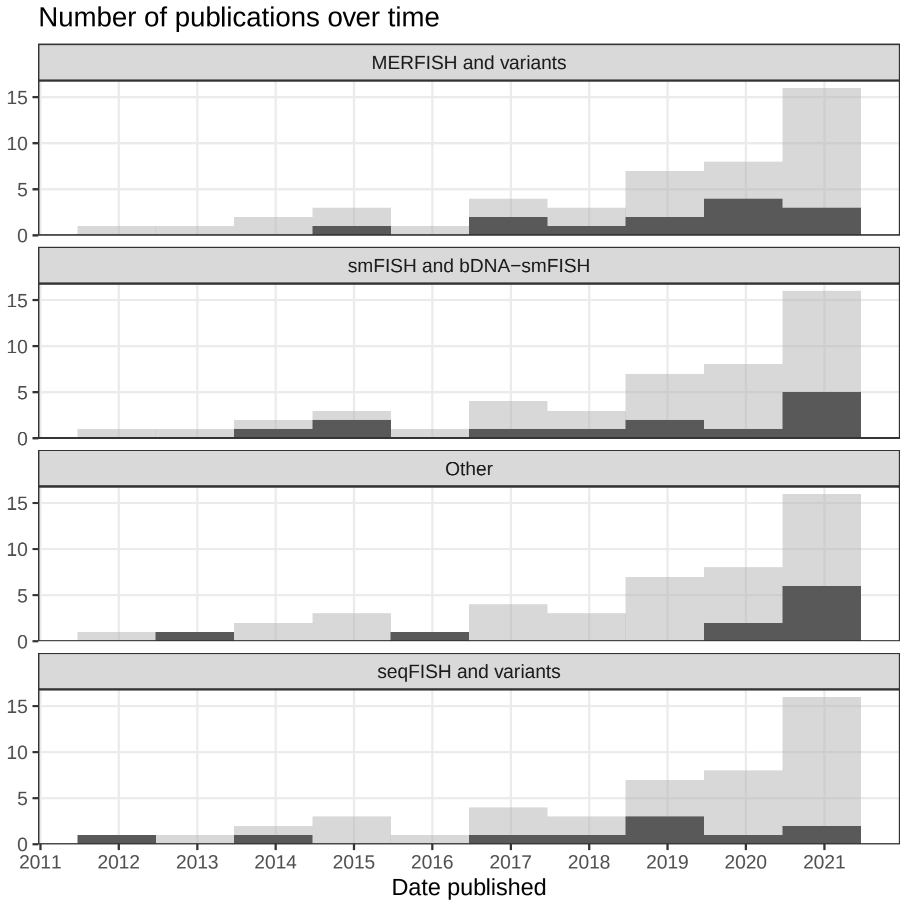

**Figure 5.22:** Number of publications over time, broken down by technique type. Preprints are included, and the gray histogram in the background is the overall trend of all smFISH based techniques.

MERFISH and seqFISH and their variants are more used and have spread, though their use still concentrates in their institutions of origin (Harvard for MERFISH and Caltech for seqFISH) (Figure 5.24). The one use of MERFISH at Caltech and the use of seqFISH at Dana-Farber and Cambridge are due to collaboration between the two institutions; the first authors analyzed the data

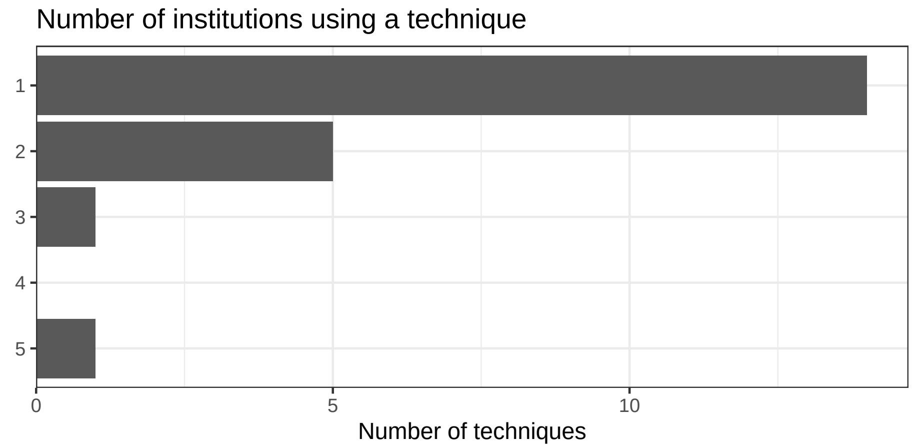

**Figure 5.23:** Number of techniques that have been used by each number of institutions; most techniques have only been used by 1 institution, i.e. the institution of origin.

while the data itself was most likely still collected at the techniques' institutions of origin. Nevertheless, this shows interest in MERFISH and seqFISH beyond their institutions of origin.

SmFISH based techniques have the following advantages. First, smFISH, especially with larger number of probes, have nearly 100% detection efficiency of transcripts (Lubeck and Cai 2012). With combinatorial barcoding, however, the efficiency is decreased. Compared to smFISH, MERFISH version 2 with HD4 code has about 95% detection efficiency on 130 genes and 92 probes per gene, although the efficiency dropped to ~25% with the HD2 code that can encode nearly 1000 genes but can only identify but not correct errors (Moffitt et al. 2016; Foreman and Wollman 2019). When scaled to 10,050 genes, MERFISH has around 79% detection efficiency (C. Xia, Babcock, et al. 2019). As for HCR-seqFISH, the efficiency is around 84% (Shah et al. 2016), and for seqFISH+, around 49% (Eng et al. 2019). Nevertheless, this is much better than the efficiency of ST, which is around 6.9% compared to smFISH (Ståhl et al. 2016). To put the 6.9% in context, from ERCC spike ins and in some cases comparison to smFISH, scRNA-seq methods such as Drop-seq, 10X, inDrop, CEL-seq, and CEL-seq2 have capture efficiency of between 3% and 25% (Macosko et al. 2015; Zheng et al. 2017; Klein et al. 2015; Hashimshony et al. 2016; Grün, Kester, and Oudenaarden 2014). Thus smFISH based spatial transcriptomics methods can be much more efficient than scRNA-seq, though efficiency of RCA based smFISH compared to regular smFISH has not been reported.

Second, since individual transcripts are imaged and counted, smFISH based methods are highly quantitative and records subcellular localization of transcripts. While most smFISH based spatial transcriptomics studies analyze data

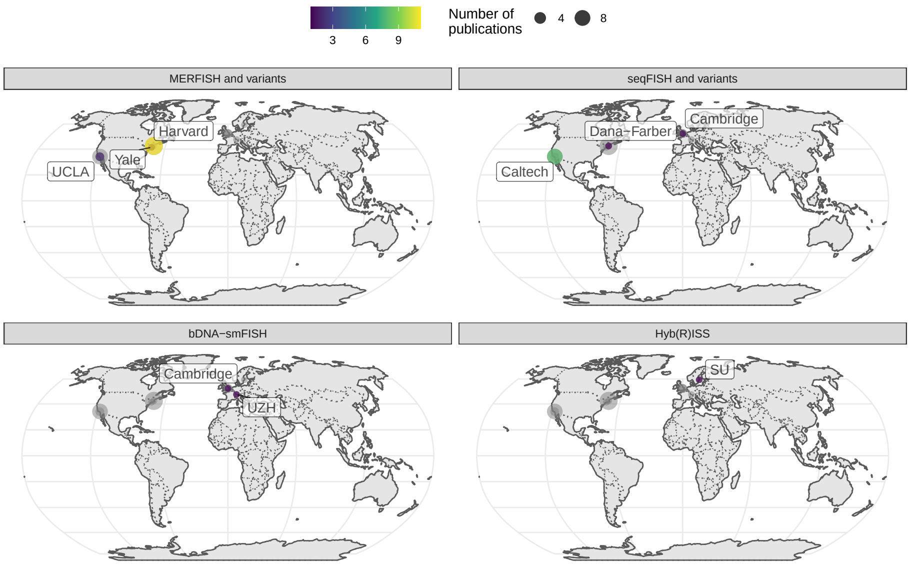

**Figure 5.24:** Geographical locations of institutions that used certain techniques. Point area is proportional to number of publication from the city of interest. Gray points in the background is all publications using smFISH based techniques. The cities and institutions labeled are those of the first author. Note that for seqFISH, the hidden Markov random field (HMRF) study at Dana Faber (Zhu et al. 2018) and the mouse embryo study (Lohoff et al. 2020) had collaboration with Long Cai's group at Caltech, so the dataset was most likely still collected at Caltech.

at the cellular gene count level, not using subcellular transcript localization, cells have been shown to show great variation in subcellular localization of transcripts of the same set of genes and a number of "archetypal" patterns have been described (Samacoits et al. 2018; Stoeger et al. 2015; Cabili et al. 2015).

The following disadvantages may explain why smFISH based spatial transcriptomics has not been widely used on large number of cells and genes (Figure 5.19, Figure 5.21), and why MERFISH is the most used technique (Figure 5.22). First, multiple rounds of hybridization and high magnification mean that data collection is time consuming. MERFISH version 2 greatly sped up imaging, as version 1 requires higher magnification and needs to photobleach fields of view (FOV) one at a time; one FOV in version 1 is 40 $\mu$m $\times$ 40 $\mu$m, while one FOV in version 2 is 223 $\mu$m $\times$ 223 $\mu$m. Version 2 also cut imaging time in half by using 2 colors, targeting 2 bits per round. This way, for 130 genes and 40,000 cells, MERFISH took about 18 hours (Moffitt et al. 2016), while HCR-seqFISH would take days because of overnight hybridization after probes are stripped for each round of hybridization although the seqFISH barcode is much shorter.

When scaled to 10,000 genes, MERFISH takes 23 rounds of hybridization (C. Xia, Fan, et al. 2019), while seqFISH+ takes 80 rounds (Eng et al. 2019), although because ExM was used for MERFISH in this case to reduce optical crowding, expanding the area to be images ~4 folds, the actual imaging time of ExM-MERFISH and seqFISH+ here may have been comparable. Perhaps MERFISH has been scaled to larger number of cells and used in more studies beyond the institution of origin (Figure 5.24) because of the higher detection efficiency and shorter imaging time.

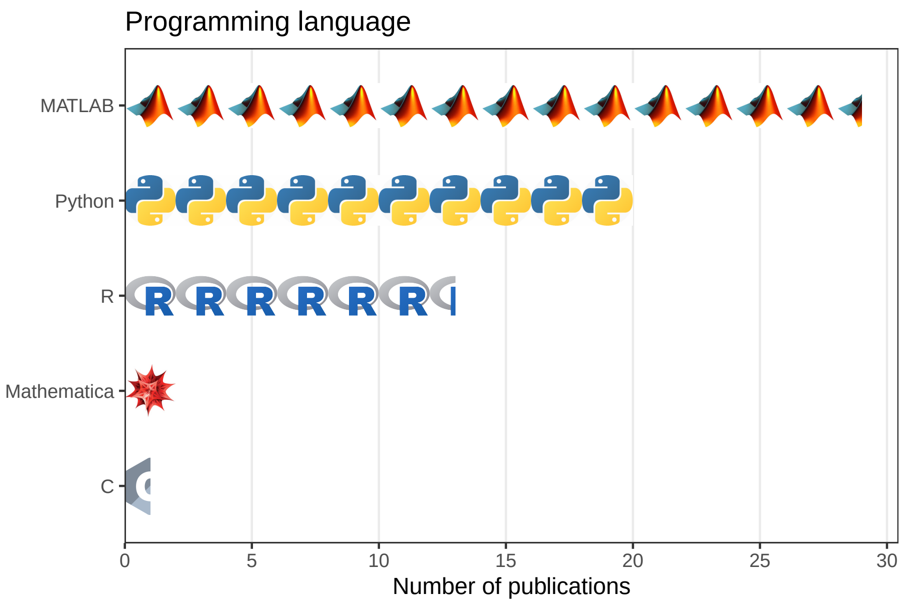

**Figure 5.25:** Number of publications using smFISH based techniques that used each of the 5 most common programming languages. Each icon stands for 2 publications.

Second, with increasing area of tissue and number of genes covered, smFISH based spatial transcriptomics generates terabytes of images – for each FOV, there is an image for each channel, z-plane, and round of hybridization. Images from the MERFISH dataset of 40,000 cells and 130 genes took 2 to 3 days to process on a multi-core server, although the number of cores was not stated (Moffitt et al. 2016). In contrast, it takes hours, or even just minutes, to process the fastq files of a scRNA-seq dataset to get the gene count matrix (Melsted et al. 2021), nor do the fastq files take up so much disk space. Until 2019, software to process such images and to decode the combinatorial barcodes was typically written in the proprietary programming language MATLAB (Figure 5.25), and poorly documented, so it was difficult for people outside the lab of origin to use.

More recently, Python is replacing MATLAB as the programming language of choice to write such image processing software. The Chan Zuckerberg Initiative developed starfish[7] in Python as a unified framework to process smFISH based spatial transcriptomics data (Axelrod et al., n.d.). However, `starfish` is still immature and not sufficiently efficient, nor is the functionality to integrate multiple FOVs developed. Perhaps because of this, `starfish` has not been widely used and image processing pipelines specific to each technology have been developed instead, such as MERlin for MERFISH (C. Xia, Fan, et al. 2019) and IRIS for ISS (Zhou et al. 2020). In contrast, for scRNA-seq, there are popular data processing tools that apply across technologies, such as STAR (wrapped by Cell Ranger) (Dobin et al. 2012), alevin (Srivastava et al. 2019), and kallisto (Melsted et al. 2021). Furthermore, even with an open source and interoperable image processing pipeline, cell segmentation, which is essential to obtaining the gene count matrix commonly used in data analysis, is challenging.

Third, custom flow cells have been used for the numerous rounds of hybridization (Eng et al. 2019; Moffitt et al. 2016; Codeluppi et al. 2018). These custom flow cell and pump systems are not commercially available and need to be built by any lab that wishes to adopt the smFISH based technologies. To the best of our knowledge, there are no core facilities that perform smFISH based spatial transcriptomics. Thus for the user, adopting an smFISH based spatial transcriptomics technique means not only learning a new syntax to process images, made difficult in some cases by the cost of MATLAB and lack of documentation, but also setting up a complex custom flow cell system, which may not be feasible with microscopy cores. Finally, smFISH based techniques require a pre-defined list of genes and probes, so unlike in RNA-seq, novel transcripts would be missed.

So far we have reviewed studies that showcase new techniques and technical improvements such as signal amplification and resolving optical crowding. Some smFISH based techniques have been used in studies that focus on biological problems rather than new techniques. HCR-seqFISH has been used twice in biological studies, in chicken neural tube (35 genes) (Lignell et al. 2017) and mouse T cell precursors (65 genes) (Zhou et al. 2019) though both were conducted within Caltech, the institution of origin. Moreover, spatial location of cells is not necessarily a reason to use HCR-seqFISH; Zhou et al. used HCR-seqFISH because of the high detection efficiency compared to scRNA-seq in dissociated FACS sorted T cell progenitors, so when spatial information is already lost. More recently, pseudocolor seqFISH was used in a mouse embryo atlas at University of Cambridge (though Long Cai is still a coauthor), finally moving beyond the stage of testing into new biological research (Lohoff et al. 2020). A new version of seqFISH, par-seqFISH, was developed to profile 105 genes in the biofilm bacterium *Pseudomonas aeruginosa* (Dar et al. 2021). This may open the way to spatial transcriptomics in not only biofilms, but in the microbiome in general.

MERFISH has been used more broadly in biological studies. Within Harvard,

[7] `https://spacetx-starfish.readthedocs.io/en/stable/`

**Table 5.1:** Pros and cons of smFISH based techniques.

| Technique | Pro | Con |
|---|---|---|
| HCR-seqFISH | Relatively high efficiency (84%), fewer rounds of hybridization, error correction | Lower efficiency than MERFISH, time consuming to re-hybridize probes to target after stripping |
| seqFISH+ | Avoids optical crowding, scalable | Lower efficiency (49%), numerous rounds of hybridization |
| MERFISH | High efficiency (95%) with HD4 code, error correction, version 2 relatively fast, scalable | Numerous rounds of hybridization, numerous probes requiring long transcripts though this is resolved by bDNA signal amplification |
| ExM-MERFISH | Avoids optical crowding, clears tissue | Each FOV contains less of the original tissue |
| HybISS | Only 5 probes per gene, applicable to isoform specific exons, padlock probe reduces background, lower magnification when imaging (20x and 40x, while MERFISH uses 60x), can discern SNPs | Error correction not reported, amplicon takes up space and might drift away |
| SCRINSHOT | Similar to HybISS, but designed for FFPE tissue | Same as HybISS |
| HybRISS | Avoids inefficiency of reverse transcription, better signal to noise ratio and more transcripts detected then HybISS. | Padlock probe sequences are proprietary to CARTANA |
| bDNA-smFISH | Commercial RNAscope kit, reduces background and amplifies signal, amplified moiety does not grow indefinitely | Except for bDNA-MERFISH, it has not been used in a highly multiplexed setting |

the institution of origin, MERFISH has been used to create atlases of the hypothalamic preoptic region (155 genes) (Moffitt et al. 2018) and the primary motor cortex (MOp) (258 genes) (Zhang et al. 2020) in mice, and adapted to stain for chromatin conformation and transcription foci (introns) (Su and Song 2020). Outside Harvard (Figure 5.24), MERFISH has been used to study how gene expression variability relates to cell state in cell culture (Foreman and Wollman 2019) and used in conjunction with smFISH based chromatin tracing to study the relationships between chromatin compartmentalization and gene expression (M. Liu et al. 2020).

After its inception, HybISS became part of a single cell atlas of the developing mouse nervous system (Manno et al. 2020). This atlas is mostly scRNA-seq data, but 119 genes were stained with HybISS to validate secondary organizers discovered via scRNA-seq.

#### 5.3 In situ sequencing

In contrast to smFISH based techniques, techniques reviewed in this section determine the sequences of the target transcript or the gene specific barcode by *in situ* sequencing by ligation (SBL) or sequencing by synthesis (SBS) to distinguish between transcripts of different genes. This section reviews 3 *in situ* SBL strategies, SOLiD, cPAL, and SEDAL, and the spatial transcriptomics techniques using them.

SBL relies on the specificity of the DNA ligase, so ligation only occurs when both sequences to ligate match the template in the vicinity of the site of ligation. Prior to SBL, this specificity was used to detect SNVs that would otherwise be missed as ISH probes can tolerate some mismatches. A technique using ligation of two oligonucleotides to detect SNVs was introduced in 1988 by Ulf Landegren (Landegren et al. 1988), laying the foundation of SBL (Figure 4.3). The padlock probe came in the same tradition of SNV detection, and Mats Nilsson worked with Landergren when creating the padlock probe (Nilsson et al. 1994).

Almost all spatial transcriptomics techniques based on SBL require *in situ* reverse transcription of the mRNAs as ligation with RNA as template is inefficient. As already mentioned in Section 5.1.1.2, IVT amplification of transcripts from single cells for expression profiling originated in the Eberwine group (Van Gelder et al. 1990; Eberwine et al. 1992), where rather than LCM, the cDNAs from the single cells were reverse transcribed *in situ* during electrophysiological recording before the cellular content was aspirated for IVT amplification. This was built upon the *in situ* reverse transcription technique from the Eberwine group in 1988 (Tecott, Barchas, and Eberwine 1988), where the cDNA of proopiomelanocortin (POMC) was radiolabeled so the spatial distribution of the mRNA was visualized on an autoradiograph. This made most *in situ* SBL techniques possible, which instead of radioactivity, use gene barcodes to locate the transcripts in a multiplexed and safer way.

##### 5.3.1 SOLiD and FISSEQ

The earliest proposal of SBL we are able to locate is a patent filed in 1995 describing a method similar to sequencing by oligo ligation detection (SOLiD). An initiator oligonucleotide hybridizes to the template to be sequenced, and is extended by ligation to a 9-mer probe with a label such as a fluorophore that indicates one or two nucleotides of the probe (Macevicz 1995). The probe has a blocking moiety so only one probe is ligated in each cycle. Then the blocking moiety is removed so the initiator can be further extended by ligation in the next cycle. As mismatches in the probe inhibit ligation, the nucleotide of interest in the probe can be read off from the label after probes that are not ligated are removed. This can determine every 9th nucleotide in the template, and with 9 different initiators, each out of phase by one nucleotide, the sequence of the entire template can be determined. However, this method existed only on paper, while since 2006, Applied Biosystems (Applera) seemed to have developed SOLiD independently from that patent after acquiring Agencourt, which developed a sequencing by ligation method that would be the foundation of SOLiD (Alsup 2009).

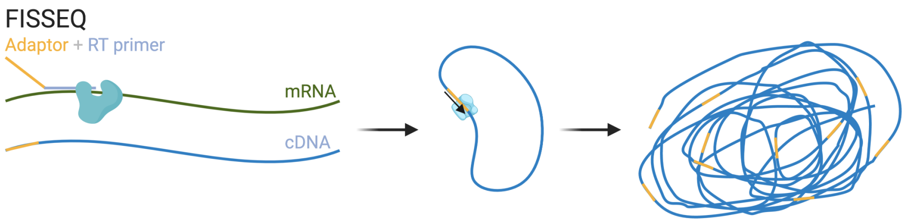

**Figure 5.26:** Schematic of RCA in FISSEQ.

In 2014, single cell resolution and transcriptome wide spatial transcriptomics was far out of reach (Figure 5.17). An attempt to reach this goal was fluorescent in situ sequencing (FISSEQ) (Lee et al. 2014). A universal adapter and random hexamer reverse transcription (RT) primer was hybridized to the mRNAs to reverse transcribe them into cDNA (Figure 5.26). Then the cDNA, now with the adaptor on the 5' end, is circularized, and amplified with RCA with a primer complementary to the adaptor. Then again, with sequencing primers receding into the adaptor, SOLiD is used to sequence the cDNA amplicons in situ.

In SOLiD, color of the fluorophore encodes the two 3'-most bases of 8-mer probes with other bases degenerate (Figure 5.27). Once a probe perfectly matching the target right after the primer, the probe is ligated to the primer and the fluorescent signal is recorded. Then the fluorophore and the nearest 3 bases of the probe are cleaved off. In the next cycle, a new matching probe is ligated to the now extended primer. This is continued until the end of the target, for 7 cycles per primer in the case of FISSEQ (Lee et al. 2015). For the first 7 cycles, the primer matches the adaptor (N). Then the primer N, extended for 7 cycles is stripped, and a new primer receding one nucleotide to the 5' end of the

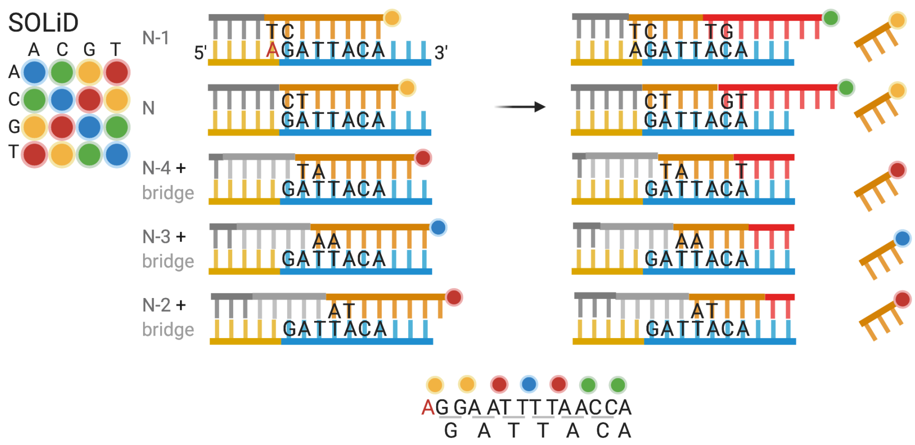

**Figure 5.27:** Schematic of SOLiD sequencing, determining the sequence GATTACA. The rows are arranged in the order of 5' to 3' positions of the first fluorescent probe, but the actual hybridization and ligation can take a different order. As part of the constant region, the 'A' highlighted in red is known.

adaptor (N-1) is added in cycle 8. Again 7 cycles of ligation are performed and the extended primer N-1 is stripped after cycle 14 to make room for N-2. For N-2, N-3, and N-4, a bridge oligo is used so the target with unknown sequence, rather than the adaptor with known sequence, is interrogated by the probes. With N through N-4, the entire target is covered. With the fluorescent signals recorded from the rounds of ligation, and the knowledge of the last nucleotide of the adaptor interrogated by the first ligation to primer N-1, the sequence of the target can be determined. Figure 5.27 shows how SOLiD determines the sequence "GATTACA". As already mentioned in the smFISH section, with increasing number of genes profiled, optical crowding is increasingly a problem. To mitigate optical crowding, the primer N can have one or more degenerate bases at the 5' end reaching into the target; with one degenerate base, only 1/4 of the amplicons are sequenced. With two bases, this would be 1/16. This is repeated to cover all transcripts, but increases imaging time.

While FISSEQ may seem a promising approach to reach the goal of single cell resolution and transcriptome wide spatial transcriptomics that unlike smFISH based techniques, is not limited by pre-defined gene panels, it has been largely dormant since its inception due to the following disadvantages. First, SOLiD has fallen out of favor because of limited read length when used in situ (5-30 nt), propagation of errors from previous cycles (Alon et al. 2020), and difficulty in sequencing panlindromic sequences (Giani et al. 2020). SOLiD was chosen for FISSEQ because it works well at room temperature; though SBS supports longer read lengths, it requires a heated stage (Lee et al. 2015). Second, FISSEQ is extremely inefficient, over 20 times less sensitive than scRNA-seq and two orders of magnitude less sensitive than 2013 Nilsson ISS (discussed later in this section)

(Lee et al. 2015), in part because of inefficiency of random RT priming (Lee et al. 2014) and tight packing of amplicons (Alon et al. 2020). Furthermore, as ribosomal RNA (rRNA) is not depleted, ~40-80% of FISSEQ reads are rRNA (Lee et al. 2014, 2015). Third, highly abundant genes involved in translation and splicing is depleted in FISSEQ compared to bulk RNA-seq (Lee et al. 2014). Finally, FISSEQ imaging is time consuming, taking 2 to 3 weeks if performed manually (Lee et al. 2015).

With expansion microscopy, the idea of FISSEQ was revived in ExSeq (Alon et al. 2020). Just like in ExM-MERFISH, transcripts are incorporated into a polyelectrolyte gel, which is expanded, so the amplicons are no longer so tightly packed. This eliminated the depletion of highly abundant genes compared to bulk RNA-seq, and the detection efficiency and proportion of rRNA reads of ExSeq seem on par with randomly primed bulk RNA-seq of adjacent sections. In addition to SOLiD sequencing as in FISSEQ, the amplicons are also sequenced ex situ with Illumina SBS. The in situ sequences are matched to ex situ sequences and only unique matches are kept, to more effectively align amplicons to the genome and to localize mRNA sequence variations such as alternative splicing that are more difficult to detect with SOLiD's short read length. There is also a targeted version of ExSeq, in which padlock probes with gene specific barcodes are RCA amplified and the barcodes are sequenced in situ by either SOLiD or Illumina SBS, profiling up to 297 genes; the detection efficiency is 62% compared to smFISH (i.e. for select genes in the same cell types, the number of transcripts detected is about 62% compared to smFISH), which is high compared to ~5% for 2013 Nilsson ISS but lower than that of MERFISH (HD4) and HCR-seqFISH (Alon et al. 2020; Lein, Borm, and Linnarsson 2017). Eight probes were designed for each gene, and the transcripts must be at least 960 nt long, shorter than required by MERFISH (without bDNA) and seqFISH variants. To our best knowledge, ExSeq has yet been used to collect new datasets after its inception.

##### 5.3.2 cPAL and ISS

An alternative SBL scheme is combinatorial probe anchor ligation (cPAL), which to our best knowledge, was first demonstrated in 2005 (Shendure 2005). In cPAL, an anchor primer is hybridized to a constant region immediately adjacent to the target. T4 DNA ligase requires matching base pairing up to 6 bases from the ligation junction when ligating from 5' to 3' and 7 bases when ligating from 3' to 5'. The first base of the target 5' to the constant region is interrogated by a 9-mer probe whose 5' most base is represented by the color of a fluorophore and ligated to the primer if a perfect match is present (Figure 5.28). Then the ligated construct is stripped and a new primer is hybridized to the constant region. The second base is interrogated by a 9-mer probe whose second 5' most base is represented by the fluorophore. This can carry on until the 6th base on the 5' direction. When the constant region is 5' to the target, bases

3' to the constant region are interrogated in a similar fashion. With constant regions flanking a target so primers bind in both direction, a 13 nt target can be sequenced this way, and the read length can be somewhat increased by adding degenerate bases to the anchor primer extending into the target (Drmanac et al. 2010).

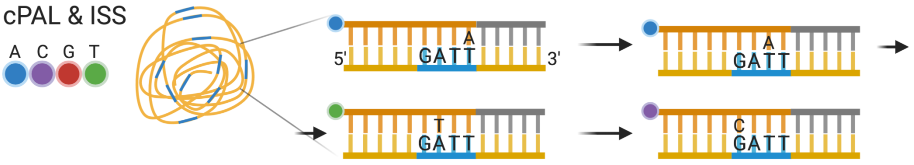

**Figure 5.28:** Schematic of cPAL as used in ISS.

The only in situ sequencing method that was reused after its inception was originally demonstrated in 2013 by Mats Nilsson's group (Ke et al. 2013), which we call ISS here (Figure 4.7). First, padlock probes are hybridized to in situ reverse transcribed cDNAs and RCA amplified (Figure 5.14). The padlock probe can carry a gene specific 4 nt barcode (barcode version), or leave a 4 nt gap between the ends of the probe after it's hybridized to the cDNA to be filled when the probe is circularized (gap filling version). Then the barcode or the filled gap is sequenced in situ, with an anchor primer binding 3' to the target, with cPAL. Because of limited read length of cPAL, short sequences uniquely identify each gene and isoform for the gap filling approach becomes difficult to find with increasing number of genes and isoforms. In contrast, a barcode with length $n$ can encode $4^n$ genes and isoforms. As a result, the barcode approach was repeatedly used after the inception of ISS and was commercialized by CARTANA, which was recently acquired by 10X Genomics.

The barcode approach was initially used to profile 39 genes (Ke et al. 2013), but has been used to profile up to 222 genes in human brains affected by Alzheimer's disease (Chen et al. 2020). Although, as already mentioned, ISS has much lower detection efficiency than smFISH based methods, because of RCA and this low detection efficiency, the density of imaged amplicons is lower, allowing for imaging at lower resolution (20x; MERFISH uses 60x) and thus facilitating profiling large areas of tissues such as whole mouse brain coronal sections (Qian et al. 2020; Partel et al. 2019). ISS has also been used in conjunction with spatial transcriptomics techniques that are transcriptome wide but lack single cell resolution, such as ST. Panels of usually fewer than 100 genes of interest are selected from ST and scRNA-seq data, to be profiled with ISS for more in depth characterization of these genes (Chen et al. 2020; Asp et al. 2019). In addition, because of the specificity conferred by the padlock probe and the small number of probes required per gene (usually 5 per gene but can be fewer), ISS has been used to quantify isoforms from isoform specific exons and exon-exon junctions (Lebrigand et al. 2020).

The number of genes that can be profiled by ISS is limited by the barcode length. Just like in seqFISH, only a small subset of all possible barcodes given a barcode length is used for error correction. As a result, to profile the entire transcriptome of over 20,000 genes, the barcode should be at least 8 nt long (65,536 barcodes), while in one direction, cPAL can only sequence 6 or 7 nt and degenerate bases. It is possible in theory to lengthen the barcode to up to 13 nt by sequencing from both ends of the barcode as in the original 2005 method (Shendure 2005). However, with increasing number of transcripts comes the problem of optical crowding, which is exacerbated by the physical size of the RCA amplicon. Perhaps ExM can be used here to mitigate optical crowding just like in ExSeq. To address the limitation in barcode length, HybISS, i.e. hybridization-based in situ sequencing, was devised (Gyllborg et al. 2020) so the now seqFISH-like barcode can be arbitrarily lengthened by increasing the number of rounds of hybridization. HybISS has already been reviewed in Section5.2.2.2; despite the "ISS" in its name, HybISS is classified as smFISH based because it does not involve SBL or SBS. HybISS also has up to 5 fold higher signal to noise ratio than ISS, and has somewhat higher detection efficiency than ISS though the improvement is less than 2 fold (Gyllborg et al. 2020). Comparison between HybISS and smFISH has not been reported. Nevertheless, HybISS has not yet been scaled to more than 120 genes and ExM may still be needed for transcriptome wide profiling.

##### 5.3.3 SEDAL and STARmap

Both SOLiD and cPAL have some drawbacks. As the gene specific barcode does not have to be long to encode all genes in the genome, when the barcode is used, limits in read length is not a major limitation. Because one color encodes two bases, SOLiD is very accurate (Liu et al. 2012), but error in one cycle propagates to later cycles. At least in the mouse brain, SOLiD also has high background (Wang, Moffitt, and Zhuang 2018). In contrast, cPAL does not have an inbuilt error rejection mechanism; the barcode must be elongated to allow for error correction, much like in the error correction scheme of seqFISH. Furthermore, in ISS, the mRNA is first reverse transcribed into cDNA because ligation of the padlock probe is inefficient when the template is RNA (Larsson et al. 2010). However, the efficiency of RT depends on the gene of interest and the variability of RT efficiency depends on RNA concentration (Schwaber, Andersen, and Nielsen 2019; Bustin et al. 2015).

A new method of in situ sequencing, namely sequencing with error-reduction by dynamic annealing and ligation (SEDAL) in spatially-resolved transcript amplicon readout mapping (STARmap), was devised to address these shortcomings (Wang, Moffitt, and Zhuang 2018). In STARmap, the specific amplification of nucleic acids via intramolecular ligation (SNAIL) probe is a derivative of the original padlock probe that avoids RT altogether. A primer partially hybridizes to the mRNA, and partially to the padlock probe (Figure 5.29). The padlock probe carrying a 5 nt gene specific barcode hybridizes to the mRNA adjacent to

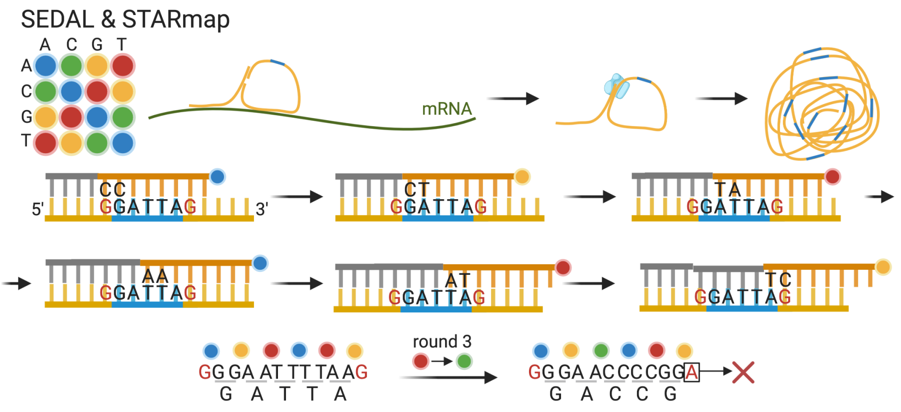

**Figure 5.29:** Schematic of RCA of SNAIL probe and SEDAL. Also showing error propagation and identification of 2 base encoding. As part of the constant region, the 'G' highlighted in red is known.

the primer, but both ends of the padlock probe hybridize to the primer instead, so when the ends are ligated together, the template is DNA rather than RNA, thus avoiding both RT and inefficiency of ligation with RNA template, and then the primer is used to initiate RCA. As both the primer and the padlock probe must match the mRNA template for RCA to occur, SNAIL probes are specific and background of non-specific binding is eliminated. To reduce background autofluorescence and prevent the RCA amplicons from moving, the amplicons are crosslinked into a hydrogel and the tissue is cleared of proteins and lipids.

Then SEDAL is used to sequence the gene specific barcodes. The sequences flanking the gene barcode are known. In the first round an anchor or reading probe binds to the constant region 5' to the barcode, one base away from the barcode (Figure 5.29). The decoding probes are 8-mers labeled with a fluorophore at the 5' end whose color represents the 2 nucleotides at the 3' end that interrogates the barcode; the other bases are degenerate. If the decoding probe matches the barcode, then it is ligated to the reading probe and the fluorescent signal is recorded. In the first round, the decoding probe interrogates the last base of the constant region and the first base of the barcode, as the last base of the constant region is necessary to decode the sequence of colors. Then the reading and decoding probes are stripped. In the second round, the reading probe stops right where the barcode starts. In the third round, the reading probe has a degenerate base extending into the barcode. Reading probes of the following rounds extend further into the barcode with degenerate bases. In the last round, the decoding probe interrogates the last base of the barcode and the first base of the following constant region. Like in SOLiD, with 2 base encoding, an error in a previous round propagates into later rounds; with propagation, when there is an error when decoding, then the first base of the constant region after the barcode would be incorrectly decoded, so the error is identified and rejected. Comparison of detection efficiency of STARmap with that of smFISH has not been reported; the efficiency is reported to be somewhat better, at least not worse, than that of scRNA-seq, suggesting that STARmap is perhaps more efficient than ISS, but most likely much less efficient than MERFISH (HD4) and seqFISH.

##### 5.3.4 Sequencing by synthesis

While most *in situ* sequencing techniques use SBL, some use SBS, indeed with a heated stage to perform SBS in situ. Because Illumina SBS is much more well-known and widely used than SBL for NGS, we will not recap it here. SBS has been tried to sequence DNA barcodes of antibodies in highly multiplexed immunofluorescence (Kohman and Church 2020). BARseq (Chen et al. 2019), a method to trace neuron projections is also based on SBS. In BARseq, the gap filling version of ISS (Section @ref{cpal}) is used and the filled gap that is the projection tracing barcode is sequenced with Illumina SBS chemistry. BARseq has also been adapted to profile endogenous transcripts (up to 79 genes as of writing) and image projection barcodes in the same neurons (BARseq2) (Sun et al. 2020; S. Chen et al. 2021); gene expression and projection can be correlated in some though not all cells. For endogenous transcripts, the mRNA is first reverse transcribed, and the barcode version of ISS (Section @ref{cpal}) is used to amplify the barcodes (in the padlock probe but not the cDNA) with RCA, which are then sequenced *in situ* with SBS. For transcripts, BARseq2 detects slightly more copies of mRNAs than 10X v3 scRNA-seq for the same gene in the same tissue.

#### 5.4 In situ array capture

This section reviews techniques that capture transcripts from a permeabilized tissue section on a spatially organized array for RNA-seq. These techniques are similar to 3' based scRNA-seq, with amplification and sequencing handle, barcode, UMI, and poly-T to capture polyadenylated transcripts, except that each spot in the array has its own barcode, rather than each droplet. These techniques can be transcriptome wide, but do not have single cell resolution; the resolution is the size and shape of the spots. In ST and Visium, the array is constructed by printing the capture sequences onto commercial microarray slides, so the 5' end of the sequences are attached to the slide; where each spatial barcode is placed is known. Alternatively, the capture sequences can be attached to beads like in droplet scRNA-seq, as in Slide-seq and HDST. The beads are randomly placed on a slide in a single layer, and the location of barcodes are determined before library preparation when the capture sequences and transcripts are released from the slide.

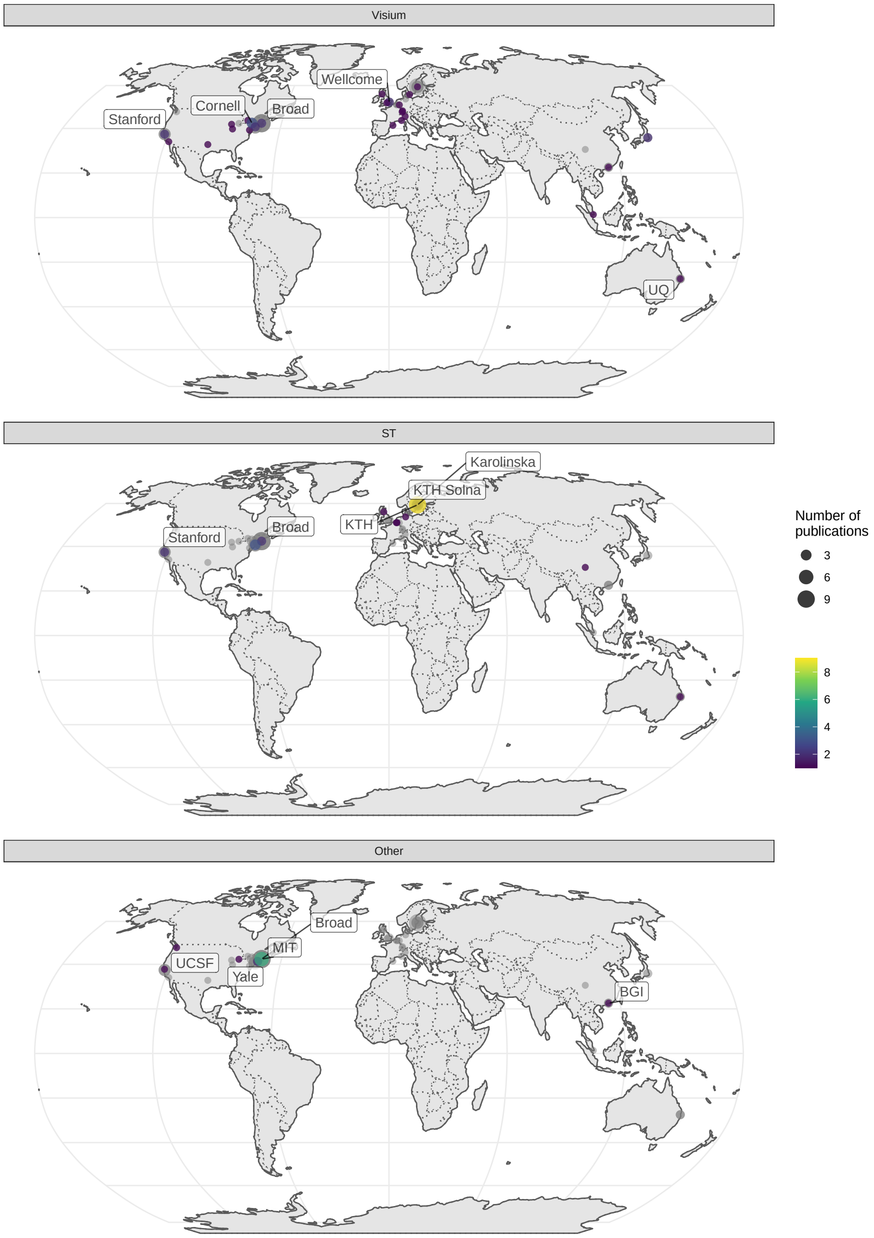

**Figure 5.30:** Cities and institutions using ST and Visium. Preprints are included.

ST (Ståhl et al. 2016) and Visium are the most widely used current era technique after LCM (Figure 4.7, Figure 5.30). In ST, the printed spots have diameter of 100 $\mu$m and are 200 $\mu$m apart from center to center (Figure 5.31). Multiple sections can be mounted to the same slide, separated by a rubber mask. For each section, there are 1007 spots covering an area of 6200 $\times$ 6600 $\mu$m. The 5' end of the capture sequence is a linker to be cleaved to release the transcripts, followed by amplification and sequencing handle, an 18 nt spatial barcode, a 9 nt UMI, and poly-T (Figure 5.32). For the genes quantified with smFISH, ST's detection efficiency is around 6.9% compared to smFISH, within the range of the efficiency of scRNA-seq techniques. Despite the low resolution, ST is popular probably due to transcriptome wide profiling, ease to apply to larger area of tissue, not requiring specialized equipment such as SRM and custom flow cells, commercial kits, possible automation of library preparation (Jemt et al. 2016), availability of a documented and open source data preprocessing pipeline called ST Pipeline (Navarro et al. 2017), and the extra information from H&E staining before library preparation.

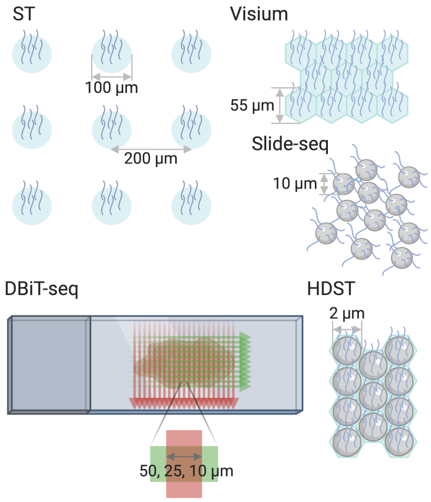

**Figure 5.31:** Schematic of spot construction and size of array based techniques.

After its inception, ST has been used in a wide range of clinical pathological tissues, such as heart after heart failure (Asp et al. 2017), peritonitis-affected gingival tissue (Lundmark et al. 2018), prostate cancer (Berglund et al. 2018), breast cancer (He et al. 2020), arthritic joint biopsies (Carlberg et al. 2019), lymph nodes affected by melanoma metastasis (Thrane et al. 2018), spinal cords (Maniatis et al. 2019) and cerebellums (Gregory et al. 2020) affected by amyotrophic lateral sclerosis (ALS), and squamous cell carcinoma (Ji et al. 2020). ST has also been used to construct gene expression atlases of healthy

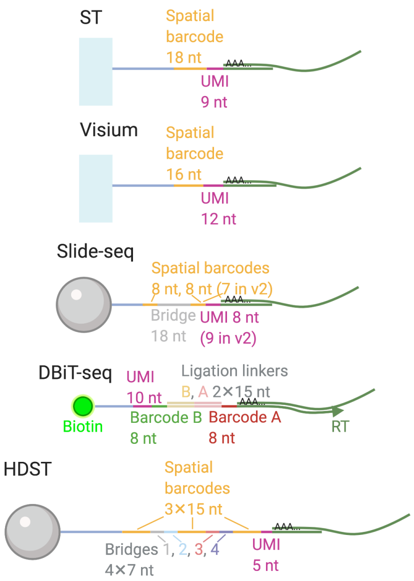

**Figure 5.32:** Barcode and UMI structure and lengths of array based techniques.

tissues such as the developing human heart (Asp et al. 2019) and the mouse brain (Ortiz et al. 2020). In addition, ST is the only current era technique other than LCM and manual microdissection that has been adapted to plants (Giacomello et al. 2017). Common downstream data analyses include identifying differentially expressed (DE) genes between diseased and healthy regions, gene set enrichment analysis (GSEA) among DE genes, and cell type deconvolution of the spots by integrating ST and scRNA-seq data. Data analysis methods designed specifically for ST or Visium will be reviewed in more detail in Chapter 7.

Although introduced fairly recently, after 10X Genomics acquired ST in December 2018, the 10X Visium has quickly gained popularity and spread to multiple institutions, and is used by many studies that utilize an array method in late 2020 and 2021 (Figure 5.30, Figure 5.33). While usage of ST seems concentrated in Sweden, where ST comes from, usage of Visium is more decentralized (Figure 5.30). Visium is similar to ST and shares the advantages of ST, but with higher spatial resolution. The spots are tiled hexagons, each with a diameter of 55 $\mu$m (Figure 5.31). After adjusting for spot area, Visium seems to capture somewhat more transcripts and genes compared to ST (Y. Liu et al. 2020). In addition, Visium's growth in popularity may also be due to core facilities at multiple institutions providing Visium services ("10X VISIUM SPATIAL TRANSCRIPTOMICS" n.d.; "ADVANCED GENOMICS CORE PRICING" n.d.; "SpaRTAN" n.d.). As a new version of ST, Visium was originally designed for fresh frozen OCT embedded tissue and 3' Illumina sequencing. However, Vi-

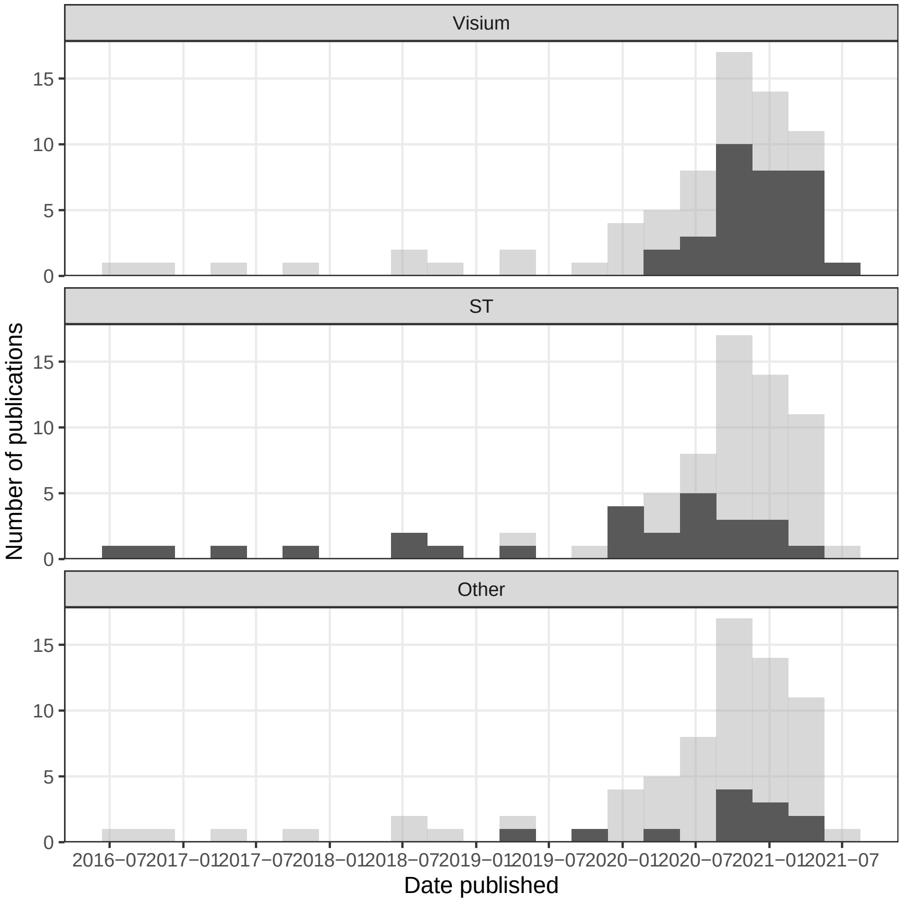

**Figure 5.33:** Number of publications over time, broken down by technique. The facets are ordered by recent usage of the technique. Bin width is 90 days.

sium has more recently been adapted to FFPE tissue (Villacampa et al. 2020), as well as to Nanopore long read sequencing to quantify isoforms (Lebrigand et al. 2020; Joglekar et al. 2020).

In response to the low resolution of ST, Slide-seq was developed to increase the resolution of array based spatial transcriptomics (Rodriques et al. 2019). Beads like those used in Drop-seq (Macosko et al. 2015) with diameter 10 $\mu$m are spread on a slide in a single layer, not necessarily in a regular grid, and bead barcodes are generated with 16 rounds of split pool, each round adding one nucleotide, broken into 2 blocks of 8 nt (2 blocks of 8 and 7 nt in version 2) (Figure 5.31, Figure 5.32). As the location of each barcode is not pre-determined, the slide is imaged and the barcodes are sequenced in situ with SOLiD. Then the OCT frozen tissue section is mounted on the layer of beads on the slide and the beads are removed for library preparation. The first version of Slide-seq is very inefficient; for the genes compared, the Slide-seq only detects 2 to 3 orders of magnitude fewer transcripts per cell than smFISH and about 2.7% compared to Drop-seq (Rodriques et al. 2019).

In the second version of Slide-seq (Slide-seq2) (Stickels et al. 2020), the barcodes are sequenced by SEDAL (like in Figure 5.29, but with one color per base) rather than SOLiD, which increased the efficiency of spatial mapping of Illumina reads, probably because of error propagation in the 2 base encoding of SOLiD. Moreover, bead synthesis is further optimized and a second strand synthesis step is added to the library preparation to increase the number of cDNAs for PCR amplification. Efficiency is improved in Slide-seq2, which is ~9.3x higher than version 1, about on par with Drop-seq, 1 order of magnitude lower than that of smFISH, and somewhat better than Visium in the dataset chosen. The official software to process the in situ sequencing images is written in MATLAB, which is proprietary. Although the size of the bead is close to the size of a single cell, Slide-seq does not have single cell resolution as one bead can capture transcripts from more than one cells nearby, so cell type deconvolution of beads is still needed. After its inception, Slide-seq2 has been used on mouse and human testes, at the institution of origin (Chen et al. 2020).

Spatial resolution of array based techniques has been further increased with HDST, with a resolution of 2 $\mu$m (Vickovic et al. 2019), which is smaller than a single cell. Like in Slide-seq, beads like those used in droplet scRNA-seq are used. The diameter of each bead is 2 $\mu$m, and hexagonal wells with diameter 2.05 $\mu$m are carved into a slides so each well contains one bead (Figure 5.31). The spatial barcodes are generated by 3 rounds of split-pool, each round adding 15 nt from the barcode pool (Figure 5.32). The UMI is only 5 nt but such a small area does not contain that many transcripts. As the beads are randomly placed in the wells, the locations of barcodes need to be determined. Four rounds of FISH, with combinations of red, green, and no color, encode each of the 3 barcodes on each bead. Again, HDST was originally designed for fresh frozen OCT embedded tissue rather than FFPE. HDST is very inefficient; for the genes compared, the detection efficiency is only ~1.3% compared to smFISH per bead. To our best knowledge, HDST has not been used for new datasets after its inception.

In response to the low efficiency and complicated procedure to localize barcodes of Slide-seq and HDST, Deterministic Barcoding in Tissue for spatial omics sequencing (DBiT-seq) was developed, with resolution up to 10 $\mu$m (Y. Liu et al. 2020). Let $i, j$ denote the index of channel in each direction. Barcode $A_i$, attached to poly-T, is flown across the slide in microfluidic channels and RT is performed (Figure 5.31). Then barcode $B_j$, attached to the UMI, PCR handle, and biotin, is flown across the slide in microfluidic channels perpendicular to those that delivered barcode $A_i$, and barcode $B_j$ is ligated to barcode $A_i$ and the cDNA (Figure 5.31, Figure 5.32). Then the ligated barcodes and cDNA can be purified by streptavidin-coated magnetic beads. Each microfluidic channel carries a different barcode, so where the channels for barcodes $A_i$ and $B_j$ intersect, an array is created and the location of each spot is encoded by $i, j$. The resolution is limited by the width of the channels and the spacing between them; widths of 50, 25, and 10 $\mu$m have been tested. Per unit spot area, DBiT-seq seems to detect at least 3 times more genes and UMIs than ST and Visium and the improvement is even starker at the 10 $\mu$m resolution. For the genes compared, DBiT-seq's detection efficiency is ~15.5% of that of smFISH, making it relative more sensitive among the array based methods reviewed here. To our best knowledge, DBiT-seq has not been used for new datasets after its inception.

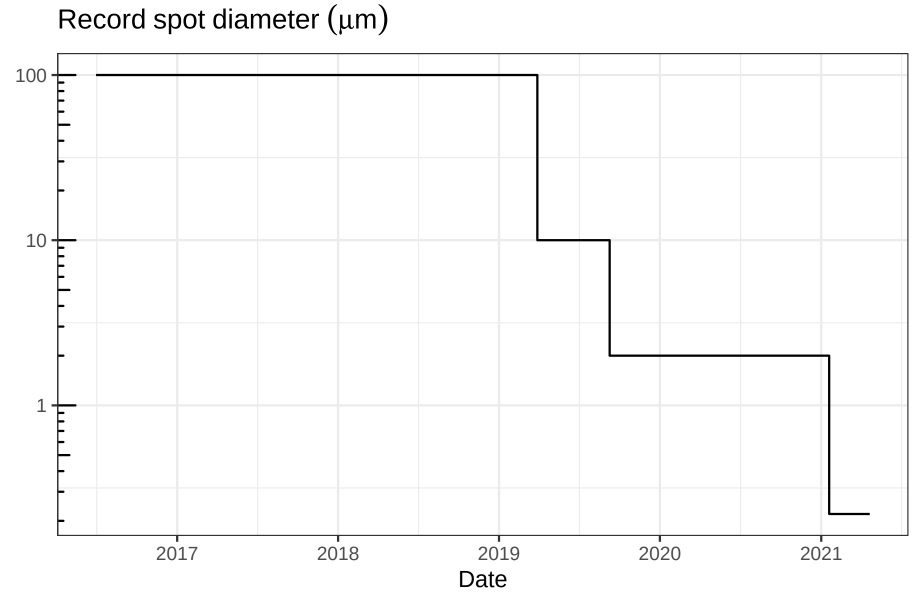

**Figure 5.34:** Record spot diameter of array based methods over time.

The record resolution of array based techniques is ever increasing (Figure 5.34); sub-micron techniques are appearing in 2021. The record is broken by Stereo-seq in January 2021, reporting a spot diameter of 220 nm although the distance between spots is 500 or 715 nm (A. Chen et al. 2021). In Stereo-seq, circularized DNA containing a random 25 nt barcode is RCA amplified and deposited into an etched grid. The barcode is sequenced and then oligos with polyT and molecular ID are hybridized to the barcode to capture polyA transcripts from the mounted tissue. The reported capture efficiency is around 170 transcripts per 100 $\mu m^2$ in mouse brain, on par with that of the Visium mouse brain dataset from the 10X website reanalyzed in the same study.

Another sub-micron array capture method is Seq-Scope (Cho et al. 2021), which creates clusters of polyT capture sequences each with its own spatial barcode (20-32 nt) from Illumina bridge amplification on a repurposed Illumina flow cell.

The spatial barcode is sequenced with SBS. Then the flow cell is dismantled so the tissue can be mounted for transcript capture. The captured transcripts are then sequenced with NGS. The clusters can have a diameter down to 0.5 $\mu$m, and the clusters are randomly seeded, not distributed in a grid. The reported capture efficiency is around 1000 and up to 2000 transcripts per 100 $\mu m^2$ in mouse colon, much higher than that of Stereo-seq, although we are not sure whether colon data is comparable to brain data here.

A more recent nearly sub-micron technique is PIXEL-seq (Fu et al. 2021). Again, as in the Illumina flow cell, PIXEL-seq amplifies each randomly seeded spatial barcode (24 nt) and polyT capture sequence into polonies. However, here a crosslinked polyacrylamide gel (rather than a linear one in Illumina) is used, to form continuous polonies without much space between their "territories" rather than discrete clusters. The spatial barcodes are also first sequenced with SBS before the tissue is mounted for transcript capture. On average, the polony is around 1.17 $\mu m^2$ in area, so assuming it is circular, then the diameter is 1.22 $\mu$m. The reported capture efficiency is around 1000 transcripts per $\mu m^2$ in mouse brain, which might be comparable to that of Seq-Scope.

While such sub-micron techniques have subcellular resolution, in practice, the data is binned into much larger grids for standard scRNA-seq analysis, such as $36\mu m \times 36\mu m$ in Stereo-seq and $10\mu m \times 10\mu m$ or $7\mu m \times 7\mu m$ or $5\mu m \times 5\mu m$ in Seq-Scope. The subcelluar information was not directly use in the analyses, although even with binning, the resolution is still higher than that of ST and Visium.

In summary, a putative ranking, from high to low, of capture efficiencies of current era techniques is:

smFISH (~100%) > MERFISH (HD4, ~95%) > HCR-seqFISH (~86%) > ExSeq (targeted, 62%) > seqFISH+ (~49%) > (maybe) Seq-Scope ~ PIXEL-seq > (maybe) DBiT-seq (~15%) ~ Visium ~ Stereo-seq > (maybe) HybRISS > HybISS ~ (maybe) STARmap ~ (maybe) scRNA-seq ~ slide-seq2 ~ ST (~6.9%) ~ ISS (~5%) > HDST > slide-seq1 > FISSEQ

This is putative because this is based on reports in the main text. There are conflicting reports of capture efficiency of Visium and DBiT-seq. Furthermore, comparison of different tissues and different genes from those studies may be problematic. For some of the technologies, the capture efficiency is compared to that of smFISH with only a few genes. Multiple datasets from each technology for as similar a tissue as possible for the same set of genes should be compared to get a better idea about the capture efficiency of each technique. Moreover, other factors such as tissue handling, sequencing depth, and data processing software may influence the results.

All these array based techniques reviewed so far capture polyadenylated transcripts. While miRNAs form a major topic in LCM literature (Figure 6.3) and are profiled in some prequel era ISH atlases, current era techniques mostly preclude miRNA quantification. For smFISH based techniques, miRNAs are way too short to accommodate the large number of probes, even with signal amplification. While RCA has been adapted to miRNA, to our best knowledge, it has not been demonstrated to show individual miRNAs as discrete puncta like in smFISH and ISS, nor has it been demonstrated in a highly multiplexed fashion (Neubacher and Arenz 2009; Kim, Kim, and Kim 2020; Zhou et al. 2020). Without a poly-A tail, miRNAs are precluded by the other array based techniques as well. To quantify miRNAs in space, an array based technique was developed as an alternative to LCM and designed for FFPE tissues (Nagarajan et al. 2020). The tissue is pixelated, and each pixel is 300 $\mu$m $\times$ 300 $\mu$m. Within each pixel is a smaller 3 $\times$ 3 array, each spot of which has probes for one miRNA; The locations of the spots within each pixel can be easily discerned with a fluorescent microscope. This way, up to 9 miRNAs can be profiled in the same tissue section at the same time, although the 9 miRNAs are from somewhat nearby cells but not the same cells.

#### 5.5 No imaging

The techniques reviewed above, involve either imaging (e.g. LCM, smFISH, ISS, Slide-seq, and HDST) or prior knowledge of locations (e.g. Tomo-seq, ST, Visium, and DBiT-seq). Some spatial transcriptomics techniques have been developed that require neither imaging nor prior knowledge of locations, and we review these in this section.

It is possible to reconstruct relative locations of cells or transcripts from colocalization without imaging, albeit imperfectly. These techniques are reviewed in more details in (Boulgakov, Ellington, and Marcotte 2020); we will only briefly summarize techniques that do not require DNA bound to a surface so they can be applied in cells and tissues. An early method to do so is Puzzle Imaging, published in 2015 (Glaser et al. 2015). Here "colocalization" can mean whether two neurons have axons in the same voxel or whether two neurons are synaptically connected. The spatial reconstruction is framed as a dimension reduction problem; each voxel is represented as a vector with $n$ dimensions, where $n$ stands for the number of neurons, and these vectors are to be projected into 2 or 3 dimensions, representing spatial dimensions, for reconstruction. Puzzle Imaging was only demonstrated in synthetic datasets, but not real biological datasets. Such reconstruction was made possible for transcripts with DNA microscopy (Weinstein, Regev, and Zhang 2019) Transcripts are reverse transcribed in situ, and the cDNA, with an UMI added, is PCR amplified in situ. The amplified products diffuse and encounter amplified products from other transcripts. The nearby cDNAs are concatenated with overlap extension PCR, with additional random sequences in the overlapping primers to encode each concatenation event, called unique event identifier (UEI). When the concatenated cDNAs are sequenced, the two UMIs and the UEI are recorded. Because amplified products from two nearby transcripts are more likely to be concatenated than those from two transcripts that are far apart, the number of UEIs between two UMIs can be used to reconstruct relative distance between transcripts.

Techniques have also been developed to quantify transcripts from subcellular compartments, such as APEX-RIP (Kaewsapsak et al. 2017) and APEX-seq (Fazal et al. 2019). Although these techniques do not record or reconstruct spatial coordinates, they are included in this review because the publications describing them described them with terms such as “spatial”, “localization”, and “spatial transcriptome”. APEX is an engineered ascorbate peroxidase, which can be targeted to specific cellular compartments by expressing a fusion of APEX and a protein targeted to the compartment of interest. With substrates $H_2O_2$ and biotin-phenol (BP), APEX catalyzes formation of biotin-phenoxyl radicals that can biotinylate nearby proteins, which can be isolated with streptavidin. In APEX-RIP, mRNAs are cross linked to nearby proteins and thus isolated after isolating biotinylated proteins. In contrast, in APEX-seq, the mRNAs are directly biotinylated. Compared to APEX-RIP, APEX-seq better discerns transcript localization in compartments not bound by membrane. However, both APEX-RIP and APEX-seq were originally designed for bulk rather than single cell samples and was tested only on cell culture. Also, because a fusion protein is required, they cannot be performed in human tissue sections.

Rare cell types are difficult to characterize with most spatial transcriptomics techniques. ST and Visium lack single cell resolution and signal from rare cell types may be diluted by signal from common cell types in the same spot. LCM is still typically not used on single cells and rare cell types may or may not be easily discernible with H&E. SmFISH based techniques and targeted ISS require a pre-defined panel of genes, often selected from scRNA-seq and well-known markers, but such selection is more challenging for rare cell types, which may not be well-studied enough to begin with due to challenges in other transcriptomics techniques. However, spatial pattern of genes expressed in rare cell types can be characterized by deliberately creating doublets or multiplets involving both common and rare cell types, as in paired cell sequencing (Halpern et al. 2018) and ClumpSeq (Manco et al. 2020). Spatial patterns of genes expressed in common cell types such as hepatocytes and small intestine enterocytes are already known from smFISH or LCM and spatial reconstruction of scRNA-seq data (Halpern et al. 2017; Moor et al. 2018). Genes expressed in the rare cell types are identified from genes much more highly expressed in the multiplet than in individual cells from common cell types in scRNA-seq, or markers of rare cell types from scRNA-seq if such data exists. Then the multiplets are mapped to spatial locations with patterned genes expressed by common cell types and existing smFISH or LCM data as reference. Then rare cell types and their characteristic gene programs are mapped to spatial locations as well and their patterns can be characterized without directly imaging these cells.

#### 5.6 Spatial multi-omics

Some spatial transcriptomics techniques have been adapted to collect data of other modalities, such as proteomics, neuron projection, and 3D chromatin conformation. These modalities can give a fuller picture of cell state than transcriptomics alone. In both MERFISH (Wang, Moffitt, and Zhuang 2018) and GeoMx DSP (Merritt et al. 2019), a panel of proteins can be quantified with oligonucleotide tagged secondary antibodies, and the oligo tag is detected and counted as spots just like mRNA. More recently, oligo tagged antibodies are also incorporated into ST as SM-Omics (Vickovic et al. 2020). We have already mentioned adaptation of MERFISH targeting introns and genomic DNA to determine 3D chromatin conformation (M. Liu et al. 2020; Su and Song 2020), and pseudocolor seqFISH and seqFISH+ have been used for this purpose in cell culture as well (Shah et al. 2018; Takei et al. 2020).

In MERFISH, traditional Nissl or poly-A based staining miss cellular processes, but neuron projection tracing can be performed prior to MERFISH. In the mouse motor cortex MERFISH atlas (Zhang et al. 2020), axons are first visualized by injecting cholera toxin subunit b (CTb) conjugated to 3 different dyes into 3 cortical areas as a retrograde tracer, tracing from terminals of the axons to the cell bodies. After imaging the axons, transcripts are imaged and quantified with MERFISH so neuronal projection can be related to the transcriptome. Viruses can be used for anterograde tracing, i.e. from cell bodies to axon terminals (Xu et al. 2020), and can in theory be performed prior to MERFISH imaging. Axonal projections are traced in BARseq(2). Electrophysiological recordings from cultured cardiomyocytes in space has been coupled to STARmap with electro-seq; the recording is performed before the cells are fixed and cleared for STARmap (Li et al. 2021).

#### 5.7 Databases of the current era

The database holding various spatial gene expression data was proposed early in the prequel era (1990s), when enhancer and gene trap data was proliferating and major WMISH atlas projects were in progress. In contrast, in the current era, databases only emerged after datasets from various techniques have already proliferated. One of such databases is SpatialDB (Fan, Chen, and Chen 2020), published in late 2019, which holds gene count matrices from ST, LCM, Tomo-seq, and etc. and spatially variable genes identified with SpatialDE (Svensson, Teichmann, and Stegle 2018) and trendsceek (Edsgärd, Johnsson, and Sandberg 2018). In addition, the SpatialDB website provides interactive visualization of gene expression in space. Data can be queried by gene symbols, species, and data collection techniques.

Another database by the Brain Research through Advancing Innovative Neurotechnologies (BRAIN) Initiative - Cell Census Network (BICCN) is under construction as of late 2020 (Adkins et al. 2020). This is an international collaboration providing and generating multi-modal data for the mouse, human, and non-human primate brain, collected with scRNA-seq, ATAC-seq, neuron projection tracing, MRI, IHC, MERFISH (Zhang et al. 2020), osmFISH, seqFISH, and etc. The database website is hosted by the Allen Institute, and thus may be considered a continuation of the ABA. Data can be queried by species, technique, modality, and the lab that generated the data, but not by gene symbols.

While current era mouse brain atlases still reference the prequel ABA ontologies (Ortiz et al. 2020; Chen et al. 2020; Vickovic et al. 2020; Lohoff et al. 2020), data cannot be queried by ontology in the current era databases, nor by a reference gene expression pattern as in the prequel database FlyExpress (Kumar et al. 2017). With more quantitative and comprehensive data, the traditional ontology may need to be revised. Unlike prequel databases such as ABA, EMAGE, and FlyExpress, to the best of our knowledge, current era spatial data has not been systematically registered to a 3D model for integrative analysis across datasets and for visualization.

### Chapter 6

### Text mining LCM transcriptomics abstracts

To analyze trends in LCM followed by microarray or RNA-seq, abstracts were downloaded from the PubMed API, with search term `"((laser capture microdissection) OR (laser microdissection)) AND ((microarray) OR (transcriptome) OR (RNA-seq))"`. For preprints, abstracts from the search term "laser microdissection" were downloaded from bioRxiv. Because bioRiv's advanced search does not acknowledge parentheses, a more complicated search term was not used. Upon random inspection, the retrieved abstracts mostly seem relevant. The number of LCM transcriptomics search results dwarfs the number of publications for other methods of spatial transcriptomics and seems to show two peaks, one around 2012, and the other in 2020 and 2021 (Figure 6.1); the LCM corpus contains 2252 abstracts as of March 26, 2021, while there are between 500 and 600 papers in the curated database.

LCM transcriptomics is also more geographically diffuse and spread out into many less well-known institutions and some developing countries, though some elite institutions are among the top contributors, such as Harvard Medical School and Massachusetts General Hospital (Boston), Columbia University, NYU, Rockefeller, and Sloan-Kettering (New York), NIH (Bethesda), and Cambridge University (Cambridge, UK) (Figure 8B).

After identifying common and relevant phrases in the abstracts, the abstracts were tokenized into unigrams. We used the `stm` R package (Roberts, Stewart, and Tingley 2019) to identify topics. The cities in which the research was conducted, date published or posted on bioRxiv (linear, not transformed), and journal (including bioRxiv) were used as covariates for topic prevalence, because labs and journals may have preferred topics and city is a proxy to institution, and it's reasonable to assume that prevalence of at least some topic changes through time, such as due to evolution of technology. Cities and journals with fewer

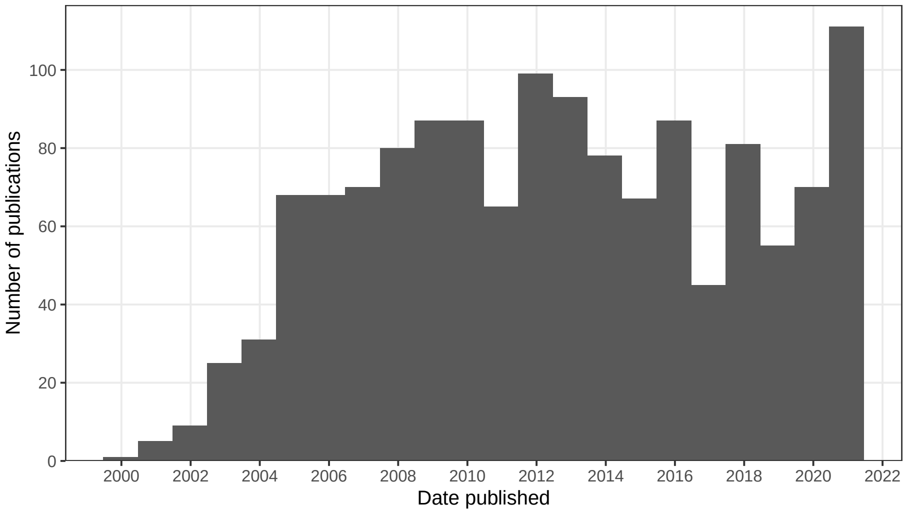

**Figure 6.1:** Number of publications in LCM transcriptomics PubMed search results over time. Bin width is 365 days.

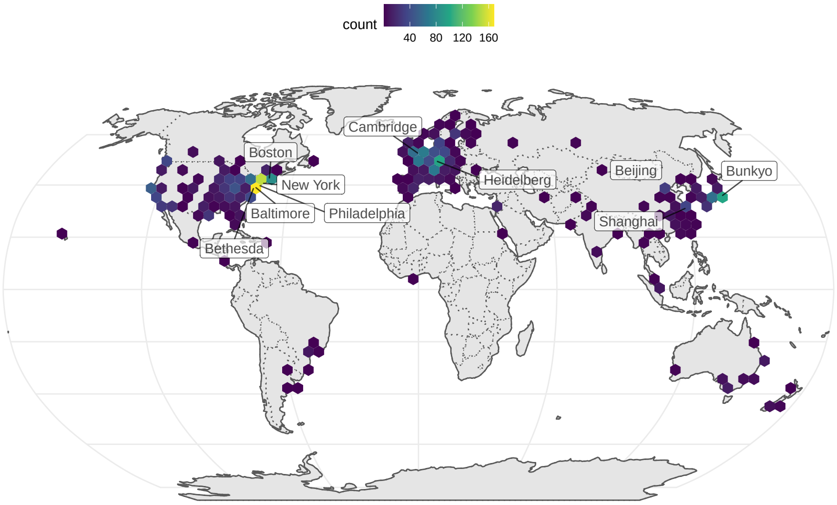

**Figure 6.2:** Geographic distribution of LCM transcriptomics research, with top 10 cities labeled. Number of publications is binned over longitude and latitude.

than 5 papers were lumped into "Other". From a trade off between held out likelihood and residual, and between topic exclusivity and semantic coherence, we chose 50 topics. Code used to find this can be found here[1].

Here `stm` stands for structural text mining. A generative model of word counts is fitted with the word counts in each abstract as well as abstract level covariates, here date, city, and journal. Among parameters of the model estimated are the proportion of each topic in each abstract after accounting for covariates ($\theta$), topic proportions in the corpus ($\gamma$), and probability of getting each word from each topic ($\beta$). See the `stm` vignette[2] for more details. `stm` can not only detect topics without having a human read all the abstracts, but also find how covariates relate to topic prevalence.

#### 6.1 Topic modeling

As already mentioned, microarray was first demonstrated on LCM samples in 1999, profiling 477 cDNAs from rat neurons (Luo et al. 1999). Since then, LCM transcriptomics has been used on many research topics, such as various aspects of cancer (topics 5, 6, 8, 10, 11, 13, 16, 20, 24, 27, 34, 44, 50), botany (topics 9, 15, 21, 40, 43, 45), developmental biology (topics 1, 3, 17, 18, 29, 35, 39), neuroscience (topics 7, 14, 19, 23, 25, 32, 33, 36, 47), immunology (topic 12, 22, 48), miRNA (topic 5), and technical issues related to LCM (topics 4, 28, 37, 41) (Figure 6.3).

In most cases, the top 5 words in each topic give us a decent idea what the topic is about. We can also plot the probability to get top words ($\beta$) in each topic.

While in most cases, the topic is apparent from the top words, some topics are less apparent (e.g. topic 49). From the top words and quick glances of abstracts with the highest proportion of each topic, the 50 topics are summarized here in more human readable terms:

1. Stem cell and fetal development
2. GWAS, genetic screens, and genetics of complex phenotypes
3. Biomechanics, ECM, eye lens, muscles, and morphogenesis
4. Data analysis, especially of RNA-seq, but also of 3D genome structure and microarray
5. miRNAs in cancer
6. Quantitative analyses of cancer, clinical and bioinformatic
7. Hippocampus and Alzheimer's disease, sometimes related to Down syndrome

[1] `https://github.com/pachterlab/museumst/blob/master/data-raw/lcm_text_mining.Rmd`

[2] `https://cran.r-project.org/web/packages/stm/vignettes/stmVignette.pdf`

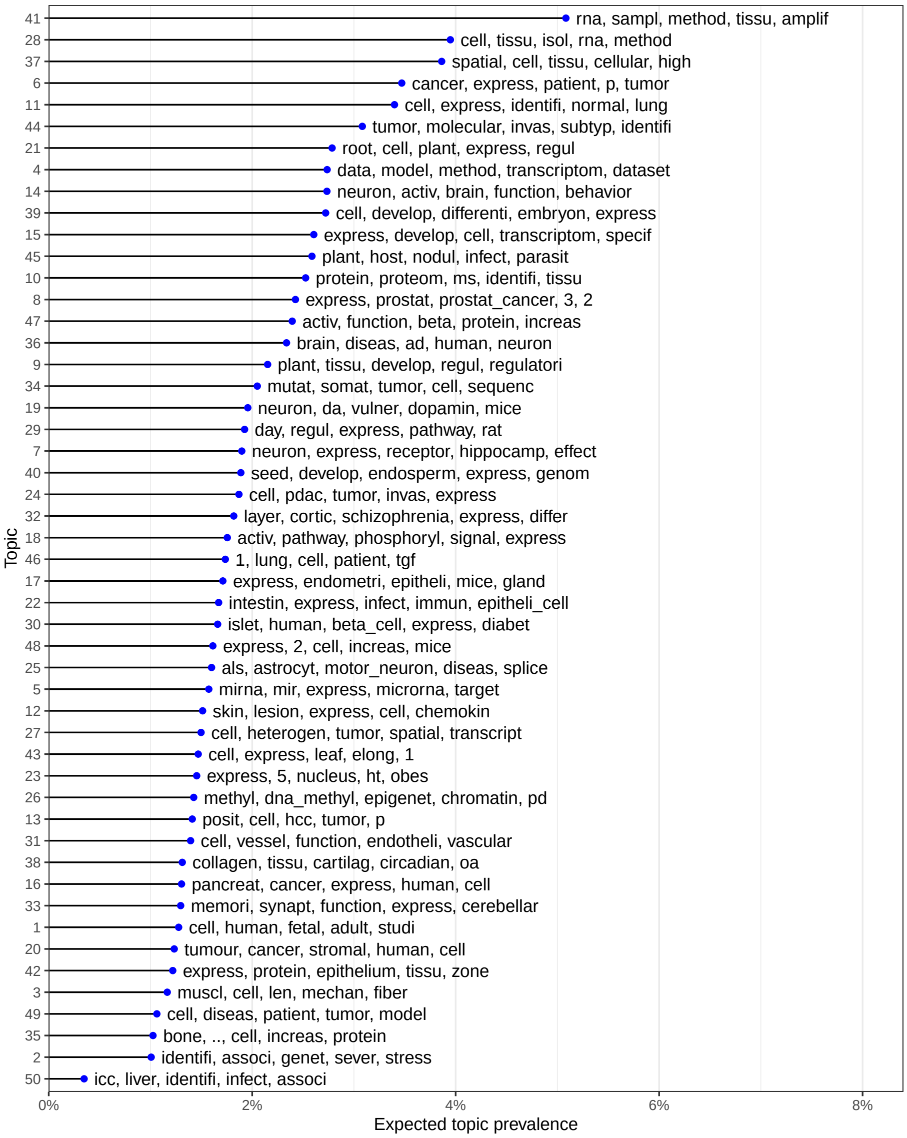

**Figure 6.3:** Top words for each of the 50 topics, ordered by expected topic prevalence and showing top 5 words contributing to each topic.

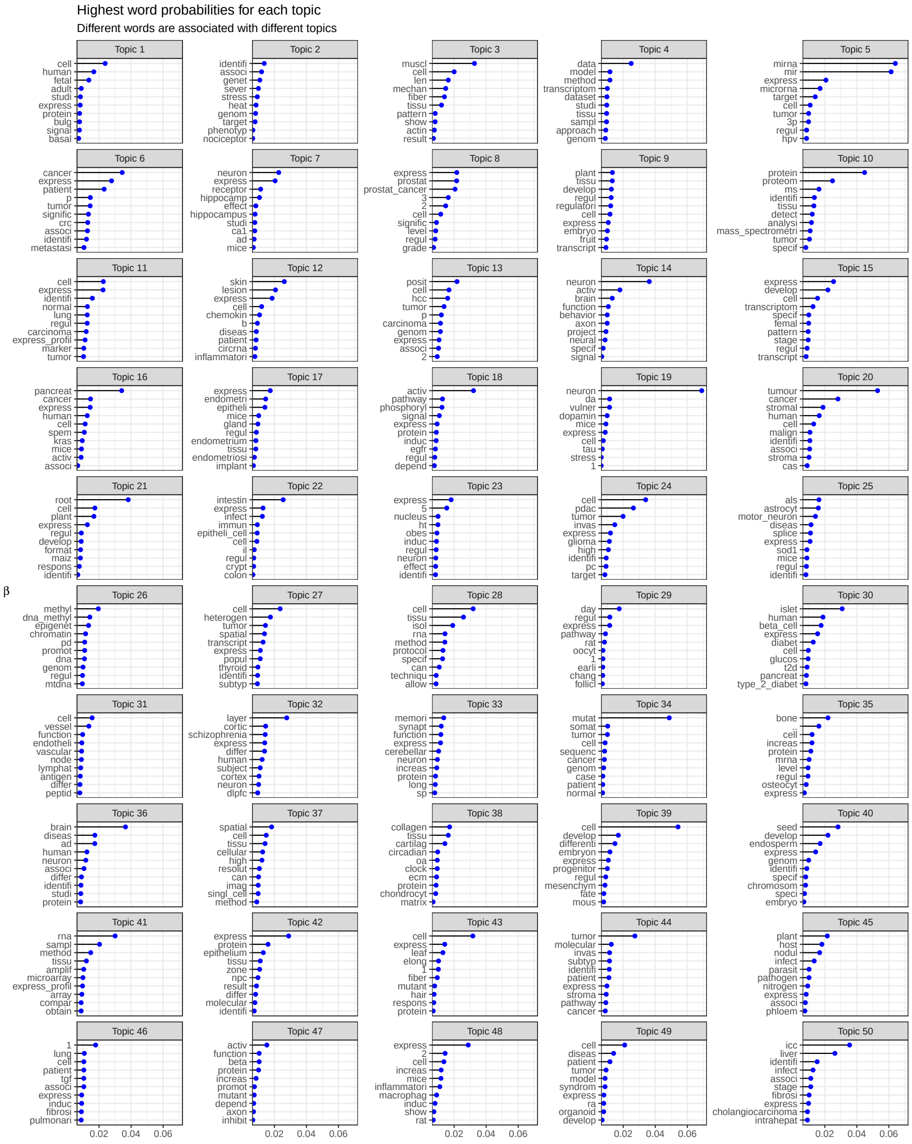

**Figure 6.4:** Probability of top 10 words in each topic.

8. Prostate cancer and other stuff in molecular biology and biochemistry, probably because some prostate cancer papers have an emphasis on molecular biology
9. Plant embryos, plant development, and some stuff about evolution and ecology related to plants
10. Proteomics, especially in cancer
11. Cancer progression and diagnostics, especially lung cancer
12. Inflammation and immunology, especially in skin diseases
13. Breast cancer and liver cancer, with an emphasis in data analysis
14. Neural circuitry, neural plasticity, brain injury, and behavior
15. Plant gamitogenesis and reproduction
16. Spasmolytic polypeptide-expressing metaplasia (SPEM), oncogenes, KRAS
17. Endometrium and implantation. Somehow the top 2 entries are about hearing loss. Why? Epithelium?
18. Cell cycle, also hepatic zonation and circadian rhythm (the latter is also a cycle)
19. Neurons, especially dopaminergic
20. Tumor stroma and microenvironment
21. Plant roots
22. Intestine, especially microbiome and immune response
23. Hypothalamus, obesity, and appetite
24. PDAC, and some stuff about glioma and prostate cancer
25. ALS, and other neurodegenerative diseases affecting motor neurons
26. Epigenetics
27. Tumor single cell profiling and cellular heterogeneity
28. Tissue isolation and preparation
29. Bone growth plate, especially recovery after radiotherapy, and some other stuff like oocytes, glaucoma, and epithelial injury
30. Pancreas and diabetes, especially T2D
31. Lymphocytes, lymphatic and blood vessels
32. Prefrontal cortex and schizophrenia
33. Synapses, dendritic spines, neuron potentiation, sometimes related to memory
34. Cancer genomics, mutations, and phylogeny
35. Bone formation, but also some other stuff about cancer and kidneys
36. Neurodegenerative diseases, Alzheimer's, Parkinson's, and multiple system atrophy
37. Spatial single cell techniques and imaging
38. Connective tissues and ECM, and some other stuff about circadian rhythms
39. Stem cells and development
40. Plant seed development and reproduction
41. RNA extraction and amplification, especially in microarray, but also in RNA-seq
42. Lots of different stuff about epithelium

43. Plant leaves, but also other stuff about gamitogenesis
44. Cancer pathway analyses and molecular and cellular mechanisms
45. Plant nitrogen fixation and soil microbiome
46. Lots of different stuff related to fibrosis and fibroblasts, such as in lung diseases and graft rejection
47. Neuron morphogenesis, axon guidance, somehow also angiogenesis, protein signaling
48. Inflammation, immune response, especially in atherosclerosis, though there's some other stuff about blood vessels
49. Model organisms and in vitro model systems
50. Intrahepatic cholangiocarcinoma (ICC)

Some of them might not really be related to LCM (e.g. GWAS), and some seem to be a mixture of different topics recognized by humans but seemingly united by something else in common. There are very likely more than 50 topics present, depending on how a topic is defined. The topics can be broadly categorized into Botany, Cancer, Development, Immunology, Neuroscience, Technical, and Other, though these categories can overlap. Some of the "Other" topics seem like mixtures of multiple topics, such as topic 29, while some are very specific and relevant, such as topic 30 (pancreas and diabetes). The broad categories will be used in further analyses.

Clusters of related topics can be seen in the topic correlation plot. See documentation of `topicCorr` in the `stm` package[3] for more details. Here we use a high-dimensional undirected graphical (HUGE) model (Zhao et al. 2012) to estimate the topic correlation graph. The topic proportions ($\theta$) are assumed to be multivariate Gaussian, and HUGE tries to identify edges connecting topics that are not independent from each other conditioned on everything else, while trying to keep the graph sparse (few edges). While $\theta$ is not Gaussian, the results from HUGE aren't unreasonable.

Indeed, cancer, botany, neuroscience, and technical topics tend to cluster together, although this is not the case for immunology and development.

#### 6.2 Changes of word usage through time

We binned dates into years and tested for association of word proportion in each year with the year by fitting a logistic regression model and checking significance of the coefficient for year; word frequency per year since 2001 for the significant words (after Benjamini-Hochberg multiple testing correction) are shown in Figure 6.6. Because too many words are significant, only top 10 from words with decreasing frequency and top 10 with increasing frequency are plotted.

[3] `https://rdrr.io/cran/stm/man/topicCorr.html`

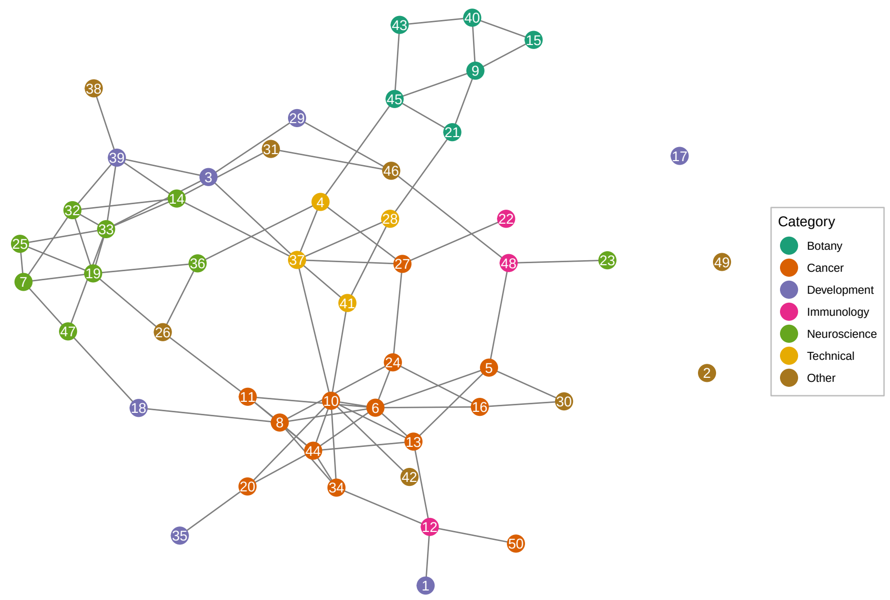

**Figure 6.5:** Correlation between topics.

Here we see that words and phrases associated with microarray and RNA amplification have declined in frequency, while words associated with RNA-seq, single cell, as well as words discussing molecular mechanisms have increased in frequency (Figure 6.6). While transcripts from LCM samples from recent studies were still amplified, the relevant terms decreased in frequency probably because more recent studies, such as ones in the curated database, tend to cite established protocols and kits of library preparation that do the amplification such as Smart-seq2 rather than discussing amplification directly. The “spatial” is associated with current era techniques. Such trends can also be clustered and shown in a heatmap.

Some words have increased in frequency, especially since 2015 (Figure 6.7). Some words sharply decreased in frequency in the early 2000s. However, some words have increased in frequency, peaking in the late 2000s and early 2010s, before declining. Among the terms whose frequency peaked around the early 2010s are “microarray” and “microarray analysis”, perhaps because while RNA-seq was introduced in 2008, microarray did not immediately become obsolete, or perhaps because microarray results are often compared to RNA-seq results, though perhaps wordings changed through the 2000s so the “cDNA” in “cDNA microarray” was omitted (Figure 6.6). Frequency of “real time PCR” also declined, probably because real time PCR was often performed along side microarray but not scRNA-seq to corroborate microarray results (e.g. (Cunnea et al.

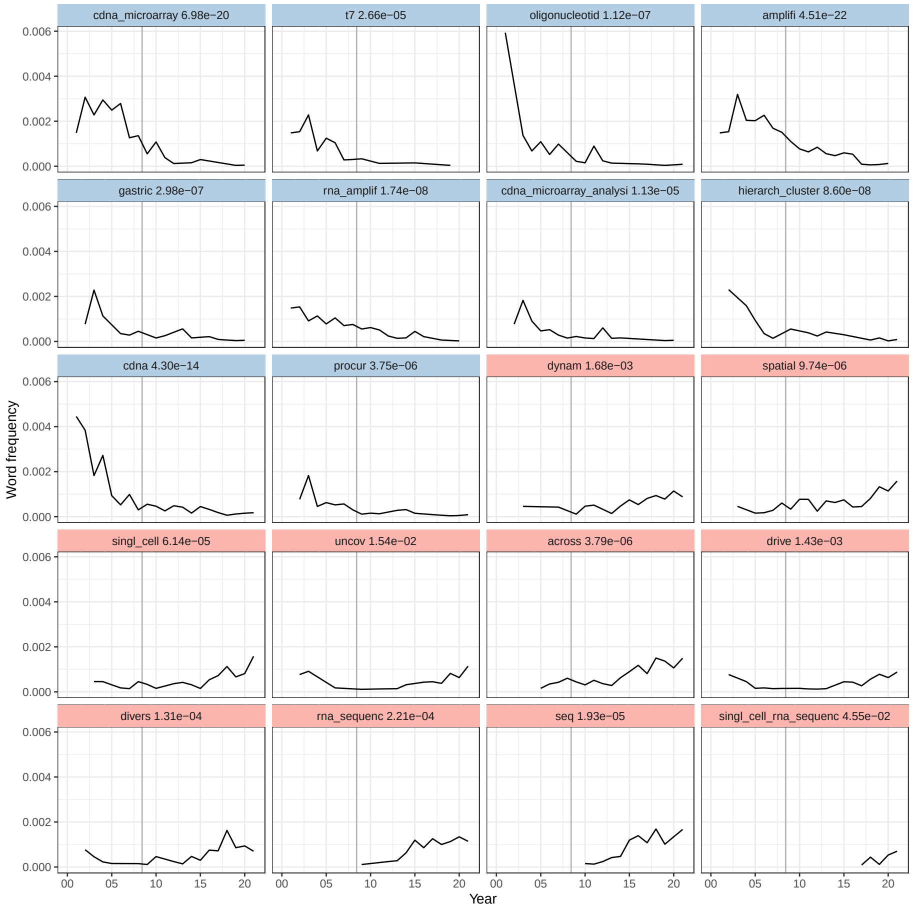

**Figure 6.6:** Word frequency over time since 2001 for words significantly associated with time, sorted from the most decreasing to the most increasing in frequency in time according to the slope in the model. The adjusted p-value of each word is shown. Vertical line marks June 6, 2008, when the first paper about RNA-seq was published (Nagalakshmi et al. 2008).

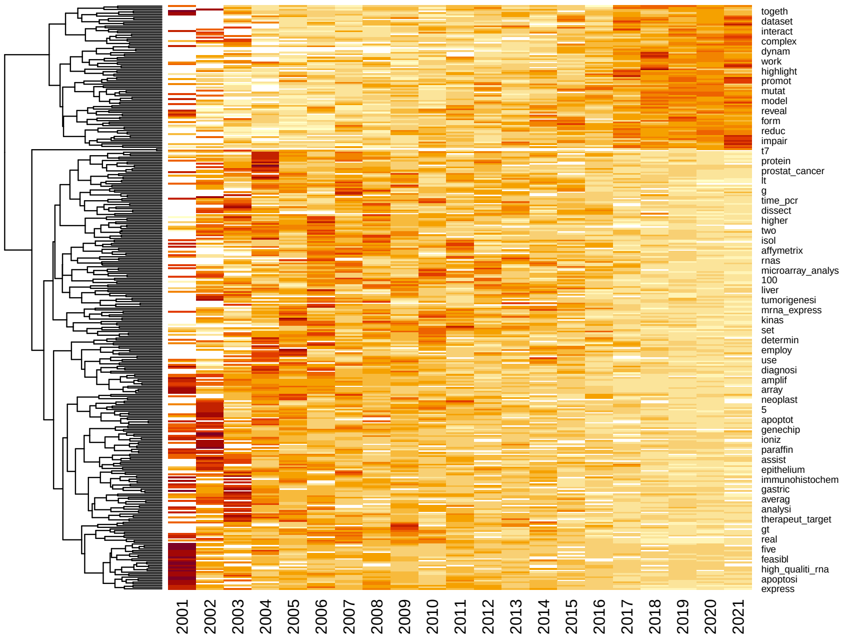

**Figure 6.7:** Heat map clustering changes in word frequency over time. The rows of the matrix are normalized, only showing trend rather than frequency.

2010; Kitamura et al. 2017)), so usage of this term declined with the decline of the cDNA microarray. Besides microarray related terms, some of the words that decreased in frequency are biological terms related to cancer. The "frequency" here is the proportion of all words from all abstracts of a year taken up by a word; the decline in proportion can either be due to decline in interest in the topics that use the word or growth in other topics that don't use the word. This will be explored further in the next section.

#### 6.3 Changes of topic prevalence through time

We tested for association of prevalence of each of the 50 topics with time using the `estimateEffect` function in the `stm` package. Samples of the parameters were taken from the variational posterior of the `stm` model to estimate the variances of the slopes of the linear model of topic prevalence vs. date published, as well as to test whether topic prevalence is significantly associated with time. The p-values of the slopes were corrected for multiple hypothesis testing with the Benjamini-Hochberg method. While the linear model only captures monotonous changes, a more flexible model, such as b-spline transform of the date, was not used because of the modest size of this corpus – on average, each topic has only 45 abstracts, though some topics are larger and some smaller.

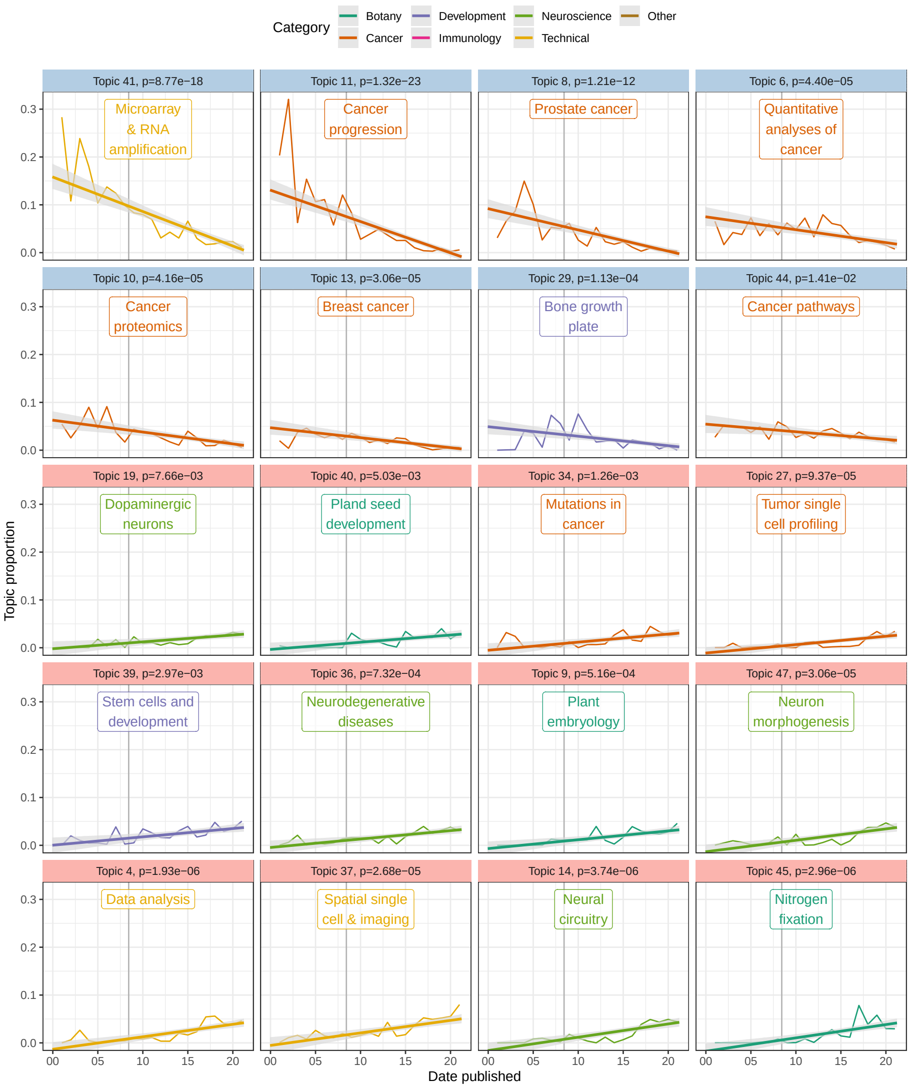

**Figure 6.8:** Topic prevalence over time since 2001 with fitted linear model. Gray ribbon indicates 95% confidence interval (CI) of the slope, estimated from the samples of the variational posterior of the `stm` model. Vertical line indicates advent of RNA-seq in 2008. Light blue facet strip means decreasing trend with adjusted $p < 0.05$, and pink strip means increasing.

As many topics have statistically significant associations with time, only the top 10 most decreasing and top 10 most decreasing topics are plotted here (that's what I intended, but there were only 8 significantly decreasing topics, so top 12 increasing topics are shown). In the early 2000s, a major topic of research about LCM was reliability of T7-based PCR amplification of the small amount of transcripts from samples for microarray, but the prevalence of this topic (topic 41) has declined over time (Figure 6.6, Figure 6.8). The reason for such decline can be a combination of the following: First, other topics in neuroscience and botany emerged and grew (Figure 6.8); some of them are now among the most prevalent topics (Figure 6.3). Second, usage of terms related to microarray and RNA amplification for microarray declined while usage of terms related to RNA-seq increased after 2008 due to the advent of RNA-seq because the latter replaced microarray as the transcriptomics method of choice, so the decline is expected (Figure 6.6). Also as expected, prevalence of topics in data analysis (topic 4) and spatial single cell and imaging technologies (topic 37) increased. Interestingly, cancer topics are among the most significantly decreasing (Figure 6.7, Figure 6.8). Because unlike cDNA microarray, these topics are still relevant today, such decline is puzzling.

Next, we checked whether whether the rise of topics not directly related to cancer may be relevant to the decline of proportions of cancer topics. In `stm`, the abstracts are not hard assigned to topics. Rather, each abstract has a proportion of each topic, and abstracts often have over 90% of one topic. Here, for simplicity, we say an abstract "has" a topic if the proportion of the topic in the abstract is at least 25%.

When the number of abstracts with each topic is plotted, the declines are less drastic or reversed while the increases became much more drastic, especially after 2015, perhaps due to the rise of scRNA-seq, whose library preparation methods made it possible to quantify transcripts from small amount of tissues from LCM (Figure 6.9). These trends don't necessarily correspond to the overall trend across the corpus (Figure 6.1). Then we see in recent years a diversification of topics that may be related to LCM from search results, resulting into decrease of proportion of some older topics the interest in which might not have drastically decreased if not somewhat increased, though not increasing as quickly as other topics. Nevertheless, it is clear that some cancer topics have decreased even in counts. However, remember that some of the `stm` topics seem to be mixtures of multiple topics recognizable by humans and these `stm` topics might have picked up aspects of the abstracts less readily noticed by humans. In other words, it might not be that interest in some cancers decreased per se, but thanks to scRNA-seq, the way these cancers are discussed changed, using words that contributed to other, growing topics. Furthermore, because so many different topics are drastically growing in recent years, the increase in proportion of each of them became less drastic.

Now return to the topic correlation graph, and all the 50 topics, along with their trends, are shown (Figure 6.10). Overall, cancer topics tend to be decreasing in proportion. As already seen in Figure 6.9, this is in part due to growth in non-cancer topics but in part due to decline in some cancer topics. Botany and

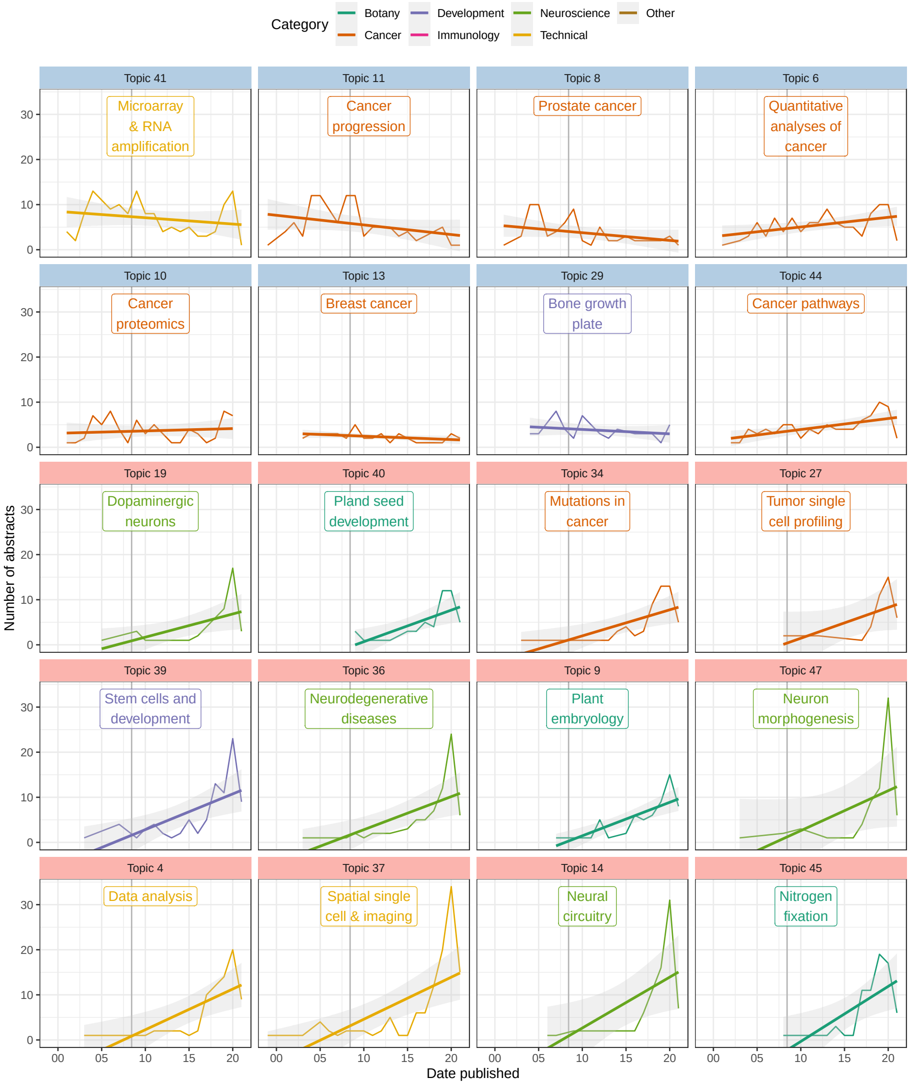

**Figure 6.9:** Number of abstracts with each topic whose proportions changed the most in time. Gray ribbon is the 95% CI of the line fitted to the count per year.

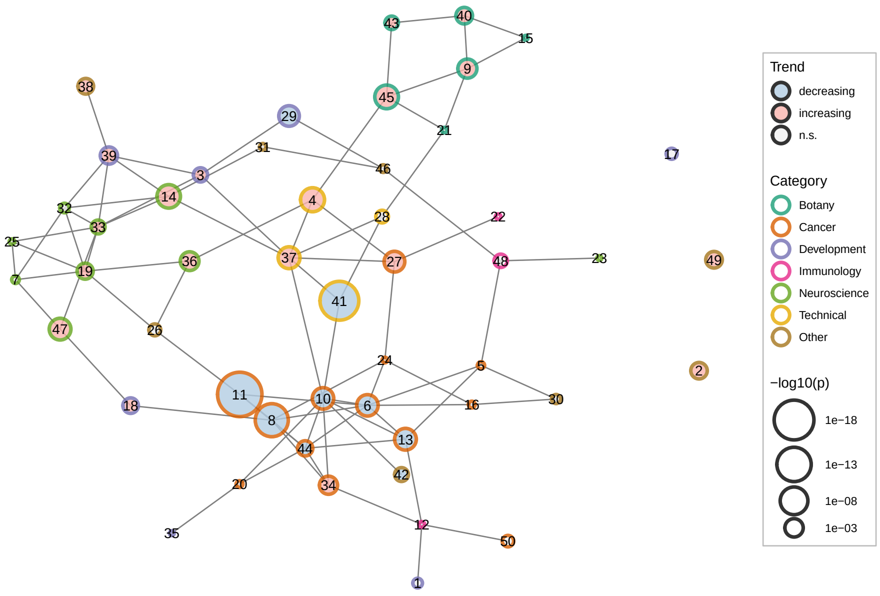

**Figure 6.10:** Correlation between topics colored by both broad categories of the topics and whether its proportion increased, decreased, or did not significantly change (n.s.).

neuroscience topics tend to increase in proportion. This trend is also evident in the topic correlations. Microarray and RNA amplification (topic 41) is correlated with a cancer topic, while spatial single cell and imaging (topic 37) and data analysis (topic 4) are correlated with neuroscience topics. Topic 27, which is about single cell profiling of tumors, has grown, perhaps due to the growth of scRNA-seq. Possibly, as cancer is still relevant, the decline in some cancer topics fed into topic 27 as tumors are examined at the single cell level.

#### 6.4 Association of topics with city

Again, with the `estimateEffects` function, we identify cities associated with certain topics. Some topics might be more spread out, while some some may be confined to a few institutions, which are approximated by city here because it's more difficult to automatically extract institutions from the author address on PubMed than cities. Some institutions might specialize in certain topics. Also note that while for PubMed papers, the cities of the first author are used, because the first author has greater contribution to the paper, only the address of the corresponding author is available from the bioRxiv API. Furthermore, multiple institutions across continents may collaborate on one paper, so the cities here only give a rough idea where LCM related research takes place. Here only the names of the cities are used, with the state and country they are in to distinguish between cities with the same name, without the longitude and latitude, because we don't expect an association between topic and the coordinates in and of themselves, nor do we expect spatial autocorrelation of the topics.

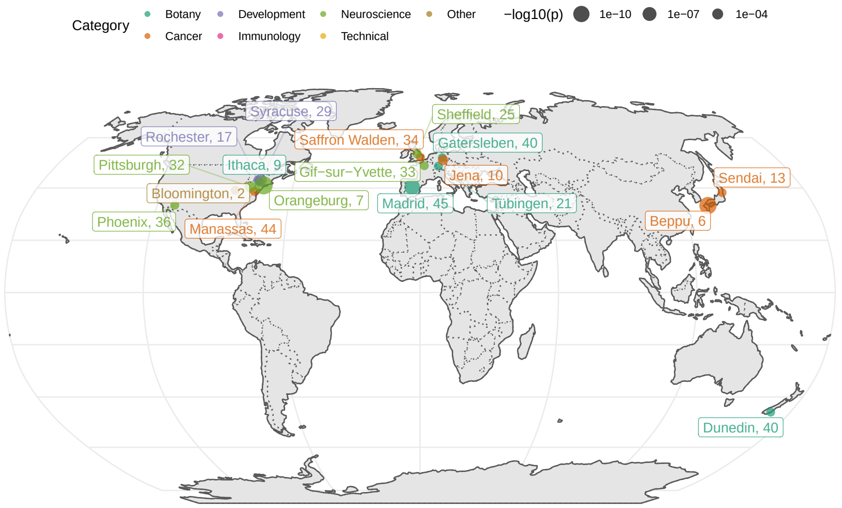

**Figure 6.11:** Cities associated with topics ($p < 0.005$) shown on a map.

Here we note that Center for Dementia Research, Nathan Kline Institute in Orangeburg has greatly contributed to research in hippocampal CA1 pyramidal neurons in Alzheimer's disease and Down syndrome (topic 7) (Figure 6.11). This is the first time I heard of Nathan Kline. Department of Plant Biology at Cornell, Ithaca has greatly contributed to study of plant development (topic 9). Topic 17 is a mixture of topics recognizable by humans; besides the endometrium, some of the top entries are about hearing loss, which come from University of Rochester. George Mason University in Manassas, Virginia contributed several papers about cancer pathway analysis (topic 44). University of Pittsburgh has disproportionate contribution to the study of prefrontal cortex and schizophrenia (topic 32), dating back to 2007. Centro de Biotecnologia y Genomica de Plantas (UPM-INIA), Madrid has disproportionate contribution to the study of soil microbiome and nitrogen fixation (topic 45). University of Sheffield has a long history and disproportionate contribution to the study of neurodegenerative diseases affecting motor neurons (topic 25), dating back to 2007.

Association of a topic with an institution that used to greatly contribute to the topic but then stopped might also explain why some topics declined in prevalence over time although drastic growth in other topics might be a better explanation (Figure 6.8, 6.9). Topic 29 prominently features the bone growth plate though this `stm` topic has entries for other biological systems as well. These bone growth plate papers come from Upstate Medical University in Syracuse, New York, from 2005 to 2010. Decline in topic 29 might be related to cessation of study of the growth plate at this institution after 2010, though other institutions have not picked up this topic afterwards. Institute of Human Genetics and Anthropology, Friedrich-Schiller-University in Jena, Germany greatly contributed to cancer proteomics (topic 10) between 2004 and 2011 but then stopped, though other institutions carried on studying this topic. Kyushu University Beppu Hospital in Japan greatly contributed to quantitative analyses in cancer (topic 6) from 2005 to 2014, although other institutions continue contributing to this topic, whose paper count actually increased over time although the topic's proportion decreased due to drastic growth in other topics (6.9). The vast majority of LCM related publications from Sendai, Japan are about breast cancer (topic 13), from between 2007 to 2017, which is why Sendai is associated with this city although this topic is widespread.

Association of a city with a topic can also be visualized with topic proportion in each city from `estimateEffect` (Figure 6.12). Here topic 45 (soil microbiome and nitrogen fixation) is plotted, but readers on RStudio Cloud can try other topics.

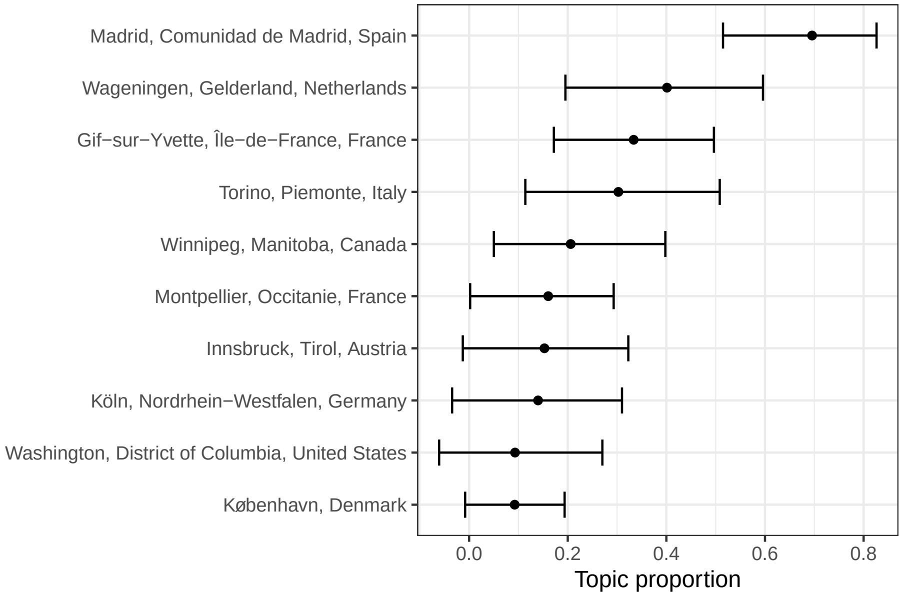

**Figure 6.12:** Proportion of topic 45 in each city. Error bars are 95% CI of the point estimate.

Here "disproportionate" means disproportionate within this corpus of LCM related search results. Institutions with "disproportionate" contribution to a topic do not necessarily dominate such topic although the topic may dominate the institution, i.e. the topic takes up a very large proportion of abstracts from this institution within this corpus. Nor are these institutions necessarily elite; this analysis might be an interesting way to discover labs from not so well-known institutions that may be outstanding in some topics. The institutions are often not elite because elite institutions often greatly contribute to many topics, weakening the association of the institution to the topic. Except for growth plate in Syracuse, we have not identified topics largely confined to an institution.

#### 6.5 GloVe word embedding

We used global vector (GloVe) embedding to identify linear substructures in the word vector space of the LCM transcriptomics abstract corpus and to identify contexts (Pennington, Socher, and Manning, n.d.). In GloVe embedding, words are represented by vectors. Words with similar meanings tend to be closer together in this vector space, and differences between word vectors can encode meaning as well. The "meanings" come from the context, or word co-occurrence. GloVe was devised to find a word embedding with properties like "king" - "man" + "woman" = "queen" or "ice" - "solid" + "gas" = "steam", and related words like "cancer" and "tumor" are close together but both are far from unrelated words like "flower".

This corpus was used to train a 125 dimensional embedding, and the embeddings of words occurring more than 30 times in the corpus were projected to lower dimensions with principal component analysis (PCA) to find axes explaining the most variance in the embedding, hopefully identifying dominant axes of meaning within this corpus. The words are also Louvain clustered in the embedding space to find clusters of words related in meaning.

The first principal component (PC) explains over 5% of the variance, and then the "elbow" is at PC5.

Words more positive in PC1 are often gene names, parts of gene names, or acronyms, and names of specific biological entities or processes. In contrast, words more negative in PC1 tend to be more general and more widely used. PC2 separates the technical (clusters 2 and 7, top) from the biological (clusters 1, 4 to 6) (Figure 6.14). As expected, "cancer", "tumor", and "disease" are not far from each other (bottom left), and "malignant" and "invasive" are close (bottom center). PC1 explains more variance than all other PCs; though it's only 5.5%, it picked up a very important dimension in word meanings in this corpus. PCs are arranged in decreasing order of variance explained.

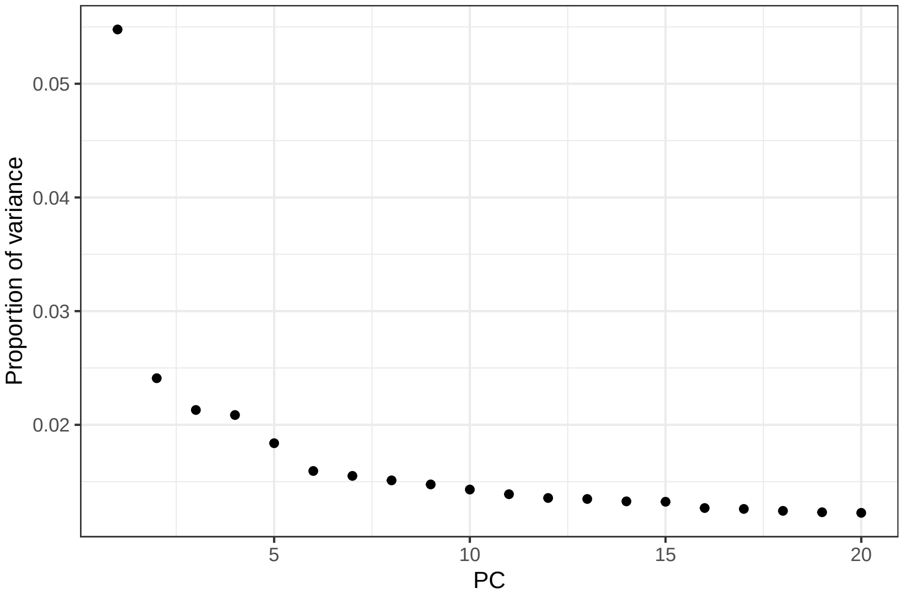

**Figure 6.13:** Proportion of variance explained by each of the first 20 principal components (PC).

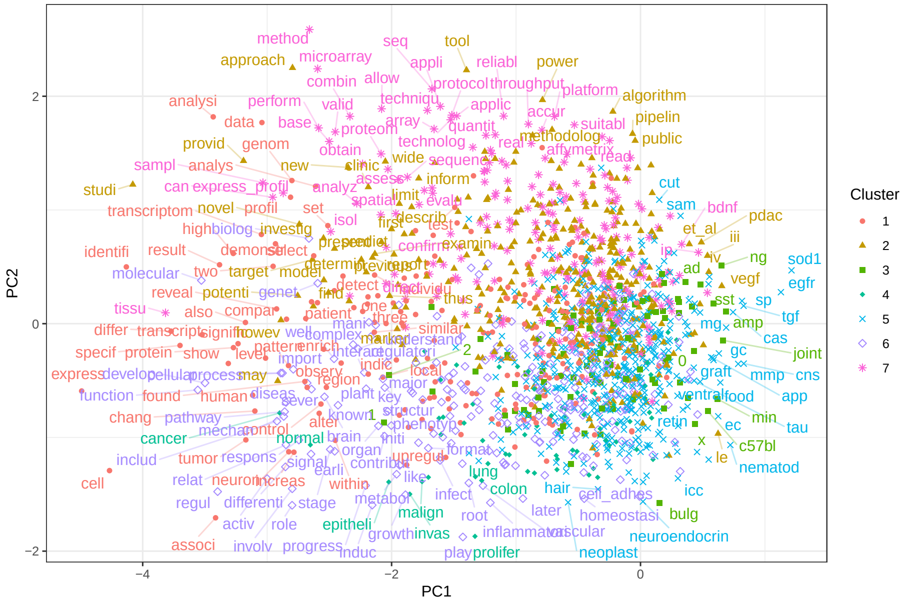

**Figure 6.14:** Projection of word embeddings into the first 2 PCs. Each point is a word occuring over 30 times in the corpus. Not all words are labeled to avoid overlaps in the labels. Words and points are colored by Louvain clusters.

Note that when the PCA plot is made on different computers, the signs of PCs might flip, because the sign does not affect the magnitude of the eigenvalue (i.e. variance explained). PCs are eigenvectors of the covariance matrix of the GloVe dimensions; an eigenvector multiplied by a scalar is still an eigenvector with the same eigenvalue.

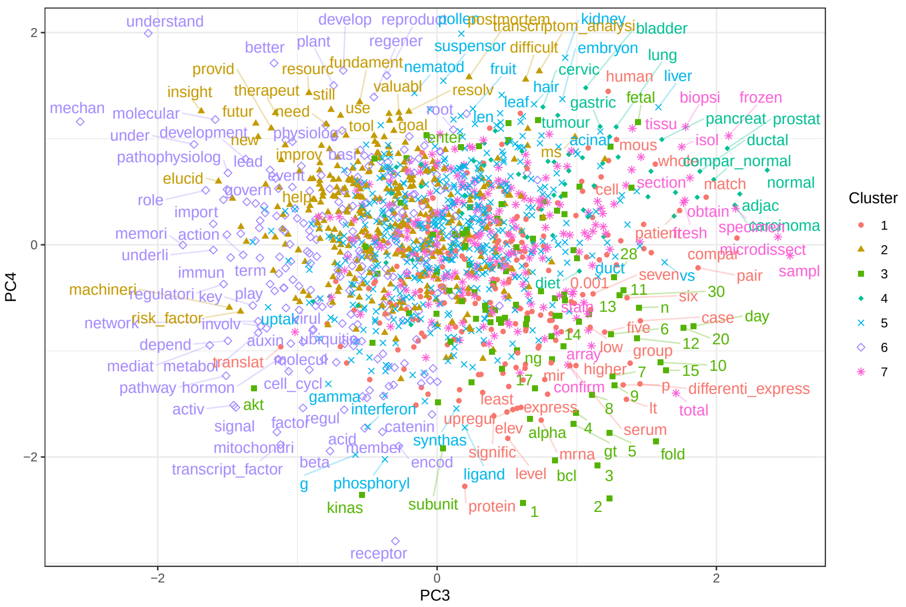

**Figure 6.15:** Projection of word embeddings into the 3rd and 4th PCs.

PC3 separates processes and interactions (cluster 6, left) from entities of samples, tissues, organs, and diseases (clusters 3, 4, 7, right). PC4 separates the molecular and cellular (bottom left) and the quantitative (clusters 1 and 3, bottom right) from the qualitative (top). Some of the qualitative terms are used to discuss implications of results of the papers (clusters 2 and 6, top left) (Figure 6.15).

Now we have seen some important axes of meanings and types of words, which are not surprising given familiarity with the general structure of abstracts and applications of LCM. There must be more axes of meaning, as the first 4 PCs only explain about 12% of the total variance of word embeddings (Figure 6.13). The clusters of words can be better visualized with UMAP, which is a non-linear dimension reduction method that tries to preserve distances between points but is most commonly used to project into 2 dimensions.

The clusters of words are easier to discern with Uniform Manifold Approximation and Projection (UMAP) (Figure 6.16). Cluster 1 is words used to describe

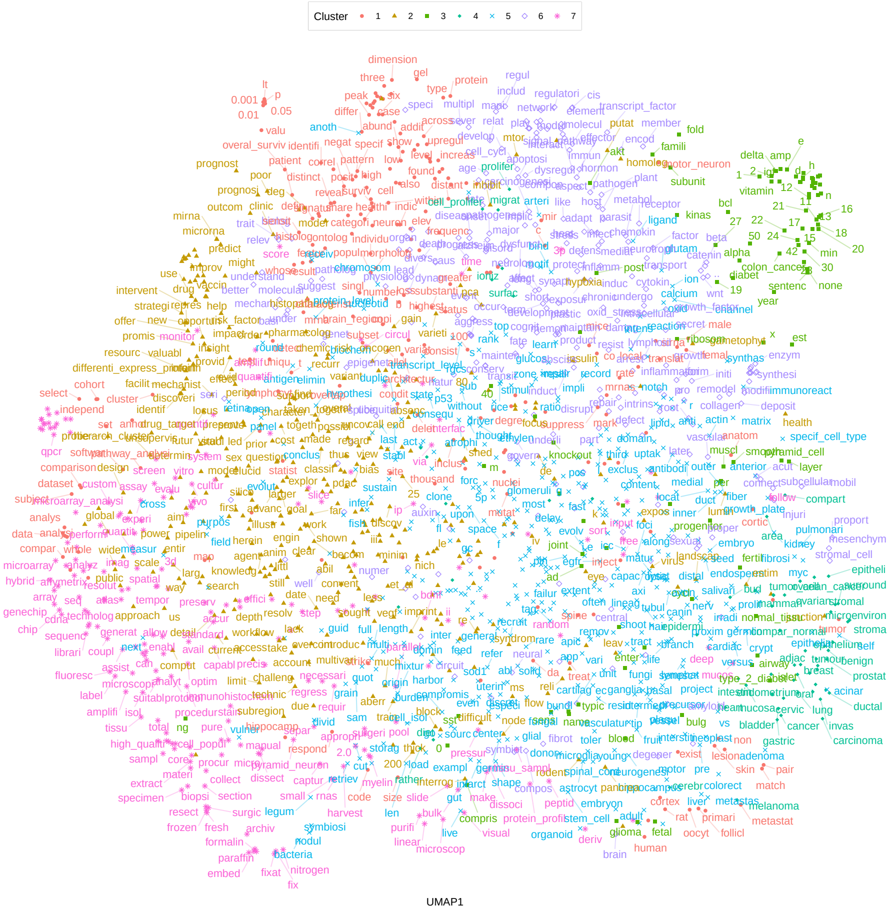

**Figure 6.16:** UMAP projection of word embeddings. Zoom in if reading the PDF version of this book.

results of studies, with many quantitative words; “p”, “0.05”, and “0.01” are found in this cluster rather than cluster 3 because p-values are results of data analyses and 0.05 and 0.01 are common thresholds of significance. Cluster 2 has many words discussing results, about data analysis and implications of results, with many clinical terms. Cluster 1 seems to be molecular and cellular processes and entities. Cluster 3 has many numbers, units, and some biological words. Cluster 4 has many words specifically about cancer. Cluster 5 is words in experimental techniques. Cluster 5 is biological terms. Cluster 6 is also biological, but with more emphasis on processes and interactions. Cluster 7 is technical. These clusters of words give some idea about topics of the studies, but unlike `stm`, these clusters also give a glimpse into different parts of the abstract, such as summary of the results and implications of the results.

### Chapter 7

### Data analysis in the current era

> Many machine learning and statistics methods are mentioned in this chapter. The names of these methods are linked to articles explaining them for those who are unfamiliar. Some of them are math heavy.

So far we have reviewed numerous techniques to collect spatial transcriptomics data. In this chapter, we review computational methods to analyze data generated by current era techniques and methods that, while only having WMISH, FISH, or ISH as spatial data, involve scRNA-seq data as well. For a publication to be included in the "Analysis" sheet of this database, it must either focus on a data analysis method, or present alongside new data, sophisticated data analysis going beyond using existing packages. While some data analysis methods originally not designed for spatial data can be used for spatial data, this chapter is about methods designed specifically with spatial data in mind. This means that the methods should be demonstrated on a spatial transcriptomic dataset in the publication, even if not explicitly using spatial coordinates.

Since 2019, there has been a sharp increase in interest in current era data analysis (Figure 7.2). If our collection of prequel data analysis literature is somewhat representative and complete, then interest in current era data analysis dwarfs the golden age of prequel data analysis from 2008 to 2014 (Figure 7.1). As already shown, interests in current era data collection increased sharply since 2018 (Figure 4.2, Figure 7.2); interest in data analysis lagged behind interest in data collection, until around 2020 (Figure 7.2).

In contrast, in the prequel era, interest in data analysis peaked after the peak for data collection, and eventually interest both eventually diminished but continues

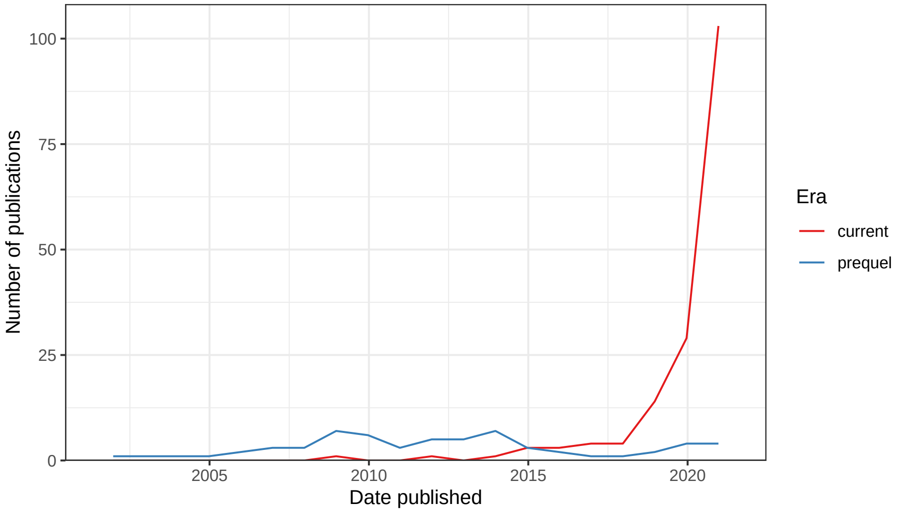

**Figure 7.1:** Number of publications over time for current era and prequel data analysis. Bin width is 365 days. Preprints are included for this figure.

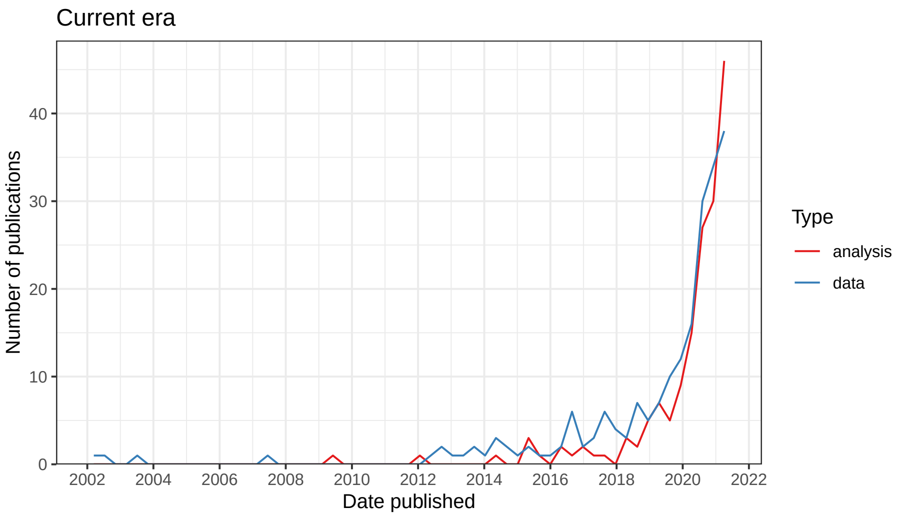

**Figure 7.2:** Number of publication over time for current era data collection and data analysis. Bin width is 120 days. Note that the count drops in 2021 because this plot was made in April 2021.

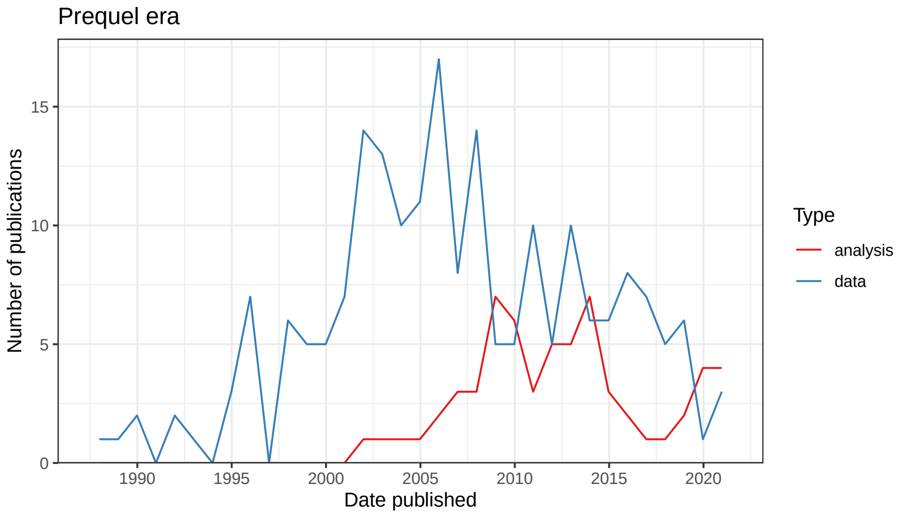

**Figure 7.3:** Number of publications over time for prequel data collection and data analysis. Bin width is 365 days.

(Figure 7.3). There are many different types of data analysis, the ones with the most interest are mapping dissociated cells in scRNA-seq to location in a spatial reference (cells to locations) and imputing expression of genes not profiled in the spatial reference according to transcriptome wide scRNA-seq data (gene expression imputation at locations) (Figure 7.4).

In 2020, several methods for cell type deconvolution in array based techniques that don't have single cell resolution were developed (cell type inference), but the drastic growth in data analysis seems to be driven by multiple categories of analyses (Figure 7.5). Top contributors to data analysis methods in the current and prequel eras are different as well. Again, the current era seems to be more of an elite club than the prequel era although some not as elite institutions have contributed as well but some less well-known institutions have joined more recently; among the top contributors in the prequel era are less famous institutions such as Arizona State University (ASU), Old Dominion University (ODU), and Lawrence Berkeley National Laboratory (LBL), which developed the BDTNP and the Fly Enhancer atlases (Figure 7.6).

In our database, we have recorded programming languages used in data analysis or package development. All programming languages that played a major role in the project were recorded. For downstream analysis, this includes languages of the user interface of existing packages used and languages of new functions written for the project. For package development, this includes any language used to write the package essential to the functioning of the package. In publications that focus on data collection, R is by far the most popular programming language used in downstream data analysis (Figure 7.7). The second most pop-

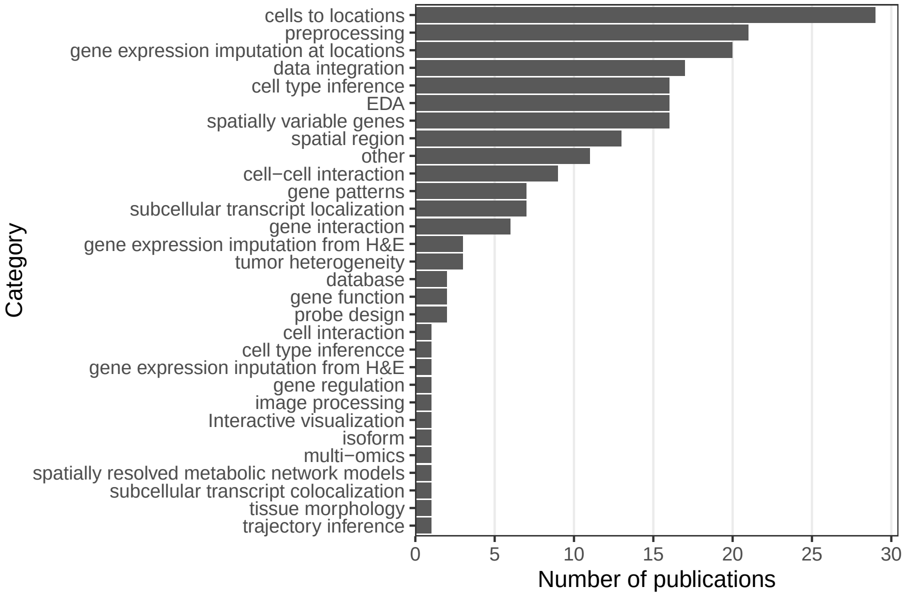

**Figure 7.4:** Number of publications for each category of data analysis; note that the same publication can fall into multiple categories.

ular is MATLAB, which is more common in smFISH (Figure 5.25) and ISS for its image processing functionality. Python follows closely, and is used for both image processing and other types of analyses. C and C++ are not as common in downstream analysis.

The same top 5 programming languages are the most common for developing data analysis packages (Figure 7.8). Python is the most popular, especially for packages involving deep learning, image processing, using Torch for optimization, or are command line tools. R follows, and is more popular for exploratory data analysis (EDA) and data visualization, but often both R and Python are used in the same package. Other languages aren't nearly as commonly used for packages reported on in our database. The above observations about usage of R and Python seem to reflect the broader cultural differences between the R and Python communities; the former caters more to the users and statisticians who do not specialize in computer science, while the latter caters more to developers and computer science specialists. MATLAB is not as commonly used for package development. While popularity of Python and R have grown (and some others such as Julia), the popularity of MATLAB seems more level (Figure 7.9). C and C++ are more common in package development than in downstream analysis, but are often used in conjunction with either R or Python or both as C and C++ are used for performance while R and Python are for user

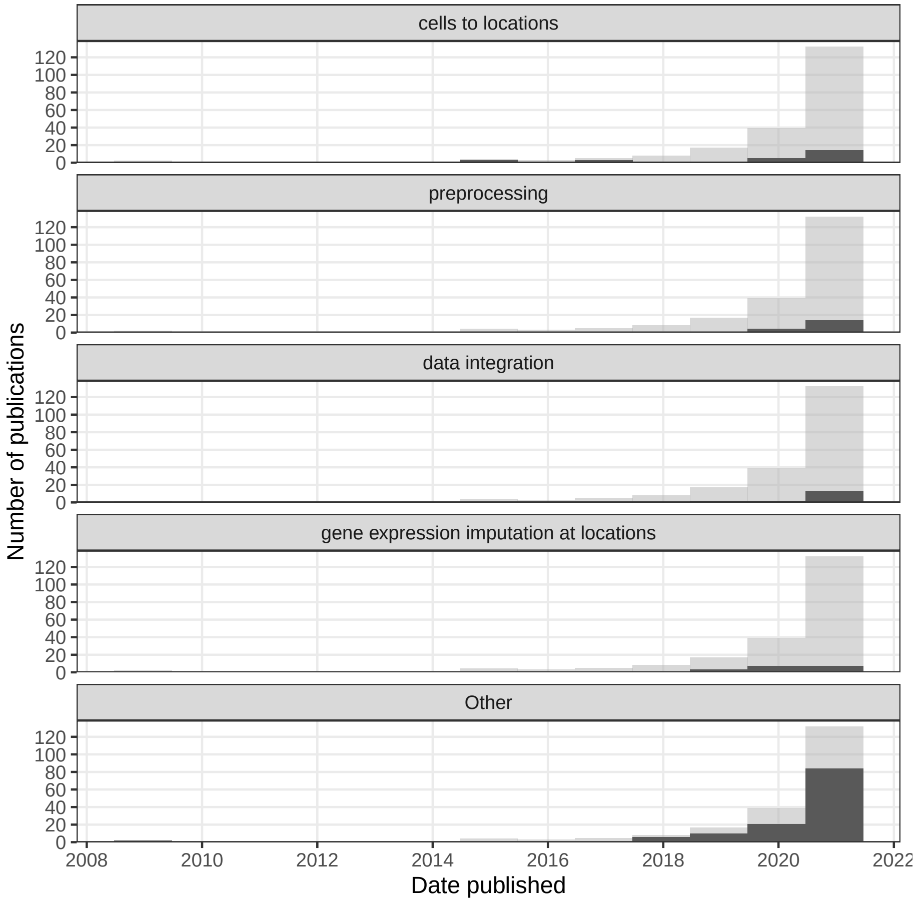

**Figure 7.5:** Number of publications over time broken down by type of data analysis. The 3 categories most popular in the past year are shown, and the others are lumped into 'Other'. Bin width is 365 days.

interface. With packages such as `reticulate`[1], `rpy2`[2], `basilisk`[3], `Rcpp`[4], and `Cython`[5], the most popular open source languages can be made interoperable to each other to some extent, making use of the best resources from each language.

We have also recorded whether the package is well documented and whether it's hosted on a public repository as a loose proxy of user friendliness and quality. Here "well documented" means at least all arguments of all functions exposed

[1] `https://rstudio.github.io/reticulate/`
[2] `https://rpy2.github.io/doc/latest/html/introduction.html`
[3] `https://bioconductor.org/packages/release/bioc/html/basilisk.html`
[4] `http://rcpp.org`
[5] `https://cython.org`

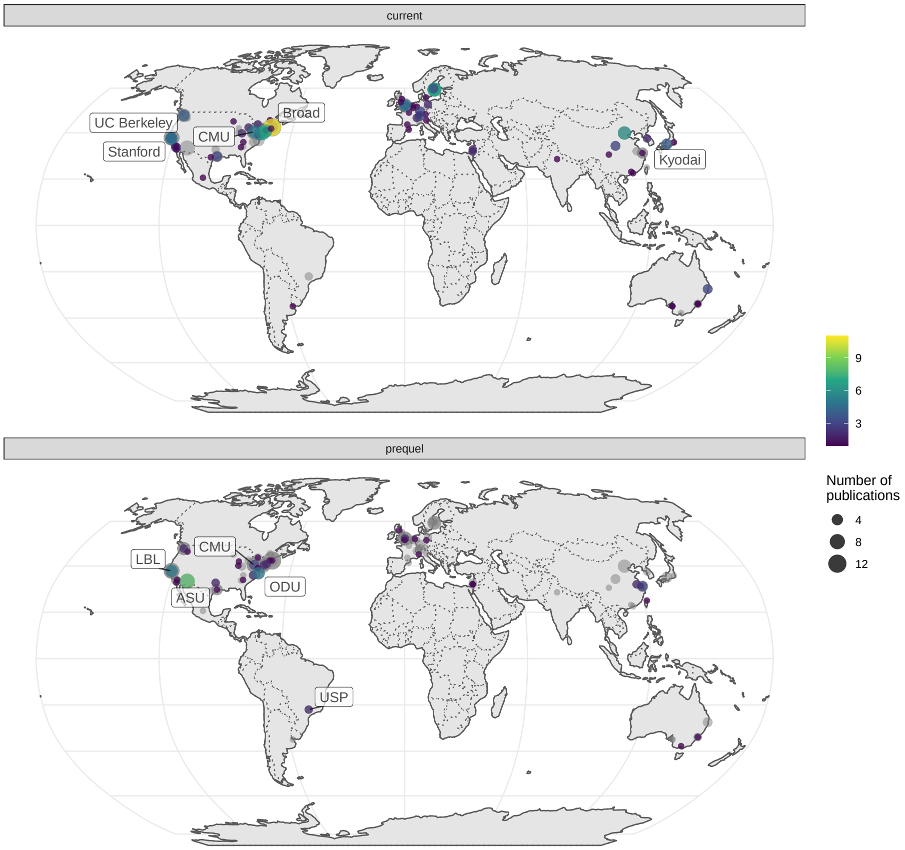

**Figure 7.6:** Map of where first authors of current era and prequel data analysis papers were located as of publication. Top 5 institutions in each era are labeled.

to the user are documented, though we consider it better when examples are included. Public repositories can to some extent indicate user friendliness and quality because the packages need to pass some sort of checking in order to be hosted on the repositories, though some repositories, such as Bioconductor, have stricter standards than others. Moreover, installation of the package is easier when the package is on a public repository. A modest majority of Python packages and the vast majority of R and C++ packages are well documented, while many MATLAB packages are not (Figure 7.9). Most packages are not on a public repository such as CRAN, Bioconductor, pip, and conda (Figure 7.10).

Some of the most popular categories of analyses (Figure 7.4) are reviewed in the rest of this section, arranged roughly in the order each task is performed in a data analysis workflow. Each category will first be defined, and the common

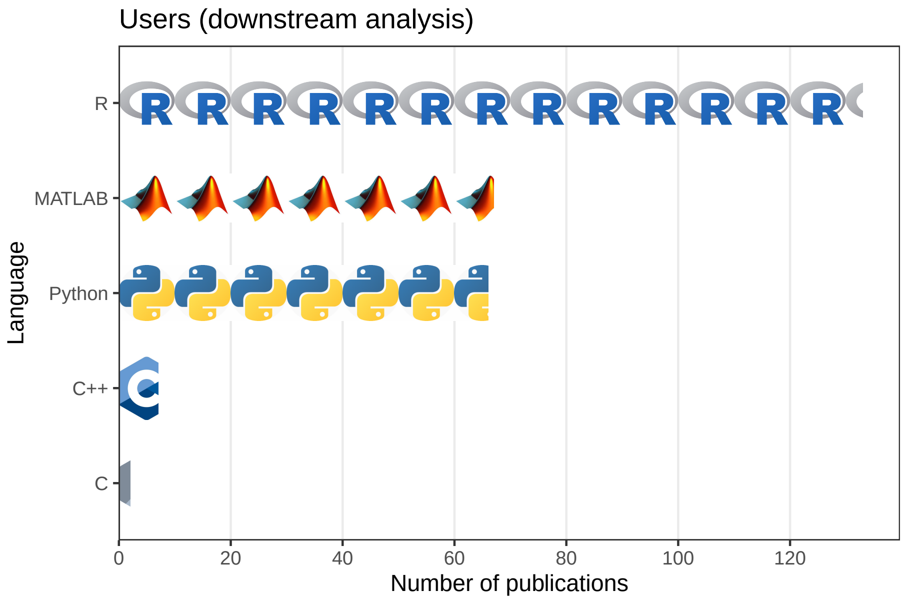

**Figure 7.7:** Number of publications for data collection using each of the 5 most popular programming languages for downstream data analysis.

core principles will be summarized.

#### 7.1 Preprocessing

By "preprocessing" we mean extracting information from raw data so common analysis methods can be applied. "Raw data" can mean any form of data, even if processed in some ways, that still needs to have information extracted for common analysis tasks to apply, such as PCA, clustering, and DE. Preprocessing for array based techniques that use NGS is similar to preprocessing for scRNA-seq. The same aligners can be used to align reads to the genome or pseudoalign to the transcriptome, and the spot barcodes can be demultiplexed just like in scRNA-seq; indeed, ST and Visium, the preprocessing pipelines ST Pipeline and Space Ranger wrap the STAR aligner. As microdissection based techniques also use NGS, preprocessing would not be very different from that of scRNA-seq or bulk RNA-seq data. However preprocessing of smFISH and ISS data is very different from that of NGS based data, and this would be the focus of this section.

The raw data is images. As mentioned earlier, preprocessing of images was typically performed with poorly documented MATLAB code difficult to decipher by users. While some switched to Python recently, such as in MERlin for

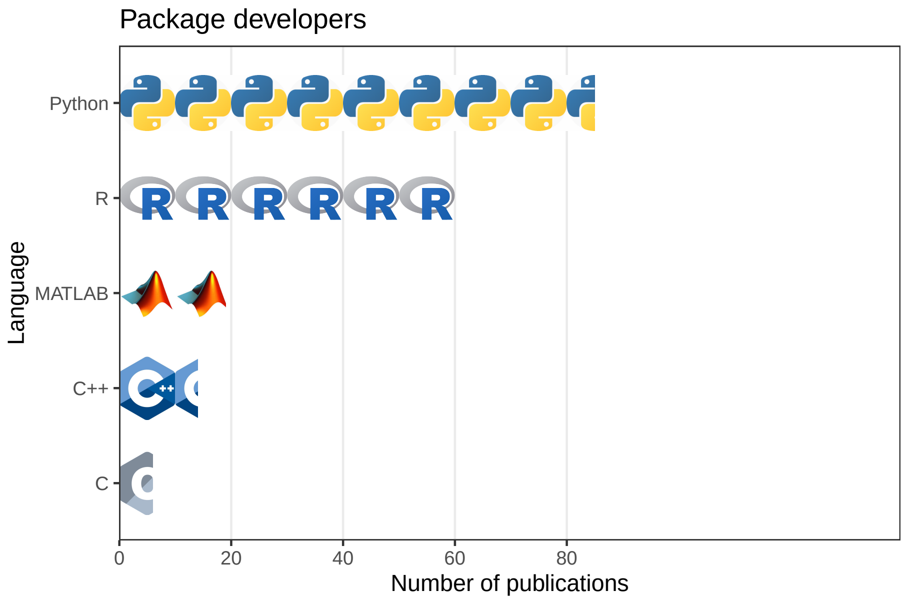

**Figure 7.8:** Number of publication for data analysis using each of the 5 most popular programming languages for package development. In this and the previous figure, each icon stands for 10 publications, and the x axes of both figures are aligned. Note that multiple programming languages can be used in one publication.

MERFISH, the preprocessing tool is still specific to the technique of interest. Also, the proprietary language MATLAB is still quite commonly used, such as for preprocessing HybISS and ISS data (Gyllborg et al. 2020; Qian et al. 2020). Some groups used GUI based tools such as Fiji, ImageJ, and CellProfiler (Shah et al. 2018; Chen et al. 2020; Sountoulidis et al. 2020). However, as the GUI based analyses are not recorded and shared or are manual, it is difficult to reproduce such analyses.

To provide a free, open source, and well-documented preprocessing tool applicable to data from multiple techniques, the Chan Zuckerberg Initiative developed the Python package `starfish` implementing image registration, spot calling, barcode calling, cell segmentation, and etc. with classical image processing methods such as thresholding, image registration by translation, top hat filtering, Laplacian of Gaussian, watershed segmentation, and etc. While a good start, it's not clear how to apply `starfish` to multiple FOVs based on its tutorials. To improve `starfish`, another Python pipeline, SMART-Q was developed, with more modularity and improvements upon `starfish` such as additional parameter to mitigate over-segmentation (individual cell or nucleus broken into too many pieces) by watershed and integration with immunofluorescence images of

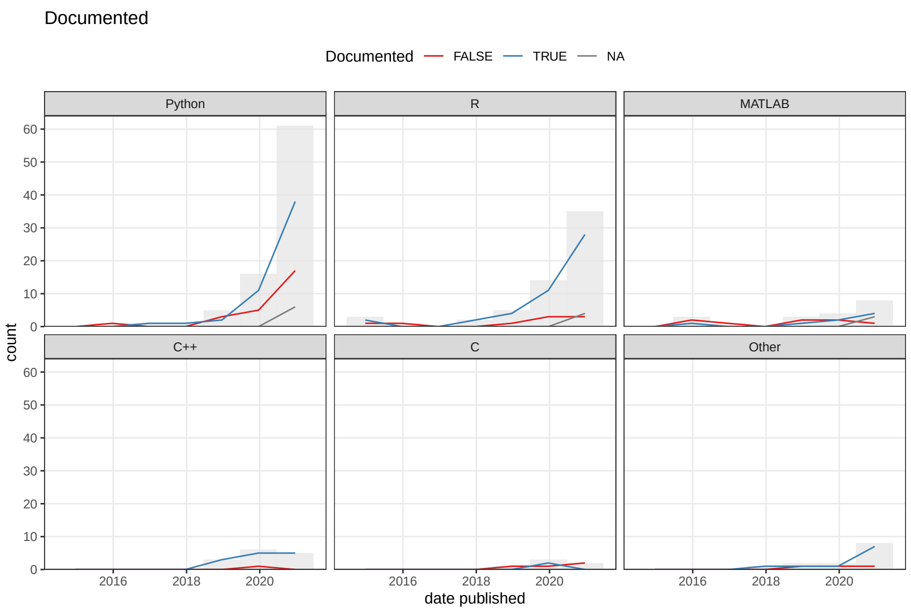

**Figure 7.9:** Among data analysis publications, the number of packages that are or are not well documented over time.

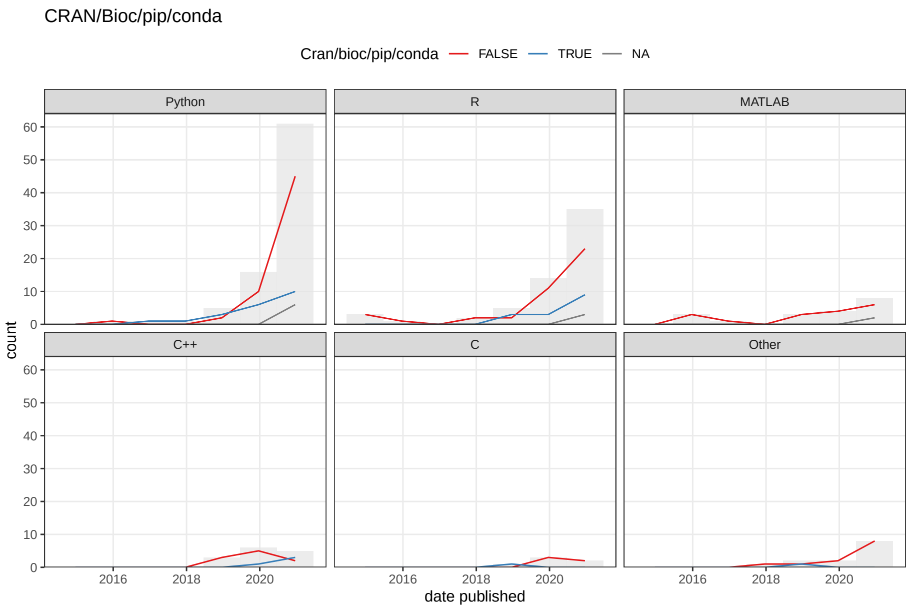

**Figure 7.10:** The number of packages that are or are not on a public repository such as CRAN, Bioconductor, pip, or conda over time. In both C and D, the bin width is 365 days. NA means the source code repository is not available.

marker genes (Yang et al. 2020). However, SMART-Q was only demonstrated in RNAscope data without combinatorial barcoding, with one FOV at a time. Another such smFISH pipeline based on classical image processing is `dotdotdot`, which is written in MATLAB but the functions are well documented (Maynard et al. 2020). Again, `dotdotdot` was only demonstrated on RNAscope without combinatorial barcoding. There are other open source tools for one or more of the preprocessing steps, but are not meant to be a comprehensive pipeline. Below we review each step in preprocessing of smFISH and ISS raw data, how this was done in the original papers of datasets with classical image processing, and alternative and improved approaches such as ones based on deep learning or Bayesian statistics.

The packages mentioned in this section are summarized in the Table 7.1. The package names link to the code repo if available, and the titles link to the paper associated with the package. Each section in this chapter has a table like this. There are relevant packages not mentioned in this book; they can be found in the database[6].

**Table 7.1:** Packages mentioned for smFISH and ISS image processing

| Name | Language | Title | Date published |
|---|---|---|---|
| corrFISH[7] | MATLAB | Dense transcript profiling in single cells by image correlation decoding[8] | 2016-06-06 |
| graph-ISS[9] | Python | Identification of spatial compartments in tissue from in situ sequencing data[10] | 2019-09-18 |
| SSAM[11] | Python; C++ | Segmentation-free inference of cell types from in situ transcriptomics data[12] | 2019-10-13 |
| pciSeq[13] | MATLAB | Probabilistic cell typing enables fine mapping of closely related cell types in situ[14] | 2019-11-18 |
| SMART-Q[15] | Python | SMART-Q: An Integrative Pipeline Quantifying Cell Type-Specific RNA Transcription[16] | 2020-04-29 |
| JSTA[17] | Python; C | JSTA: joint cell segmentation and cell type annotation for spatial transcriptomics[18] | 2020-09-20 |
| spage2vec[19] | Python | Spage2vec: Unsupervised detection of spatial gene expression constellations[20] | 2020-09-25 |

[6]`https://docs.google.com/spreadsheets/d/1sJDb9B7AtYmfKv4-m8XR7uc3XXw_k4kGSout8cqZ8bY/edit#gid=1424019374`

| | | | |
|---|---|---|---|
| Baysor[21] | Julia | Bayesian segmentation of spatially resolved transcriptomics data[22] | 2020-10-06 |
| deepBlink[23] | Python | deepBlink: Threshold-independent detection and localization of diffraction-limited spots[24] | 2020-12-15 |
| ISTDECO[25] | Python | ISTDECO: In Situ Transcriptomics Decoding by Deconvolution[26] | 2021-03-02 |
| BarDensr[27] | Python | BARcode DEmixing through Non-negative Spatial Regression (BarDensr)[28] | 2021-03-08 |

##### 7.1.1 Image registration

First, images of each FOV from different rounds of hybridization must be aligned; this is image registration. The images can be aligned to a reference of fiducial beads or DAPI staining, which is especially useful when "no fluorescence" is part of the barcode (K. H. Chen et al. 2015; Eng et al. 2019). If "no fluorescence" is not involved, then the reference can be a particular round of hybridization (Shah et al. 2016; Wang, Moffitt, and Zhuang 2018). Image registration is usually affine, i.e. images are translated, scaled, or rotated to match the reference, and often only translation is used. However, non-linear registration has been used in case the sample does not lie flat and chromatic aberration shifts spots in different channels (Qian et al. 2020).

##### 7.1.2 Spot and barcode calling

Then the spots representing individual transcripts are identified (spot calling). The background of autofluorescence and non-specific hybridization is often removed by thresholding or top hat filtering, only preserving brighter pixels. Spots can be identified with multi-Gaussian fitting[29] with fixed width, which can distinguish between partially overlapping spots (K. H. Chen et al. 2015), or tightened by Lucy-Richardson deconvolution[30] (Moffitt et al. 2018), or by identifying local maxima in intensity after identifying potential spots with Laplacian of Gaussian[31] (Shah et al. 2016; Wang, Moffitt, and Zhuang 2018). The spots can also be identified with deep learning. In Python package graph-ISS (Partel et al. 2019), a convolutional neural network (CNN)[32] is pretrained on manually

[29]http://www.inf.ed.ac.uk/teaching/courses/inf2b/learnSlides/inf2b12-learnlec09.pdf
[30]https://en.wikipedia.org/wiki/Richardson-Lucy_deconvolution
[31]http://fourier.eng.hmc.edu/e161/lectures/gradient/node8.html
[32]https://towardsdatascience.com/a-comprehensive-guide-to-convolutional-neural-networks-the-eli5-way-3bd2b1164a53

annotated candidate signal spots from another dataset, and probability that a new candidate obtained after top hat filtering[33] and h-maxima transform[34] is a signal is returned by the last softmax layer of the CNN. Another CNN based spot calling tool is deepBlink (Eichenberger et al. 2020), which builds on the popular U-net architecture.

Once spots are called in each round of hybridization, spots that most likely to correspond to the same transcript are read as barcode and decoded to identify the gene encoded by the barcode (barcode calling). As image registration is imperfect, the spot coming from the same transcript may still be slightly shifted between rounds of hybridization. To identify the barcode from the rounds of hybridization, the spot in one round of hybridization is typically identified with a spot in another round if the spatial distance between the two is sufficiently small, such as less than between 1 and 3 pixels, or smaller than the distance to a barcode that contains error (Shah et al. 2016; Wang, Moffitt, and Zhuang 2018; Moffitt et al. 2016; Eng et al. 2019).

In graph-ISS (Partel et al. 2019), spots identified from CNN from different rounds of hybridization are connected in a graph, with each spot in each round of hybridization a node and the edge weight decreases with increasing distance between spots across rounds up to a maximum distance. Edges connecting spots not from consecutive rounds are removed. The barcode is called by maximum flow of minimum costs between the sink and the source of the graph. Then a quality score is calculated for the barcode according to the CNN probability of spots and distance between spots from different rounds. Although graph-ISS was originally designed for ISS data, it might be adapted to seqFISH, HybISS, STARmap, and SCRINSHOT as well. However, for MERFISH and seqFISH+, in which a transcript may not have signal in some rounds of hybridization, graph-ISS would need to be altered. Alteration would also be required to decode STARmap's 2 base encoding.

For MERFISH specifically, transcript counts have been statistically modeled in the Rust package MERFISHtools, which takes errors in barcode calling into account (Köster, Brown, and Liu 2019). While MERFISH's inbuilt error correction (HD4) accounts for 1 to 0 error, which is more common, 0 to 1 errors can still occur, and there are still barcodes with so many errors that they can't be matched to genes (dropout). The errors are modeled as a multinomial distribution[35] with event probabilities as probabilities of identifying transcripts of a gene correctly with and without the inbuilt correction, misidentifying transcripts of a gene as those from each other gene with and without the inbuilt correction, and dropouts, with actual transcript counts, number of correct and incorrect identifications, and dropouts as latent variables to be estimated by Bayesian inference. The flat prior is used for now.

[33] `https://micro.magnet.fsu.edu/primer/java/digitalimaging/russ/tophatfilter/index.html`

[34] `https://en.wikipedia.org/wiki/H-maxima_transform`

[35] `https://online.stat.psu.edu/stat504/node/40/`

Computational methods to overcome optical crowding and to deconvolute spots were summarized in Section 5.2.3: corrFISH, BarDensr, and ISTDECO. The above mentioned spot calling methods all treat spot detection and decoding as separate tasks. In contrast, in both BarDensr and ISTDECO, the two related tasks are performed jointly.

##### 7.1.3 Cell segmentation

To assign transcript spots to cells, the cells need to be segmented and spots within the segmented boundary of a cell must be assigned to that cell. For neurons, Nissl staining, which stains the cell body and dendrites but not axons, has been used for cell segmentation (Shah et al. 2016; Wang, Moffitt, and Zhuang 2018). Without Nissl staining, total poly-A staining can be used instead, and segmented with watershed transform, although poly-a staining concentrates in the cell body and misses cellular processes such as dendrites (Moffitt et al. 2018). This misses some interesting biological information; dendrites can have different transcriptomes from the cell body of the same neuron, both in vitro and in vivo (Middleton, Eberwine, and Kim 2019; Ciolli Mattioli et al. 2019; Farris et al. 2019). Cell segmentation can be done manually as automated methods may not be sufficiently reliable and would still require manual inspection and correction, or automated with machine learning models trained by manual segmentation of smaller number of cells such as the random forest model in Ilastik (Wang, Moffitt, and Zhuang 2018; Lohoff et al. 2020) and CNN models such as DeepCell (Van Valen et al. 2016) and U-net (Ronneberger, Fischer, and Brox 2015). Watershed segmentation[36] is more commonly used.

Without seeing the actual extent of the cell, the quality of manual segmentation is questionable, especially in regions with high cell density, thus limiting the performance of machine learning models. Sometimes problematic methods were used to segment cells, such as 3D Voroni tessellation[37] (Shah et al. 2016) and convex hull[38] of Nissl staining based segmentation (Wang, Moffitt, and Zhuang 2018); these are problematic because cells need not to take a convex shape so such segmentation may mis-assign transcripts from other cells, or to be conservative about mis-assigning transcripts from other cells, miss transcripts that in fact belong to the cell of interest. However, one study did specifically stain for membrane bound proteins for the actual extent of the plasma membrane and accurate cell segmentation (Lohoff et al. 2020).

To address the challenges of cell segmentation, segmentation methods utilizing scRNA-seq data with annotated cell types have been developed recently. One such method is Python package JSTA (Littman et al. 2020), in which a deep neural network (DNN) learns a segmentation and cell type annotation using

[36] https://www.mathworks.com/company/newsletters/articles/the-watershed-transform-strategies-for-image-segmentation.html
[37] https://en.wikipedia.org/wiki/Voronoi_diagram
[38] https://en.wikipedia.org/wiki/Convex_hull

the information from a scRNA-seq reference with cell type annotations. First, watershed is used for an initial cell segmentation, both MERFISH and scRNA-seq data are scaled and centered. Then a DNN is trained on the scRNA-seq data to predict cell type from gene expression. Then a separate DNN is trained to refine the cell boundaries iteratively with expectation maximization (EM)[39]: The cell type classifier is applied on the watershed segmented MERFISH data to classify putative cells (E). Then a random subset of the pixels are used to train the pixel classifier, maximizing a loss function comparing the new pixel cell type probabilities to the initial/previous assignment (M). The new cell type probabilities are then scaled per pixel according to distance to nuclei. Only probabilities of cell types of neighboring cells are kept and the other cell types are assigned probability 0. The new cell type probabilities of each pixel is then used as event probabilities of a multinomial distribution and randomly assign a new cell type label to the pixel. Then the new cell type assignment to pixels is used to train the pixel classifier again, until the cell type assignments converge. This may refine boundaries between neighboring cells of different types, and the initial watershed boundaries are kept for neighboring cells of the same type. A problem with this package is that inhomogeneous transcript localization is not taken into account.

##### 7.1.4 Alternatives to cell segmentation

Due to the challenges in accurate cell segmentation, some analysis methods did away with cell segmentation altogether, directly using the transcript locations. In the Julia package Baysor (Petukhov et al. 2020), based on Markov random field (MRF)[40], which encourages nearby transcripts to take the same label. A spatial neighborhood graph is constructed with Delaunay triangulation[41] with each transcript as a node. The probability of each transcript taking each label is modeled with a MRF and initial edge weights decrease with distance. This package first distinguishes between intracellular transcripts and extracellular background. Then it can also assign transcripts to cell types without cell segmentation, with a scRNA-seq reference with cell type annotations; as locations of the transcripts are known, this amounts to annotating tissue regions with cell types. It can also segment cells, with existing segmentation and staining (e.g. Nissl, DAPI, and poly-A) as priors. Cell segmentation can also be informed by cell type labels, so transcripts from different cell types are not assigned to the same cells. Each of the three functionalities, identifying intracellular transcripts, cell type annotation of transcripts, and cell segmentation, is based on a different MRF model. The parameters of the model, such as edge weights, labels of other transcripts, and etc. are estimated with EM. The drawbacks of

[39] `http://ai.stanford.edu/~chuongdo/papers/em_tutorial.pdf`

[40] `https://ocw.mit.edu/courses/electrical-engineering-and-computer-science/6-867-machine-learning-fall-2006/lecture-notes/lec23.pdf`

[41] `https://towardsdatascience.com/delaunay-triangulation-228a86d1ddad`

this package are that its current implementation is limited to 2D and it does not take inhomogeneous subcellular transcript localization into account.

Besides cell type annotation of transcripts based on MRF, another segmentation-free method is also described in the Baysor paper (Petukhov et al. 2020), in which the $k$ nearest neighbors of each transcript are taken to be a pseudo-cell and analyzed by standard scRNA-seq data analysis methods such as clustering, PCA, and UMAP. For ISS, transcripts can be probabilistically assigned to cells and cells to cell types, with pciSeq (Qian et al. 2020). Briefly, spatial locations of transcripts are modeled by a Poisson point process[42] whose intensity is scaled by a term following Gamma distribution[43] to give the negative binomial distribution[44] of transcript counts in cells. The intensity for each gene and each cell is also informed by distance between transcripts and nucleus centroids (from DAPI), scRNA-seq data of the cell type this cell belongs to, and the detection efficiency of ISS. The data consists of locations of transcripts and the genes they come from. The unknown parameters, such as probability of each transcript to come from each cell and each cell from each cell type, are estimated by variational Bayesian inference[45]. Cell types and spatial domains can also be identified without scRNA-seq cell type annotations as well.

In the Python package SSAM (Park et al. 2019), transcript density is first estimated with Gaussian kernel[46] density, which is then projected into a square lattice. Local maxima of transcript density are taken as pseudo-cells and clustered to infer *de novo* cell types. Then tissue domains are identified by clustering sliding windows of spatial cell type maps. Tissue domains can also be identified without appealing to cell types.

In the Python package spage2vec (Partel and Wählby 2020), graphs are constructed by connecting each transcript spot to its neighbors within a certain distance such that at 97% of all transcript spot are connected to at least one neighbor. Then the transcript spots with these graphs are projected by a graph neural network (GNN)[47] into a 50 dimensional space which is informed by the graphs and thus local neighborhoods of transcripts. The transcript spots in the 50 dimensional space can then be clustered or projected to 2 or 3 dimensions with UMAP[48] to show tissue domains.

#### 7.2 Exploratory data analysis

[42] https://hpaulkeeler.com/poisson-point-process/
[43] https://en.wikipedia.org/wiki/Gamma_distribution
[44] https://en.wikipedia.org/wiki/Negative_binomial_distribution
[45] https://omarelb.github.io/variational-bayes/
[46] https://wiki.analytica.com/index.php/Kernel_Density_Smoothing
[47] https://blog.exxactcorp.com/a-friendly-introduction-to-graph-neural-networks/
[48] https://umap-learn.readthedocs.io/en/latest/how_umap_works.html

**Table 7.2:** Packages mentioned for EDA

| Name | Language | Title | Date published |
|---|---|---|---|
| Spaniel[49] | R | Spaniel: analysis and interactive sharing of Spatial Transcriptomics data[50] | 2019-05-05 |
| Seurat3[51] | R | Comprehensive Integration of Single-Cell Data[52] | 2019-06-13 |
| SpatialCPie[53] | R | SpatialCPie: an R/Bioconductor package for spatial transcriptomics cluster evaluation[54] | 2020-04-29 |
| STUtility[55] | R | Seamless integration of image and molecular analysis for spatial transcriptomics workflows[56] | 2020-07-16 |
| SPATA[57] | R | Inferring spatially transient gene expression pattern from spatial transcriptomic studies[58] | 2020-10-21 |
| SpatialExperiment[59] | R | SpatialExperiment: infrastructure for spatially resolved transcriptomics data in R using Bioconductor[60] | 2021-01-27 |
| Squidpy[61] | Python | Squidpy: a scalable framework for spatial single cell analysis[62] | 2021-02-20 |
| Giotto[63] | R | Giotto, a pipeline for integrative analysis and visualization of single-cell spatial transcriptomic data[64] | 2021-03-08 |

After data preprocessing, as described above, for array or microdissection based data, we get a gene count matrix with locations of voxels, and for smFISH and ISS based data, we get locations of transcripts, and if cell segmentation is performed, a gene count matrix and cell boundaries as well. For scRNA-seq, Seurat (Stuart et al. 2019), scanpy, and packages surrounding SingleCellExperiment on Bioconductor such as scran and scater implement further preprocessing of the gene count matrix, such as data normalization and scaling, as well as basic EDA methods to inspect and create an overview of the data, such as quality control (QC), data visualization, finding highly variable genes, dimension reduction, and clustering, and have user friendly tutorials, consistent user interface, and decent documentation. Such integrative EDA packages, as well as more specialized data visualization packages, have emerged for spatial transcriptomics as well, and are reviewed in this section.

In practice, spatial transcriptomics data is often analyzed with standard scRNA-seq analysis at the EDA stage, with one or more of PCA, tSNE[65], UMAP, clustering cells or spots, and finding marker genes for clusters, and differential expression (DE) between case and control (Shah et al. 2016; Moffitt et al. 2018; Zhang et al. 2020; Moncada et al. 2020; Berglund et al. 2018). For ST and Visium, the data is also often normalized like in scRNA-seq with CPM[66] or classical Seurat log normalization[67] and scaling[68] (Moncada et al. 2020; Ji et al. 2020; Berglund et al. 2018). Seurat also implements data integration, which has been used to transfer cell type labels from scRNA-seq to Visium for cell type deconvolution (Mantri et al. 2020), and can potentially be used to impute gene expression in non-transcriptome wide spatial data from scRNA-seq (discussed in Section 7.3). Then the clusters, marker genes, and genes of interest from scRNA-seq are often visualized within spatial context, and some studies proceed to other analyses that utilize the spatial information. Due to the relevance of scRNA-seq data normalization, EDA, and data integration to spatial data, the existing scRNA-seq ecosystems of Seurat, scanpy (spatial part in Squidpy (Palla et al. 2021)), and SingleCellExperiment (spatial part in SpatialExperiment (Righelli et al. 2021)) are adapting to the rise of spatial transcriptomics, with new data structures, visualization of gene expression and cell metadata (e.g. total UMI counts, cluster, and cell type) on the spatial coordinantes, with H&E as background for ST and Visium, and perhaps other spatial functionalities such as spatial neighborhood graphs and spatially variable genes.

There are other EDA packages not originating from an existing scRNA-seq EDA ecosystem as well. R packages Giotto (Dries et al. 2019), STUtility (Bergenstråhle et al. 2020), and SPATA (Kueckelhaus et al. 2020) not only support basic QC and EDA functionalities like those in Seurat, but also spatial analyses not supported by Seurat. These packages are well documented, but are not (yet?) on CRAN or Bioconductor.

Giotto has two main parts: Giotto Analyzer and Giotto Viewer. Besides basic Seurat functionalities and spatial data visualization, Giotto Analyzer implements several types of spatial analyses to be reviewed in more detail in the rest of this section: cell type enrichment in spatial data without single cell resolution, identifying spatially variable genes, gene co-expression patterns, cellular neighborhoods, interactions between cell types and ligand-receptor pairs in such interactions, and genes whose expression is associated with cell type interactions. However, the methods implemented in Giotto tend to have simpler principles than those of more specialized packages for each of the above tasks, such as hypergeometric test for cell type enrichment and spatially coherent genes, though Giotto wraps specialized packages such as SpatialDE (Svensson, Teichmann, and Stegle 2018), trendsceek (Edsgärd, Johnsson, and Sandberg 2018) for spatially variable genes, and smfishhmrf (Zhu et al. 2018) to identify spatial cellular neighborhoods. Giotto Viewer provides interactive visualization

[65] `https://en.wikipedia.org/wiki/T-distributed_stochastic_neighbor_embedding`
[66] `https://reneshbedre.github.io/blog/expression_units.html`
[67] `https://rdrr.io/cran/Seurat/man/NormalizeData.html`
[68] `https://rdrr.io/cran/Seurat/man/ScaleData.html`

of the data. As Giotto uses its own object class to store data, interoperability with other single cell and spatial software becomes more challenging given the popularity of Seurat and SingleCellExperiment.

In contrast, STUtility develops upon the Seurat class, so is interoperable with other Seurat functionalities. STUtility is specific to ST and Visium, while Giotto applies to all spatial technologies with cell or spot level data. Beyond Seurat, STUtility enables masking the array to remove spots outside the tissue, alignment of multiple sections, manual annotation and alignment with `shiny`[69], visualization of the aligned sections in 3D, finding neighbors of spots of a given type, and using NMF to identify archetypal gene expression patterns. While Giotto and STUtility might not have the most sophisticated spatial analysis methods, their main advantage is akin to that of Seurat and SingleCellExperiment, namely that multiple analysis tasks, often with a variety of algorithms for each task, can be done with the same object class and user interface, saving the time and trouble on learning new syntax and converting objects to new classes.

SPAtial Transcriptomic Analysis (SPATA), while implementing its own class, uses Seurat for data normalization and dimension reduction. SPATA also implements functions to visualize spatial data and a `shiny` app for not only interactive data visualization but also manually setting spatial trajectories and annotation of spatial regions. It also wraps Monocle 3 (Cao et al. 2019) for pseudotime analysis and SPARK (S. Sun, Zhu, and Zhou 2020) for finding spatially variable genes. In addition, SPATA implements its own method of finding spatially variable genes, reviewed in Section 7.5.

Some R packages have also been written for specific visualization tasks, but not the entire EDA process. Spaniel is a package that builds on Seurat and SingleCellExperiment for interoperability and implements QC plots that help the user to remove ST or Visium spots outside the tissue. However, Spaniel's main difference from STUtility is that Spaniel can create a `shiny` app for interactive visualization and exploration of the data. While this may make Spaniel sound unremarkable, it was written about a year before Seurat supported spatial data. Another specialized package is SpatialCPie (Bergenstråhle, Bergenstråhle, and Lundeberg 2020), which also uses `shiny` for interactive visualization. SpatialCPie cluster ST or Visium data at multiple resolutions and plots a graph showing how clusters from one resolution relates to those from other resolutions. It also plots a pie chart at each ST or Visium spot, on top of an H&E background, showing similarity of each spot to each cluster, to give a more nuanced view than simply coloring the spots by cluster. Both packages are on Bioconductor.

#### 7.3 Spatial reconstruction of scRNA-seq data

It may be fair to say that the holy grail of spatial transcriptomics is to profile the whole transcriptome at single cell resolution and without dropouts. We

[69] `https://shiny.rstudio.com`

have already seen that, with seqFISH+ and ExM-MERFISH, this goal seems to possibly be within reach. However, the goal may be further than is seemingly the case, as the smFISH based techniques are still not generally applied to more than a few dozens to a few hundreds of genes, in the order of 10,000 cells (Figure 5.19, Figure 5.21), which only covers a small area of tissue. Meanwhile, techniques without single cell resolution and with lower detection efficiency but can cover large swaths of tissue have grown in popularity (Figure 5.33). Hence spatial transcriptomics has not supplanted scRNA-seq – which has also grown tremendously in popularity in recent years (Svensson, da Veiga Beltrame, and Pachter 2020) – but remains a complement. Spatial data that is not transcriptome wide can be complemented by scRNA-seq for information of other genes; this section reviews computational methods that map cells from scRNA-seq to spatial locations with a small panel of landmark genes and/or to impute gene expression not profiled by the spatial reference in space, or in short spatial reconstruction of scRNA-seq data. These are the most common types of data analysis(Figure 7.4). The two tasks are related but distinct, as when cells from scRNA-seq are mapped to spatial locations, spatial patterns of the genes expressed in the cells are also predicted. However, gene expression can also be predicted at spatial locations without mapping cells to the locations. Spatial data that does not have single cell resolution can be complemented by scRNA-seq for cell type deconvolution of the spots (Section 7.4). In turn, spatial data complements scRNA-seq with spatial information such as gene expression patterns and cell neighborhoods.

Attempts at spatial reconstruction of single cell data date back to 2014, when growth in the popularity of scRNA-seq started to pick up pace (Svensson, da Veiga Beltrame, and Pachter 2020). Early (2014-2017) methods tend to fall in three categories: direct dimension reduction with PCA, ad hoc scoring, and pseudotime projected into space. The first two have been by and large abandoned due to their limitations, and the third isn't commonly used. Another category is generative modeling, which we consider intermediate due to its early origin and lasting legacy as some later methods involve more sophisticated generative modeling. Later (2018-present) methods commonly involve a lower dimensional latent space shared by the scRNA-seq and the spatial data, and many different approaches have been tried to obtain the shared latent space and project it back into the higher dimensional space of gene expression. However, other principles were used as well, such as optimal transport, nonlinear direct dimension reduction, black box machine learning, mixture of experts model, and etc.

**Table 7.3:** Packages mentioned for spatial reconstruction of scRNA-seq data

| Name | Language | Title | Date published |
|---|---|---|---|
| Seurat1[70] | R | Spatial reconstruction of single-cell gene expression data[71] | 2015-04-13 |

| | | | |
|---|---|---|---|
| DistMap[72] | R | The Drosophila embryo at single-cell transcriptome resolution[73] | 2017-10-13 |
| gimVI[74] | Python | A joint model of unpaired data from scRNA-seq and spatial transcriptomics for imputing missing gene expression measurements[75] | 2019-05-06 |
| Seurat3[76] | R | Comprehensive Integration of Single-Cell Data[77] | 2019-06-13 |
| LIGER[78] | R; C++ | Single-Cell Multi-omic Integration Compares and Contrasts Features of Brain Cell Identity[79] | 2019-06-13 |
| SPRESSO[80] | R; Python | Novel computational model of gastrula morphogenesis to identify spatial discriminator genes by self-organizing map (SOM) clustering[81] | 2019-08-29 |
| Harmony[82] | R; C; C++ | Fast, sensitive and accurate integration of single-cell data with Harmony[83] | 2019-11-18 |
| novoSpaRc[84] | Python | Gene expression cartography[85] | 2019-11-20 |
| sstGPLVM[86] | Python | A Bayesian nonparametric semi-supervised model for integration of multiple single-cell experiments[87] | 2020-01-21 |
| SpaOTsc[88] | Python | Inferring spatial and signaling relationships between cells from single cell transcriptomic data[89] | 2020-04-29 |
| st_analysis[90] | Python | Molecular atlas of the adult mouse brain[91] | 2020-06-26 |
| GLISS[92] | NA | Integrative Spatial Single-cell Analysis with Graph-based Feature Learning[93] | 2020-08-13 |
| Tangram[94] | Python | Deep learning and alignment of spatially-resolved whole transcriptomes of single cells in the mouse brain with Tangram[95] | 2020-08-30 |
| SpaGE[96] | Python | SpaGE: Spatial Gene Enhancement using scRNA-seq[97] | 2020-09-21 |
| FIST[98] | MATLAB | Imputation of Spatially-resolved Transcriptomes by Graph-regularized Tensor Completion[99] | 2021-04-07 |

| | | | |
|---|---|---|---|
| LIGER[100] | R; C++ | Iterative single-cell multi-omic integration using online learning[101] | 2021-04-19 |

##### 7.3.1 Direct dimension reduction

As already mentioned in our summary of Puzzle Imaging, spatial reconstruction of dissociated tissue can be considered a dimension reduction problem. Here with scRNA-seq, the high dimensional gene expression data is directly projected to 1 to 3 dimensions that correspond to the spatial dimensions.

One of the earliest reconstruction methods (2014) maps single cell qPCR data onto a sphere that mimics the developing mouse otocyst (Durruthy-Durruthy et al. 2014). Ninety six genes were profiled with qPCR in single cells, and the gene expression profiles were projected to the first 3 principal components (PCs), which are then projected onto the surface of a sphere. The sphere is oriented on the dorsal-ventral (DV), anterior-posterior (AP), and left-right (LR) axes by expression of marker genes known to be expressed in one end of those axes. At least for the otocyst, this approach seemed to recapitulate expression patterns of many genes, at least qualitatively, at the resolution of octants. This approach was later adapted to reconstruct the human (Durruthy-Durruthy et al. 2016) and mouse (Mori et al. 2017) blastocysts. A one dimensional version of this approach was also adapted to spatially reconstruct cells from the organ of Corti along the apical and basal axis, though the PCA was performed only on DE genes between apical and basal cells and 2 PCs were projected to 1 dimension (Waldhaus, Durruthy-Durruthy, and Heller 2015).

Direct dimension reduction is still used after 2018, with dimension reductions other than PCA. Another form of dimension reduction for spatial reconstruction is the self-organinzing map (SOM)[102] as in the package SPRESSO (Mori et al. 2019). The Geo-seq mid-gastrula mouse embryo data (Peng et al. 2016) was reconstructed in 3D with genes selected from GO terms; 18 genes selected from a few GO terms could place all microdissected samples into the correct AP/LR quadrant with SOM. However, such genes were found by checking the SOM projections from thousands of GO combinations against the Geo-seq ground truth and may not apply to other biological systems. Also, the spatial reconstruction along the DV axis was not checked, though in Geo-seq, the samples were microdissected along the DV axis with a cryotome in addition to dissection into AP/LR quadrants with LCM.

A more recent, graph based dimension reduction is GLISS (Zhu and Sabatti 2020). After using a Laplacian score[103] based method to identify landmark genes from spatial data (to be reviewed in Section 7.5), a graph is constructed for the

[102] https://en.wikipedia.org/wiki/Self-organizing_map

[103] https://proceedings.neurips.cc/paper/2005/file/b5b03f06271f8917685d14cea7c6c50a-Paper.pdf

scRNA-seq data based on similarity in expression profiles of the landmark genes among cells as a proxy to spatial locations. With this graph, a new set of genes whose expression depend on the structure of the graph, or spatially variable genes, are identified, and added to the landmark genes. A new similarity graph is then constructed with both the landmark genes and spatially variable genes, and the dimension reduction is the eigenvectors of the graph Laplacian[104] of this graph, starting from the second eigenvector. One dimensional projection would be the second eigenvector. Two dimensional projection would be the second and third, and so on.

Ligand-receptor (L-R) pairs have also been used for direct dimension reduction, in CSOmap (Ren et al. 2020). Expression of L-R pairs in scRNA-seq cells is used to construct a cell-cell affinity matrix, with higher affinity meaning that two cells are more likely to be close to each other. Then an algorithm similar to tSNE is used to project the affinity matrix into 3 dimensions, corresponding to the physial dimensions. The Kullback–Leibler (KL) divergence[105] between the affinity and probability of the two cells to be neighbors is minimized, with constraints of the minimum physical size of the cell and the amount of space available.

##### 7.3.2 *Ad hoc* scoring

The methods above tend to only capture simple spatial patterns with simple gradients along axes, or have low resolution that is effectively restricted to octants or quadrants. More complex patterns with higher resolution can be reconstructed qualitatively with some score that measures similarity between each cell in scRNA-seq and each location in a spatial reference for the genes present in both datasets and favors genes more specific to a subset of cells. The spatial pattern of the score is the predicted gene expression pattern. As the score is qualitative and does not utilize statistical modeling of the data, this is called ad hoc scoring. The spatial reference is FISH (not smFISH) data of a panel of genes, with images for different genes registered onto a common coordinate system. As FISH is not very quantitative, both the spatial and the scRNA-seq data are binarized into "on" and "off" for each gene, and the predicted gene expression patterns based on the score is binarized as well since the score is only qualitative. Such approach is simple to implement, but the binarization misses quantitative nuances of gene expression patterns.

Ad hoc scoring has been used in *Platynereis dumerilii* brains; the FISH atlas was broken into voxels 3 $\mu$m on each side, smaller than the average single cell, and 98 landmark genes in the atlas used to predict patterns of other genes in scRNA-seq with a score (Achim et al. 2015). A different method, DistMap, uses a score based on Matthew correlation coefficient (MCC)[106] was used to soft assign

[104] `https://csustan.csustan.edu/~tom/Clustering/GraphLaplacian-tutorial.pdf`
[105] `https://en.wikipedia.org/wiki/Kullback-Leibler_divergence`
[106] `https://en.wikipedia.org/wiki/Matthews_correlation_coefficient`

cells from scRNA-seq to locations in the BDTNP atlas with 84 landmark genes and to predict expression patterns of the other genes (Karaiskos et al. 2017). The latter method inspired the DREAM Single-cell transcriptomics challenge in 2018 (Tanevski et al. 2020), a competition in which participants select the most informative genes and predict cell locations with 60, 40, and 20 of the 84 BDTNP landmark genes. At least some participating teams adapted the scoring method used in the original DistMap after selecting genes with their own methods (Alonso, Carrea, and Diambra 2020; Pham et al. 2020).

##### 7.3.3 Generative models

Many areas in spatial transcriptomics data analysis describe the data with a plausible statistical model and fit such a model to the data. Generative models have several advantages. First, uncertainties in parameter estimates and model predictions can be computed. Second, the model is more explainable, i.e. that humans may understand contributions of variables to the fitted model. Explainability plays an important role in models identifying spatially variable genes. As already mentioned, some of the segmentation-free smFISH or ISS analysis packages, such as pciSeq, rely on generative models. Generative models are used for spatial reconstruction of scRNA-seq data as well.

The popular scRNA-seq EDA package Seurat originated from spatial reconstruction of scRNA-seq data in 2015, to map cells from scRNA-seq to a WMISH reference with 47 landmark genes (Satija et al. 2015). The WMISH images were mostly obtained from ZFIN, and divided into 128 bins, which was then collapsed into 64 due to LR symmetry. As WMISH is not very quantitative, the WMISH reference was binarized. Due to the sparsity of scRNA-seq data, the normalized scRNA-seq data was smoothed. Then a mixture of 2 Gaussian distributions was fitted to each gene, for the "on" and the "off" states. With such distributions, the posterior probability that each cell comes from each bin can be calculated with the probability that the cell is "on" of "off" like in the bin for the 47 genes, although cells can very well have intermediate and more nuanced gene expression. The spatial centroid of each cell is the center of mass of the spatial map of the posterior probabilities. So far, the landmark genes have been assumed to be independent, which is unrealistic. Centroids that are close to actual bins are then used to calculate a covariance matrix of a subset of the landmark genes for each bin, with which the Gaussian mixture models and posterior probabilities are updated. While this model seems reasonable, it is no longer used, likely because of the advances in highly multiplexed smFISH and ISS that produced quantitative spatial references that do no need binarization for some tissues, especially the mouse brain. Nevertheless, the scRNA-seq part of Seurat lived on. As already mentioned, WMISH or ISH atlases are the only spatial transcriptomics resources available for some biological systems and most of the atlases are not transcriptome wide, so this method can still be useful.

A different generative model was used to map scRNA-seq cells to a smFISH atlas in the mouse liver (Halpern et al. 2017). Six marker genes known to be patterned in the portal-central axis of the hepatic lobule were profiled with smFISH. Then the smFISH data was binned into 9 zone, normalized, and each gene in each zone was modeled with a gamma distribution, which was then multiplied by coefficients correcting for the fact that only part of the cell is in the tissue section for the $\lambda$ of a Poisson distribution to form a negative binomial distribution. The negative binomial distribution was sampled and normalized for the whole cells in scRNA-seq and proportion of UMIs from the gene of interest, which would approximate the distribution of a cell in each zone having expression levels of the gene of interest. The prior probability of a hepatocyte originating from each zone seems to be the relative area of the concentric ring that is each zone, centered on the central vein. With the prior and the sampled distribution of expression of marker genes, the posterior probability of each cell from each zone can be calculated with Bayes rule. To impute expression of genes other than the 6 markers in each zone, the gene count matrix is multiplied to the posterior probability matrix (after weighing the probabilities). Here the 6 markers are assumed to be independent, which might not be realistic. The same approach is still used by the same lab for more recent liver datasets (Halpern et al. 2018; Droin et al. 2020), although we are unaware of its use outside that lab.

Some of the shared latent space methods are based on generative models as well, with the latent space as part of the model. In gimVI (Lopez et al. 2019), which is adapted from scVI specifically to impute gene expression in space by integrating spatial and scRNA-seq data, gene expression in scRNA-seq is modeled with the negative binomial (NB) or zero inflated negative binomial (ZINB) distribution, and the spatial data is modeled with the Poisson or NB distribution (depending on the technology and dataset). The scRNA-seq and spatial data are modeled as coming from a shared latent lower dimensional space, which is decoded back to the higher dimensional gene expression space by a neural network to capture nonlinear structures as part of the mean parameters of the NB, ZINB, or Poisson distributions. The latent space is estimated when the model is fitted with variational Bayesian inference. To impute gene expression in space, the latent space is sampled and passed through the decoding neural network to get the mean parameters of the gene expression distributions for spatial data.

Another generative model with a shared latent space is semi-supervised t-distributed Gaussian process latent variable model (sstGPLVM) (Verma and Engelhardt 2020). The scRNA-seq or spatial data is modeled as coming from a noisy sample in high dimension from a lower dimensional shared latent space. The latent space can be concatenated to fixed covariates such as batch, technology used to collect data, spatial coordinates, and etc. and is estimated with black box variational inference. Missing data in gene expression and covariates can be estimated from the latent space, thus enabling mapping scRNA-seq cells to spatial coordinates and imputing gene expression, and the latent space can be collapsed across a covariate to remove its effect. The latent space has a Gaussian prior with identity variance. The prior of the high dimensional noiseless space is a Gaussian process with covariance between cells defined by a kernel that is a weighted sum of Matern 1/2[107] and Gaussian kernels to allow for a non-smooth manifold that better represents data. The input to the kernel is a weighted sum (length scales of kernel) of l1 distance between the cells in the latent space (including the covariates). The noise added to the noiseless high dimensional space to model actual data is a heavy tailed Student's t distribution[108], to account for overdispersion and non-Gaussian distribution of the data. This method is not specifically designed for spatial data, but can be used to integrate different scRNA-seq datasets as well.

##### 7.3.4 Shared latent space

There are some additional methods that project scRNA-seq and spatial data into a shared latent space to impute gene expression in space but without generative modeling. Some of them are designed for data integration in general, but included here the authors demonstrated integration of scRNA-seq and spatial data, seeming to intend their packages for such usage.

In version 3 of Seurat (Stuart et al. 2019), the scRNA-seq and spatial datasets are projected into a shared latent space by canonical correlation analysis (CCA)[109], which finds a low dimensional space that maximizes correlation between the two dataset, or by projecting one dataset into a low dimensional PCA space of the other dataset. Then anchor cells are identified, as cells in the two datasets with sufficient shared neighborhood, and the weight of each anchor on each cell in the spatial dataset is calculated by ad hoc scoring favoring closeness in the latent space and more similar shared neighborhood to the anchor. Gene expression is then simply transferred from scRNA-seq to spatial data by multiplying the normalized gene count matrix of genes absent from the spatial data in scRNA-seq with the anchor weight matrix.

LIGER (Welch et al. 2019) is a different data integration method, of which a Seurat wrapper has been implemented. The latent space is inferred by integrative NMF, which finds a set of factors unique to the scRNA-seq or the spatial dataset, and a set of factors shared by both. Gene expression is imputed in spatial data by averaging the expression of genes of interest in the $k$ (50) nearest neighbors (kNN) from the scRNA-seq data in the space spanned by the shared factors.

In SpaGE (Abdelaal et al. 2020), a common latent space is inferred as such: gene shared by the spatial dataset and scRNA-seq are used to do PCA independently for the two datasets. Then the cosine similarity matrix of the PCs of the two dataset is passed to singular value decomposition (SVD)[110]. Then the left and

[107] https://en.wikipedia.org/wiki/Matérn_covariance_function
[108] https://en.wikipedia.org/wiki/Student%27s_t-distribution
[109] https://en.wikipedia.org/wiki/Canonical_correlation
[110] https://en.wikipedia.org/wiki/Singular_value_decomposition

and right singular vectors are used to align the PCs to a common latent space of principal vectors. The original data is projected into the space spanned by the principal vectors of the scRNA-seq data. Then kNN is used to project gene expression from scRNA-seq to spatial data.

In Harmony (Korsunsky et al. 2019), the data, with different batches, is first PCA projected. Then the PCA projection is clustered with an altered k-means clustering[111] algorithm that assigns cells probabilistically to clusters and maximizes diversity in batches in each cluster. Then the batch correction is found by mixture of expert model[112]. In each cluster, the PCA projection is modeled by a linear combination of variables in the design matrix (containing batch information), with an intercept term for batch free variation in each cluster. The batch correction term is a weighted sum of the linear model predictions excluding the intercept term, weighted by the probabilistic assignment of each cell to each cluster. Then the batch correction term is subtracted from the original PCA projection. The clustering and correction are repeated until convergence. This way, the cells from scRNA-seq and spatial data are aligned in a common latent space. Then gene expression is imputed in spatial data with kNN.

##### 7.3.5 Other principles

Approaches that do not fall into the categories reviewed above are reviewed in this subsection, including projecting pseudotime into space, black box machine learning, and optimal transport.

In some biological systems, cell differentiation corresponds to physical locations of the cells, so pseudotime, which supposedly arranges cells along differentiation trajectories, have been mapped to space, thus placing dissociated cells in space. For instance, in the bone growth place, cells at different stages of differentiation are physically arranged along the length of the bone in a cylinder, so the pseudotime trajectory of the cells was simply warped into a straight line for spatial reconstruction (Li et al. 2016). Similarly, in Drosophila larva, cell differentiation corresponds to the proximal-distal axis in the antenna disk and the AP axis in the eye disk, so cells from both scRNA-seq and scATAC-seq were binned according to pseudotime and assigned to the corresponding bins in the eye-antenna disk (Bravo González-Blas et al. 2020). However, this would not work in tissues without such neat correspondence, such as the Drosophila embryo, in which some genes are expressed in periodic patterns to specify segments.

Deep learning libraries such as PyTorch also made it more effective to predict locations for scRNA-seq cells without a pre-conceived statistical model of the data. For instance, after data normalization and batch correction, a deep neural network can be trained on ST data with annotations of spatial regions to

[111] `https://medium.com/analytics-vidhya/k-means-clustering-explained-419ee66d095e`
[112] `https://people.cs.pitt.edu/~milos/courses/cs2750-Spring04/lectures/class22.pdf`

predict spatial regions for scRNA-seq data (Ortiz et al. 2020). In addition, PyTorch's gradient-based optimization has been used to probabilistically map scRNA-seq cells to spatial locations in Tangram (Biancalani et al. 2020). The spatial reference is voxelated, and a mapping matrix of probability of each cell mapping to each voxel is inferred by minimizing KL divergence between mapped and actual cell density in each voxel and favoring stronger correlation between mapped data and the spatial reference in expression of each gene across voxels and gene expression profiles of each voxel.

Thus far, the reconstruction methods do not take spatial autocorrelation, i.e. that cells physically closer to each other are more likely to have more similar gene expression profiles, in the spatial data into account. Optimal transport[113], i.e. finding a way to transport a pile of dirt from one place to others with minimum cost, has been used to exploit spatial autocorrelation to map scRNA-seq cells to spatial locations. In novosparc (Nitzan et al. 2019), neighborhood graphs are constructed for scRNA-seq in gene expression space and for spatial reference data in physical space. Then assuming spatial autocorrelation, optimal transport is used to place cells in locations to make the two graphs match. This can be done without gene expression data in the spatial grid, but can be improved with a spatial gene expression reference. In SpaOTsc (Cang and Nie 2020), first an optimal transport plan from scRNA-seq cells to spatial locations is inferred with gene expression dissimilarity matrices between scRNA-seq cells and between cells and locations and a spatial distance matrix between spatial locations. Then a spatial distance matrix for scRNA-seq cells is imputed based on that optimal transport plan. The plan can also be used to impute gene expression in space. SpaOTsc also uses optimal transport to infer cell-cell interaction, to be reviewed in the Cell-cell Interaction section. A drawback of this kind of method is that because different cell types can mix in the same spatial neighborhood, such as hepatocytes and Kupffer cells in the liver, spatial autocorrelation is not absolute.

Spatial autocorrelation can also be utilized without optimal transport, but with tensor completion in Canonical Polyadic Decomposition (CPD)[114] form as in FIST (Li et al. 2020). The spatial data can be viewed as a 3 dimensional tensor, with the x and y coordinates and gene expression at each location ((or 4 with z coordinate). CPD is used to improve computational efficiency. In CPD, the tensor is approximated with a sum of rank 1 tensors, i.e. cross products of 3 vectors, one for each dimension. This decomposition, with extra dimensions for unknown gene expressions, is found by minimizing the difference between the reconstructed tensor with the existing tensor for known genes and by favoring spatial autocorrelation of gene expression on a neighborhood graph and favoring similarity of expression of genes with similar functions in a protein-protein interaction graph.

[113] https://www.stat.cmu.edu/~larry/=sml/Opt.pdf
[114] https://www.tensorlab.net/doc/cpd.html

#### 7.4 Cell type deconvolution

There is another aspect to how spatial and scRNA-seq data complement each other. In array based techniques that do not have single cell resolution, the cell type composition of each spot can be estimated with scRNA-seq data. Perhaps because of the increasing popularity of ST and Visium, several cell type deconvolution methods have been developed in the past year, falling into three categories: part of other packages, NMF, and statistical modeling. While any tool designed for cell type deconvolution of bulk RNA-seq data can be used, this section specifically focuses on cell type deconvolution tools designed with spatial data in mind.

**Table 7.4:** Packages mentioned for cell type deconvolution

| Name | Language | Title | Date published |
|---|---|---|---|
| NMFreg[115] | Python; MATLAB | Slide-seq: A scalable technology for measuring genome-wide expression at high spatial resolution[116] | 2019-03-29 |
| Seurat3[117] | R | Comprehensive Integration of Single-Cell Data[118] | 2019-06-13 |
| RCTD[119] | R | Robust decomposition of cell type mixtures in spatial transcriptomics[120] | 2020-05-08 |
| Tangram[121] | Python | Deep learning and alignment of spatially-resolved whole transcriptomes of single cells in the mouse brain with Tangram[122] | 2020-08-30 |
| stereoscope[123] | Python | Single-cell and spatial transcriptomics enables probabilistic inference of cell type topography[124] | 2020-10-09 |
| DSTG[125] | Python | DSTG: Deconvoluting Spatial Transcriptomics Data through Graph-based Artificial Intelligence[126] | 2021-01-22 |
| SPOTlight[127] | R | SPOTlight:Seeded NMF regression to Deconvolute Spatial Transcriptomics Spots with Single-Cell Transcriptomes[128] | 2021-02-05 |

| | | | |
|---|---|---|---|
| Giotto[129] | R | Giotto, a pipeline for integrative analysis and visualization of single-cell spatial transcriptomic data[130] | 2021-03-08 |

Some of the packages already mentioned in previous sections have cell type deconvolution functionalities as well. For instance, Seurat's data transfer based on anchors between datasets can also be used to transfer cell type annotations, and the *ad hoc* score for the transferred cell types has been used as a qualitative measure of cell type composition in Visium spots (Mantri et al. 2020). Giotto implements 3 methods for qualitative cell type deconvolution: First, a score based on fold change in expression of marker genes in a spot compared to the mean across spots. Second, another score scoring genes for specificity in both scRNA-seq cell types and ST or Visium spots and the sum of the top 100 gene scores is the cell type enrichment score for each spot. For these two methods, p-values are calculated from permutation testing[131]. Third, given a fixed set of cell type marker genes, a hypergeometric test[132] is used to test for enrichment of marker genes among top 5% expressed genes of the spot. In Tangram, the cell mapping matrix from scRNA-seq to ST or Visium can be inferred as the ground truth cell density per spot can be measured from H&E staining. When cells from scRNA-seq are mapped to spots in ST and Visium, the cell type annotations are also mapped.

In both the prequel and current era, NMF[133] is quite popular among data analysis methods as the factors (cell embeddings) and the gene loadings tend to exhibit block like structures and the values of the basis and the loadings are enforced to be non-negative, corresponding to the non-negative nature of gene expression data and making the results more interpretable. The blocks in the factors may reflect cell types or clusters, and the blocks in gene loadings may reflect cell type marker genes. NMF has been used for cell type deconvolution as well. To address slide-seq (version 1)'s lack of single cell resolution and poor efficiency, NMFreg was developed to reconstruct the expression profile of each spot as a weighted sum of cell type signatures from scRNA-seq (Rodriques et al. 2019). First, scRNA-seq gene count matrix of cell types of interest and cell type annotations is decomposed with NMF, and each factor is assigned to a cell type and one cell type can have multiple factors. Then non-negative least squares[134] is used to compute the weights of the weighted sum of the factors for each spot. As such weights tend not to cleanly assign spots to cell types, perhaps due to the sparsity of scRNA-seq and slide-seq data, the weights are then thresholded. The threshold is the maximum cell loading of cells not assigned to the cell type of interest among in the factors of this cell type. The weights for this cell type

[131] https://www.r-bloggers.com/2019/04/what-is-a-permutation-test/
[132] https://brilliant.org/wiki/hypergeometric-distribution/
[133] http://www.cs.cmu.edu/~11755/lectures/Lee_Seung_NMF.pdf
[134] https://www.r-bloggers.com/2019/11/non-negative-least-squares/

is only kept if the $l_2$ norm[135] of the weight vector for these factors exceed the threshold. Another NMF based method, SPOTlight (Elosua et al. 2020), uses a very similar principle.

Cell type deconvolution can also be performed by explicitly modeling spot level gene expression in terms of individual cell types. In stereoscope (Andersson et al. 2019), a negative binomial distribution is fit to the expression of each gene in each cell type in scRNA-seq data. Then at each spot, gene expression is modeled as a weighted sum of the negative binomial distributions from each cell type, and the weights are estimated by maximum likelihood estimation (MLE)[136]. In Robust Cell Type Decomposition (RCTD) (Cable et al. 2020), gene expression at each spot is modeled as a Poisson distribution, whose mean is an expected rate scaled by total transcript count at the spot. The log rate is the sum of the log of weighted sum of mean gene expression for each cell type from a scRNA-seq reference, a fixed spot specific effect term, a gene specific platform random effect, and another gene specific random effect term for overdispersion[137]. The parameters, including cell type weights, are then estimated with MLE.

More recently, the graph convolutional neural network (GCN)[138] has been applied to cell type deconvolution, in DSTG (Su and Song 2020). First, scRNA-seq cells are randomly assigned to "spots" of 2 to 8 cells, forming a pseudo-ST dataset. Then the pseudo-ST and real ST data are projected to a CCA space, and a mutual $k$ nearest neighbor graph is built in this space. After that both the pseudo and real ST data and the graph are fed into a GCN, trained to minimize cross entropy between imputed cell composition and actual cell composition in the pseudo-ST spots. Finally, the trained model is used to predict cell composition in real ST data.

#### 7.5 Spatially variable genes

Some genes, such as house keeping genes, are ubiquitously expressed. Such genes, while highly variable at the single cell level, may be interspersed in space so they may not show a spatial trend. Expression of some genes depends on spatial location, which can be due to cell type localization or variation within or independent from cell types. One of the goals of early prequel studies was to identify spatially variable genes, which was done manually, which can be inconsistent and labor intensive. With more quantitative data and data analysis methods, the current era brought identification of spatially variable genes to the next level. Simple methods to identify such genes include dividing the extent of the tissue into a grid and use Fisher's exact test to test for non-random distribution of transcript counts in the grid, or to run DE between one region — be

[135] `https://mathworld.wolfram.com/L2-Norm.html`

[136] `https://brilliant.org/wiki/maximum-likelihood-estimation-mle/`

[137] `http://biometry.github.io/APES/LectureNotes/2016-JAGS/Overdispersion/OverdispersionJAGS.html`

[138] `https://tkipf.github.io/graph-convolutional-networks/`

it a grid cell or a manually annotated histological region — and another region. However, some more sophisticated methods have been developed that avoid the potential arbitrariness of grids and manual annotation, take advantages of increased resolution of spatial transcriptomics. This section reviews these computational methods that identifies genes with expression that depends on spatial locations. Two principles are the most common. One is Gaussian process regression[139] and generalization to discrete distributions with the log mean parameter modeled as Gaussian process. Another centers on Laplacian scores of graphs. There are also some additional methods using other principles.

**Table 7.5:** Packages mentioned for spatially variable genes

| Name | Language | Title | Date published |
|---|---|---|---|
| trendsceek[140] | R | Identification of spatial expression trends in single-cell gene expression data[141] | 2018-03-19 |
| SpatialDE[142] | Python | SpatialDE: identification of spatially variable genes[143] | 2018-03-19 |
| scGCO[144] | Python | Identification of spatially variable genes with graph cuts[145] | 2018-12-09 |
| Seurat3[146] | R | Comprehensive Integration of Single-Cell Data[147] | 2019-06-13 |
| RayleighSelection[14 | R; C++ | Clustering-independent analysis of genomic data using spectral simplicial theory[149] | 2019-11-22 |
| SPARK[150] | R; C++ | Statistical analysis of spatial expression patterns for spatially resolved transcriptomic studies[151] | 2020-01-27 |
| GPcounts[152] | Python | Non-parametric modelling of temporal and spatial counts data from RNA-seq experiments[153] | 2020-07-30 |
| GLISS[154] | NA | Integrative Spatial Single-cell Analysis with Graph-based Feature Learning[155] | 2020-08-13 |
| singleCellHaystack[1 | R | A clustering-independent method for finding differentially expressed genes in single-cell transcriptome data[157] | 2020-08-28 |
| SPATA[158] | R | Inferring spatially transient gene expression pattern from spatial transcriptomic studies[159] | 2020-10-21 |

[139]`https://bookdown.org/rbg/surrogates/chap5.html`

| | | | |
|---|---|---|---|
| Giotto[160] | R | Giotto, a pipeline for integrative analysis and visualization of single-cell spatial transcriptomic data[161] | 2021-03-08 |
| SOMDE[162] | Python | SOMDE: A scalable method for identifying spatially variable genes with self-organizing map[163] | 2021-03-24 |

##### 7.5.1 Gaussian process regression

Gene expression in space can be modeled as a 2D Gaussian process. Spatial dependence of gene expression from any finite collection of locations in space can be modeled with a joint multivariate Gaussian distribution, whose covariance matrix can be defined with a kernel, which is typically defined so spatially closer cells or spots have higher covariance.

SpatialDE (Svensson, Teichmann, and Stegle 2018) is one of the more popular methods to identify spatially variable genes. Spatial gene expression is modeled as a Gaussian process, in which the mean is the mean expression level of the gene, and the covariance matrix has a spatial and non-spatial component. The spatial component uses the Gaussian kernel, in which the covariance decays exponentially with squared distance between cells or spots, with rate of decay controlled by a length scale parameter. In the null model, the gene expression follows a Gaussian distribution without covariance between cells or spots. Then the model likelihood of the fitted full model and the null model are compared with log likelihood ratio test[164]. The log likelihood ratios under null model are asymptotically $\chi^2$ distributed, and this distribution is used to calculate the p-values of the test. If a gene is found to be significantly spatially variable, then the full model can be fitted with two other kernels, linear and periodic, and compared to the Gaussian kernel with Bayesian Information Criterion (BIC)[165] to discover linear and periodic patterns. As gene expression is discrete and not Gaussian, the data needs to be normalized before applying SpatialDE; even then, data normalization does not make the data Gaussian.

The discrete, non-Gaussian distribution of gene expression is directly modeled by SPARK (S. Sun, Zhu, and Zhou 2020). Gene expression is modeled by a Poisson distribution, with a rate parameter scaled by total transcript count at the spot or cell of interest. The log rate parameter contains a linear model for non-spatial variation in gene expression and can include cell or spot level covariates such as cell type, with non-spatial residuals. The spatial dependence is modeled by a zero mean Gaussian process with either a Gaussian or cosine kernel for the covariance matrix and 5 different length scale parameters are tried for each kernel type, so 10 kernels are tried. The model is fitted with one

[164] `https://www.probabilitycourse.com/chapter8/8_4_5_likelihood_ratio_tests.php`
[165] `https://en.wikipedia.org/wiki/Bayesian_information_criterion`

kernel at a time, with a penalized quasilikelihood algorithm[166]. The p-values are estimated by Satterthwaite method[167], and the p-values from the 10 kernels are combined with the Cauchy p-value combination rule[168].

Gene expression data may better be modeled with NB or ZINB, which is done in GPcounts (BinTayyash et al. 2020). The log of the mean parameter of the NB or ZINB, scaled by total transcript count at the cell or spot, is modeled with a Gaussian process with Gaussian kernel for covariance. For ZINB, the dropout probability is related to the NB mean by a Michaelis-Menten equation[169]. For one sample, the null hypothesis a constant model, a Gaussian with fixed mean and no covariance between cells or spots, i.e. gene expression does not vary in space. Spatially variable genes are identified with the log likelihood ratio test as in SpatialDE. For two samples, the null hypothesis is that two samples have the same gene expression pattern, and the alternative hypothesis is that two different Gaussian processes are required to model the two samples. Three models are fitted, one for each sample and another fit with both samples as replicates, and the SpatialDE likelihood ratio test is used to compare the separate models to the shared one. The models are fitted with a sparse approximation of variational Bayesian inference.

The size of the covariance matrix of the cells or spots grows quadratically with the number of cells or spots. To speed up computation, SMODE aggregates cells or spots into nodes with SOM, reducing the size of the covariance matrix, before proceeding to a SpatialDE-like test (Hao, Hua, and Zhang 2021).

##### 7.5.2 Laplacian score

GLISS (Zhu and Sabatti 2020) has already been mentioned as a method to reconstruct scRNA-seq data in space by projecting scRNA-seq data into a 1 to 3 dimensions that stand for spatial dimensions. The first step of GLISS is to identify spatially variable genes in the spatial reference as landmark genes. In 2005, the Laplacian score was proposed as a method of feature selection, which favors features that preserves the local structure of the data in the feature space and has large variance (He et al. 2020). In GLISS, a spatial neighborhood graph is constructed on the spatial reference; two cells or spots have larger edge weight if they are physically close to each other. By default, the graph is a mutual nearest neighbor graph, in which cells or spots are nodes and an edge connects two nodes if they are mutual $k$ nearest neighbors. Then for each gene, a Laplacian score is computed using the gene of interest and the

[166] https://academic.oup.com/bioinformatics/article/35/3/487/5055584
[167] https://www.jstor.org/stable/3002019?casa_token=qXYt_raEK8IAAAAA%3A3bbUKlAh6ruVm7KRvoEpjPHfPM_qf4tnaRUadXujACmbEiXoUPINFhTMrE3m7GUTYjgPEYp1i8nIZ1ktuKs3Z5aX2alhiH0pMSnpKB5tiS8dWxA1Cg&se[illegible]1#metadata_info_tab_contents
[168] https://www.jstor.org/stable/43974709?casa_token=GvHZB2UWAJIAAAAA%3A4h2Wa4vbQog_b_oIXg0uFh4HJeSp-2qHllLvx9K3m6of8C86U0Be_Y8CIGyZ7q0JMl2JPMeKvX1wMHSlxKVedW2-KMzR-wx74wEyqO1Rg74piqmtaw&seq=1#metadata_info_tab_contents
[169] https://en.wikipedia.org/wiki/Michaelis-Menten_kinetics

graph Laplacian of the spatial neighborhood graph. Genes with low Laplacian scores are chosen as landmark genes, as a low score favors similarity of gene expression in nearby cells or spots and large variance among the spots, which means spatially coherent regions with high and low expression of the gene. The p-value of the gene is computed by permuting expression of the gene of interest among cells and recomputing the score.

The simplical complex[170] is a generalization of the graph that not only includes nodes and edges but also triangles, tetrahedrons, and their higher dimensional generalizations. RayleighSelection implements generalizations of the Laplacian score for simplical complexes for clustering-free DE (Govek, Yamajala, and Camara 2019). The 1-dimensional Laplacian score, a generalization in which gene expression values are attributed to edges rather than nodes, has been used for DE in scRNA-seq data. The nodes here are clusters of cells and two nodes are connected by an edge when they intersect, as in topological data analysis (TDA)[171] (Rizvi et al. 2017). P-values of genes were computed by permutation test, permuting expression of a gene of interest among cells. For spatial data, the spatial neighborhood graph was created as the Vietoris-Rips complex[172]. The 0-dimensional, which is the same as the original Laplacian score, was used to identify spatially variable genes. The graph was also created for cells from pairs of cell types and the Laplacian score, with feature as cell type label, was used to identify cell type colocalization.

##### 7.5.3 Other principles

A spatial point pattern is the observed spatial locations of things or events, and a point process is a stochastic mechanism that generated the point pattern. As already mentioned, in pciSeq, transcript spot locations are modeled by a Poisson point process whose intensity itself is modeled with a Gamma distribution. Cell locations can also be modeled as a point process, which is done in trendsceek (Edsgärd, Johnsson, and Sandberg 2018). Each point in a spatial point process can have additional properties other than location, such as gene expression and cell type, which are called marks. If the marks are completely randomly distributed in space, then points with one mark would not be more or less likely to be near points with the same (for categorical marks) or similar (quantitative marks) marks than to points with dissimilar marks. To identify spatial distribution of gene expression that deviates from complete randomness, trendsceek uses 4 mark-segregation summary statistics, which are functions of distance between two points, taking the expected value of a summary statistics on the marks of every pair of points sepearated by the given distance: Stoyan's mark-correlation function (squared geometric mean of marks of two points normalize by squared mean of marks), mean-mark function (average of marks in two

[170] https://www2.cs.duke.edu/courses/fall06/cps296.1/Lectures/sec-III-1.pdf
[171] https://medium.com/@varad.deshmukh/topological-data-analysis-a-very-short-introduction-611d3238a0bd
[172] https://en.wikipedia.org/wiki/Vietoris-Rips_complex

points), variance-mark function (variance of the marks given distance between points), and mark-variogram[173] (squared difference of marks of two points). Permutation testing is used to calculate p-values. Regions of interest in the tissue are the regions with $p < 0.05$ from the permutation testing. Perhaps due to the permutation, trendsceek seems to be less scalable and less sensitive than SpatialDE and SPARK (S. Sun, Zhu, and Zhou 2020; Zhang, Feng, and Wang 2018).

Seurat's spatial functionalities include finding spatially variable genes, which currently provides two methods, one of this is mark-variogram, inspired by trendsceek. The other is Moran's I[174], which is a common summary statistics of spatial autocorrelation, as spatially patterned genes also exhibit autocorrelation. However, the problem with Moran's I is that when almost all cells take very similar values in gene expression, such as when a gene is constitutively expressed or expressed by very few cells, Moran's I will still indicate strong spatial autocorrelation although the gene is not actually spatially variable.

Giotto (Dries et al. 2019) implements 3 simple and fast methods to find spatially variable genes in addition to wrapping SpatialDE and trendsceek. First, a spatial neighborhood graph is constructed, which can be mutual $k$ nearest neighbors graph, a graph placing an edge when two cells are within a certain distance, or Delaunay triangulation. Then the gene expression is binarized. The first method uses the silhouette score[175]. In clustering, a measure of whether each point should be assigned to its current cluster or it should better be assigned to a neighboring cluster. The mean silhouette score indicates how tight and segregated the cluster are. Here the clusters are cells expressing the gene of interest and those not expressing. Then a high silhouette score means that cells expressing the gene and those not expressing are well-segregated in space, which means the gene is spatially variable. The second and third method only differ in the way gene expression is binarized. The second uses k-means with $k = 2$, and the third uses a threshold. Then a contingency table $M$ is constructed from neighboring cells in the graph expressing or not expressing the gene; each row is whether a cell expresses the gene, and each column is whether its neighbor also expresses it, so $M_{1,1}$ is the number of distinct pairs of cells both expressing the gene, $M_{1,2}$ is the number of pairs of cells in which source cell is expressing the gene and target cell is not, and so on. Fisher's exact test[176] is used to test for dependency in gene expression on whether cells are neighbors.

The KL divergence is a measure of difference between two probability distributions. In singleCellHaystack (Vandenbon and Diez 2020), the cell density in the tissue (or a PCA, tSNE, or UMAP space) is estimated at grid points with Gaussian kernel density, and normalized to form a probability distribution

[173] https://rdrr.io/cran/spatstat/man/markvario.html
[174] https://pro.arcgis.com/en/pro-app/tool-reference/spatial-statistics/h-how-spatial-autocorrelation-moran-s-i-spatial-st.htm
[175] https://en.wikipedia.org/wiki/Silhouette_%28clustering%29
[176] https://mathworld.wolfram.com/FishersExactTest.html

of locations of cells. Then the probability distribution of whether a gene of interest is expressed at each grid point is compared to the cell density distribution with KL divergence. P-values are computed by permuting gene expression among cells. Again, this is a cluster-free DE method, not designed specifically for spatial data but can be applied to spatial data.

We have already mentioned Markov random field (MRF) models for partitioning a tissue section into cell types and cells. MRF has also been used to identify spatially variable genes, as in scGCO (single-cell graph cuts optimization) (Zhang, Feng, and Wang 2018). Expression values of a gene are binned into 2 to 10 categories with Gaussian mixture model clustering. Then a graph connecting cells in space is constructed over the tissue by Delaunay triangulation, and the graph, with the expression category of the gene, is modeled with a MRF. Then as the model favors neighbors in the graph with the same category, edges of the graph are cut to maximize the likelihood of the model, thus identifying not only regions of tissue with an expression category of the gene, but also genes forming such regions. As MRF enforces coherent regions of the tissue to take the same category, while when only gene expression, without spatial information, is considered, not all cells in the region warrant the category. Then the number of cells that truly deserve the category in each region is used to calculate statistical significance of the gene's spatial variability. The null hypothesis is a homogeneous Poisson point process, in which cells (points) are completely randomly distributed in space and the location of one cell is independent from the location of any other cell. The smallest p-value of any category and any region is reported for the gene of interest.

So far the methods identifying spatially variable genes based on Gaussian process regression commonly use the Gaussian kernel for the covariance matrix, which assumes that the gene expression modeled is weakly stationary, i.e. covariance only depends on distance between cells or spots. This does not take into account anisoptropy[177], i.e. spatial dependence of gene expression is different in different directions, observed in tissues such as the brain cortex and the hepatic lobule in which cell functions are primarily stratified along one direction or axis. SPATA (Kueckelhaus et al. 2020) implements a method to find spatially variable genes for such primary axis. With the interactive `shiny` app, the user defines this axis, which may or may not be a straight line, and cells within a certain distance from the axis are included for further analysis. Then among the included cells, gene expression and cell type annotations along the axis can be visualized in the `shiny` app. Then SPATA fits a variety of functions with known forms, e.g. linear or nonlinear descent or ascend, peaks, periodic, and etc. to the gene expression along the axis. For each function, the sum of the residuals is calculated and compared to find functions that better represent the change in gene expression along the axis to identify patterns.

[177]`https://arxiv.org/abs/1712.01634v2`

#### 7.6 Gene patterns

**Table 7.6:** Packages mentioned for gene patterns

| Name | Language | Title | Date published |
|---|---|---|---|
| SpatialDE[178] | Python | SpatialDE: identification of spatially variable genes[179] | 2018-03-19 |
| std-nb[180] | C++ | Charting Tissue Expression Anatomy by Spatial Transcriptome Decomposition[181] | 2018-12-28 |
| stLearn[182] | Python | stLearn: integrating spatial location, tissue morphology and gene expression to find cell types, cell-cell interactions and spatial trajectories within undissociated tissues[183] | 2020-05-31 |
| GLISS[184] | NA | Integrative Spatial Single-cell Analysis with Graph-based Feature Learning[185] | 2020-08-13 |

When spatially variable genes are identified, a question naturally arises: Are there archetypal patterns among these spatially variable genes? As already reviewed in Section 3, comparing and classifying gene expression patterns was a major topic in the prequel era. Such interest persists in the current era, although we find no evidence that current era gene pattern analysis is significantly influenced by the prequel antecedents, although factor analysis and NMF have been used in both eras.

The most straightforward way to identify archetypal gene patterns is to cluster the gene expression patterns and obtain the cluster centers to represent the cluster. This has been used to analyze mouse brain voxelation data in 2009 (An et al. 2009). Wavelet transform[186] was applied to the data and the Euclidean distance between the wavelet feature vectors was used to measure gene similarity. Gene similarity between pre-defined "typical" genes and other genes was one way to find groups of similar genes and k-means clustering is another. Some package already reviewed also have functionality to identify archetypal gene patterns. In DistMap and SPARK (Karaiskos et al. 2017; S. Sun, Zhu, and Zhou 2020), the gene patterns are clustered with hierarchical clustering[187], and the individual clusters are obtained by tree cut. In Giotto (Dries et al. 2019), a gene-gene correlation matrix (by default Pearson) is computed, which is then

[186] https://en.wikipedia.org/wiki/Wavelet_transform
[187] https://towardsdatascience.com/understanding-the-concept-of-hierarchical-clustering-technique-c6e8243758ec

hierarchically clustered. Then the mean or centroid of each cluster is taken to represent that cluster. SpatialDE (Svensson, Teichmann, and Stegle 2018) also clusters gene expression patterns, in automatic expression histology (AEH), which implements a Gaussian process generalization of Gaussian mixture model clustering[188]. The number of components is set by the user, and the model is fitted to infer the mean pattern of each component. In GLISS (Zhu and Sabatti 2020), the archetypal patterns are identified in the reconstructed latent space as gene expression in the latent space is spline smoothed[189], and the spline coefficients are clustered.

Beyond clustering, a common way to identify archetypal gene patterns is factor analysis[190]. This has already been done in the prequel era (Pruteanu-Malinici, Mace, and Ohler 2011), but is further developed in the current era. Factor analysis tries to model higher dimensional data as a linear combination of a smaller number of variables called "factors", and PCA is a type of factor analysis. A prostate cancer ST dataset has been modeled with Poisson factor analysis (Berglund et al. 2018). The observed UMI counts at each spot is modeled as a sum of factors, each of which is Poisson distributed, with its own rate parameter, which in turn depends on Gamma distributed factor, gene, and spot level parameters that may account for overdispersion though this model does not entirely capture the mean-variance relationship of NB. The parameters are estimated from MCMC[191] sampling of the posterior of this model. Once the parameters are estimated, the individual factors can be calculated from the parameters based on the model. The factors seem to indicate regions in the tumor, such as cancer, stroma, and regions with immune cell infiltration. As NB may describe gene expression better than the Poisson distribution, a NB adaptation of the above Poisson factor analysis model has been developed (Maaskola et al. 2018). The observed UMI count at each spot is modeled as a sum of NB factors, whose rate parameter can incorporate gene, spot, and experiment level covariates. The package stLearn (Pham et al. 2020), which also implements methods to identify cell-cell interactions and spatial regions, uses PCA, ICA[192], and factor analysis to detect microenvironments in the tissue as again, the factors can correspond to specific regions in the tissue.

#### 7.7 Spatial regions

As already mentioned in trendsceek and scGCO, the problem of identifying spatially variable genes is closely related to identifying regions in tissue defined by gene expression. When archetypal gene patterns are identified, a related question arises: Do the patterns define novel anatomical regions in the tissue?

[188]https://towardsdatascience.com/gaussian-mixture-models-d13a5e915c8e
[189]https://en.wikipedia.org/wiki/Smoothing_spline
[190]https://en.wikipedia.org/wiki/Factor_analysis
[191]https://en.wikipedia.org/wiki/Markov_chain_Monte_Carlo
[192]http://wwwf.imperial.ac.uk/~nsjones/TalkSlides/HyvarinenSlides.pdf

As seen in the previous section, archetypal gene patterns, such as in factors, can reflect tissue regions. There are also methods that identify such regions without first identifying spatially variable genes and/or archetypal gene patterns. In the prequel era (Chapter 3), some studies clustered the voxels based on gene expression to identify spatial regions in the tissue, with either k-means clustering or co-clustering (Zhang et al. 2013; Bohland et al. 2010; Ko et al. 2013), or with Potts model (Pettit et al. 2014). More sophisticated clustering methods have been developed in the current era tot identify spatial regions. However, as different cell types can reside in the same spatial neighborhood, and conversely, cells from one cell type can reside in different regions of the tissue, MRF has been used to find spatially coherent regions that can contain multiple cell types.

**Table 7.7:** Packages mentioned for spatial regions

| Name | Language | Title | Date published |
|---|---|---|---|
| smfishHmrf[193] | R; Python; C | Identification of spatially associated subpopulations by combining scRNAseq and sequential fluorescence in situ hybridization data[194] | 2018-10-29 |
| SSAM[195] | Python; C++ | Segmentation-free inference of cell types from in situ transcriptomics data[196] | 2019-10-13 |
| stLearn[197] | Python | stLearn: integrating spatial location, tissue morphology and gene expression to find cell types, cell-cell interactions and spatial trajectories within undissociated tissues[198] | 2020-05-31 |
| BayesSpace[199] | R; C++ | BayesSpace enables the robust characterization of spatial gene expression architecture in tissue sections at increased resolution[200] | 2020-09-05 |
| Baysor[201] | Julia | Bayesian segmentation of spatially resolved transcriptomics data[202] | 2020-10-06 |

SSAM (Park et al. 2019), already reviewed in Section 7.1, also uses clustering to identify tissue domains in smFISH or ISS data but without cell segmentation. StLearn (Pham et al. 2020) develops further on top of clustering. First, a pretrained CNN is used to extract a 2048 dimensional feature vector from the H&E image behind each ST or Visium spot. The cosine similarity between the feature vectors from neighboring spots is then calculated. To normalize data, the gene expression data is smoothed in space, and the smoothing is weighted by the cosine similarity of feature vectors between spots. Then the spots are clustered with Louvain[203] or k-means. A spatial $k$ nearest neighbor graph is constructed, and used to refine the clustering. If a gene expression based cluster is broken into multiple pieces in space, then those pieces would become subclusters. Singleton spots are merged with a nearby cluster if the singletons have enough spatial neighbors in that cluster.

BayesSpace (Zhao et al. 2020) incorporates both Gaussian mixture model clustering and MRF. The ST or Visium data is first projected to a low dimensional space, such as by PCA. Then for each spot, the low dimensional projection of that spot is modeled with a Gaussian mixture model, with a pre-defined number of components or clusters. The spatial neighborhood graph is simply the square grid of spots for ST and the hexagonal grid for Visium. The model has a MRF prior to encourage neighboring spots to be assigned to the same cluster. The cluster assignment is initiated with non-spatial clustering, and the parameters of the model are estimated by MCMC. In addition, BayesSpace can increase the resolution of ST and Visium. Each spot is subdivided and initiated with the dimension reduction values at the spot, and an additional parameter is added to the model that nudges the dimension reduction values at each sub-spot while preserving the sum at the spot level. The nudging parameters are estimated by MCMC along with with other parameters.

As already reviewed in Section 7.1, Baysor (Petukhov et al. 2020) uses MRF to delineate cell type regions in the tissue without cell segmentation. MRF is used to identify spatial regions for cell or spot level data as well. Like in the 2014 *Platynereis dumereilii* atlas (Pettit et al. 2014), smfishHmrf (Zhu et al. 2018) also uses Potts model for dependence of label on neighborhood. As seqFISH data is quantitative, gene expression of each cell is modeled with a Gaussian mixture model, with as many components as there are there are region labels. The data needs to be normalized, although data normalization methods don't typically turn the distribution of gene expression Gaussian. Again, the parameters, i.e. the label assignment, and mean and covariance matrices for each Gaussian component, are estimated by EM, initiated with k-means clustering of the cells.

#### 7.8 Cell-cell interaction

Related to spatial regions is cell-cell interaction: Suppose a distinct neighborhood of the tissue has been identified with one of the methods in the previous section, and the neighborhood contains different cell types. Then it's natural to ask whether these cell types interact by their spatial proximity. Such information is lost in scRNA-seq. The composition of tissue neighborhoods can be characterized with existing tools. For instance, in smFISH or ISS data, we can count the number of cell types within a certain distance from each cell, as was done in the hypothalamus and the motor cortex MERFISH studies (Moffitt

[203] https://en.wikipedia.org/wiki/Louvain_method

et al. 2018; Zhang et al. 2020). We can model the data as a marked spatial point process, in which each point is a cell, with cell type annotations as marks, and use cross-type K or L function[204] to find cell types that colocalize; the cross-type L function[205] has been used in spicyR (Canete and Patrick 2020) for this purpose. For ST and Visium, we can use one of the cell type deconvolution methods to find the number and proportion of cell types per unit area in each tissue region and cell type colocalization. When two cell types colocalize, they might interact with secreted ligands or ligands and receptors bound to the membrane. Expression of ligand-receptor (L-R) pairs in neighboring cells is often used to identify cell-cell interaction in spatial data, and the CellPhoneDB (Efremova et al. 2020) database of ligands, receptors, and their interactions is often used to identify such L-R pairs. Another type of analysis going beyond colocalization tests for effects of cell-cell interaction or cell type colocalization on gene expression.

**Table 7.8:** Packages mentioned for cell-cell interactions

| Name | Language | Title | Date published |
|---|---|---|---|
| SVCA[206] | R; Python; C; C++; Fortran | Modeling Cell-Cell Interactions from Spatial Molecular Data with Spatial Variance Component Analysis[207] | 2019-10-01 |
| SpaOTsc[208] | Python | Inferring spatial and signaling relationships between cells from single cell transcriptomic data[209] | 2020-04-29 |
| MISTy[210] | R | Explainable multi-view framework for dissecting inter-cellular signaling from highly multiplexed spatial data[211] | 2020-05-10 |
| stLearn[212] | Python | stLearn: integrating spatial location, tissue morphology and gene expression to find cell types, cell-cell interactions and spatial trajectories within undissociated tissues[213] | 2020-05-31 |
| DIALOGUE[214] | R | Mapping multicellular programs from single-cell profiles[215] | 2020-08-11 |
| GCNG[216] | Python | GCNG: Graph convolutional networks for inferring cell-cell interactions[217] | 2020-12-10 |

[204]https://www.rdocumentation.org/packages/spatstat/versions/1.64-1/topics/Kest
[205]https://www.rdocumentation.org/packages/spatstat/versions/1.64-1/topics/Lcross

| | | | |
|---|---|---|---|
| Giotto[218] | R | Giotto, a pipeline for integrative analysis and visualization of single-cell spatial transcriptomic data[219] | 2021-03-08 |

##### 7.8.1 Ligand-receptor pairs

In stLearn (Pham et al. 2020), CellPhoneDB is used to identify L-R coexpression in neighboring spots, and the p-value of the coexpression is computed by permutation testing. Then regions with diverse cell types (from Seurat label transfering or cell type deconvolution) and L-R coexpression in neighboring spots are identified as regions where cells are likely to be signaling to each other. A similar strategy is used in Giotto. Giotto identifies cell type colocalization by labeling edges of the spatial neighborhood graph as homo- or heterotypic and permutes cell type labels to find whether the cell types are more or less likely to colocalize than expected from completely random cell type localization. L-R coexpression in neighboring cells on the spatial neighborhood graph from two cell types is identified and the p-values of the coexpression scores are computed by permutation testing, permuting locations of cells within each cell type.

While MRF, stLearn, and Giotto only use the immediate neighbors on the spatial neighborhood graph, there is a method that can capture higher order structures of the graph. In GCNG (Yuan and Bar-Joseph 2019), the spatial neighborhood graph is constructed as an edge connects a cell to its 3 nearest neighbors. Then both the gene count matrix and the normalized Laplacian of the neighborhood graph are fed into a graph convolutional neural network (GCN), which is trained on known L-R pairs. The GCN can then predict novel pairs of genes involved in signaling, and if trained on the direction of interaction in the L-R pairs, it can also predict the direction of causality in the novel pairs.

SpaOTsc has already been mentioned in Section 7.3. To recapitulate, SpaOTsc uses optimal transport from scRNA-seq cells to spatial locations to impute a spatial cell-cell distance matrix for scRNA-seq cells, and the optimal transport plan can be used to impute gene expression in space. With the cell-cell distance matrix, another optimal transport plan from ligands to receptors can be inferred, interpreted as how likely one cell communicates with another. A disadvantage of spatial neighborhood graph is that common ways of construction are somewhat arbitrary. For instance, $k$ nearest neighbor is a common way to construct the graph, but this $k$ is somewhat arbitrary, although cell signaling can occur over a distance with secreted ligands. Here no such graph is used; the length scale of interaction is inferred by random forest[220]. Random forest models are trained with expression of the ligand and genes correlated with a downstream target gene within a certain distance from the cells expressing the target gene are the input features. Receptor expression is the sample weights, and the target gene

[220] https://towardsdatascience.com/random-forest-3a55c3aca46d

is to be predicted by the random forest model. Several different length scales are tried, and the one resulting into the most feature importance of the ligand is used. When L-R information is unavailable, interactions between genes can be inferred by partial information decomposition, i.e. how much unique information can a source gene provide on a target gene in a spatial neighborhood.

With a very different model, DIALOGUE (Jerby-Arnon and Regev 2020) identifies genes that may be involved in interactions between cell types. In a niche in a tissue, different cell types can respond to the same environmental cue in a concerted manner though each cell type changes gene expression in a different way. DIALOGUE aims to identify such concerted gene programs in each cell type. First, the gene expression data is projected into a lower dimensional space in which correlation between all pairs of cell types across niches is maximized, and the basis of this space is ordered in descending strength of correlation. This is similar to CCA, but with a penalty term to enforce sparsity in gene loading. Here the niche is a patch of cells in space with a predefined number of cells. Then each cell type has a rotation matrix that projects cells into this lower dimensional space, and different cell types from the same niche should be close to each other in this space. In this projection, for each dimension, a gene is added to the multicellular program (MCP) of each cell type if its expression among cells of this cell type correlates with the projection of this cell type in this dimension and is significantly associated with the projection of other cell types while accounting for cell type level and niche level covariates such as sample, age, and gender. Thus the MCPs could be cell type specific co-regulated gene programs. Putative signaling between cell types can be identified by finding known L-R pairs in the MCPs: Each cell type is added the L-R graph as a node, and is connected to a gene if the gene is present in the MCP for this cell type. Then a path connecting one cell type to a ligand to a receptor and then to another cell type suggests signaling between the two cell types.

##### 7.8.2 Genes associated with cell-cell interaction

Gene expression can be affected by several different factors, including cell type, local environment, interaction with other cells, and so on. Some packages have been developed to identify genes whose expression is associated with one or more of these factors, without using L-R databases. Within one cell type, Giotto uses classical DE (Student's t-test, Wilcoxon rank sum test[221], limma[222], and permutation of spatial locations) to find DE genes between neighbors of cells of another cell type and non-neighbors. Other packages implement more complex models that account for more of these factors associated with gene expression.

Spatial variance component analysis (SVCA) (Arnol et al. 2019) models the expression of each gene of interest among the cells as a 0 mean Gaussian process. The covariance has the following components: First, the intrinsic variability,

[221] `https://en.wikipedia.org/wiki/Mann-Whitney_U_test`
[222] `https://bioconductor.org/packages/release/bioc/html/limma.html`

which can be cell types or continuous cell states. In the latter case, the covariance matrix of this component is the covariance between cells with genes other than the gene of interest that is modeled. Second, the spatial neighborhood. A neighborhood graph is not constructed, and the covariance matrix of this component is computed with the Gaussian kernel in which covariance decreases with distance between cells. Third, cell-cell interaction. The covariance matrix of this term is the covariance between cells weighed by a Gaussian kernel for distance between cells, so gene expression in nearby cells contributes more to this component. Finally, the residual has an identity covariance matrix, so in the residual, cells are independent from each other. The parameter to be estimated are weights of each of these components and the length scale parameter of the Gaussian kernel, which are estimated with MLE. Significance of the cell-cell interaction component is calculated by likelihood ratio test between the full model and a reduced model without the cell-cell interaction component. Again, as gene expression is modeled as Gaussian, the data needs to be normalized before using this method.

Like SVCA, Multiview Intercellular SpaTial modeling framework (MISTy) (Tanevski et al. 2020) also models expression of each gene of interest among the cells, but with ensemble learning, in which any machine learning method that is explainable (i.e. feature importance can be extracted) and suitable for ensemble learning[223] can be used. In each view, which can be intrinsic cell state, spatial neighborhood (juxtaview), or wider tissue structure (paraview), features are extracted from gene expression that represent the view and used in machine learning methods such as random forest to predict expression of a gene of interest. For intrinsic cell state, the features are expression of other genes. For juxtaview, the features are sum of expression of other genes in neighboring cells in the spatial neighborhood graph. For paraview, the feature are sum of expression of other genes in all cells in the tissue weighed by distance to each cell with a Gaussian kernel. Other views, with other feature engineering, can also be used. The full model is a linear combination of predictions of each view. In other words, the contribution of each view is determined by linear regression with prediction of each view as a covariate to predict the expression of the gene of interest. For each view, the importance of each feature is assessed as the z-score of the feature importance (e.g. from random forest) multiplied by 1 minus the p-value of the coefficient of this view in the linear regression model, so views that contribute significantly to the ensemble model and features in each of these views that are more important than other features in the same views stand out. This way, interaction among genes at different spatial scales can be identified.

#### 7.9 Gene-gene interaction

[223] `https://www.analyticsvidhya.com/blog/2015/08/introduction-ensemble-learning/`

**Table 7.9:** Packages mentioned for gene-gene interactions

| Name | Language | Title | Date published |
|---|---|---|---|
| SpaOTsc[224] | Python | Inferring spatial and signaling relationships between cells from single cell transcriptomic data[225] | 2020-04-29 |
| MISTy[226] | R | Explainable multi-view framework for dissecting inter-cellular signaling from highly multiplexed spatial data[227] | 2020-05-10 |
| scHOT[228] | R | Investigating higher-order interactions in single-cell data with scHOT[229] | 2020-07-13 |
| MESSI[230] | Python | Identifying signaling genes in spatial single cell expression data[231] | 2020-09-04 |
| GCNG[232] | Python | GCNG: Graph convolutional networks for inferring cell-cell interactions[233] | 2020-12-10 |

Some of the packages already reviewed can also infer interactions between genes, such as GCNG, SpaOTsc, and MISTy. GCNG and SpaOTsc predict potential L-R pairs, and MISTy identify genes whose expression at a given spatial scale is associated with another gene of interest. The package scHOT (Ghazanfar et al. 2020) tests for association of correlation between genes with pseudotime or spatial locations by permutation testing, permuting locations of cells along pseudotime or in space. The package Mixture of Experts for Spatial Signaling genes Identification (MESSI) (Li et al. 2020) uses a mixture of experts model to predict expression of response genes with a set of features. A spatial neighborhood graph of the cells is constructed with Delaunay triangulation The features include all genes quantified in the dataset that are also found in a L-R database, expression of genes in the L-R database in neighboring cells, cell type of neighboring cells, and etc. The response genes are all genes quantified other than the L-R genes used as features. Each cell is assigned to exactly one "expert", i.e. subtype. For each expert, expression of response genes in each cell is modeled with linear regression with the features as covariates. The parameters of the linear models and assignment of cells to experts are estimated with MLE, where the log likelihood is maximized with EM. This model can be trained in a control sample and used to predict gene expression in experimental samples. If expression of a gene is as well predicted as in the control, then signaling may not have changed in the experimental condition. If prediction becomes worse, then there may be a change in signaling involving this gene, and the experts whose coefficients significantly differ between the control and experimental models suggest cell populations involved in the signaling change.

#### 7.10 Subcellular transcript localization

**Table 7.10:** Packages mentioned for subcellular transcript localization

| Name | Language | Title | Date published |
|---|---|---|---|
| FISH_quant[234] | MATLAB | A computational framework to study sub-cellular RNA localization[235] | 2018-11-02 |

So far, except for segmentation free data analysis methods of smFISH and ISS images, all analysis methods are at the cellular or spot level. However, transcripts do show inhomogeneous subcellular localization that can be biologically relevant, such as whether the transcripts are translated in the endoplasmic reticulum (ER) or the cytoplasm. Thirty four lncRNAs have been manually classified into 5 types of subcellular patterns: one or two large foci in nucleus, large foci and dispersed single molecules in nucleus, no foci in nucleus, nucleus and cytoplasm, and cytoplasmic (Cabili et al. 2015). The bDNA-smFISH study in 2013 that profiled 928 genes in cultured cells, though each gene was profiled in different cells, generated features that characterize subcellular transcript localization (mRNAs of protein coding genes) which were used to cluster cells (Battich, Stoeger, and Pelkmans 2013). These features include closest distance of a transcript spot to cell outline, distance to cell centroid, distance to nuclear centroid, radius to include 5%, 10%, 15%, 25%, 50%, and 75% of all spots in the cell, mean distance of a spot to other spots (related to Ripley's K or L function), and variance of distance to other spots. The package FISHquant (Samacoits et al. 2018) uses additional features derived from Ripley's L function of subcellular transcript localization. Then it uses these features to simulate smFISH data, cluster cells, and classify transcript localization patterns.

Whether transcripts are located in the nucleus or in the cytoplasm has also been used for RNA velocity in a MERFISH study (C. Xia, Babcock, et al. 2019). In traditional RNA velocity based on scRNA-seq (La Manno et al. 2018), when there are more transcripts with intronic reads — i.e. nascent transcripts not yet spliced — than expected from steady state in a cell, then the gene of interest is up regulated, and conversely, if there are fewer transcripts with intronic reads, then the gene may be down regulated. In other words, intronic reads not yet spliced out gives a glimpse into a near future transcriptome of the cell. In this MERFISH study, instead of introns that require separate probes from exons, transcripts inside the nucleus are taken to be nascent, i.e. not yet exported from the nucleus, and used in lieu of intronic reads as in scRNA-seq for RNA velocity. In this study, the ER was also stained for and segmented and genes with transcripts enriched in the ER were also identified.

These studies analyzing subcellular transcript localization were all performed on cultured cells rather than in tissues, so there is no highly multiplexed smFISH data on in vivo subcellular transcript localization yet. Furthermore, as the cells cultures used in these studies grow on a plate in single layer, while cells stack on top of each other through the thickness of the section, cell segmentation in tissue can be more challenging. Some of the features used to characterize subcellular transcript localization, such as distance to cell outline and Ripley's L function (with edge correction), depend on accurate cell segmentation, which as already explained in Section 7.1, is challenging. Subcellular transcript location can be modeled as a spatial point pattern in 3D or collapsed into 2D, and analyses such as finding effects of covariates such as whether the spot is in the nucleus, distance from the nucleus, distance from the cell outline, and etc., and whether the pattern exhibits clustering (e.g. foci) or inhibition (i.e. more spaced out than expected from CSR[236]). However, the observational window of the point process, i.e. cell segmentation, can greatly affect results of spatial point pattern analysis. For instance, when the convex hull of some spots are taken to be the observational window, then the point pattern may not appear clustered. However, if the actual observational window is much larger than the convex hull, then the point process is in fact clustered. Hence accurate cell segmentation is important to analyses of subcellular transcript localization patterns. Furthermore, some smFISH or ISS datasets only provide 2D cell segmentations, and resolution in the z axis tends to be lower than that in the x and y axes. The implications of collapsing 3D into 2D, and when 3D segmentation is available, the lower resolution in the z axis are yet to be determined.

#### 7.11 Gene expression imputation from H&E

**Table 7.11:** Packages mentioned for gene expression imputation from H&E

| Name | Language | Title | Date published |
|---|---|---|---|
| Xfuse[237] | Python | Super-resolved spatial transcriptomics by deep data fusion[238] | 2020-03-13 |
| ST-Net[239] | Python | Integrating Spatial Gene Expression and Breast Tumour Morphology via Deep Learning[240] | 2020-06-22 |
| PathoMCH[241] | Python | Spatial transcriptomics inferred from pathology whole-slide images links tumor heterogeneity to survival in breast and lung cancer[242] | 2020-11-02 |

[236] https://en.wikipedia.org/wiki/Complete_spatial_randomness

Although ST and Visium do not have single cell resolution, the tissue sections can be H&E stained prior to library preparation, thus the transcriptomes of the spots can be mapped to H&E tissue morphology. H&E is also commonly used in clinical pathology, while ST and Visium are not used for diagnostic purposes. The package ST-Net was developed to use a pretrained CNN to extract features from H&E images behind the ST spots, and a dense neural net is trained on the extracted features and ST data to predict gene expression based on H&E images from held out patients as log normalized UMI counts (He et al. 2020). Another method to predict gene expression from H&E is PathoMCH (Levy-Jurgenson et al. 2020). TGCA transcriptomics data is normalized and the corresponding H&E slides are labeled with the percentile of expression of each gene of interest. Then the whole slide images are broken into small tiles, all of which take the percentile label of the slide. Then the Inception v3 classification neural network is trained with the tiles and labels with very high or very low expression, and when predicting on held out images, it gives a score of gene expression from low to high in each tile. Such gene expression prediction methods can give pathologists a more nuanced view of the tissue beyond morphology.

While cell segmentation is difficult in H&E images, H&E images do have enough resolution to exhibit subcellular details. The H&E image is used in Xfuse to increase resolution of the spatial transcriptome from ST (Bergenstråhle et al. 2020). The H&E image and the corresponding transcriptomes are modeled to come from a shared latent space. Intensity of each channel at each pixel is modeled as Gaussian, and gene expression at each pixel is modeled as NB so the observed value at each spot are the sums of values at the pixels in the spot. Parameters of these distributions are mapped from the latent space through a generator CNN. The parameters are estimated with variational Bayesian inference. With the parameters, gene expression at each pixel can be predicted, thus increasing the resolution of ST.

### Part III

### Future perspectives

### Chapter 8

### From the past to the present to the future

The quest to profile the transcriptome in space with high resolution is not new. It started with the enhancer and gene trap screens in the late 1980s and the 1990s, before the genomes of metazoans were sequenced. However, in the prequel era, challenges with the existing technology made the dream of profiling the transcriptome in space hard to reach, as the technologies were not highly-multiplexed and not very quantitative. Over 30 years later, this dream seems to be more within reach, though with some caveats. We have come so far, because of so many strands of ideas and technologies coming together since the late 2010s. Highly multiplexed smFISH that can profile 10000 genes at a time would not have been possible without the reference genome sequence to screen for off target binding, the reference transcriptome and genome annotation with which to design the probes, the technology to synthesize DNA oligos, smFISH, confocal microscopy, digital photography, combinatorial barcoding, and the computing power to store and process terabytes of images. ST and Visium would not have been possible without microarray technology, scRNA-seq techniques designed for small amount of RNA from each spot, NGS, and the computing power to process the data. Some of these strands are older than others, and each of them would not have been possible without more preceding strands coming together. For instance, smFISH would not have been possible without the development of non-radioactive FISH in the late 1970s and the 1980s and techniques to synthesize fluorophore labeled probes. The field of spatial transcriptomics has grown tremendously since the late 2010s, as this is the time when a wide array of technologies truly started to add up to more than the sum of their parts.

Spatial transcriptomics still faces many challenges. First, there still is the trade off between quantity and quality. ST and Visium, which have limited resolution and low detection efficiency, can be more easily applied to larger areas of tissue and the whole transcriptome. ISS has been applied to whole mouse brain sections, because while it has lower detection efficiency than smFISH, the amplified and less crowded signals can be detected at lower magnification. In contrast, while smFISH based techniques have subcellular resolution and often over 80% detection efficiency, the efficiency is compromised when applied to 10000 genes and these techniques are more difficult to apply to larger areas of tissue. As there are still challenges, new techniques to collect data are constantly being developed. Second, compared to the prequel era, the current era is more elitist. While commercial LCM, ST, and Visium have spread far and wide, the various high quality smFISH based techniques mostly failed to spread beyond their usually elite institutions of origin. This might be due to difficulty in building custom flow cells, challenges in customizing the protocols to different tissues, limits in number of genes and cells profiled, lack of core facilities for these techniques, and lack of unified, efficient, open source, and well documented software platform to process the data.

Data analysis has also come a long way, from PCA and ICA in the early 2000s to much more sophisticated techniques today. Many ideas that originated in other fields such as computer vision, machine learning, and statistics, including geospatial statistics, have been adapted to spatial transcriptomics in recent years. Ideas from computer vision include SIFT, NMF, CNN, and to some extent also PCA and ICA. Ideas from machine learning include SVM, neural networks, bag of words, variational autoencoders (for some cases of latent space), mixture of experts model, $k$ nearest neighbor, and clustering. Ideas from statistics include CCA, permutation testing, MCMC, factor analysis, generalized linear models, and hierarchical modeling. Ideas from geospatial statistics include Gaussian process model (usually used for kriging), spatial point process, and MRF. Other ideas include Laplacian score and optimal transport. Conceivably, more ideas can be adapted to spatial transcriptomics. For instance, spatiotemporal statistics can be adapted to analyze multiple aligned sections of the same tissue to address the difference in covariance between the z axis and the x and y axes. Well established methods in geospatial statistics, such as the semivariogram, J function, G function, and other point process models are also primising for spatial transcriptomics.

We have reviewed many different types of data analysis, using a diverse arsenal of principles. However, integrated analysis pipelines like Seurat are still immature for spatial transcriptomics; Seurat only supports the most rudimentary analyses and the user still needs to learn different syntax and convert data to different formats to use many of the other more specialized and advanced tools, many of which are not well documented. However, the open source culture is flourishing and growing. Most prequel data analysis publications did not link to a repository of the implementation of the software, while most current era data analysis publications do. While the proprietary MATLAB language is still in use, most, especially more recent, current era publication use R, Python, C++, and in some cases Julia and Rust, which are open source and free. Open source software and freely available data may enable less privileged individuals and institutions to perform data analysis and develop new data analysis tools.

What would an ideal future of spatial transcriptomics look like? Data collection would have subcellular resolution, be transcriptome wide, have nearly 100% detection efficiency, and is scalable to large areas of tissues in 3D. Even better, it's multiomic, profiling not only transcriptome, but also epigenome, proteome, metabolome, and etc., with equally high quality and throughput for the other omics. Moreover, the data collection technique is easy to use, such as coming in easy to use kits, and affordable, so it can spread far and wide into non-elite institutions. It should also be open source and transparent, so it would be easier for others to improve it. While we have reviewed many data analysis methods, a comprehensive benchmark of the methods for each analysis task and evaluation of user experience, like in dynverse for scRNA-seq pseudotime analysis (Saelens et al. 2019), would be helpful for users to choose a method to use and for developers to compare their new methods to existing methods.

Data analysis would have the same user-friendly user interface for different data types and different methods for the same task. Also, the package should be modular, so dependencies are only installed if needed. It should also be extensible, so users can add additional modules or additional tools for existing tasks to the integrative framework. This would be like SeuratWrappers, which provides Seurat interfaces to data integration and RNA velocity methods not implemented by Seurat. Or like caret and tidymodels, which provide a uniform user interface to numerous machine learning methods. This can be achieved with guidelines such as those used by Bioconductor, encouraging developers to reuse existing data structures and methods in Bioconductor rather than reinventing the wheel. It should also be effective at its task, scalable, well documented, open source, unit tested, easy to install, and portable, again, as enforced to some extent by the Bioconductor guideline. It should be implemented in easy to read code, so developers can more easily fix bugs and improve the package. In addition, it should be interoperable, so tools written in different programming languages can be integrated, combining their strengths and bridging cultural differences between the programming language communities. It should have elegant data visualization, both static for publications and interactive for data exploration and sharing. The data visualization should also be accessible, such as using redundant encoding and colorblind friendly palettes and providing alternatives to those who are visual impaired. Finally, it should be integrated with a graphical user interface (GUI) like iSee so the data can be shared with colleagues who do not code.

### References

"10X VISIUM SPATIAL TRANSCRIPTOMICS." n.d. Accessed October 7, 2020. https://biotech.illinois.edu/htdna/applications/10x-visium-spatial-transcriptomics.

Abdelaal, Tamim, Soufiane Mourragui, Ahmed Mahfouz, and Marcel J T Reinders. 2020. "SpaGE: Spatial Gene Enhancement using scRNA-seq." *Nucleic Acids Research* 48 (18): e107–e107. https://doi.org/10.1093/nar/gkaa740.

Abed-Esfahani, Pegah, Benjamin C Darwin, Derek Howard, Nick Wang, Ethan Kim, Jason Lerch, and Leon French. 2021. "Evaluation of deep convolutional neural networks for in situ hybridization gene expression image representation." *bioRxiv*, January, 2021.01.22.427860. https://doi.org/10.1101/2021.01.22.427860.

Aboobaker, A. Aziz, Pavel Tomancak, Nipam Patel, Gerald M. Rubin, and Eric C. Lai. 2005. "Drosophila microRNAs exhibit diverse spatial expression patterns during embryonic development." *Proceedings of the National Academy of Sciences* 102 (50): 18017–22. https://doi.org/10.1073/pnas.0508823102.

Achim, Kaia, Jean Baptiste Pettit, Luis R. Saraiva, Daria Gavriouchkina, Tomas Larsson, Detlev Arendt, and John C. Marioni. 2015. "High-throughput spatial mapping of single-cell RNA-seq data to tissue of origin." *Nature Biotechnology* 33 (5): 503–9. https://doi.org/10.1038/nbt.3209.

Adkins, Ricky S, Andrew I Aldridge, Shona Allen, Seth A Ament, Xu An, Ethan Armand, Giorgio A Ascoli, et al. 2020. "A multimodal cell census and atlas of the mammalian primary motor cortex." *bioRxiv*, January, 2020.10.19.343129. https://doi.org/10.1101/2020.10.19.343129.

"ADVANCED GENOMICS CORE PRICING." n.d. Accessed October 7, 2020. https://brcf.medicine.umich.edu/cores/advanced-genomics/price-list/.

Agapite, Julie, Laurent-Philippe Albou, Suzi Aleksander, Joanna Argasinska, Valerio Arnaboldi, Helen Attrill, Susan M. Bello, et al. 2020. "Alliance of Genome Resources Portal: unified model organism research platform." *Nucleic Acids Research* 48 (D1): D650–D658. https://doi.org/10.1093/nar/gkz813.

Aguila, Julio, Shangli Cheng, Nigel Kee, Ming Cao, Qiaolin Deng, and Eva Hedlund. 2018. "Spatial transcriptomics and in silico random pooling identify novel markers of vulnerable and resistant midbrain dopamine neurons." *bioRxiv*, October, 334417. `https://doi.org/10.1101/334417`.

Ahmed, Ayisha, Nicole J Ward, Simon Moxon, Sara Lopez-Gomollon, Camille Viaut, Matthew L Tomlinson, Ilya Patrushev, et al. 2015. "A Database of microRNA Expression Patterns in Xenopus laevis." *PLOS ONE* 10 (10): e0138313. `https://doi.org/10.1371/journal.pone.0138313`.

Allen, Nicholas D., David G. Cran, Sheila C. Barton, Simon Hettle, Wolf Reik, and M. Azim Surani. 1988. "Transgenes as probes for active chromosomal domains in mouse development." *Nature* 333 (6176): 852–55. `https://doi.org/10.1038/333852a0`.

Alon, Shahar, Daniel R Goodwin, Anubhav Sinha, Asmamaw T Wassie, Fei Chen, Evan R Daugharthy, Yosuke Bando, et al. 2020. "Expansion Sequencing: Spatially Precise <em>In Situ</em> Transcriptomics in Intact Biological Systems." *bioRxiv*, January, 2020.05.13.094268. `https://doi.org/10.1101/2020.05.13.094268`.

Alonso, A M, A Carrea, and L Diambra. 2020. "Prediction of cell position using single-cell transcriptomic data: an iterative procedure [version 2; peer review: 2 approved]." *F1000Research* 8 (1775). `https://doi.org/10.12688/f1000research.20715.2`.

Alsup, William H. 2009. "Applera v. Illumina." `https://www.leagle.com/decision/infco20100325236`.

"A Mouse for All Reasons." 2007. *Cell* 128 (1): 9–13. `https://doi.org/10.1016/j.cell.2006.12.018`.

An, Li, Hongbo Xie, Mark H Chin, Zoran Obradovic, Desmond J Smith, and Vasileios Megalooikonomou. 2009. "Analysis of multiplex gene expression maps obtained by voxelation." *BMC Bioinformatics* 10 (S4): S10. `https://doi.org/10.1186/1471-2105-10-S4-S10`.

Andersson, Alma, Joseph Bergenstråhle, Michaela Asp, Ludvig Bergenstråhle, Aleksandra Jurek, José Fernández Navarro, and Joakim Lundeberg. 2019. "Spatial mapping of cell types by integration of transcriptomics data." *bioRxiv*, December, 2019.12.13.874495. `https://doi.org/10.1101/2019.12.13.874495`.

Andersson, Axel, Ferran Diego, Fred A Hamprecht, and Carolina Wählby. 2021. "ISTDECO: In Situ Transcriptomics Decoding by Deconvolution." *bioRxiv*, January, 2021.03.01.433040. `https://doi.org/10.1101/2021.03.01.433040`.

Andonian, Alexander, Daniel Paseltiner, Travis J. Gould, and Jason B. Castro. 2019. "A deep learning based method for large-scale classification, registration, and clustering of in-situ hybridization experiments in the mouse olfactory bulb." *Journal of Neuroscience Methods* 312 (January): 162–68. `https://doi.org/10.1016/j.jneumeth.2018.12.003`.

"Arcturus XT Laser Capture Microdissection (LCM) Instrument - US." n.d. //www.thermofisher.com/us/en/home/life-science/gene-expression-analysis-genotyping/laser-capture-microdissection/arcturus-laser-capture-microdissection-lcm-instrument.html.

Ardini-Poleske, Maryanne E., Robert F. Clark, Charles Ansong, James P. Carson, Richard A. Corley, Gail H. Deutsch, James S. Hagood, et al. 2017. "LungMAP: The Molecular Atlas of Lung Development Program." *American Journal of Physiology-Lung Cellular and Molecular Physiology* 313 (5): L733–L740. https://doi.org/10.1152/ajplung.00139.2017.

Arnol, Damien, Denis Schapiro, Bernd Bodenmiller, Julio Saez-Rodriguez, and Oliver Stegle. 2019. "Modeling Cell-Cell Interactions from Spatial Molecular Data with Spatial Variance Component Analysis." *Cell Reports* 29 (1): 202–211.e6. https://doi.org/10.1016/j.celrep.2019.08.077.

Asp, Michaela, Stefania Giacomello, Ludvig Larsson, Chenglin Wu, Daniel Fürth, Xiaoyan Qian, Eva Wärdell, et al. 2019. "A Spatiotemporal Organ-Wide Gene Expression and Cell Atlas of the Developing Human Heart." *Cell* 179 (7): 1647–1660.e19. https://doi.org/10.1016/j.cell.2019.11.025.

Asp, Michaela, Fredrik Salmén, Patrik L Ståhl, Sanja Vickovic, Ulrika Felldin, Marie Löfling, José Fernandez Navarro, et al. 2017. "Spatial detection of fetal marker genes expressed at low level in adult human heart tissue." *Scientific Reports* 7 (1): 12941. https://doi.org/10.1038/s41598-017-13462-5.

Axelrod, Shannon, Ambrose J Carr, Jeremy Freeman, Deep Ganguli, Brian Long, Tony Tung, and Others. n.d. "{Starfish}: Open Source Image Based Transcriptomics and Proteomics Tools." http://github.com/spacetx/starfish.

Baccin, Chiara, Jude Al-Sabah, Lars Velten, Patrick M. Helbling, Florian Grünschläger, Pablo Hernández-Malmierca, César Nombela-Arrieta, Lars M. Steinmetz, Andreas Trumpp, and Simon Haas. 2020. "Combined single-cell and spatial transcriptomics reveal the molecular, cellular and spatial bone marrow niche organization." *Nature Cell Biology* 22 (1): 38–48. https://doi.org/10.1038/s41556-019-0439-6.

Baharlou, Heeva, Nicolas P. Canete, Anthony L. Cunningham, Andrew N. Harman, and Ellis Patrick. 2019. "Mass Cytometry Imaging for the Study of Human Diseases—Applications and Data Analysis Strategies." *Frontiers in Immunology* 10 (November): 2657. https://doi.org/10.3389/fimmu.2019.02657.

Bakken, Trygve E., Jeremy A. Miller, Song Lin Ding, Susan M. Sunkin, Kimberly A. Smith, Lydia Ng, Aaron Szafer, et al. 2016. "A comprehensive transcriptional map of primate brain development." *Nature* 535 (7612): 367–75. https://doi.org/10.1038/nature18637.

Baldock, Richard, Jonathan Bard, Matt Kaufman, and Duncan Davidson. 1992.

"What's New? A real mouse for your computer." *BioEssays* 14 (7): 501–2. https://doi.org/10.1002/bies.950140713.

Baner, J., Mats Nilsson, Maritha Mendel-Hartvig, and Uif Landegren. 1998. "Signal amplification of padlock probes by rolling circle replication." *Nucleic Acids Research* 26 (22): 5073–8. https://doi.org/10.1093/nar/26.22.5073.

Bassell, G J, C M Powers, K L Taneja, and R H Singer. 1994. "Single mRNAs visualized by ultrastructural in situ hybridization are principally localized at actin filament intersections in fibroblasts." *The Journal of Cell Biology* 126 (4): 863–76. https://doi.org/10.1083/jcb.126.4.863.

Bates, Mark, Graham T. Dempsey, Kok Hao Chen, and Xiaowei Zhuang. 2012. "Multicolor Super-Resolution Fluorescence Imaging via Multi-Parameter Fluorophore Detection." *ChemPhysChem* 13 (1): 99–107. https://doi.org/10.1002/cphc.201100735.

Battich, Nico, Thomas Stoeger, and Lucas Pelkmans. 2013. "Image-based transcriptomics in thousands of single human cells at single-molecule resolution." *Nature Methods* 10 (11): 1127–36. https://doi.org/10.1038/nmeth.2657.

Bayraktar, Omer Ali, Theresa Bartels, Staffan Holmqvist, Vitalii Kleshchevnikov, Araks Martirosyan, Damon Polioudakis, Lucile Ben Haim, et al. 2020. "Astrocyte layers in the mammalian cerebral cortex revealed by a single-cell in situ transcriptomic map." *Nature Neuroscience* 23 (4): 500–509. https://doi.org/10.1038/s41593-020-0602-1.

Becker, Ingrid, Karl Friedrich Becker, Michael H. Röhrl, Günter Minkus, Karin Schütze, and Heinz Höfler. 1996. "Single-cell mutation analysis of tumors from stained histologic slides." *Laboratory Investigation* 75 (6): 801–7. https://pubmed.ncbi.nlm.nih.gov/8973475/.

Bell, George W., Tatiana A. Yatskievych, and Parker B. Antin. 2004. "GEISHA, a whole-mount in situ hybridization gene expression screen in chicken embryos." *Developmental Dynamics* 229 (3): 677–87. https://doi.org/10.1002/dvdy.10503.

Bellen, H. J., C. J. O'Kane, C. Wilson, U. Grossniklaus, R. K. Pearson, and W. J. Gehring. 1989. "P-element-mediated enhancer detection: a versatile method to study development in Drosophila." *Genes & Development* 3 (9): 1288–1300. https://doi.org/10.1101/gad.3.9.1288.

Belmamoune, Mounia, and Fons J. Verbeek. 2008. "Data Integration for Spatio-Temporal Patterns of Gene Expression of Zebrafish development: the GEMS database." *Journal of Integrative Bioinformatics* 5 (2): 35–48. https://doi.org/10.1515/jib-2008-92.

Belyavsky, A, T Vinogradova, and K Rajewsky. 1989. "PCR-based cDNA library construction: general cDNA libraries at the level of a few cells." *Nucleic Acids Research* 17 (8): 2919–32. https://doi.org/10.1093/nar/17.8.2919.

Bergenstråhle, Joseph, Ludvig Bergenstråhle, and Joakim Lundeberg. 2020. "SpatialCPie: an R/Bioconductor package for spatial transcriptomics cluster evaluation." *BMC Bioinformatics* 21 (1): 161. https://doi.org/10.1186/s12859-020-3489-7.

Bergenstråhle, Ludvig, Bryan He, Joseph Bergenstråhle, Alma Andersson, Joakim Lundeberg, James Zou, and Jonas Maaskola. 2020. "Super-resolved spatial transcriptomics by deep data fusion." *bioRxiv*, March, 2020.02.28.963413. https://doi.org/10.1101/2020.02.28.963413.

Berglund, Emelie, Jonas Maaskola, Niklas Schultz, Stefanie Friedrich, Maja Marklund, Joseph Bergenstråhle, Firas Tarish, et al. 2018. "Spatial maps of prostate cancer transcriptomes reveal an unexplored landscape of heterogeneity." *Nature Communications*. https://doi.org/10.1038/s41467-018-04724-5.

Berns, Michael W., Roberts S. Olson, and Donald E. Rounds. 1969. "In vitro Production of Chromosomal Lesions with an Argon Laser Microbeam." *Nature* 221 (5175): 74–75. https://doi.org/10.1038/221074a0.

Bettenhausen, Berthold, and Achim Gossler. 1995. "Efficient Isolation of Novel Mouse Genes Differentially Expressed in Early Postimplantation Embryos." *Genomics* 28 (3): 436–41. https://doi.org/10.1006/geno.1995.1172.

Biancalani, Tommaso, Gabriele Scalia, Lorenzo Buffoni, Raghav Avasthi, Ziqing Lu, Aman Sanger, Neriman Tokcan, et al. 2020. "Deep learning and alignment of spatially-resolved whole transcriptomes of single cells in the mouse brain with Tangram." *bioRxiv*, January, 2020.08.29.272831. https://doi.org/10.1101/2020.08.29.272831.

Bidarimath, Mallikarjun, Andrew K. Edwards, and Chandrakant Tayade. 2015. "Laser Capture Microdissection for Gene Expression Analysis." In *Methods in Molecular Biology*, 1219:115–37. Humana Press Inc. https://doi.org/10.1007/978-1-4939-1661-0_10.

Bier, E., H. Vaessin, S. Shepherd, K. Lee, K. McCall, S. Barbel, L. Ackerman, R. Carretto, T. Uemura, and E. Grell. 1989. "Searching for pattern and mutation in the Drosophila genome with a P-lacZ vector." *Genes & Development* 3 (9): 1273–87. https://doi.org/10.1101/gad.3.9.1273.

BinTayyash, Nuha, Sokratia Georgaka, S T John, Sumon Ahmed, Alexis Boukouvalas, James Hensman, and Magnus Rattray. 2020. "Non-parametric modelling of temporal and spatial counts data from RNA-seq experiments." *bioRxiv*, January, 2020.07.29.227207. https://doi.org/10.1101/2020.07.29.227207.

Birchall, Philip S, Rita M Fishpool, and Donna G Albertson. 1995. "Expression patterns of predicted genes from the C. elegans genome sequence visualized by FISH in whole organisms," 314–20. https://doi.org/10.1038/ng1195-314.

Blackshaw, Seth, Sanjiv Harpavat, Jeff Trimarchi, Li Cai, Haiyan Huang, Winston P Kuo, Griffin Weber, et al. 2004. "Genomic Analysis of Mouse Retinal Development." Edited by William Harris. *PLoS Biology* 2 (9): e247. https://doi.org/10.1371/journal.pbio.0020247.

Blitz, Ira L., Kitt D. Paraiso, Ilya Patrushev, William T. Y. Chiu, Ken W. Y. Cho, and Michael J. Gilchrist. 2017. "A catalog of Xenopus tropicalis transcription factors and their regional expression in the early gastrula stage embryo." *Developmental Biology* 426 (2): 409–17. https://doi.org/10.1016/j.ydbio.2016.07.002.

Boer, B. A. de, Jan M. Ruijter, F. P. J. M. Voorbraak, and A. F. M. Moorman. 2009. "More than a decade of developmental gene expression atlases: where are we now?" *Nucleic Acids Research* 37 (22): 7349–59. https://doi.org/10.1093/nar/gkp819.

Bohland, Jason W., Hemant Bokil, Sayan D. Pathak, Chang-Kyu Lee, Lydia Ng, Christopher Lau, Chihchau Kuan, Michael Hawrylycz, and Partha P. Mitra. 2010. "Clustering of spatial gene expression patterns in the mouse brain and comparison with classical neuroanatomy." *Methods* 50 (2): 105–12. https://doi.org/10.1016/j.ymeth.2009.09.001.

Boulgakov, Alexander A, Andrew D Ellington, and Edward M Marcotte. 2020. "Bringing Microscopy-By-Sequencing into View." *Trends in Biotechnology* 38 (2): 154–62. https://doi.org/https://doi.org/10.1016/j.tibtech.2019.06.001.

Bowes, Jeff B., Kevin A. Snyder, Erik Segerdell, Chris J. Jarabek, Kenan Azam, Aaron M. Zorn, and Peter D. Vize. 2009. "Xenbase: Gene expression and improved integration." *Nucleic Acids Research* 38 (SUPPL.1): D607–D612. https://doi.org/10.1093/nar/gkp953.

Brady, Gerard, Mary Barbara, and Norman N. Iscove. 1990. "Representative in Vitro cDNA Amplification From Individual Hemopoietic Cells and Colonies." *Methods in Molecular and Cellular Biology* 2: 17–25.

Brady, Lauren, Michelle Kriner, Ilsa Coleman, Colm Morrissey, Martine Roudier, Lawrence D True, Roman Gulati, et al. 2021. "Inter- and intra-tumor heterogeneity of metastatic prostate cancer determined by digital spatial gene expression profiling." *Nature Communications* 12 (1): 1426. https://doi.org/10.1038/s41467-021-21615-4.

Bravo González-Blas, Carmen, Xiao-Jiang Quan, Ramon Duran-Romaña, Ibrahim Ihsan Taskiran, Duygu Koldere, Kristofer Davie, Valerie Christiaens, et al. 2020. "Identification of genomic enhancers through spatial integration of single-cell transcriptomics and epigenomics." *Molecular Systems Biology* 16 (5): e9438. https://doi.org/10.15252/msb.20209438.

Brink, Susanne C. van den, Anna Alemany, Vincent van Batenburg, Naomi Moris, Marloes Blotenburg, Judith Vivié, Peter Baillie-Johnson, et al. 2020. "Single-cell and spatial transcriptomics reveal somitogenesis in gastruloids." *Nature* 582 (7812): 405. https://doi.org/10.1038/s41586-020-2024-3.

Brown, Vanessa M. 2002. "High-Throughput Imaging of Brain Gene Expression." *Genome Research* 12 (2): 244–54. https://doi.org/10.1101/gr.204102.

Brown, V. M., A. Ossadtchi, A. H. Khan, S. Yee, G. Lacan, W. P. Melega, S. R. Cherry, R. M. Leahy, and D. J. Smith. 2002. "Multiplex Three-Dimensional Brain Gene Expression Mapping in a Mouse Model of Parkinson's Disease." *Genome Research* 12 (6): 868–84. https://doi.org/10.1101/gr.229002.

Buchberger, Amanda Rae, Kellen DeLaney, Jillian Johnson, and Lingjun Li. 2018. "Mass Spectrometry Imaging: A Review of Emerging Advancements and Future Insights." *Analytical Chemistry* 90 (1): 240–65. https://doi.org/10.1021/acs.analchem.7b04733.

Burgess, Darren J. 2019. "Spatial transcriptomics coming of age." *Nature Reviews Genetics* 20 (6): 317–17. https://doi.org/10.1038/s41576-019-0129-z.

Burkhard, Silja Barbara, and Jeroen Bakkers. 2018. "Spatially resolved RNA-sequencing of the embryonic heart identifies a role for Wnt/$\beta$-catenin signaling in autonomic control of heart rate." *eLife* 7 (February). https://doi.org/10.7554/eLife.31515.

Bustin, Stephen, Harvinder S Dhillon, Sara Kirvell, Christina Greenwood, Michael Parker, Gregory L Shipley, and Tania Nolan. 2015. "Variability of the Reverse Transcription Step: Practical Implications." *Clinical Chemistry* 61 (1): 202–12. https://doi.org/10.1373/clinchem.2014.230615.

Butler, Daniel, Christopher Mozsary, Cem Meydan, Jonathan Foox, Joel Rosiene, Alon Shaiber, David Danko, et al. 2021. "Shotgun transcriptome, spatial omics, and isothermal profiling of SARS-CoV-2 infection reveals unique host responses, viral diversification, and drug interactions." *Nature Communications* 12 (1): 1660. https://doi.org/10.1038/s41467-021-21361-7.

Cabili, Moran N., Margaret C. Dunagin, Patrick D. McClanahan, Andrew Biaesch, Olivia Padovan-Merhar, Aviv Regev, John L. Rinn, and Arjun Raj. 2015. "Localization and abundance analysis of human lncRNAs at single-cell and single-molecule resolution." *Genome Biology* 16 (1): 20. https://doi.org/10.1186/s13059-015-0586-4.

Cable, Dylan M., Evan Murray, Luli S. Zou, Aleksandrina Goeva, Evan Z. Macosko, Fei Chen, and Rafael A. Irizarry. 2020. "Robust decomposition of cell type mixtures in spatial transcriptomics." *bioRxiv*, May, 2020.05.07.082750. https://doi.org/10.1101/2020.05.07.082750.

Campiteli, Monica Guimarães, Cesar Henrique Comin, Luciano Da Fontoura Costa, M. Madan Babu, and Roberto Marcondes Cesar. 2013. "A methodology to infer gene networks from spatial patterns of expression – an application to fluorescence in situ hybridization images." *Molecular BioSystems* 9 (7): 1926. https://doi.org/10.1039/c3mb25475e.

Canete, Nicolas, and Ellis Patrick. 2020. *spicyR: Spatial analysis of in situ cytometry data*. https://doi.org/10.18129/B9.bioc.spicyR.

Cang, Zixuan, and Qing Nie. 2020. "Inferring spatial and signaling relationships between cells from single cell transcriptomic data." *Nature Communications* 11 (1): 2084. https://doi.org/10.1038/s41467-020-15968-5.

Cankaya, Murat, Ana Hernandez, Mehmet Ciftci, Sukru Beydemir, Hasan Ozdemir, Harun Budak, Ilhami Gulcin, et al. 2007. "An analysis of expression patterns of genes encoding proteins with catalytic activities." *BMC Genomics* 8 (1): 232. https://doi.org/10.1186/1471-2164-8-232.

Cao, Junyue, Malte Spielmann, Xiaojie Qiu, Xingfan Huang, Daniel M Ibrahim, Andrew J Hill, Fan Zhang, et al. 2019. "The single-cell transcriptional landscape of mammalian organogenesis." *Nature* 566 (7745): 496–502. https://doi.org/10.1038/s41586-019-0969-x.

Carlberg, Konstantin, Marina Korotkova, Ludvig Larsson, Anca I Catrina, Patrik L Ståhl, and Vivianne Malmström. 2019. "Exploring inflammatory signatures in arthritic joint biopsies with Spatial Transcriptomics." *Scientific Reports* 9 (1): 18975. https://doi.org/10.1038/s41598-019-55441-y.

Carson, James P., Christina Thaller, and Gregor Eichele. 2002. "A transcriptome atlas of the mouse brain at cellular resolution." *Current Opinion in Neurobiology* 12 (5): 562–65. https://doi.org/10.1016/S0959-4388(02)00356-2.

Carter, Mark G. 2003. "In Situ-Synthesized Novel Microarray Optimized for Mouse Stem Cell and Early Developmental Expression Profiling." *Genome Research* 13 (5): 1011–21. https://doi.org/10.1101/gr.878903.

Chen, Ao, Sha Liao, Mengnan Cheng, Kailong Ma, Liang Wu, Yiwei Lai, Jin Yang, et al. 2021. "Large field of view-spatially resolved transcriptomics at nanoscale resolution." *bioRxiv*, January, 2021.01.17.427004. https://doi.org/10.1101/2021.01.17.427004.

Chen, Fei, Paul W. Tillberg, and Edward S. Boyden. 2015. "Expansion microscopy." *Science* 347 (6221): 543–48. https://doi.org/10.1126/science.1260088.

Chen, Fei, Asmamaw T. Wassie, Allison J. Cote, Anubhav Sinha, Shahar Alon, Shoh Asano, Evan R. Daugharthy, et al. 2016. "Nanoscale imaging of RNA with expansion microscopy." *Nature Methods* 13 (8): 679–84. https://doi.org/10.1038/nmeth.3899.

Chen, Kok Hao, Alistair N. Boettiger, Jeffrey R. Moffitt, Siyuan Wang, and Xiaowei Zhuang. 2015. "Spatially resolved, highly multiplexed RNA profiling in single cells." *Science*. https://doi.org/10.1126/science.aaa6090.

Chen, Shuonan, Jackson Loper, Xiaoyin Chen, Alex Vaughan, Anthony M Zador, and Liam Paninski. 2021. "BARcode DEmixing through Non-negative Spatial Regression (BarDensr)." *PLOS Computational Biology* 17 (3): e1008256. https://doi.org/10.1371/journal.pcbi.1008256.

Chen, Wei-Ting, Ashley Lu, Katleen Craessaerts, Benjamin Pavie, Carlo Sala Frigerio, Nikky Corthout, Xiaoyan Qian, et al. 2020. "Spatial Transcriptomics and In Situ Sequencing to Study Alzheimer's Disease." *Cell* 0 (0). https://doi.org/10.1016/j.cell.2020.06.038.

Chen, Xiaoyin, Yu-Chi Sun, Huiqing Zhan, Justus M Kebschull, Stephan Fischer, Katherine Matho, Z Josh Huang, Jesse Gillis, and Anthony M Zador. 2019. "High-Throughput Mapping of Long-Range Neuronal Projection Using <em>In Situ</em> Sequencing." *Cell* 179 (3): 772–786.e19. https://doi.org/10.1016/j.cell.2019.09.023.

Chin, Mark H., Alex B. Geng, Arshad H. Khan, Wei-Jun Qian, Vladislav A. Petyuk, Jyl Boline, Shawn Levy, et al. 2007. "A genome-scale map of expression for a mouse brain section obtained using voxelation." *Physiological Genomics* 30 (3): 313–21. https://doi.org/10.1152/physiolgenomics.00287.2006.

Cho, Chun-Seok, Jingyue Xi, Sung-Rye Park, Jer-En Hsu, Myungjin Kim, Goo Jun, Hyun-Min Kang, and Jun Hee Lee. 2021. "Seq-Scope: Submicrometer-resolution spatial transcriptomics for single cell and subcellular studies." *bioRxiv*, January, 2021.01.25.427807. https://doi.org/10.1101/2021.01.25.427807.

Ciolli Mattioli, Camilla, Aviv Rom, Vedran Franke, Koshi Imami, Gerard Arrey, Mandy Terne, Andrew Woehler, Altuna Akalin, Igor Ulitsky, and Marina Chekulaeva. 2019. "Alternative 3 UTRs direct localization of functionally diverse protein isoforms in neuronal compartments." *Nucleic Acids Research* 47 (5): 2560–73. https://doi.org/10.1093/nar/gky1270.

Clarkson, Melissa D. 2016. "Representation of anatomy in online atlases and databases: a survey and collection of patterns for interface design." *BMC Developmental Biology* 16 (1): 18. https://doi.org/10.1186/s12861-016-0116-y.

Cleary, Brian, Brooke Simonton, Jon Bezney, Evan Murray, Shahul Alam, Anubhav Sinha, Ehsan Habibi, et al. 2021. "Compressed sensing for highly efficient imaging transcriptomics." *Nature Biotechnology*. https://doi.org/10.1038/s41587-021-00883-x.

Codeluppi, Simone, Lars E. Borm, Amit Zeisel, Gioele La Manno, Josina A. van Lunteren, Camilla I. Svensson, and Sten Linnarsson. 2018. "Spatial organization of the somatosensory cortex revealed by osmFISH." *Nature Methods*. https://doi.org/10.1038/s41592-018-0175-z.

Combs, Peter A., and Michael B. Eisen. 2013. "Sequencing mRNA from Cryo-Sliced Drosophila Embryos to Determine Genome-Wide Spatial Patterns of Gene Expression." Edited by Barbara Jennings. *PLoS ONE* 8 (8): e71820. https://doi.org/10.1371/journal.pone.0071820.

Combs, Peter A., and Hunter B. Fraser. 2018. "Spatially varying cis-regulatory divergence in Drosophila embryos elucidates cis-regulatory logic." Edited by Claude Desplan. *PLOS Genetics* 14 (11): e1007631. https://doi.org/10.1371/journal.pgen.1007631.

Coskun, Ahmet F., and Long Cai. 2016. "Dense transcript profiling in single cells by image correlation decoding." *Nature Methods* 13 (8): 657–60. https://doi.org/10.1038/nmeth.3895.

Coudry, Renata A., Sibele I. Meireles, Radka Stoyanova, Harry S. Cooper, Alan Carpino, Xianqun Wang, Paul F. Engstrom, and Margie L. Clapper. 2007. "Successful Application of Microarray Technology to Microdissected Formalin-Fixed, Paraffin-Embedded Tissue." *The Journal of Molecular Diagnostics* 9 (1): 70–79. https://doi.org/10.2353/jmoldx.2007.060004.

Crosetto, Nicola, Magda Bienko, and Alexander Van Oudenaarden. 2015. "Spatially resolved transcriptomics and beyond." https://doi.org/10.1038/nrg3832.

Cunnea, Paula, Jill McMahon, Enda O'Connell, Kaveh Mashayekhi, Una Fitzgerald, and Stephen McQuaid. 2010. "Gene expression analysis of the microvascular compartment in multiple sclerosis using laser microdissected blood vessels." *Acta Neuropathologica* 119 (5): 601–15. https://doi.org/10.1007/s00401-009-0618-9.

"Dana-Farber Core Facilities." n.d. Accessed October 14, 2020. https://www.dana-farber.org/research/core-facilities/.

Dar, Daniel, Nina Dar, Long Cai, and Dianne K Newman. 2021. "In situ single-cell activities of microbial populations revealed by spatial transcriptomics." *bioRxiv*, January, 2021.02.24.432792. https://doi.org/10.1101/2021.02.24.432792.

Darnell, Diana K., Simran Kaur, Stacey Stanislaw, Jay K. Konieczka, Tatiana A. Yatskievych, and Parker B. Antin. 2006. "MicroRNA expression during chick embryo development." *Developmental Dynamics* 235 (11): 3156–65. https://doi.org/10.1002/dvdy.20956.

De Giovanni, Marco, Valeria Cutillo, Amir Giladi, Eleonora Sala, Carmela G Maganuco, Chiara Medaglia, Pietro Di Lucia, et al. 2020. "Spatiotemporal regulation of type I interferon expression determines the antiviral polarization of CD4+ T cells." *Nature Immunology* 21 (3): 321–30. https://doi.org/10.1038/s41590-020-0596-6.

Delorey, Toni M, Carly G K Ziegler, Graham Heimberg, Rachelly Normand, Yiming Yang, Asa Segerstolpe, Domenic Abbondanza, et al. 2021. "A single-cell and spatial atlas of autopsy tissues reveals pathology and cellular targets of SARS-CoV-2." *bioRxiv*, January, 2021.02.25.430130. https://doi.org/10.1101/2021.02.25.430130.

Diez-Roux, Graciana, Sandro Banfi, Marc Sultan, Lars Geffers, Santosh Anand, David Rozado, Alon Magen, et al. 2011. "A High-Resolution Anatomical Atlas of the Transcriptome in the Mouse Embryo." Edited by Gregory S. Barsh. *PLoS Biology* 9 (1): e1000582. https://doi.org/10.1371/journal.pbio.1000582.

Dirks, Robert M., and Niles A. Pierce. 2004. "From The Cover: Triggered amplification by hybridization chain reaction." *Proceedings of the National Academy of Sciences* 101 (43): 15275–8. https://doi.org/10.1073/pnas.0407024101.

Dobin, Alexander, Carrie A Davis, Felix Schlesinger, Jorg Drenkow, Chris Zaleski, Sonali Jha, Philippe Batut, Mark Chaisson, and Thomas R Gingeras. 2012. "STAR: ultrafast universal RNA-seq aligner." *Bioinformatics* 29 (1): 15–21. https://doi.org/10.1093/bioinformatics/bts635.

Dries, Ruben, Qian Zhu, Chee-Huat Linus Eng, Arpan Sarkar, Feng Bao, Rani E George, Nico Pierson, Long Cai, and Guo-Cheng Yuan. 2019. "Giotto, a pipeline for integrative analysis and visualization of single-cell spatial transcriptomic data." *bioRxiv*, May, 701680. https://doi.org/10.1101/701680.

Drmanac, R., Andrew B. Sparks, Matthew J. Callow, Aaron L. Halpern, Norman L. Burns, Bahram G. Kermani, Paolo Carnevali, et al. 2010. "Human Genome Sequencing Using Unchained Base Reads on Self-Assembling DNA Nanoarrays." *Science* 327 (5961): 78–81. https://doi.org/10.1126/science.1181498.

Droin, Colas, Jakob El Kholtei, Keren Bahar Halpern, Clémence Hurni, Milena Rozenberg, Sapir Muvkadi, Shalev Itzkovitz, and Felix Naef. 2020. "Space-time logic of liver gene expression at sublobular scale." *bioRxiv*, January, 2020.03.05.976571. https://doi.org/10.1101/2020.03.05.976571.

"DSP Technology Access Program (TAP)." n.d. Accessed March 30, 2021.

Dulac, Catherine, and Richard Axel. 1995. "A novel family of genes encoding putative pheromone receptors in mammals." *Cell* 83 (2): 195–206. https://doi.org/https://doi.org/10.1016/0092-8674(95)90161-2.

Durruthy-Durruthy, Jens, Mark Wossidlo, Sunil Pai, Yusuke Takahashi, Gugene Kang, Larsson Omberg, Bertha Chen, Hiromitsu Nakauchi, Renee Reijo Pera, and Vittorio Sebastiano. 2016. "Spatiotemporal Reconstruction of the Human Blastocyst by Single-Cell Gene-Expression Analysis Informs Induction of Naive Pluripotency." *Developmental Cell* 38 (1): 100–115. https://doi.org/10.1016/j.devcel.2016.06.014.

Durruthy-Durruthy, Robert, Assaf Gottlieb, Byron H. Hartman, Jörg Waldhaus, Roman D. Laske, Russ Altman, and Stefan Heller. 2014. "Reconstruction of the Mouse Otocyst and Early Neuroblast Lineage at Single-Cell Resolution." *Cell* 157 (4): 964–78. https://doi.org/10.1016/j.cell.2014.03.036.

Ebbing, Annabel, Abel Vertesy, Marco Betist, Bastiaan Spanjaard, Jan Philipp Junker, Eugene Berezikov, Alexander van Oudenaarden, and Hendrik Korswagen. 2018. "Spatial transcriptomics of C. elegans males and hermaphrodites identifies novel fertility genes." *bioRxiv*, June, 348201. https://doi.org/10.1101/348201.

Eberwine, J, H Yeh, K Miyashiro, Y Cao, S Nair, R Finnell, M Zettel, and P Coleman. 1992. "Analysis of gene expression in single live neurons." *Proceedings of the National Academy of Sciences* 89 (7): 3010 LP–3014. `https://doi.org/10.1073/pnas.89.7.3010`.

Edsgärd, Daniel, Per Johnsson, and Rickard Sandberg. 2018. "Identification of spatial expression trends in single-cell gene expression data." *Nature Methods* 15 (5): 339–42. `https://doi.org/10.1038/nmeth.4634`.

Efremova, Mirjana, Miquel Vento-Tormo, Sarah A Teichmann, and Roser Vento-Tormo. 2020. "CellPhoneDB: inferring cell–cell communication from combined expression of multi-subunit ligand–receptor complexes." *Nature Protocols* 15 (4): 1484–1506. `https://doi.org/10.1038/s41596-020-0292-x`.

Eichenberger, Bastian Th., YinXiu Zhan, Markus Rempfler, Luca Giorgetti, and Jeffrey A Chao. 2020. "deepBlink: Threshold-independent detection and localization of diffraction-limited spots." *bioRxiv*, January, 2020.12.14.422631. `https://doi.org/10.1101/2020.12.14.422631`.

Elosua, Marc, Paula Nieto, Elisabetta Mereu, Ivo Gut, and Holger Heyn. 2020. "SPOTlight: Seeded NMF regression to Deconvolute Spatial Transcriptomics Spots with Single-Cell Transcriptomes." *bioRxiv*, June, 2020.06.03.131334. `https://doi.org/10.1101/2020.06.03.131334`.

Emmert-Buck, Michael R., Robert F. Bonner, Paul D. Smith, Rodrigo F. Chuaqui, Zhengping Zhuang, Seth R. Goldstein, Rhonda A. Weiss, and Lance A. Liotta. 1996. "Laser Capture Microdissection." *Science* 274 (5289): 998–1001. `https://doi.org/10.1126/science.274.5289.998`.

Eng, Chee Huat Linus, Michael Lawson, Qian Zhu, Ruben Dries, Noushin Koulena, Yodai Takei, Jina Yun, et al. 2019. "Transcriptome-scale super-resolved imaging in tissues by RNA seqFISH+." *Nature* 568 (7751): 235–39. `https://doi.org/10.1038/s41586-019-1049-y`.

Eng, Chee-Huat Linus, Sheel Shah, Julian Thomassie, and Long Cai. 2017. "Profiling the transcriptome with RNA SPOTs." *Nature Methods* 14 (12): 1153–5. `https://doi.org/10.1038/nmeth.4500`.

Fan, Zhen, Runsheng Chen, and Xiaowei Chen. 2020. "SpatialDB: a database for spatially resolved transcriptomes." *Nucleic Acids Research* 48 (D1): D233–D237. `https://doi.org/10.1093/nar/gkz934`.

Farris, Shannon, James M Ward, Kelly E Carstens, Mahsa Samadi, Yu Wang, and Serena M Dudek. 2019. "Hippocampal Subregions Express Distinct Dendritic Transcriptomes that Reveal Differences in Mitochondrial Function in CA2." *Cell Reports* 29 (2): 522–539.e6. `https://doi.org/10.1016/j.celrep.2019.08.093`.

Fazal, Furqan M., Shuo Han, Kevin R. Parker, Pornchai Kaewsapsak, Jin Xu, Alistair N. Boettiger, Howard Y. Chang, and Alice Y. Ting. 2019. "Atlas of

Subcellular RNA Localization Revealed by APEX-Seq." *Cell* 178 (2): 473–490.e26. https://doi.org/10.1016/j.cell.2019.05.027.

Femino, Andrea M, Fredric S Fay, Kevin Fogarty, and Robert H Singer. 1998. "Visualization of Single RNA Transcripts in Situ." *Science* 280 (5363): 585 LP–590. https://doi.org/10.1126/science.280.5363.585.

Fire, Andrew, and Si Qun Xu. 1995. "Rolling replication of short DNA circles." *Proceedings of the National Academy of Sciences* 92 (10): 4641–5. https://doi.org/10.1073/pnas.92.10.4641.

Foley, Joseph W., Chunfang Zhu, Philippe Jolivet, Shirley X. Zhu, Peipei Lu, Michael J. Meaney, and Robert B. West. 2019. "Gene expression profiling of single cells from archival tissue with laser-capture microdissection and Smart-3SEQ." *Genome Research* 29 (11): 1816–25. https://doi.org/10.1101/gr.234807.118.

Foreman, Robert, and Roy Wollman. 2019. "Mammalian gene expression variability is explained by underlying cell state." *bioRxiv*. https://doi.org/10.1101/626424.

Forrester, Lesley M., Andras Nagy, Mehran Sam, Alistair Watt, Lois Stevenson, Alan Bernstein, Alexandra L. Joyner, and Wolfgang Wurst. 1996. "An induction gene trap screen in embryonic stem cells: Identification of genes that respond to retinoic acid in vitro." *Proceedings of the National Academy of Sciences of the United States of America* 93 (4): 1677–82. https://doi.org/10.1073/pnas.93.4.1677.

Fowlkes, Charless C., Cris L. Luengo Hendriks, Soile V. E. Keränen, Gunther H. Weber, Oliver Rübel, Min-Yu Huang, Sohail Chatoor, et al. 2008. "A Quantitative Spatiotemporal Atlas of Gene Expression in the Drosophila Blastoderm." *Cell* 133 (2): 364–74. https://doi.org/10.1016/j.cell.2008.01.053.

Friedrich, G., and P. Soriano. 1991. "Promoter traps in embryonic stem cells: a genetic screen to identify and mutate developmental genes in mice." *Genes & Development* 5 (9): 1513–23. https://doi.org/10.1101/gad.5.9.1513.

Frise, Erwin, Ann S Hammonds, and Susan E Celniker. 2010. "Systematic image-driven analysis of the spatial Drosophila embryonic expression landscape." *Molecular Systems Biology* 6 (1): 345. https://doi.org/10.1038/msb.2009.102.

Frohman, M. A., M. K. Dush, and G. R. Martin. 1988. "Rapid production of full-length cDNAs from rare transcripts: amplification using a single gene-specific oligonucleotide primer." *Proceedings of the National Academy of Sciences* 85 (23): 8998–9002. https://doi.org/10.1073/pnas.85.23.8998.

Fu, Xiaonan, Li Sun, Jane Y Chen, Runze Dong, Yiing Lin, Richard D Palmiter, Shin Lin, and Liangcai Gu. 2021. "Continuous Polony Gels for Tissue Mapping with High Resolution and RNA Capture Efficiency." *bioRxiv*, January, 2021.03.17.435795. https://doi.org/10.1101/2021.03.17.435795.

Gall, Joseph G, Mary Lou, Pardue Kline, and Norman H Giles. 1969. "Formation and Detecction of RNA-DNA Hybrid Molecules in Cytological Preparations." *PNAS* 63 (2): 378–83. https://doi.org/10.1073/pnas.63.2.378.

Gawantka, Volker, Nicolas Pollet, Hajo Delius, Martin Vingron, Ralf Pfister, Rebecca Nitsch, Claudia Blumenstock, and Christof Niehrs. 1998. "Gene expression screening in Xenopus identifies molecular pathways, predicts gene function and provides a global view of embryonic patterning." *Mechanisms of Development* 77 (2): 95–141. https://doi.org/10.1016/S0925-4773(98)00115-4.

Gebhardt, Rolf. 2014. "Liver zonation: Novel aspects of its regulation and its impact on homeostasis." *World Journal of Gastroenterology* 20 (26): 8491. https://doi.org/10.3748/wjg.v20.i26.8491.

Geffers, Lars, Benjamin Tetzlaff, Xiao Cui, Jun Yan, and Gregor Eichele. 2012. "METscout: a pathfinder exploring the landscape of metabolites, enzymes and transporters." *Nucleic Acids Research* 41 (D1): D1047–D1054. https://doi.org/10.1093/nar/gks886.

"Gene Expression Panels | NanoString Technologies." n.d. Accessed September 15, 2020. https://www.nanostring.com/products/gene-expression-panels/gene-expression-panels-overview.

Ghazanfar, Shila, Yingxin Lin, Xianbin Su, David Ming Lin, Ellis Patrick, Ze-Guang Han, John C Marioni, and Jean Yee Hwa Yang. 2020. "Investigating higher-order interactions in single-cell data with scHOT." *Nature Methods* 17 (8): 799–806. https://doi.org/10.1038/s41592-020-0885-x.

Giacomello, Stefania, Fredrik Salmén, Barbara K. Terebieniec, Sanja Vickovic, José Fernandez Navarro, Andrey Alexeyenko, Johan Reimegård, et al. 2017. "Spatially resolved transcriptome profiling in model plant species." *Nature Plants*. https://doi.org/10.1038/nplants.2017.61.

Giani, Alice Maria, Guido Roberto Gallo, Luca Gianfranceschi, and Giulio Formenti. 2020. "Long walk to genomics: History and current approaches to genome sequencing and assembly." *Computational and Structural Biotechnology Journal* 18 (January): 9–19. https://doi.org/10.1016/j.csbj.2019.11.002.

Gilchrist, Michael J., Mikkel B. Christensen, Odile Bronchain, Frédéric Brunet, Albert Chesneau, Ursula Fenger, Timothy J. Geach, et al. 2009. "Database of queryable gene expression patterns for Xenopus." *Developmental Dynamics* 238 (6): 1379–88. https://doi.org/10.1002/dvdy.21940.

Giolai, Michael, Walter Verweij, Ashleigh Lister, Darren Heavens, Iain Macaulay, and Matthew D. Clark. 2019. "Spatially resolved transcriptomics reveals plant host responses to pathogens." *Plant Methods* 15 (1): 114. https://doi.org/10.1186/s13007-019-0498-5.

Glaser, Joshua I., Bradley M. Zamft, George M. Church, and Konrad P. Kording. 2015. "Puzzle Imaging: Using Large-Scale Dimensionality Reduction

Algorithms for Localization." Edited by Joseph Najbauer. *PLOS ONE* 10 (7): e0131593. https://doi.org/10.1371/journal.pone.0131593.

Goh, Jolene Jie Lin, Nigel Chou, Wan Yi Seow, Norbert Ha, Chung Pui Paul Cheng, Yun Ching Chang, Ziqing Winston Zhao, and Kok Hao Chen. 2020. "Highly specific multiplexed RNA imaging in tissues with split-FISH." *Nature Methods* 17 (7): 689–93. https://doi.org/10.1038/s41592-020-0858-0.

Gong, Shiaoching, Chen Zheng, Martin L. Doughty, Kasia Losos, Nicholas Didkovsky, Uta B. Schambra, Norma J. Nowak, et al. 2003. "A gene expression atlas of the central nervous system based on bacterial artificial chromosomes." *Nature* 425 (6961): 917–25. https://doi.org/10.1038/nature02033.

Gossler, Achim, A. Joyner, Janet Rossant, and W. Skarnes. 1989. "Mouse embryonic stem cells and reporter constructs to detect developmentally regulated genes." *Science* 244 (4903): 463–65. https://doi.org/10.1126/science.2497519.

Govek, Kiya W, Venkata S Yamajala, and Pablo G Camara. 2019. "Clustering-independent analysis of genomic data using spectral simplicial theory." *PLOS Computational Biology* 15 (11): e1007509. https://doi.org/10.1371/journal.pcbi.1007509.

Grange, Pascal, Jason W. Bohland, Benjamin W. Okaty, Ken Sugino, Hemant Bokil, Sacha B. Nelson, Lydia Ng, Michael Hawrylycz, and Partha P. Mitra. 2014. "Cell-type-based model explaining coexpression patterns of genes in the brain." *Proceedings of the National Academy of Sciences* 111 (14): 5397–5402. https://doi.org/10.1073/pnas.1312098111.

Gregory, J M, K McDade, M R Livesey, I Croy, S Marion de Proce, T Aitman, S Chandran, and C Smith. 2020. "Spatial transcriptomics identifies spatially dysregulated expression of GRM3 and USP47 in amyotrophic lateral sclerosis." *Neuropathology and Applied Neurobiology* 46 (5): 441–57. https://doi.org/10.1111/nan.12597.

Greulich, Karl Otto. 1999. "Introduction: The history of using light as a working tool." In *Micromanipulation by Light in Biology and Medicine*, 1–6. Boston, MA: Birkhäuser Boston. https://doi.org/10.1007/978-1-4612-4110-2_1.

Grün, Dominic, Lennart Kester, and Alexander van Oudenaarden. 2014. "Validation of noise models for single-cell transcriptomics." *Nature Methods* 11 (6): 637–40. https://doi.org/10.1038/nmeth.2930.

Gurunathan, Rajalakshmi, Bernard Van Emden, Sethuraman Panchanathan, and Sudhir Kumar. 2004. "Identifying spatially similar gene expression patterns in early stage fruit fly embryo images: Binary feature versus invariant moment digital representations." *BMC Bioinformatics* 5 (1): 202. https://doi.org/10.1186/1471-2105-5-202.

Gyllborg, Daniel, Christoffer Mattsson Langseth, Xiaoyan Qian, Sergio Marco Salas, Markus Hilscher, Ed Lein, and Mats Nilsson. 2020. "Hybridization-based In Situ Sequencing (HybISS): spatial transcriptomic detection in human and mouse brain tissue." *bioRxiv*, February, 2020.02.03.931618. https://doi.org/10.1101/2020.02.03.931618.

Halpern, Keren Bahar, Rom Shenhav, Hassan Massalha, Beata Toth, Adi Egozi, Efi E. Massasa, Chiara Medgalia, et al. 2018. "Paired-cell sequencing enables spatial gene expression mapping of liver endothelial cells." *Nature Biotechnology* 36 (10): 962. https://doi.org/10.1038/nbt.4231.

Halpern, Keren Bahar, Rom Shenhav, Orit Matcovitch-Natan, Beáta Tóth, Doron Lemze, Matan Golan, Efi E. Massasa, et al. 2017. "Single-cell spatial reconstruction reveals global division of labour in the mammalian liver." *Nature* 542 (7641): 1–5. https://doi.org/10.1038/nature21065.

Hammonds, Ann S., Christopher A. Bristow, William W. Fisher, Richard Weiszmann, Siqi Wu, Volker Hartenstein, Manolis Kellis, Bin Yu, Erwin Frise, and Susan E. Celniker. 2013. "Spatial expression of transcription factors in Drosophila embryonic organ development." *Genome Biology* 14 (12): R140. https://doi.org/10.1186/gb-2013-14-12-r140.

Hanchuan Peng, Fuhui Long, M. B. Eisen, and E. W. Myers. 2006. "Clustering Gene Expression Patterns of Fly Embryos." In *3rd Ieee International Symposium on Biomedical Imaging: Macro to Nano, 2006.*, 2006:1144–7. IEEE. https://doi.org/10.1109/ISBI.2006.1625125.

Hao, Minsheng, Kui Hua, and Xuegong Zhang. 2021. "SOMDE: A scalable method for identifying spatially variable genes with self-organizing map." *bioRxiv*, January, 2020.12.10.419549. https://doi.org/10.1101/2020.12.10.419549.

Harding, Simon D., Chris Armit, Jane Armstrong, Jane Brennan, Ying Cheng, Bernard Haggarty, Derek Houghton, et al. 2011. "The GUDMAP database - an online resource for genitourinary research." *Development* 138 (13): 2845–53. https://doi.org/10.1242/dev.063594.

Harrison, P. R., D. Conkie, J. Paul, and K. Jones. 1973. "Localisation of cellular globin messenger RNA by in situ hybridisation to complementary DNA." *FEBS Letters* 32 (1): 109–12. https://doi.org/10.1016/0014-5793(73)80749-5.

Hashimshony, Tamar, Naftalie Senderovich, Gal Avital, Agnes Klochendler, Yaron de Leeuw, Leon Anavy, Dave Gennert, et al. 2016. "CEL-Seq2: sensitive highly-multiplexed single-cell RNA-Seq." *Genome Biology* 17 (1): 77. https://doi.org/10.1186/s13059-016-0938-8.

Hashimshony, Tamar, Florian Wagner, Noa Sher, and Itai Yanai. 2012. "CEL-Seq: Single-Cell RNA-Seq by Multiplexed Linear Amplification." *Cell Reports* 2 (3): 666–73. https://doi.org/https://doi.org/10.1016/j.celrep.2012.08.003.

Haudry, Yannick, Hugo Berube, Ivica Letunic, P.-D. Weeber, Julien Gagneur, Charles Girardot, Misha Kapushesky, et al. 2007. "4DXpress: a database for cross-species expression pattern comparisons." *Nucleic Acids Research* 36 (Database): D847–D853. https://doi.org/10.1093/nar/gkm797.

Hawrylycz, Michael J., Ed S. Lein, Angela L. Guillozet-Bongaarts, Elaine H. Shen, Lydia Ng, Jeremy A. Miller, Louie N. van de Lagemaat, et al. 2012. "An anatomically comprehensive atlas of the adult human brain transcriptome." *Nature* 489 (7416): 391–99. https://doi.org/10.1038/nature11405.

He, Bryan, Ludvig Bergenstråhle, Linnea Stenbeck, Abubakar Abid, Alma Andersson, Åke Borg, Jonas Maaskola, Joakim Lundeberg, and James Zou. 2020. "Integrating spatial gene expression and breast tumour morphology via deep learning." *Nature Biomedical Engineering*, June, 1–8. https://doi.org/10.1038/s41551-020-0578-x.

Heffel, Andreas, Peter F. Stadler, Sonja J. Prohaska, Gerhard Kauer, and Jens-Peer Kuska. 2008. "Process flow for classification and clustering of fruit fly gene expression patterns." In *2008 15th Ieee International Conference on Image Processing*, 721–24. IEEE. https://doi.org/10.1109/ICIP.2008.4711856.

Hemby, Scott E, Stephen D Ginsberg, Brian Brunk, Steven E Arnold, John Q Trojanowski, and James H Eberwine. 2002. "Gene Expression Profile for Schizophrenia: Discrete Neuron Transcription Patterns in the Entorhinal Cortex." *Archives of General Psychiatry* 59 (7): 631–40. https://doi.org/10.1001/archpsyc.59.7.631.

Henrich, Thorsten. 2003. "MEPD: a Medaka gene expression pattern database." *Nucleic Acids Research* 31 (1): 72–74. https://doi.org/10.1093/nar/gkg017.

Hiwatashi, Yuji, Tomoaki Nishiyama, Tomomichi Fujita, and Mitsuyasu Hasebe. 2001. "Establishment of gene-trap and enhancer-trap systems in the moss Physcomitrella patens." *The Plant Journal* 28 (1): 105–16. https://doi.org/10.1046/j.1365-313X.2001.01121.x.

Hope, I A. 1991. "'Promoter trapping' in Caenorhabditis elegans." Vol. 113. https://dev.biologists.org/content/develop/113/2/399.full.pdf.

Howe, Douglas G., Yvonne M. Bradford, Anne Eagle, David Fashena, Ken Frazer, Patrick Kalita, Prita Mani, et al. 2017. "The Zebrafish Model Organism Database: New support for human disease models, mutation details, gene expression phenotypes and searching." *Nucleic Acids Research* 45 (D1): D758–D768. https://doi.org/10.1093/nar/gkw1116.

Huber, D., L. Voith von Voithenberg, and G. V. Kaigala. 2018. "Fluorescence in situ hybridization (FISH): History, limitations and what to expect from micro-scale FISH?" *Micro and Nano Engineering* 1 (November): 15–24. https://doi.org/10.1016/j.mne.2018.10.006.

Hunt-Newbury, Rebecca, Ryan Viveiros, Robert Johnsen, Allan Mah, Dina Anastas, Lily Fang, Erin Halfnight, et al. 2007. "High-Throughput In Vivo Analysis of Gene Expression in Caenorhabditis elegans." Edited by John Sulston. *PLoS Biology* 5 (9): e237. https://doi.org/10.1371/journal.pbio.0050237.

Hwang, William L, Karthik A Jagadeesh, Jimmy A Guo, Hannah I Hoffman, Payman Yadollahpour, Rahul Mohan, Eugene Drokhlyansky, et al. 2020. "Single-nucleus and spatial transcriptomics of archival pancreatic cancer reveals multi-compartment reprogramming after neoadjuvant treatment." *bioRxiv*, August, 2020.08.25.267336. https://doi.org/10.1101/2020.08.25.267336.

Islam, Saiful, Una Kjällquist, Annalena Moliner, Pawel Zajac, Jian-Bing Fan, Peter Lönnerberg, and Sten Linnarsson. 2011. "Characterization of the single-cell transcriptional landscape by highly multiplex RNA-seq." *Genome Research* 21 (7): 1160–7. https://doi.org/10.1101/gr.110882.110.

Jagalur, Manjunatha, Chris Pal, Erik Learned-Miller, R Thomas Zoeller, and David Kulp. 2007. "Analyzing in situ gene expression in the mouse brain with image registration, feature extraction and block clustering." *BMC Bioinformatics* 8 (Suppl 10): S5. https://doi.org/10.1186/1471-2105-8-S10-S5.

Janning, Wilfried. 1997. "FlyView, aDrosophilaimage database, and other-Drosophiladatabases." *Seminars in Cell & Developmental Biology* 8 (5): 469–75. https://doi.org/https://doi.org/10.1006/scdb.1997.0172.

Jayaraman, Karthik, Sethuraman Panchanathan, and Sudhir Kumar. 2001. "Classification and Indexing of Gene Expression Images."

Jemt, Anders, Fredrik Salmén, Anna Lundmark, Annelie Mollbrink, José Fernández Navarro, Patrik L. Ståhl, Tülay Yucel-Lindberg, and Joakim Lundeberg. 2016. "An automated approach to prepare tissue-derived spatially barcoded RNA-sequencing libraries." *Scientific Reports*. https://doi.org/10.1038/srep37137.

Jenett, Arnim, Gerald M. Rubin, Teri T. B. Ngo, David Shepherd, Christine Murphy, Heather Dionne, Barret D. Pfeiffer, et al. 2012. "A GAL4-Driver Line Resource for Drosophila Neurobiology." *Cell Reports* 2 (4): 991–1001. https://doi.org/10.1016/j.celrep.2012.09.011.

Jerby-Arnon, Livnat, and Aviv Regev. 2020. "Mapping multicellular programs from single-cell profiles." *bioRxiv*, January, 2020.08.11.245472. https://doi.org/10.1101/2020.08.11.245472.

Ji, Andrew L, Adam J Rubin, Kim Thrane, Sizun Jiang, David L Reynolds, Robin M Meyers, Margaret G Guo, et al. 2020. "Multimodal Analysis of Composition and Spatial Architecture in Human Squamous Cell Carcinoma." *Cell* 182 (2): 497–514.e22. https://doi.org/https://doi.org/10.1016/j.cell.2020.05.039.

Ji, Shuiwang, Ying Xin Li, Zhi Hua Zhou, Sudhir Kumar, and Jieping Ye. 2009. "A bag-of-words approach for Drosophila gene expression pattern annotation." *BMC Bioinformatics* 10 (1): 119. https://doi.org/10.1186/1471-2105-10-119.

Ji, Shuiwang, Liang Sun, Rong Jin, Sudhir Kumar, and Jieping Ye. 2008. "Automated annotation of Drosophila gene expression patterns using a controlled vocabulary." *Bioinformatics* 24 (17): 1881–8. https://doi.org/10.1093/bioinformatics/btn347.

Ji, Shuiwang, Lei Yuan, Ying-Xin Li, Zhi-Hua Zhou, Sudhir Kumar, and Jieping Ye. 2009. "Drosophila gene expression pattern annotation using sparse features and term-term interactions." In *Proceedings of the 15th Acm Sigkdd International Conference on Knowledge Discovery and Data Mining - Kdd '09*, 407. New York, New York, USA: ACM Press. https://doi.org/10.1145/1557019.1557068.

Joglekar, Anoushka, Andrey Prjibelski, Ahmed Mahfouz, Paul Collier, Susan Lin, Anna Katharina Schlusche, Jordan Marrocco, et al. 2020. "Cell-type, single-cell, and spatial signatures of brain-region specific splicing in postnatal development." *bioRxiv*, January, 2020.08.27.268730. https://doi.org/10.1101/2020.08.27.268730.

John, H. A., M. L. Birnstiel, and K. W. Jones. 1969. "RNA-DNA hybrids at the cytological level." *Nature* 223 (5206): 582–87. https://doi.org/10.1038/223582a0.

"Johns Hopkins Cell Imaging Core Facility." n.d. Accessed October 14, 2020. https://www.hopkinsmedicine.org/kimmel%7B/_%7Dcancer%7B/_%7Dcenter/research/shared%7B/_%7Dresources/cell%7B/_%7Dimaging.html.

Johnson, Alexander A. T., Julian M. Hibberd, Céline Gay, Pauline A. Essah, Jim Haseloff, Mark Tester, and Emmanuel Guiderdoni. 2005. "Spatial control of transgene expression in rice (Oryza sativa L.) using the GAL4 enhancer trapping system." *The Plant Journal* 41 (5): 779–89. https://doi.org/10.1111/j.1365-313X.2005.02339.x.

Junker, Jan Philipp, Emily S. Noël, Victor Guryev, Kevin A. Peterson, Gopi Shah, Jan Huisken, Andrew P. McMahon, Eugene Berezikov, Jeroen Bakkers, and Alexander Van Oudenaarden. 2014. "Genome-wide RNA Tomography in the Zebrafish Embryo." *Cell*. https://doi.org/10.1016/j.cell.2014.09.038.

Kacharmina, Janet Estee, Peter B Crino, and James B T - Methods in Enzymology Eberwine. 1999. "[1] Preparation of cDNA from single cells and subcellular regions." In *CDNA Preparation and Characterization*, 303:3–18. Academic Press. https://doi.org/https://doi.org/10.1016/S0076-6879(99)03003-7.

Kaewsapsak, Pornchai, David Michael Shechner, William Mallard, John L. Rinn, and Alice Y. Ting. 2017. "Live-cell mapping of organelle-associated RNAs via proximity biotinylation combined with protein-RNA crosslinking." *eLife* 6 (December). https://doi.org/10.7554/eLife.29224.

Kamme, Fredrik, Ranelle Salunga, Jingxue Yu, Da-Thao Tran, Jessica Zhu, Lin Luo, Anton Bittner, et al. 2003. "Single-Cell Microarray Analysis in Hippocampus CA1: Demonstration and Validation of Cellular Heterogeneity." *The Journal of Neuroscience* 23 (9): 3607 LP–3615. https://doi.org/10.1523/JNEUROSCI.23-09-03607.2003.

Karaiskos, Nikos, Philipp Wahle, Jonathan Alles, Anastasiya Boltengagen, Salah Ayoub, Claudia Kipar, Christine Kocks, Nikolaus Rajewsky, and Robert P. Zinzen. 2017. "The Drosophila embryo at single-cell transcriptome resolution." *Science* 358 (6360): 194–99. https://doi.org/10.1126/science.aan3235.

Karali, Marianthi, Ivana Peluso, Vincenzo A. Gennarino, Marchesa Bilio, Roberta Verde, Giampiero Lago, Pascal Dollé, and Sandro Banfi. 2010. "miRNeye: a microRNA expression atlas of the mouse eye." *BMC Genomics* 11 (1): 715. https://doi.org/10.1186/1471-2164-11-715.

Kawakami, Koichi, Gembu Abe, Tokuko Asada, Kazuhide Asakawa, Ryuichi Fukuda, Aki Ito, Pradeep Lal, et al. 2010. "ZTrap: Zebrafish gene trap and enhancer trap database." *BMC Developmental Biology* 10 (1): 105. https://doi.org/10.1186/1471-213X-10-105.

Kawashima, T. 2000. "MAGEST: MAboya Gene Expression patterns and Sequence Tags." *Nucleic Acids Research* 28 (1): 133–35. https://doi.org/10.1093/nar/28.1.133.

Ke, Rongqin, Marco Mignardi, Alexandra Pacureanu, Jessica Svedlund, Johan Botling, Carolina Wählby, and Mats Nilsson. 2013. "In situ sequencing for RNA analysis in preserved tissue and cells." *Nature Methods* 10 (9): 857–60. https://doi.org/10.1038/nmeth.2563.

Kerman, Ilan A., Bradley J. Buck, Simon J. Evans, Huda Akil, and Stanley J. Watson. 2006. "Combining laser capture microdissection with quantitative real-time PCR: Effects of tissue manipulation on RNA quality and gene expression." *Journal of Neuroscience Methods* 153 (1): 71–85. https://doi.org/10.1016/j.jneumeth.2005.10.010.

Kim, Minhee, Dong-Min Kim, and Dong-Eun Kim. 2020. "Label-free fluorometric detection of microRNA using isothermal rolling circle amplification generating tandem G-quadruplex." *Analyst* 145 (18): 6130–7. https://doi.org/10.1039/D0AN01329C.

Kitahara, Osamu, Yoichi Furukawa, Toshihiro Tanaka, Chikashi Kihara, Kenji Ono, Renpei Yanagawa, Marcelo E Nita, Toshihisa Takagi, Yusuke Nakamura, and Tatsuhiko Tsunoda. 2001. "Alterations of Gene Expression during Colorectal Carcinogenesis Revealed by cDNA Microarrays after Laser-Capture Microdissection of Tumor Tissues and Normal Epithelia." *Cancer Research* 61 (9): 3544 LP–3549. http://cancerres.aacrjournals.org/content/61/9/3544.abstract.

Kitamura, S, T Tanahashi, E Aoyagi, T Nakagawa, K Okamoto, T Kimura, H Miyamoto, et al. 2017. "Response Predictors of S-1, Cisplatin, and Docetaxel Combination Chemotherapy for Metastatic Gastric Cancer: Microarray Analysis of Whole Human Genes." *Oncology* 93 (2): 127–35. https://doi.org/10.1159/000464329.

Klein, Allon M., Linas Mazutis, Ilke Akartuna, Naren Tallapragada, Adrian Veres, Victor Li, Leonid Peshkin, David A. Weitz, and Marc W. Kirschner. 2015. "Droplet Barcoding for Single-Cell Transcriptomics Applied to Embryonic Stem Cells." *Cell* 161 (5): 1187–1201. https://doi.org/10.1016/j.cell.2015.04.044.

Klein, Christoph A, Stefan Seidl, Karina Petat-Dutter, Sonja Offner, Jochen B Geigl, Oleg Schmidt-Kittler, Nicole Wendler, et al. 2002. "Combined transcriptome and genome analysis of single micrometastatic cells." *Nature Biotechnology* 20 (4): 387–92. https://doi.org/10.1038/nbt0402-387.

Kloosterman, Wigard P., Erno Wienholds, Ewart de Bruijn, Sakari Kauppinen, and Ronald H A Plasterk. 2006. "In situ detection of miRNAs in animal embryos using LNA-modified oligonucleotide probes." *Nature Methods* 3 (1): 27–29. https://doi.org/10.1038/nmeth843.

Ko, M S H, S B H Ko, N Takahashi, K Nishiguchi, and K Abe. 1990. "Unbiased amplification of a highly complex mixture of DNA fragments by 'lone linker'-tagged PCR." *Nucleic Acids Research* 18 (14): 4293. https://doi.org/10.1093/nar/18.14.4293.

Ko, Younhee, Seth A. Ament, James A. Eddy, Juan Caballero, John C. Earls, Leroy Hood, and Nathan D. Price. 2013. "Cell type-specific genes show striking and distinct patterns of spatial expression in the mouse brain." *Proceedings of the National Academy of Sciences* 110 (8): 3095–3100. https://doi.org/10.1073/pnas.1222897110.

Kohman, Richie E, and George M Church. 2020. "Fluorescent in situ sequencing of DNA barcoded antibodies." *bioRxiv* 20 (April): 2020.04.27.060624. https://doi.org/10.1101/2020.04.27.060624.

Kopczynski, Casey C., Jasprina N. Noordermeer, Thomas L. Serano, W.-Y. Chen, John D. Pendleton, Suzanna Lewis, Corey S. Goodman, and Gerald M. Rubin. 1998. "A high throughput screen to identify secreted and transmembrane proteins involved in Drosophila embryogenesis." *Proceedings of the National Academy of Sciences* 95 (17): 9973–8. https://doi.org/10.1073/pnas.95.17.9973.

Korsunsky, Ilya, Nghia Millard, Jean Fan, Kamil Slowikowski, Fan Zhang, Kevin Wei, Yuriy Baglaenko, Michael Brenner, Po-ru Loh, and Soumya Raychaudhuri. 2019. "Fast, sensitive and accurate integration of single-cell data with Harmony." *Nature Methods* 16 (12): 1289–96. https://doi.org/10.1038/s41592-019-0619-0.

Köster, Johannes, Myles Brown, and X. Shirley Liu. 2019. "A Bayesian model for single cell transcript expression analysis on MERFISH data." *Bioinformatics*. https://doi.org/10.1093/bioinformatics/bty718.

Kudo, Lili C., Nancy Vi, Zhongcai Ma, Tony Fields, Nuraly K. Avliyakulov, Michael J. Haykinson, Anatol Bragin, and Stanislav L. Karsten. 2012. "Novel Cell and Tissue Acquisition System (CTAS): Microdissection of Live and Frozen Brain Tissues." Edited by Rafael Aldabe. *PLoS ONE* 7 (7): e41564. https://doi.org/10.1371/journal.pone.0041564.

Kueckelhaus, Jan, Jasmin von Ehr, Vidhya M Ravi, Paulina Will, Kevin Joseph, Juergen Beck, Ulrich G Hofmann, Daniel Delev, Oliver Schnell, and Dieter Henrik Heiland. 2020. "Inferring spatially transient gene expression pattern from spatial transcriptomic studies." *bioRxiv*, January, 2020.10.20.346544. https://doi.org/10.1101/2020.10.20.346544.

Kumar, Sudhir, Karthik Jayaraman, Sethuraman Panchanathan, Rajalakshmi Gurunathan, Ana Marti-Subirana, and Stuart J Newfeld. 2002. "BEST: A Novel Computational Approach for Comparing Gene Expression Patterns From Early Stages of <em>Drosophila melanogaster</em> Development." *Genetics* 162 (4): 2037 LP–2047. http://www.genetics.org/content/162/4/2037.abstract.

Kumar, Sudhir, Charlotte Konikoff, Maxwell Sanderford, Li Liu, Stuart Newfeld, Jieping Ye, and Rob J. Kulathinal. 2017. "FlyExpress 7: An Integrated Discovery Platform To Study Coexpressed Genes Using in situ Hybridization Images in Drosophila." *G3; Genes|Genomes|Genetics* 7 (8): 2791–7. https://doi.org/10.1534/g3.117.040345.

Kvon, Evgeny Z., Tomas Kazmar, Gerald Stampfel, J. Omar Yáñez-Cuna, Michaela Pagani, Katharina Schernhuber, Barry J. Dickson, and Alexander Stark. 2014. "Genome-scale functional characterization of Drosophila developmental enhancers in vivo." *Nature* 512 (7512): 91–95. https://doi.org/10.1038/nature13395.

Lacraz, Grégory P. A., Jan Philipp Junker, Monika M. Gladka, Bas Molenaar, Koen T. Scholman, Marta Vigil-Garcia, Danielle Versteeg, et al. 2017. "Tomo-Seq Identifies SOX9 as a Key Regulator of Cardiac Fibrosis During Ischemic Injury." *Circulation* 136 (15): 1396–1409. https://doi.org/10.1161/CIRCULATIONAHA.117.027832.

La Manno, Gioele, Ruslan Soldatov, Amit Zeisel, Emelie Braun, Hannah Hochgerner, Viktor Petukhov, Katja Lidschreiber, et al. 2018. "RNA velocity of single cells." *Nature* 560 (7719): 494–98. https://doi.org/10.1038/s41586-018-0414-6.

Landegren, U, R Kaiser, J Sanders, and L Hood. 1988. "A ligase-mediated gene detection technique." *Science* 241 (4869): 1077 LP–1080. https://doi.org/10.1126/science.3413476.

Lander, Eric S., Lauren M. Linton, Bruce Birren, Chad Nusbaum, Michael C. Zody, Jennifer Baldwin, Keri Devon, et al. 2001. "Initial sequencing and analysis of the human genome." *Nature* 409 (6822): 860–921. https://doi.org/10.1038/35057062.

Langer-Safer, P. R., M. Levine, and D. C. Ward. 1982. "Immunological method for mapping genes on Drosophila polytene chromosomes." *Proceedings of the National Academy of Sciences* 79 (14): 4381–5. https://doi.org/10.1073/pnas.79.14.4381.

Larsson, Chatarina, Ida Grundberg, Ola Söderberg, and Mats Nilsson. 2010. "In situ detection and genotyping of individual mRNA molecules." *Nature Methods* 7 (5): 395–97. https://doi.org/10.1038/nmeth.1448.

Lebrigand, Kevin, Joseph Bergenstråhle, Kim Thrane, Annelie Mollbrink, Pascal Barbry, Rainer Waldmann, and Joakim Lundeberg. 2020. "The spatial landscape of gene expression isoforms in tissue sections." *bioRxiv*, August, 2020.08.24.252296. https://doi.org/10.1101/2020.08.24.252296.

Lee, Hower, Sergio Marco Salas, Daniel Gyllborg, and Mats Nilsson. 2020. "Direct RNA targeted transcriptomic profiling in tissue using Hybridization-based RNA In Situ Sequencing (HybRISS)." *bioRxiv*, January, 2020.12.02.408781. https://doi.org/10.1101/2020.12.02.408781.

Lee, Je Hyuk, Evan R. Daugharthy, Jonathan Scheiman, Reza Kalhor, Thomas C. Ferrante, Richard Terry, Brian M. Turczyk, et al. 2015. "Fluorescent in situ sequencing (FISSEQ) of RNA for gene expression profiling in intact cells and tissues." *Nature Protocols* 10 (3): 442–58. https://doi.org/10.1038/nprot.2014.191.

Lee, Je Hyuk, Evan R Daugharthy, Jonathan Scheiman, Reza Kalhor, Joyce L Yang, Thomas C Ferrante, Richard Terry, et al. 2014. "Highly Multiplexed Subcellular RNA Sequencing in Situ." *Science* 343 (6177): 1360 LP–1363. https://doi.org/10.1126/science.1250212.

Leighton, Philip A., Kevin J. Mitchell, Lisa V. Goodrich, Xiaowei Lu, Kathy Pinson, Paul Scherz, William C. Skarnes, and Marc Tessier-Lavigne. 2001. "Defining brain wiring patterns and mechanisms through gene trapping in mice." *Nature* 410 (6825): 174–79. https://doi.org/10.1038/35065539.

Lein, Ed, Lars E. Borm, and Sten Linnarsson. 2017. "The promise of spatial transcriptomics for neuroscience in the era of molecular cell typing." https://doi.org/10.1126/science.aan6827.

Lein, Ed S. 2004. "Defining a Molecular Atlas of the Hippocampus Using DNA Microarrays and High-Throughput In Situ Hybridization." *Journal of Neuroscience* 24 (15): 3879–89. https://doi.org/10.1523/JNEUROSCI.4710-03.2004.

Lein, Ed S., Michael J. Hawrylycz, Nancy Ao, Mikael Ayres, Amy Bensinger, Amy Bernard, Andrew F. Boe, et al. 2007. "Genome-wide atlas of gene expression in the adult mouse brain." *Nature* 445 (7124): 168–76. https://doi.org/10.1038/nature05453.

Levsky, Jeffrey M. 2002. "Single-Cell Gene Expression Profiling." *Science* 297 (5582): 836–40. https://doi.org/10.1126/science.1072241.

Levy-Jurgenson, Alona, Xavier Tekpli, Vessela N Kristensen, and Zohar Yakhini. 2020. "Spatial transcriptomics inferred from pathology whole-slide images links tumor heterogeneity to survival in breast and lung cancer." *Scientific Reports* 10 (1): 18802. https://doi.org/10.1038/s41598-020-75708-z.

Lécuyer, Eric, Hideki Yoshida, Neela Parthasarathy, Christina Alm, Tomas Babak, Tanja Cerovina, Timothy R. Hughes, Pavel Tomancak, and Henry M. Krause. 2007. "Global Analysis of mRNA Localization Reveals a Prominent Role in Organizing Cellular Architecture and Function." *Cell* 131 (1): 174–87. https://doi.org/10.1016/j.cell.2007.08.003.

Li, Junxiang, Haofei Luo, Rui Wang, Jidong Lang, Siyu Zhu, Zhenming Zhang, Jianhuo Fang, et al. 2016. "Systematic Reconstruction of Molecular Cascades Regulating GP Development Using Single-Cell RNA-Seq." *Cell Reports* 15 (7): 1467–80. https://doi.org/10.1016/j.celrep.2016.04.043.

Li, Qiang, Zuwan Lin, Ren Liu, Xin Tang, Jiahao Huang, Yichun He, Haowen Zhou, et al. 2021. "<em>In situ</em> electro-sequencing in three-dimensional tissues." *bioRxiv*, January, 2021.04.22.440941. https://doi.org/10.1101/2021.04.22.440941.

Li, Ying-Xin, Shuiwang Ji, Sudhir Kumar, Jieping Ye, and Zhi-Hua Zhou. 2009. "Drosophila Gene Expression Pattern Annotation through Multi-Instance Multi-Label Learning." *IJCAI : Proceedings of the Conference* 2009 (January): 1445–50. http://www.ncbi.nlm.nih.gov/pubmed/20824158%20http://www.pubmedcentral.nih.gov/articlerender.fcgi?artid=PMC2932460.

Li, Yujie, Hanbo Chen, Xi Jiang, Xiang Li, Jinglei Lv, Hanchuan Peng, Joe Z. Tsien, and Tianming Liu. 2017. "Discover mouse gene coexpression landscapes using dictionary learning and sparse coding." *Brain Structure and Function* 222 (9): 4253–70. https://doi.org/10.1007/s00429-017-1460-9.

Li, Zhuliu, Tianci Song, Jeongsik Yong, and Rui Kuang. 2020. "Imputation of Spatially-resolved Transcriptomes by Graph-regularized Tensor Completion." *bioRxiv*, January, 2020.08.05.237560. https://doi.org/10.1101/2020.08.05.237560.

Liang, P, and A B Pardee. 1992. "Differential display of eukaryotic messenger RNA by means of the polymerase chain reaction." *Science* 257 (5072): 967 LP–971. https://doi.org/10.1126/science.1354393.

Liao, Jie, Xiaoyan Lu, Xin Shao, Ling Zhu, and Xiaohui Fan. 2020. "Uncovering an Organ's Molecular Architecture at Single-Cell Resolution by Spatially Resolved Transcriptomics." *Trends in Biotechnology*, June. https://doi.org/10.1016/j.tibtech.2020.05.006.

Lignell, Antti, Laura Kerosuo, Sebastian J. Streichan, Long Cai, and Marianne E. Bronner. 2017. "Identification of a neural crest stem cell niche by Spatial Genomic Analysis." *Nature Communications* 8 (1): 1830. https://doi.org/10.1038/s41467-017-01561-w.

Lis, John T., Jeffrey A. Simon, and Claudia A. Sutton. 1983. "New heat shock puffs and $\beta$=galactosidase activity resulting from transformation of Drosophila with an hsp70-lacZ hybrid gene." *Cell* 35 (2): 403–10. https://doi.org/10.1016/0092-8674(83)90173-3.

Liscovitch, Noa, Uri Shalit, and Gal Chechik. 2013. "FuncISH: learning a functional representation of neural ISH images." *Bioinformatics* 29 (13): i36–i43. https://doi.org/10.1093/bioinformatics/btt207.

Littman, Russell, Zachary Hemminger, Robert Foreman, Douglas Arneson, Guanglin Zhang, Fernando Gómez-Pinilla, Xia Yang, and Roy Wollman. 2020. "JSTA: joint cell segmentation and cell type annotation for spatial transcriptomics." *bioRxiv*, January, 2020.09.18.304147. https://doi.org/10.1101/2020.09.18.304147.

Liu, Lin, Yinhu Li, Siliang Li, Ni Hu, Yimin He, Ray Pong, Danni Lin, Lihua Lu, and Maggie Law. 2012. "Comparison of Next-Generation Sequencing Systems." Edited by P J Oefner. *Journal of Biomedicine and Biotechnology* 2012: 251364. https://doi.org/10.1155/2012/251364.

Liu, Miao, Yanfang Lu, Bing Yang, Yanbo Chen, Jonathan S. D. Radda, Mengwei Hu, Samuel G. Katz, and Siyuan Wang. 2020. "Multiplexed imaging of nucleome architectures in single cells of mammalian tissue." *Nature Communications* 11 (1): 1–14. https://doi.org/10.1038/s41467-020-16732-5.

Liu, Yang, Mingyu Yang, Yanxiang Deng, Graham Su, Archibald Enninful, Cindy C Guo, Toma Tebaldi, et al. 2020. "High-Spatial-Resolution Multi-Omics Sequencing via Deterministic Barcoding in Tissue." *Cell* 183 (6): 1665–1681.e18. https://doi.org/https://doi.org/10.1016/j.cell.2020.10.026.

Liu, Zheng, S. Frank Yan, John R. Walker, Theresa A. Zwingman, Tao Jiang, Jing Li, and Yingyao Zhou. 2007. "Study of gene function based on spatial co-expression in a high-resolution mouse brain atlas." *BMC Systems Biology* 1 (1): 19. https://doi.org/10.1186/1752-0509-1-19.

Lizardi, Paul M., Xiaohua Huang, Zhengrong Zhu, Patricia Bray-Ward, David C. Thomas, and David C. Ward. 1998. "Mutation detection and single-molecule counting using isothermal rolling-circle amplification." *Nature Genetics* 19 (3): 225–32. https://doi.org/10.1038/898.

Lohoff, T, S Ghazanfar, A Missarova, N Koulena, N Pierson, J A Griffiths, E S Bardot, et al. 2020. "Highly multiplexed spatially resolved gene expression profiling of mouse organogenesis." *bioRxiv*, January, 2020.11.20.391896. https://doi.org/10.1101/2020.11.20.391896.

Long, W, T Li, Y Yang, and H -B. Shen. 2021. "FlyIT: Drosophila Embryogenesis Image Annotation based on Image Tiling and Convolutional Neural Networks." *IEEE/ACM Transactions on Computational Biology and Bioinformatics* 18 (1): 194–204. https://doi.org/10.1109/TCBB.2019.2935723.

Long, Xi, Jennifer Colonell, Allan M Wong, Robert H Singer, and Timothée Lionnet. 2017. "Quantitative mRNA imaging throughout the entire Drosophila brain." *Nature Methods* 14 (7): 703–6. https://doi.org/10.1038/nmeth.4309.

Lopez, Romain, Achille Nazaret, Maxime Langevin, Jules Samaran, Jeffrey Regier, Michael I. Jordan, and Nir Yosef. 2019. "A joint model of unpaired data from scRNA-seq and spatial transcriptomics for imputing missing gene expression measurements," May. http://arxiv.org/abs/1905.02269.

Lovatt, Ditte, Brittani K. Ruble, Jaehee Lee, Hannah Dueck, Tae Kyung Kim, Stephen Fisher, Chantal Francis, et al. 2014. "Transcriptome in vivo analysis (TIVA) of spatially defined single cells in live tissue." *Nature Methods* 11 (2): 190–96. https://doi.org/10.1038/nmeth.2804.

Lovell, Peter V., Morgan Wirthlin, Taylor Kaser, Alexa A. Buckner, Julia B. Carleton, Brian R. Snider, Anne K. McHugh, Alexander Tolpygo, Partha P. Mitra, and Claudio V. Mello. 2020. "ZEBrA: Zebra finch Expression Brain Atlas—A resource for comparative molecular neuroanatomy and brain evolution studies." *Journal of Comparative Neurology* 528 (12): 2099–2131. https://doi.org/10.1002/cne.24879.

Lowe, David G. 2004. "Distinctive Image Features from Scale-Invariant Keypoints." *International Journal of Computer Vision* 60 (2): 91–110. https://doi.org/10.1023/B:VISI.0000029664.99615.94.

Lubeck, Eric, and Long Cai. 2012. "Single-cell systems biology by super-resolution imaging and combinatorial labeling." *Nature Methods 2012 9:7* 9 (7): 743–48. https://doi.org/10.1038/nmeth.2069.

Lubeck, Eric, Ahmet F. Coskun, Timur Zhiyentayev, Mubhij Ahmad, and Long Cai. 2014. "Single-cell in situ RNA profiling by sequential hybridization." Nature Publishing Group. https://doi.org/10.1038/nmeth.2892.

Luengo Hendriks, Cris L., Soile V. E. Keränen, Charless C. Fowlkes, Lisa Simirenko, Gunther H. Weber, Angela H. DePace, Clara Henriquez, et al. 2006. "Three-dimensional morphology and gene expression in the Drosophila blastoderm at cellular resolution I: Data acquisition pipeline." *Genome Biology* 7 (12): R123. https://doi.org/10.1186/gb-2006-7-12-r123.

Lukan, Tjaša, Maruša Pompe-Novak, Špela Baebler, Magda Tušek-Žnidarič, Aleš Kladnik, Maja Križnik, Andrej Blejec, et al. 2020. "Precision transcriptomics of viral foci reveals the spatial regulation of immune-signaling genes and identifies RBOHD as an important player in the incompatible interaction between potato virus Y and potato." *The Plant Journal* 104 (3): 645–61. https://doi.org/10.1111/tpj.14953.

Lundberg, Emma, and Georg H. H. Borner. 2019. "Spatial proteomics: a powerful discovery tool for cell biology." *Nature Reviews Molecular Cell Biology* 20 (5): 285–302. https://doi.org/10.1038/s41580-018-0094-y.

Lundmark, Anna, Natalija Gerasimcik, Tove Båge, Anders Jemt, Annelie Mollbrink, Fredrik Salmén, Joakim Lundeberg, and Tülay Yucel-Lindberg. 2018. "Gene expression profiling of periodontitis-affected gingival tissue by spatial transcriptomics." *Scientific Reports* 8 (1): 9370. https://doi.org/10.1038/s41598-018-27627-3.

Luo, Lin, Ranelle C. Salunga, Hongqing Guo, Anton Bittner, K. C. Joy, Jose E. Galindo, Huinian Xiao, et al. 1999. "Gene expression profiles of laser-captured adjacent neuronal subtypes." *Nature Medicine* 5 (1): 117–22. https://doi.org/10.1038/4806.

Lynch, Andrew S., David Briggs, and Ian A. Hope. 1995. "Developmental expression pattern screen for genes predicted in the C. elegans genome sequencing project." *Nature Genetics* 11 (3): 309–13. https://doi.org/10.1038/ng1195-309.

Maaskola, Jonas, Ludvig Bergenstråhle, Aleksandra Jurek, José Fernández Navarro, Jens Lagergren, and Joakim Lundeberg. 2018. "Charting Tissue Expression Anatomy by Spatial Transcriptome Decomposition." *bioRxiv*, December, 362624. https://doi.org/10.1101/362624.

Macevicz, Stephen. 1995. "DNA sequencing by parallel olgonucleotide extensions." https://patents.google.com/patent/US5969119A/en.

Macosko, Evan Z., Anindita Basu, Rahul Satija, James Nemesh, Karthik Shekhar, Melissa Goldman, Itay Tirosh, et al. 2015. "Highly parallel genome-wide expression profiling of individual cells using nanoliter droplets." *Cell* 161 (5): 1202–14. https://doi.org/10.1016/j.cell.2015.05.002.

Manco, Rita, Inna Averbukh, Ziv Porat, Keren Bahar Halpern, Ido Amit, and Shalev Itzkovitz. 2020. "Clump sequencing exposes the spatial expression programs of intestinal secretory cells." *bioRxiv*, August, 2020.08.05.237917. https://doi.org/10.1101/2020.08.05.237917.

Maniatis, Silas, Tarmo Äijö, Sanja Vickovic, Catherine Braine, Kristy Kang, Annelie Mollbrink, Delphine Fagegaltier, et al. 2019. "Spatiotemporal dynamics of molecular pathology in amyotrophic lateral sclerosis." *Science* 364 (6435): 89 LP–93. https://doi.org/10.1126/science.aav9776.

Manno, Gioele La, Kimberly Siletti, Alessandro Furlan, Daniel Gyllborg, Elin Vinsland, Christoffer Mattsson Langseth, Irina Khven, et al. 2020. "Molecular architecture of the developing mouse brain." *bioRxiv*, July, 2020.07.02.184051. https://doi.org/10.1101/2020.07.02.184051.

Mantri, Madhav, Gaetano J Scuderi, Roozbeh Abedini Nassab, Michael F Z Wang, David McKellar, Jonathan T Butcher, and Iwijn De Vlaminck. 2020. "Spatiotemporal single-cell RNA sequencing of developing hearts reveals interplay between cellular differentiation and morphogenesis." *bioRxiv*, January, 2020.05.03.065102. https://doi.org/10.1101/2020.05.03.065102.

Margaroli, C, P Benson, N S Sharma, M C Madison, S W Robison, N Arora, K Ton, et al. 2021. "Spatial mapping of SARS-CoV-2 and H1N1 Lung Injury Identifies Differential Transcriptional Signatures." *Cell Reports. Medicine*, March, 100242. https://doi.org/10.1016/j.xcrm.2021.100242.

Marquart, Gregory D., Kathryn M. Tabor, Mary Brown, Jennifer L. Strykowski, Gaurav K. Varshney, Matthew C. LaFave, Thomas Mueller, Shawn M. Burgess, Shin-ichi Higashijima, and Harold A. Burgess. 2015. "A 3D Searchable Database of Transgenic Zebrafish Gal4 and Cre Lines for Functional Neuroanatomy Studies." *Frontiers in Neural Circuits* 9 (November): 1–17. https://doi.org/10.3389/fncir.2015.00078.

Martinelli, Sylvia D., Clive G. Brown, and Richard Durbin. 1997. "Gene expression and development databases forC. elegans." *Seminars in Cell & Developmental Biology* 8 (5): 459–67. https://doi.org/10.1006/scdb.1997.0171.

Maynard, Kristen R, Madhavi Tippani, Yoichiro Takahashi, BaDoi N Phan, Thomas M Hyde, Andrew E Jaffe, and Keri Martinowich. 2020. "dotdotdot: an automated approach to quantify multiplex single molecule fluorescent in situ hybridization (smFISH) images in complex tissues." *Nucleic Acids Research* 48 (11): e66–e66. https://doi.org/10.1093/nar/gkaa312.

McKee, Adrienne E., Emmanuel Minet, Charlene Stern, Shervin Riahi, Charles D. Stiles, and Pamela A. Silver. 2005. "A genome-wide in situ hybridization map of RNA-binding proteins reveals anatomically restricted expression in the developing mouse brain." *BMC Developmental Biology* 5 (1): 14. https://doi.org/10.1186/1471-213X-5-14.

Medaglia, Chiara, Amir Giladi, Liat Stoler-Barak, Marco De Giovanni, Tomer Meir Salame, Adi Biram, Eyal David, et al. 2017. "Spatial reconstruction of immune niches by combining photoactivatable reporters and scRNA-seq." *Science* 358 (6370): 1622–6. https://doi.org/10.1126/science.aao4277.

Meier-Ruge, W., W. Bielser, E. Remy, F. Hillenkamp, R. Nitsche, and R. Uns ld. 1976. "The laser in the Lowry technique for microdissection of freeze-dried tissue slices." *The Histochemical Journal* 8 (4): 387–401. https://doi.org/10.1007/BF01003828.

Melsted, Páll, A Sina Booeshaghi, Lauren Liu, Fan Gao, Lambda Lu, Kyung Hoi (Joseph) Min, Eduardo da Veiga Beltrame, Kristján Eldjárn Hjörleifsson, Jase Gehring, and Lior Pachter. 2021. "Modular, efficient and constant-memory single-cell RNA-seq preprocessing." *Nature Biotechnology*. https://doi.org/10.1038/s41587-021-00870-2.

Merritt, Christopher R., Giang T. Ong, Sarah Church, Kristi Barker, Gary Geiss, Margaret Hoang, Jaemyeong Jung, et al. 2019. "High multiplex, digital spatial profiling of proteins and RNA in fixed tissue using genomic detection methods." *bioRxiv* 38 (May): 559021. https://doi.org/10.1101/559021.

Middleton, Sarah A, James Eberwine, and Junhyong Kim. 2019. "Comprehensive catalog of dendritically localized mRNA isoforms from sub-cellular sequencing of single mouse neurons." *BMC Biology* 17 (1): 5. https://doi.org/10.1186/s12915-019-0630-z.

Miller, Jeremy A., Song-Lin Ding, Susan M. Sunkin, Kimberly A. Smith, Lydia Ng, Aaron Szafer, Amanda Ebbert, et al. 2014. "Transcriptional landscape of the prenatal human brain." *Nature* 508 (7495): 199–206. https://doi.org/10.1038/nature13185.

Moffitt, Jeffrey R., Dhananjay Bambah-Mukku, Stephen W. Eichhorn, Eric Vaughn, Karthik Shekhar, Julio D. Perez, Nimrod D. Rubinstein, et al. 2018. "Molecular, spatial, and functional single-cell profiling of the hypothalamic preoptic region." *Science*. https://doi.org/10.1126/science.aau5324.

Moffitt, Jeffrey R., Junjie Hao, Guiping Wang, Kok Hao Chen, Hazen P. Babcock, and Xiaowei Zhuang. 2016. "High-throughput single-cell gene-expression profiling with multiplexed error-robust fluorescence in situ hybridization." *Proceedings of the National Academy of Sciences of the United States of America*. https://doi.org/10.1073/pnas.1612826113.

Moncada, Reuben, Dalia Barkley, Florian Wagner, Marta Chiodin, Joseph C Devlin, Maayan Baron, Cristina H Hajdu, Diane M Simeone, and Itai Yanai. 2020. "Integrating microarray-based spatial transcriptomics and single-cell RNA-seq reveals tissue architecture in pancreatic ductal adenocarcinomas." *Nature Biotechnology* 38 (3): 333–42. https://doi.org/10.1038/s41587-019-0392-8.

Moor, Andreas E., Matan Golan, Efi E. Massasa, Doron Lemze, Tomer Weizman, Rom Shenhav, Shaked Baydatch, et al. 2017. "Global mRNA polarization regulates translation efficiency in the intestinal epithelium." *Science* 357 (6357): 1299–1303. https://doi.org/10.1126/science.aan2399.

Moor, Andreas E., Yotam Harnik, Shani Ben-Moshe, Efi E. Massasa, Milena Rozenberg, Raya Eilam, Keren Bahar Halpern, and Shalev Itzkovitz. 2018. "Spatial Reconstruction of Single Enterocytes Uncovers Broad Zonation along the Intestinal Villus Axis." *Cell* 175 (4): 1156–1167.e15. https://doi.org/10.1016/j.cell.2018.08.063.

Moor, Andreas E., and Shalev Itzkovitz. 2017. "Spatial transcriptomics: paving the way for tissue-level systems biology." *Current Opinion in Biotechnology* 46 (August): 126–33. https://doi.org/10.1016/j.copbio.2017.02.004.

Mori, Tomoya, Haruka Takaoka, Junko Yamane, Cantas Alev, and Wataru Fujibuchi. 2019. "Novel computational model of gastrula morphogenesis to identify spatial discriminator genes by self-organizing map (SOM) clustering." *Scientific Reports* 9 (1): 1–10. https://doi.org/10.1038/s41598-019-49031-1.

Mori, Tomoya, Junko Yamane, Kenta Kobayashi, Nobuko Taniyama, Takanori Tano, and Wataru Fujibuchi. 2017. "Development of 3D Tissue Reconstruction Method from Single-cell RNA-seq Data." *Genomics and Computational Biology; Vol 3 No 1 (2017).* https://doi.org/10.18547/gcb.2017.vol3.iss1.e53.

Moris, Naomi, Kerim Anlas, Susanne C. van den Brink, Anna Alemany, Julia Schröder, Sabitri Ghimire, Tina Balayo, Alexander van Oudenaarden, and Alfonso Martinez Arias. 2020. "An in vitro model of early anteroposterior organization during human development." *Nature* 582 (7812): 410–15. https://doi.org/10.1038/s41586-020-2383-9.

Morton, Matthew L., Xiaodong Bai, Callie R. Merry, Philip A. Linden, Ahmad M. Khalil, Rom S. Leidner, and Cheryl L. Thompson. 2014. "Identification of mRNAs and lincRNAs associated with lung cancer progression using next-generation RNA sequencing from laser micro-dissected archival FFPE tissue specimens." *Lung Cancer* 85 (1): 31–39. https://doi.org/10.1016/j.lungcan.2014.03.020.

Myers, Eugene W. 2000. "A Whole-Genome Assembly of Drosophila." *Science* 287 (5461): 2196–2204. https://doi.org/10.1126/science.287.5461.2196.

Nagalakshmi, Ugrappa, Zhong Wang, Karl Waern, Chong Shou, Debasish Raha, Mark Gerstein, and Michael Snyder. 2008. "The Transcriptional Landscape of the Yeast Genome Defined by RNA Sequencing." *Science* 320 (5881): 1344–9. https://doi.org/10.1126/science.1158441.

Nagarajan, Maxwell B, Augusto M Tentori, Wen Cai Zhang, Frank J Slack, and Patrick S Doyle. 2020. "Spatially resolved and multiplexed MicroRNA quantification from tissue using nanoliter well arrays." *Microsystems & Nanoengineering* 6 (1): 51. https://doi.org/10.1038/s41378-020-0169-8.

Nakamura, Toru, Yoichi Furukawa, Hidewaki Nakagawa, Tatsuhiko Tsunoda, Hiroaki Ohigashi, Kohei Murata, Osamu Ishikawa, et al. 2004. "Genome-wide cDNA microarray analysis of gene expression profiles in pancreatic cancers using populations of tumor cells and normal ductal epithelial cells selected for purity by laser microdissection." *Oncogene* 23 (13): 2385–2400. https://doi.org/10.1038/sj.onc.1207392.

Nakayama, Naomi, Juana M. Arroyo, Joseph Simorowski, Bruce May, Robert Martienssen, and Vivian F. Irish. 2005. "Gene Trap Lines Define Domains of Gene Regulation in Arabidopsis Petals and Stamens." *The Plant Cell* 17 (9): 2486–2506. https://doi.org/10.1105/tpc.105.033985.

Navarro, José Fernández, Joel Sjöstrand, Fredrik Salmén, Joakim Lundeberg, and Patrik L. Ståhl. 2017. "ST Pipeline: an automated pipeline for spatial mapping of unique transcripts." *Bioinformatics (Oxford, England)*. https://doi.org/10.1093/bioinformatics/btx211.

Nederlof, P. M., S. van der Flier, J. Wiegant, A. K. Raap, H. J. Tanke, J. S. Ploem, and M. van der Ploeg. 1990. "Multiple fluorescence in situ hybridization." *Cytometry* 11 (1): 126–31. https://doi.org/10.1002/cyto.990110115.

Neubacher, Saskia, and Christoph Arenz. 2009. "Rolling-Circle Amplification: Unshared Advantages in miRNA Detection." *ChemBioChem* 10 (8): 1289–91. https://doi.org/10.1002/cbic.200900116.

Nichterwitz, Susanne, Geng Chen, Julio Aguila Benitez, Marlene Yilmaz, Helena Storvall, Ming Cao, Rickard Sandberg, Qiaolin Deng, and Eva Hedlund. 2016. "Laser capture microscopy coupled with Smart-seq2 for precise spatial transcriptomic profiling." *Nature Communications* 7 (1): 12139. https://doi.org/10.1038/ncomms12139.

Nilsson, Mats, Helena Malmgren, Martina Samiotaki, Marek Kwiatkowski, B. Chowdhary, and Ulf Landegren. 1994. "Padlock probes: circularizing oligonucleotides for localized DNA detection." *Science* 265 (5181): 2085–8. https://doi.org/10.1126/science.7522346.

Nitzan, Mor, Nikos Karaiskos, Nir Friedman, and Nikolaus Rajewsky. 2019. "Gene expression cartography." *Nature* 576 (7785): 132–37. https://doi.org/10.1038/s41586-019-1773-3.

Noto, Torben, Derrick Barnagian, and Jason B. Castro. 2017. "Genome-scale investigation of olfactory system spatial heterogeneity." Edited by Hiroaki Matsunami. *PLOS ONE* 12 (5): e0178087. https://doi.org/10.1371/journal.pone.0178087.

Ohyama, H., X. Zhang, Y. Kohno, I. Alevizos, M. Posner, D. T. Wong, and R. Todd. 2000. "Laser Capture Microdissection-Generated Target Sample for High-Density Oligonucleotide Array Hybridization." *BioTechniques* 29 (3): 530–36. https://doi.org/10.2144/00293st05.

Oka, Yuma, and Thomas N Sato. 2015. "Whole-mount single molecule FISH method for zebrafish embryo." *Scientific Reports* 5 (February): 8571. https://doi.org/10.1038/srep08571.

Okamura-Oho, Yuko, Kazuro Shimokawa, Satoko Takemoto, Asami Hirakiyama, Sakiko Nakamura, Yuki Tsujimura, Masaomi Nishimura, et al. 2012. "Transcriptome Tomography for Brain Analysis in the Web-Accessible Anatomical Space." Edited by Satoru Hayasaka. *PLoS ONE* 7 (9): e45373. https://doi.org/10.1371/journal.pone.0045373.

O'Kane, C. J., and W. J. Gehring. 1987. "Detection in situ of genomic regulatory elements in Drosophila." *Proceedings of the National Academy of Sciences* *of the United States of America* 84 (24): 9123–7. https://doi.org/10.1073/pnas.84.24.9123.

Ortiz, Cantin, Jose Fernandez Navarro, Aleksandra Jurek, Antje Märtin, Joakim Lundeberg, and Konstantinos Meletis. 2020. "Molecular atlas of the adult mouse brain." *Science Advances* 6 (26): eabb3446. https://doi.org/10.1126/sciadv.abb3446.

Palla, Giovanni, Hannah Spitzer, Michal Klein, David Fischer, Anna Christina Schaar, Louis Benedikt Kuemmerle, Sergei Rybakov, et al. 2021. "Squidpy: a scalable framework for spatial single cell analysis." *bioRxiv*, January, 2021.02.19.431994. https://doi.org/10.1101/2021.02.19.431994.

Pan, Jia-Yu, André Guilherme, Ribeiro Balan, Eric P. Xing, Agma Juci Machado Traina, and Christos Faloutsos. 2006. "Automatic mining of fruit fly embryo images." In *Proceedings of the 12th Acm Sigkdd International Conference on Knowledge Discovery and Data Mining - Kdd '06*, 2006:693. New York, New York, USA: ACM Press. https://doi.org/10.1145/1150402.1150489.

Park, Jeongbin, Wonyl Choi, Sebastian Tiesmeyer, Brian Long, Lars E. Borm, Emma Garren, Thuc Nghi Nguyen, et al. 2019. "Segmentation-free inference of cell types from in situ transcriptomics data." *bioRxiv*. https://doi.org/10.1101/800748.

Park, Jiwoon, Jonathan Foox, Tyler Hether, David Danko, Sarah Warren, Youngmi Kim, Jason Reeves, et al. 2021. "Systemic Tissue and Cellular Disruption from SARS-CoV-2 Infection revealed in COVID-19 Autopsies and Spatial Omics Tissue Maps." *bioRxiv*, January, 2021.03.08.434433. https://doi.org/10.1101/2021.03.08.434433.

Partel, Gabriele, Markus Hilscher, Giorgia Milli, Leslie Solorzano, Anna Klemm, Mats Nilsson, and Carolina Wählby. 2019. "Identification of spatial compartments in tissue from in situ sequencing data." *bioRxiv*, September, 765842. https://doi.org/10.1101/765842.

Partel, Gabriele, and Carolina Wählby. 2020. "Spage2vec: Unsupervised detection of spatial gene expression constellations." *bioRxiv*, February, 2020.02.12.945345. https://doi.org/10.1101/2020.02.12.945345.

Parthasarathy, Raghuveer. 2012. "Rapid, accurate particle tracking by calculation of radial symmetry centers." *Nature Methods* 9 (7): 724–26. https://doi.org/10.1038/nmeth.2071.

Patrushev, Ilya, Christina James-Zorn, Aldo Ciau-Uitz, Roger Patient, and Michael J. Gilchrist. 2018. "New methods for computational decomposition of whole-mount in situ images enable effective curation of a large, highly redundant collection of Xenopus images." *PLoS Computational Biology* 14 (8): e1006077. https://doi.org/10.1371/journal.pcbi.1006077.

Peng, Guangdun, Shengbao Suo, Jun Chen, Weiyang Chen, Chang Liu, Fang Yu, Ran Wang, et al. 2016. "Spatial Transcriptome for the Molecular Annotation of Lineage Fates and Cell Identity in Mid-gastrula Mouse Embryo." *Developmental Cell* 36 (6): 681–97. https://doi.org/10.1016/j.devcel.2016.02.020.

Pennington, Jeffrey, Richard Socher, and Christopher D Manning. n.d. "GloVe: Global Vectors for Word Representation."

Pettit, Jean-Baptiste, Raju Tomer, Kaia Achim, Sylvia Richardson, Lamiae Azizi, and John Marioni. 2014. "Identifying Cell Types from Spatially Referenced Single-Cell Expression Datasets." Edited by Quaid Morris. *PLoS Computational Biology* 10 (9): e1003824. https://doi.org/10.1371/journal.pcbi.1003824.

Petukhov, Viktor, Ruslan A Soldatov, Konstantin Khodosevich, and Peter V Kharchenko. 2020. "Bayesian segmentation of spatially resolved transcriptomics data." *bioRxiv*, January, 2020.10.05.326777. https://doi.org/10.1101/2020.10.05.326777.

Pérez-Martín, Fernando, Fernando J. Yuste-Lisbona, Benito Pineda, María Pilar Angarita-Díaz, Begoña García-Sogo, Teresa Antón, Sibilla Sánchez, et al. 2017. "A collection of enhancer trap insertional mutants for functional genomics in tomato." *Plant Biotechnology Journal* 15 (11): 1439–52. https://doi.org/10.1111/pbi.12728.

Pham, Duy, Xiao Tan, Jun Xu, Laura F Grice, Pui Yeng Lam, Arti Raghubar, Jana Vukovic, Marc J Ruitenberg, and Quan Nguyen. 2020. "stLearn: integrating spatial location, tissue morphology and gene expression to find cell types, cell-cell interactions and spatial trajectories within undissociated tissues." *bioRxiv*, January, 2020.05.31.125658. https://doi.org/10.1101/2020.05.31.125658.

Player, Audrey N., Lu Ping Shen, Daryn Kenny, Vincent P. Antao, and Janice A. Kolberg. 2001. "Single-copy gene detection using branched DNA (bDNA) in situ hybridization." *Journal of Histochemistry and Cytochemistry* 49 (5): 603–11. https://doi.org/10.1177/002215540104900507.

Plouhinec, Jean-Louis, Sofía Medina-Ruiz, Caroline Borday, Elsa Bernard, Jean-Philippe Vert, Michael B. Eisen, Richard M. Harland, and Anne H. Monsoro-Burq. 2017. "A molecular atlas of the developing ectoderm defines neural, neural crest, placode, and nonneural progenitor identity in vertebrates." Edited by James Briscoe. *PLOS Biology* 15 (10): e2004045. https://doi.org/10.1371/journal.pbio.2004045.

Pruteanu-Malinici, Iulian, Daniel L. Mace, and Uwe Ohler. 2011. "Automatic Annotation of Spatial Expression Patterns via Sparse Bayesian Factor Models." Edited by Joel S. Bader. *PLoS Computational Biology* 7 (7): e1002098. https://doi.org/10.1371/journal.pcbi.1002098.

Puchalski, Ralph B, Nameeta Shah, Jeremy Miller, Rachel Dalley, Steve R Nomura, Jae-Guen Yoon, Kimberly A Smith, et al. 2018. "An anatomic transcriptional atlas of human glioblastoma." *Science* 360 (6389): 660 LP–663. https://doi.org/10.1126/science.aaf2666.

Puniyani, Kriti, and Eric P. Xing. 2013. "GINI: From ISH Images to Gene Interaction Networks." Edited by Gal Chechik. *PLoS Computational Biology* 9 (10): e1003227. https://doi.org/10.1371/journal.pcbi.1003227.

Qian, Xiaoyan, Kenneth D. Harris, Thomas Hauling, Dimitris Nicoloutsopoulos, Ana B. Muñoz-Manchado, Nathan Skene, Jens Hjerling-Leffler, and Mats Nilsson. 2020. "Probabilistic cell typing enables fine mapping of closely related cell types in situ." *Nature Methods* 17 (1): 101–6. https://doi.org/10.1038/s41592-019-0631-4.

Raj, Arjun, Patrick van den Bogaard, Scott A. Rifkin, Alexander van Oudenaarden, and Sanjay Tyagi. 2008. "Imaging individual mRNA molecules using multiple singly labeled probes." *Nature Methods* 5 (10): 877–79. https://doi.org/10.1038/nmeth.1253.

Ramsköld, Daniel, Shujun Luo, Yu-Chieh Wang, Robin Li, Qiaolin Deng, Omid R Faridani, Gregory A Daniels, et al. 2012. "Full-length mRNA-Seq from single-cell levels of RNA and individual circulating tumor cells." *Nature Biotechnology* 30 (8): 777–82. https://doi.org/10.1038/nbt.2282.

Ren, Xianwen, Guojie Zhong, Qiming Zhang, Lei Zhang, Yujie Sun, and Zemin Zhang. 2020. "Reconstruction of cell spatial organization based on ligand-receptor mediated self-assembly." *bioRxiv*, January, 2020.02.13.948521. https://doi.org/10.1101/2020.02.13.948521.

Righelli, Dario, Lukas M Weber, Helena L Crowell, Brenda Pardo, Leonardo Collado-Torres, Shila Ghazanfar, Aaron T L Lun, Stephanie C Hicks, and Davide Risso. 2021. "SpatialExperiment: infrastructure for spatially resolved transcriptomics data in R using Bioconductor." *bioRxiv*, January, 2021.01.27.428431. https://doi.org/10.1101/2021.01.27.428431.

Ringwald, Martin, Richard Baldock, Jonathan Bard, Matthew Kaufman, Janan T. Eppig, Joel E. Richardson, Joseph H. Nadeau, and Duncan Davidson. 1994. "A database for mouse development." *Science* 265 (5181): 2033–4. https://doi.org/10.1126/science.8091224.

Ringwald, Martin, Geoffrey L Davis, Alex G Smith, Laura E Trepanier, Dale A Begley, Joel E Richardson, and Janan T Eppig. 1997. "The mouse gene expression database GXD." *Seminars in Cell & Developmental Biology* 8 (5): 489–97. https://doi.org/https://doi.org/10.1006/scdb.1997.0177.

Ringwald, Martin, Mary E Mangan, Janan T Eppig, James A Kadin, and Joel E Richardson. 1999. "GXD: a Gene Expression Database for the laboratory mouse." *Nucleic Acids Research* 27 (1): 106–12. https://doi.org/10.1093/nar/27.1.106.

Rizvi, Abbas H, Pablo G Camara, Elena K Kandror, Thomas J Roberts, Ira Schieren, Tom Maniatis, and Raul Rabadan. 2017. "Single-cell topological

RNA-seq analysis reveals insights into cellular differentiation and development." *Nature Biotechnology* 35 (6): 551–60. https://doi.org/10.1038/nbt.3854.

Roberts, Kenny, Alexander Aivazidis, Vitalii Kleshchevnikov, Tong Li, Robin Fropf, Michael Rhodes, Joseph M Beechem, Martin Hemberg, and Omer Ali Bayraktar. 2021. "Transcriptome-wide spatial RNA profiling maps the cellular architecture of the developing human neocortex." *bioRxiv*, January, 2021.03.20.436265. https://doi.org/10.1101/2021.03.20.436265.

Roberts, Margaret E., Brandon M. Stewart, and Dustin Tingley. 2019. "Stm: An R package for structural topic models." *Journal of Statistical Software* 91. https://doi.org/10.18637/jss.v091.i02.

Rodriques, Samuel G., Robert R. Stickels, Aleksandrina Goeva, Carly A. Martin, Evan Murray, Charles R. Vanderburg, Joshua Welch, Linlin M. Chen, Fei Chen, and Evan Z. Macosko. 2019. "Slide-seq: A scalable technology for measuring genome-wide expression at high spatial resolution." *Science*. https://doi.org/10.1126/science.aaw1219.

Ronneberger, Olaf, Philipp Fischer, and Thomas Brox. 2015. "U-Net: Convolutional Networks for Biomedical Image Segmentation BT - Medical Image Computing and Computer-Assisted Intervention – MICCAI 2015." In, edited by Nassir Navab, Joachim Hornegger, William M Wells, and Alejandro F Frangi, 234–41. Cham: Springer International Publishing.

Rosen, Barry, and Rosa S. P. Beddington. 1993. "Whole-mount in situ hybridization in the mouse embryo: gene expression in three dimensions." *Trends in Genetics* 9 (5): 162–67. https://doi.org/10.1016/0168-9525(93)90162-B.

Rödelsperger, Christian, Annabel Ebbing, Devansh Raj Sharma, Misako Okumura, Ralf J Sommer, and Hendrik C Korswagen. 2020. "Spatial transcriptomics of nematodes identifies sperm cells as a source of genomic novelty and rapid evolution." *Molecular Biology and Evolution*, August. https://doi.org/10.1093/molbev/msaa207.

Saelens, Wouter, Robrecht Cannoodt, Helena Todorov, and Yvan Saeys. 2019. "A comparison of single-cell trajectory inference methods." *Nature Biotechnology* 37 (5): 547–54. https://doi.org/10.1038/s41587-019-0071-9.

Salgado, David, Gregory Gimenez, François Coulier, and Christophe Marcelle. 2008. "COMPARE, a multi-organism system for cross-species data comparison and transfer of information." *Bioinformatics* 24 (3): 447–49. https://doi.org/10.1093/bioinformatics/btm599.

Samacoits, Aubin, Racha Chouaib, Adham Safieddine, Abdel-Meneem Traboulsi, Wei Ouyang, Christophe Zimmer, Marion Peter, Edouard Bertrand, Thomas Walter, and Florian Mueller. 2018. "A computational framework to study sub-cellular RNA localization." *Nature Communications* 9 (1): 4584. https://doi.org/10.1038/s41467-018-06868-w.

Satija, Rahul, Jeffrey A. Farrell, David Gennert, Alexander F. Schier, and Aviv Regev. 2015. "Spatial reconstruction of single-cell gene expression data." *Nature Biotechnology* 33 (5): 495–502. https://doi.org/10.1038/nbt.3192.

Satou, Yutaka, Naohito Takatori, Lixy Yamada, Yasuaki Mochizuki, Makoto Hamaguchi, Hisayoshi Ishikawa, Shota Chiba, et al. 2001. "Gene expression profiles in <em>Ciona intestinalis</em> tailbud embryos." *Development* 128 (15): 2893 LP–2904. http://dev.biologists.org/content/128/15/2893.abstract.

Saviano, Antonio, Neil C. Henderson, and Thomas F. Baumert. 2020. "Single-cell genomics and spatial transcriptomics: Discovery of novel cell states and cellular interactions in liver physiology and disease biology." *Journal of Hepatology* 73 (5): 1219–30. https://doi.org/10.1016/j.jhep.2020.06.004.

Schede, Halima Hannah, Christian Gabriel Schneider, Johanna Stergiadou, Lars E Borm, Anurag Ranjak, Tracy M Yamawaki, Fabrice P A David, et al. 2020. "Spatial tissue profiling by imaging-free molecular tomography." *bioRxiv*, August, 2020.08.04.235655. https://doi.org/10.1101/2020.08.04.235655.

Schwaber, Jessica, Stacey Andersen, and Lars Nielsen. 2019. "Shedding light: The importance of reverse transcription efficiency standards in data interpretation." *Biomolecular Detection and Quantification* 17 (March): 100077. https://doi.org/10.1016/j.bdq.2018.12.002.

Seydoux, G, and A Fire. 1994. "Soma-germline asymmetry in the distributions of embryonic RNAs in Caenorhabditis elegans." *Development* 120 (10): 2823 LP–2834. http://dev.biologists.org/content/120/10/2823.abstract.

Sgroi, Dennis C, Sarena Teng, Greg Robinson, Rebbecca LeVangie, James R Hudson, and Abdel G Elkahloun. 1999. "<em>In Vivo</em> Gene Expression Profile Analysis of Human Breast Cancer Progression." *Cancer Research* 59 (22): 5656 LP–5661. http://cancerres.aacrjournals.org/content/59/22/5656.abstract.

Shah, Sheel, Eric Lubeck, Wen Zhou, and Long Cai. 2016. "In Situ Transcription Profiling of Single Cells Reveals Spatial Organization of Cells in the Mouse Hippocampus." *Neuron*. https://doi.org/10.1016/j.neuron.2016.10.001.

Shah, Sheel, Yodai Takei, Wen Zhou, Eric Lubeck, Jina Yun, Chee Huat Linus Eng, Noushin Koulena, et al. 2018. "Dynamics and Spatial Genomics of the Nascent Transcriptome by Intron seqFISH." *Cell*. https://doi.org/10.1016/j.cell.2018.05.035.

Sharma, Ankur, Justine Jia Wen Seow, Charles-Antoine Dutertre, Rhea Pai, Camille Blériot, Archita Mishra, Regina Men Men Wong, et al. 2020. "Onco-fetal Reprogramming of Endothelial Cells Drives Immunosuppressive Macrophages in Hepatocellular Carcinoma." *Cell* 183 (2): 377–394.e21. https://doi.org/10.1016/j.cell.2020.08.040.

Shcherbatyy, Volodymyr, James Carson, Murat Yaylaoglu, Katharina Jäckle, Frauke Grabbe, Maren Brockmeyer, Halenur Yavuz, and Gregor Eichele. 2015. "A Digital Atlas of Ion Channel Expression Patterns in the Two-Week-Old Rat Brain." *Neuroinformatics* 13 (1): 111–25. https://doi.org/10.1007/s12021-014-9247-0.

Shendure, Jay. 2005. "Accurate Multiplex Polony Sequencing of an Evolved Bacterial Genome." *Science* 309 (5741): 1728–32. https://doi.org/10.1126/science.1117389.

Siegert, Sandra, Brigitte Gross Scherf, Karina Del Punta, Nick Didkovsky, Nathaniel Heintz, and Botond Roska. 2009. "Genetic address book for retinal cell types." *Nature Neuroscience* 12 (9): 1197–1204. https://doi.org/10.1038/nn.2370.

Singer, R. H., and D. C. Ward. 1982. "Actin gene expression visualized in chicken muscle tissue culture by using in situ hybridization with a biotinated nucleotide analog." *Proceedings of the National Academy of Sciences of the United States of America* 79 (23 I): 7331–5. https://doi.org/10.1073/pnas.79.23.7331.

Singh, Ram P., Vanessa M. Brown, Abhijit Chaudhari, Arshad H. Khan, Alex Ossadtchi, Daniel M. Sforza, A.Ken Meadors, Simon R. Cherry, Richard M. Leahy, and Desmond J. Smith. 2003. "High-resolution voxelation mapping of human and rodent brain gene expression." *Journal of Neuroscience Methods* 125 (1-2): 93–101. https://doi.org/10.1016/S0165-0270(03)00045-1.

Skarnes, William C., Julie E. Moss, Stella M. Hurtley, and Rosa S. P. Beddington. 1995. "Capturing genes encoding membrane and secreted proteins important for mouse development." *Proceedings of the National Academy of Sciences* 92 (14): 6592–6. https://doi.org/10.1073/pnas.92.14.6592.

Skarnes, William C., Barry Rosen, Anthony P. West, Manousos Koutsourakis, Wendy Bushell, Vivek Iyer, Alejandro O. Mujica, et al. 2011. "A conditional knockout resource for the genome-wide study of mouse gene function." *Nature* 474 (7351): 337–42. https://doi.org/10.1038/nature10163.

Smith, Constance M, Terry F. Hayamizu, Jacqueline H. Finger, Susan M. Bello, Ingeborg J. McCright, Jingxia Xu, Richard M. Baldarelli, et al. 2019. "The mouse Gene Expression Database (GXD): 2019 update." *Nucleic Acids Research* 47 (D1): D774–D779. https://doi.org/10.1093/nar/gky922.

Sountoulidis, Alexandros, Andreas Liontos, Hong Phuong Nguyen, Alexandra B Firsova, Athanasios Fysikopoulos, Xiaoyan Qian, Werner Seeger, Erik Sundström, Mats Nilsson, and Christos Samakovlis. 2020. "SCRINSHOT, a spatial method for single-cell resolution mapping of cell states in tissue sections." *bioRxiv*, February, 2020.02.07.938571. https://doi.org/10.1101/2020.02.07.938571.

"SpaRTAN." n.d. Accessed October 7, 2020. https://www.ncl.ac.uk/gcf/spatiallyresolvedtranscriptomics/.

Spradling, A., and G. Rubin. 1982. "Transposition of cloned P elements into Drosophila germ line chromosomes." *Science* 218 (4570): 341–47. https://doi.org/10.1126/science.6289435.

Sprague, Judy. 2003. "The Zebrafish Information Network (ZFIN): the zebrafish model organism database." *Nucleic Acids Research* 31 (1): 241–43. https://doi.org/10.1093/nar/gkg027.

Srivastava, Avi, Laraib Malik, Tom Smith, Ian Sudbery, and Rob Patro. 2019. "Alevin efficiently estimates accurate gene abundances from dscRNA-seq data." *Genome Biology* 20 (1): 65. https://doi.org/10.1186/s13059-019-1670-y.

Stanford, William L., Georgina Caruana, Katherine A. Vallis, Maneesha Inamdar, Michihiro Hidaka, Victoria L. Bautch, and Alan Bernstein. 1998. "Expression Trapping: Identification of Novel Genes Expressed in Hematopoietic and Endothelial Lineages by Gene Trapping in ES Cells." *Blood* 92 (12): 4622–31. https://doi.org/10.1182/blood.V92.12.4622.

Stanford, William L., Jason B. Cohn, and Sabine P. Cordes. 2001. "Gene-trap mutagenesis: past, present and beyond." *Nature Reviews Genetics* 2 (10): 756–68. https://doi.org/10.1038/35093548.

Stapleton, Mark. 2002. "The Drosophila Gene Collection: Identification of Putative Full-Length cDNAs for 70% of D. melanogaster Genes." *Genome Research* 12 (8): 1294–1300. https://doi.org/10.1101/gr.269102.

Ståhl, Patrik L., Fredrik Salmén, Sanja Vickovic, Anna Lundmark, José Fernández Navarro, Jens Magnusson, Stefania Giacomello, et al. 2016. "Visualization and analysis of gene expression in tissue sections by spatial transcriptomics." *Science* 353 (6294): 78–82. https://doi.org/10.1126/science.aaf2403.

Stickels, Robert, Evan Murray, Pawan Kumar, Jilong Li, Jamie Marshall, Daniela Di Bella, Paola Arlotta, Evan Macosko, and Fei Chen. 2020. "Sensitive spatial genome wide expression profiling at cellular resolution." *bioRxiv*, March, 2020.03.12.989806. https://doi.org/10.1101/2020.03.12.989806.

Stoeger, Thomas, Nico Battich, Markus D. Herrmann, Yauhen Yakimovich, and Lucas Pelkmans. 2015. "Computer vision for image-based transcriptomics." *Methods* 85 (September): 44–53. https://doi.org/10.1016/j.ymeth.2015.05.016.

Strell, Carina, Markus M. Hilscher, Navya Laxman, Jessica Svedlund, Chenglin Wu, Chika Yokota, and Mats Nilsson. 2019. "Placing RNA in context and space – methods for spatially resolved transcriptomics." https://doi.org/10.1111/febs.14435.

Stuart, Tim, Andrew Butler, Paul Hoffman, Christoph Hafemeister, Efthymia Papalexi, William M. Mauck, Yuhan Hao, Marlon Stoeckius, Peter Smibert, and Rahul Satija. 2019. "Comprehensive Integration of Single-Cell Data." *Cell* 177 (7): 1888–1902.e21. https://doi.org/10.1016/j.cell.2019.05.031.

Su, Jing, and Qianqian Song. 2020. "DSTG: Deconvoluting Spatial Transcriptomics Data through Graph-based Artificial Intelligence." *bioRxiv*, January, 2020.10.20.347195. https://doi.org/10.1101/2020.10.20.347195.

Sulston, J., Z. Du, K. Thomas, R. Wilson, L. Hillier, R. Staden, N. Halloran, et al. 1992. "The C. elegans genome sequencing project: A beginning." *Nature* 356 (6364): 37–41. https://doi.org/10.1038/356037a0.

Sun, Qian, Sherin Muckatira, Lei Yuan, Shuiwang Ji, Stuart Newfeld, Sudhir Kumar, and Jieping Ye. 2013. "Image-level and group-level models for Drosophila gene expression pattern annotation." *BMC Bioinformatics* 14 (1): 350. https://doi.org/10.1186/1471-2105-14-350.

Sun, Shiquan, Jiaqiang Zhu, and Xiang Zhou. 2020. "Statistical analysis of spatial expression patterns for spatially resolved transcriptomic studies." *Nature Methods* 17 (2): 193–200. https://doi.org/10.1038/s41592-019-0701-7.

Sun, Yu-Chi, Xiaoyin Chen, Stephan Fischer, Shaina Lu, Jesse Gillis, and Anthony M Zador. 2020. "Integrating barcoded neuroanatomy with spatial transcriptional profiling reveals cadherin correlates of projections shared across the cortex." *bioRxiv*, January, 2020.08.25.266460. https://doi.org/10.1101/2020.08.25.266460.

Sundaresan, Venkatesan, Patricia Springer, Thomas Volpe, Samuel Haward, Jonathan D. G. Jones, Caroline Dean, Hong Ma, and Robert Martienssen. 1995. "Patterns of gene action in plant development revealed by enhancer trap and gene trap transposable elements." *Genes & Development* 9 (14): 1797–1810. https://doi.org/10.1101/gad.9.14.1797.

Svensson, Valentine, Eduardo da Veiga Beltrame, and Lior Pachter. 2020. "A curated database reveals trends in single-cell transcriptomics." *Database* 2020 (January). https://doi.org/10.1093/database/baaa073.

Svensson, Valentine, Sarah A. Teichmann, and Oliver Stegle. 2018. "SpatialDE: Identification of spatially variable genes." *Nature Methods* 15 (5): 343–46. https://doi.org/10.1038/nmeth.4636.

Şişecioğlu, Melda, Harun Budak, Lars Geffers, Murat Çankaya, Mehmet Çiftci, Christina Thaller, Gregor Eichele, Ömer İrfan Küfrevioğlu, and Hasan Özdemir. 2015. "A compendium of expression patterns of cholesterol biosynthetic enzymes in the mouse embryo." *Journal of Lipid Research* 56 (8): 1551–9. https://doi.org/10.1194/jlr.M059634.

Takei, Yodai, Jina Yun, Noah Ollikainen, Shiwei Zheng, Nico Pierson, Jonathan White, Sheel Shah, et al. 2020. "Global architecture of the nucleus in single cells by DNA seqFISH+ and multiplexed immunofluorescence." *bioRxiv*, January, 2020.11.29.403055. https://doi.org/10.1101/2020.11.29.403055.

Tanevski, Jovan, Thin Nguyen, Buu Truong, Nikos Karaiskos, Mehmet Eren Ahsen, Xinyu Zhang, Chang Shu, et al. 2020. "Gene selection for optimal prediction of cell position in tissues from single-cell transcriptomics data." *Life Science Alliance* 3 (11): e202000867. https://doi.org/10.26508/lsa.202000867.

Tang, Fuchou, Catalin Barbacioru, Yangzhou Wang, Ellen Nordman, Clarence Lee, Nanlan Xu, Xiaohui Wang, et al. 2009. "mRNA-Seq whole-transcriptome analysis of a single cell." *Nature Methods* 6 (5): 377–82. https://doi.org/10.1038/nmeth.1315.

Tang, Fuchou, Kaiqin Lao, and M Azim Surani. 2011. "Development and applications of single-cell transcriptome analysis." *Nature Methods* 8 (4 Suppl): S6–S11. https://doi.org/10.1038/nmeth.1557.

Tassy, Olivier, Delphine Dauga, Fabrice Daian, Daniel Sobral, François Robin, Pierre Khoueiry, David Salgado, et al. 2010. "The ANISEED database: Digital representation, formalization, and elucidation of a chordate developmental program." *Genome Research* 20 (10): 1459–68. https://doi.org/10.1101/gr.108175.110.

Tautz, Diethard, and Christine Pfeifle. 1989. "A non-radioactive in situ hybridization method for the localization of specific RNAs in Drosophila embryos reveals translational control of the segmentation gene hunchback." *Chromosoma* 98 (2): 81–85. https://doi.org/10.1007/BF00291041.

Tecott, L H, J D Barchas, and J H Eberwine. 1988. "In situ transcription: specific synthesis of complementary DNA in fixed tissue sections." *Science* 240 (4859): 1661 LP–1664. https://doi.org/10.1126/science.2454508.

The C. elegans Sequencing Consortium. 1998. "Genome Sequence of the Nematode C. elegans: A Platform for Investigating Biology." *Science* 282 (5396): 2012–8. https://doi.org/10.1126/science.282.5396.2012.

Thompson, Carol, Jonathan Wisor, Chang-Kyu Lee, Sayan Pathak, Dmitry Gerashchenko, Kimberly Smith, Shanna Fischer, et al. 2010. "Molecular and Anatomical Signatures of Sleep Deprivation in the Mouse Brain." https://www.frontiersin.org/article/10.3389/fnins.2010.00165.

Thrane, Kim, Hanna Eriksson, Jonas Maaskola, Johan Hansson, and Joakim Lundeberg. 2018. "Spatially resolved transcriptomics enables dissection of genetic heterogeneity in stage III cutaneous malignant melanoma." *Cancer Research*. https://doi.org/10.1158/0008-5472.CAN-18-0747.

Thut, Catherine J., Ryan B. Rountree, Michael Hwa, and David M. Kingsley. 2001. "A Large-Scale in Situ Screen Provides Molecular Evidence for the Induction of Eye Anterior Segment Structures by the Developing Lens." *Developmental Biology* 231 (1): 63–76. https://doi.org/10.1006/dbio.2000.0140.

Tietjen, Ian, Jason M Rihel, Yanxiang Cao, Georgy Koentges, Lisa Zakhary, and Catherine Dulac. 2003. "Single-Cell Transcriptional Analysis of Neuronal Progenitors." *Neuron* 38 (2): 161–75. https://doi.org/https://doi.org/10.1016/S0896-6273(03)00229-0.

Tomancak, Pavel, Amy Beaton, Richard Weiszmann, Elaine Kwan, Sheng Qiang Shu, Suzanna E. Lewis, Stephen Richards, et al. 2002. "Systematic determination of patterns of gene expression during Drosophila embryogenesis." *Genome Biology* 3 (12): research0088.1. https://doi.org/10.1186/gb-2002-3-12-research0088.

Tomancak, Pavel, Benjamin P. Berman, Amy Beaton, Richard Weiszmann, Elaine Kwan, Volker Hartenstein, Susan E. Celniker, and Gerald M. Rubin. 2007. "Global analysis of patterns of gene expression during Drosophila embryogenesis." *Genome Biology* 8 (7): R145. https://doi.org/10.1186/gb-2007-8-7-r145.

"Translational Pathology Core Laboratory (TPCL)." n.d. Accessed October 14, 2020. https://www.uclahealth.org/pathology/tpcl-services.

Tripodo, Claudio, Federica Zanardi, Fabio Iannelli, Saveria Mazzara, Mariella Vegliante, Gaia Morello, Arianna Di Napoli, et al. 2020. "A Spatially Resolved Dark- versus Light-Zone Microenvironment Signature Subdivides Germinal Center-Related Aggressive B Cell Lymphomas." *iScience* 23 (10). https://doi.org/10.1016/j.isci.2020.101562.

Tuck, Elizabeth, Jeanne Estabel, Anika Oellrich, A. K. Maguire, Hibret A. Adissu, Luke Souter, Emma Siragher, et al. 2015. "A gene expression resource generated by genome-wide lacZ profiling in the mouse." *Disease Models & Mechanisms* 8 (11): 1467–78. https://doi.org/10.1242/dmm.021238.

Tzur, Yonatan B., Eitan Winter, Jinmin Gao, Tamar Hashimshony, Itai Yanai, and Monica P. Colaiácovo. 2018. "Spatiotemporal Gene Expression Analysis of the Caenorhabditis elegans Germline Uncovers a Syncytial Expression Switch." *Genetics* 210 (2): 587–605. https://doi.org/10.1534/genetics.118.301315.

Urdea, M. S. 1993. "Synthesis and Characterization of Branched DNA (bDNA) for the Direct and Quantitative Detection of CMV, HBV, HCV, and HIV." *Clinical Chemistry* 39 (4): 725–26. https://doi.org/10.1093/clinchem/39.4.725.

Valoczi, A. 2004. "Sensitive and specific detection of microRNAs by northern blot analysis using LNA-modified oligonucleotide probes." *Nucleic Acids Research* 32 (22): e175–e175. https://doi.org/10.1093/nar/gnh171.

Vandenbon, Alexis, and Diego Diez. 2020. "A clustering-independent method for finding differentially expressed genes in single-cell transcriptome data." *Nature Communications* 11 (1): 4318. https://doi.org/10.1038/s41467-020-17900-3.

Van Gelder, R N, M E von Zastrow, A Yool, W C Dement, J D Barchas, and J H Eberwine. 1990. "Amplified RNA synthesized from limited quantities of heterogeneous cDNA." *Proceedings of the National Academy of Sciences* 87 (5): 1663 LP–1667. http://www.pnas.org/content/87/5/1663.abstract.

Van Valen, David A, Takamasa Kudo, Keara M Lane, Derek N Macklin, Nicolas T Quach, Mialy M DeFelice, Inbal Maayan, Yu Tanouchi, Euan A Ashley, and Markus W Covert. 2016. "Deep Learning Automates the Quantitative Analysis of Individual Cells in Live-Cell Imaging Experiments." *PLOS Computational Biology* 12 (11): e1005177. https://doi.org/10.1371/journal.pcbi.1005177.

Venter, J. Craig, Mark D. Adams, Eugene W. Myers, Peter W. Li, Richard J. Mural, Granger G. Sutton, Hamilton O. Smith, et al. 2001. "The Sequence of the Human Genome." *Science* 291 (5507): 1304–51. https://doi.org/10.1126/science.1058040.

"Veritas Laser Capture Microdissection (LCM) and Laser Cutting System from Applied Biosystems." n.d. Accessed October 14, 2020. https://www.unthsc.edu/research/flow-cytometry-and-laser-capture-microdissection-core-facility/beckman-coulter-cytomics-fc500-flow-cytometry-analyzer/veritas-laser-capture.

Verma, Archit, and Barbara Engelhardt. 2020. "A Bayesian nonparametric semi-supervised model for integration of multiple single-cell experiments." *bioRxiv*, January, 2020.01.14.906313. https://doi.org/10.1101/2020.01.14.906313.

Vickovic, Sanja, Gökcen Eraslan, Fredrik Salmén, Johanna Klughammer, Linnea Stenbeck, Denis Schapiro, Tarmo Äijö, et al. 2019. "High-definition spatial transcriptomics for in situ tissue profiling." *Nature Methods*. https://doi.org/10.1038/s41592-019-0548-y.

Vickovic, Sanja, Britta Lötstedt, Johanna Klughammer, Åsa Segerstolpe, Orit Rozenblatt-Rosen, and Aviv Regev. 2020. "SM-Omics: An automated platform for high-throughput spatial multi-omics." *bioRxiv*, January, 2020.10.14.338418. https://doi.org/10.1101/2020.10.14.338418.

Villacampa, Eva Gracia, Ludvig Larsson, Linda Kvastad, Alma Andersson, Joseph Carlson, and Joakim Lundeberg. 2020. "Genome-wide Spatial Expression Profiling in FFPE Tissues." *bioRxiv*, July, 2020.07.24.219758. https://doi.org/10.1101/2020.07.24.219758.

Visel, Axel, Leila Taher, Hani Girgis, Dalit May, Olga Golonzhka, Renee V. Hoch, Gabriel L. McKinsey, et al. 2013. "A high-resolution enhancer atlas of the developing telencephalon." *Cell* 152 (4): 895–908. https://doi.org/10.1016/j.cell.2012.12.041.

Waldhaus, Jörg, Robert Durruthy-Durruthy, and Stefan Heller. 2015. "Quantitative High-Resolution Cellular Map of the Organ of Corti." *Cell Reports* 11 (9): 1385–99. https://doi.org/10.1016/j.celrep.2015.04.062.

Wang, Chong, Tian Lu, George Emanuel, Hazen P. Babcock, and Xiaowei Zhuang. 2019. "Imaging-based pooled CRISPR screening reveals regulators of lncRNA localization." *Proceedings of the National Academy of Sciences of the United States of America*. https://doi.org/10.1073/pnas.1903808116.

Wang, Fay, John Flanagan, Nan Su, Li-Chong Wang, Son Bui, Allissa Nielson, Xingyong Wu, Hong-Thuy Vo, Xiao-Jun Ma, and Yuling Luo. 2012. "RNAscope." *The Journal of Molecular Diagnostics* 14 (1): 22–29. https://doi.org/10.1016/j.jmoldx.2011.08.002.

Wang, Guiping, Jeffrey R. Moffitt, and Xiaowei Zhuang. 2018. "Multiplexed imaging of high-density libraries of RNAs with MERFISH and expansion microscopy." *Scientific Reports*. https://doi.org/10.1038/s41598-018-22297-7.

Wang, Xiao, William E. Allen, Matthew A. Wright, Emily L. Sylwestrak, Nikolay Samusik, Sam Vesuna, Kathryn Evans, et al. 2018. "Three-dimensional intact-tissue sequencing of single-cell transcriptional states." *Science* 361 (6400): eaat5691. https://doi.org/10.1126/science.aat5691.

Wang, Yuhan, Mark Eddison, Greg Fleishman, Martin Weigert, Shengjin Xu, Fredrick E Henry, Tim Wang, et al. 2021. "Expansion-Assisted Iterative-FISH defines lateral hypothalamus spatio-molecular organization." *bioRxiv*, January, 2021.03.08.434304. https://doi.org/10.1101/2021.03.08.434304.

Waterston, Robert H., Kerstin Lindblad-Toh, Ewan Birney, Jane Rogers, Josep F. Abril, Pankaj Agarwal, Richa Agarwala, et al. 2002. "Initial sequencing and comparative analysis of the mouse genome." *Nature* 420 (6915): 520–62. https://doi.org/10.1038/nature01262.

Waylen, Lisa N, Hieu T Nim, Luciano G Martelotto, and Mirana Ramialison. 2020. "From whole-mount to single-cell spatial assessment of gene expression in 3D." *Communications Biology* 3 (1): 602. https://doi.org/10.1038/s42003-020-01341-1.

Weinstein, Joshua A., Aviv Regev, and Feng Zhang. 2019. "DNA Microscopy: Optics-free Spatio-genetic Imaging by a Stand-Alone Chemical Reaction." *Cell* 178 (1): 229–241.e16. https://doi.org/10.1016/j.cell.2019.05.019.

Welch, Joshua D., Velina Kozareva, Ashley Ferreira, Charles Vanderburg, Carly Martin, and Evan Z. Macosko. 2019. "Single-Cell Multi-omic Integration Compares and Contrasts Features of Brain Cell Identity." *Cell* 177 (7): 1873–1887.e17. https://doi.org/10.1016/j.cell.2019.05.006.

West, David B., Ravi K. Pasumarthi, Brian Baridon, Esi Djan, Amanda Trainor, Stephen M. Griffey, Eric K. Engelhard, et al. 2015. "A lacZ reporter gene expression atlas for 313 adult KOMP mutant mouse lines." *Genome Research* 25 (4): 598–607. https://doi.org/10.1101/gr.184184.114.

Westerfield, Monte, Eckehard Doerry, Arthur E. Kirkpatrick, Wolfgang Driever, and Sarah A. Douglas. 1997. "An on-line database for zebrafish development and genetics research." *Seminars in Cell & Developmental Biology* 8 (5): 477–88. https://doi.org/10.1006/scdb.1997.0173.

White, Jacqueline K., Anna-Karin Gerdin, Natasha A. Karp, Ed Ryder, Marija Buljan, James N. Bussell, Jennifer Salisbury, et al. 2013. "Genome-wide

Generation and Systematic Phenotyping of Knockout Mice Reveals New Roles for Many Genes." *Cell* 154 (2): 452–64. https://doi.org/10.1016/j.cell.2013.06.022.

Wienholds, Erno. 2005. "MicroRNA Expression in Zebrafish Embryonic Development." *Science* 309 (5732): 310–11. https://doi.org/10.1126/science.1114519.

Wilson, Clive, Rebecca Kurth Pearson, Hugo J Bellen, C J O'Kane, Ueli Grossniklaus, and Walter J Gehring. 1989. "P-element-mediated enhancer detection: an efficient method for isolating and characterizing developmentally regulated genes in Drosophila." *Genes & Development* 3 (9): 1301–13. https://doi.org/10.1101/gad.3.9.1301.

Wu, Siqi, Antony Joseph, Ann S. Hammonds, Susan E. Celniker, Bin Yu, and Erwin Frise. 2016. "Stability-driven nonnegative matrix factorization to interpret spatial gene expression and build local gene networks." *Proceedings of the National Academy of Sciences* 113 (16): 4290–5. https://doi.org/10.1073/pnas.1521171113.

Wurst, W, J Rossant, V Prideaux, M Kownacka, A Joyner, D P Hill, F Guillemot, S Gasca, D Cado, and A Auerbach. 1995. "A large-scale gene-trap screen for insertional mutations in developmentally regulated genes in mice." *Genetics* 139 (2). https://www.genetics.org/content/139/2/889.

"XDB3." 2004. http://xenopus.nibb.ac.jp/.

Xia, Chenglong, Hazen P. Babcock, Jeffrey R. Moffitt, and Xiaowei Zhuang. 2019. "Multiplexed detection of RNA using MERFISH and branched DNA amplification." *Scientific Reports* 9 (1): 1–13. https://doi.org/10.1038/s41598-019-43943-8.

Xia, Chenglong, Jean Fan, George Emanuel, Junjie Hao, and Xiaowei Zhuang. 2019. "Spatial transcriptome profiling by MERFISH reveals subcellular RNA compartmentalization and cell cycle-dependent gene expression." *Proceedings of the National Academy of Sciences of the United States of America*. https://doi.org/10.1073/pnas.1912459116.

Xu, Xiangmin, Todd C Holmes, Min-Hua Luo, Kevin T Beier, Gregory D Horwitz, Fei Zhao, Wenbo Zeng, May Hui, Bert L Semler, and Rozanne M Sandri-Goldin. 2020. "Viral Vectors for Neural Circuit Mapping and Recent Advances in Trans-synaptic Anterograde Tracers." *Neuron* 107 (6): 1029–47. https://doi.org/https://doi.org/10.1016/j.neuron.2020.07.010.

Yang, Xiangdong W., Peter Model, and Nathaniel Heintz. 1997. "Homologous recombination based modification in Esherichia coli and germline transmission in transgenic mice of a bacterial artificial chromsome." *Nature Biotechnology* 15 (9): 859–65. https://doi.org/10.1038/nbt0997-859.

Yang, Xiaoyu, Seth Bergenholtz, Lenka Maliskova, Mark-Phillip Pebworth, Arnold R Kriegstein, Yun Li, and Yin Shen. 2020. "SMART-Q: An Integrative

Pipeline Quantifying Cell Type-Specific RNA Transcription." *PLOS ONE* 15 (4): e0228760. https://doi.org/10.1371/journal.pone.0228760.

Yang, Yang, Qingwei Fang, and Hong-Bin Shen. 2019. "Predicting gene regulatory interactions based on spatial gene expression data and deep learning." Edited by Smita Krishnaswamy. *PLOS Computational Biology* 15 (9): e1007324. https://doi.org/10.1371/journal.pcbi.1007324.

Yaylaoglu, Murat Burak, Andrew Titmus, Axel Visel, Gonzalo Alvarez-Bolado, Christina Thaller, and Gregor Eichele. 2005. "Comprehensive expression atlas of fibroblast growth factors and their receptors generated by a novel robotic in situ hybridization platform." *Developmental Dynamics* 234 (2): 371–86. https://doi.org/10.1002/dvdy.20441.

Yoda, Takuya, Masahito Hosokawa, Kiyofumi Takahashi, Chikako Sakanashi, Haruko Takeyama, and Hideki Kambara. 2017. "Site-specific gene expression analysis using an automated tissue micro-dissection punching system." *Scientific Reports* 7 (1): 4325. https://doi.org/10.1038/s41598-017-04616-6.

Yokoyama, Shigetoshi, Yoshiaki Ito, Hiroe Ueno-Kudoh, Hirohito Shimizu, Kenta Uchibe, Sonia Albini, Kazuhiko Mitsuoka, et al. 2009. "A Systems Approach Reveals that the Myogenesis Genome Network Is Regulated by the Transcriptional Repressor RP58." *Developmental Cell* 17 (6): 836–48. https://doi.org/10.1016/j.devcel.2009.10.011.

Yoshikawa, Toshiyuki, Yulan Piao, Jinhui Zhong, Ryo Matoba, Mark G. Carter, Yuxia Wang, Ilya Goldberg, and Minoru S. H. Ko. 2006. "High-throughput screen for genes predominantly expressed in the ICM of mouse blastocysts by whole mount in situ hybridization." *Gene Expression Patterns* 6 (2): 213–24. https://doi.org/10.1016/j.modgep.2005.06.003.

Yuan, Lei, Alexander Woodard, Shuiwang Ji, Yuan Jiang, Zhi-Hua Zhou, Sudhir Kumar, and Jieping Ye. 2012. "Learning Sparse Representations for Fruit-Fly Gene Expression Pattern Image Annotation and Retrieval." *BMC Bioinformatics* 13 (1): 107. https://doi.org/10.1186/1471-2105-13-107.

Yuan, Ye, and Ziv Bar-Joseph. 2019. "GCNG: Graph convolutional networks for inferring cell-cell interactions." *bioRxiv*, January, 2019.12.23.887133. https://doi.org/10.1101/2019.12.23.887133.

Zechel, Sabrina, Pawel Zajac, Peter Lönnerberg, Carlos F. Ibáñez, and Sten Linnarsson. 2014. "Topographical transcriptome mapping of the mouse medial ganglionic eminence by spatially resolved RNA-seq." *Genome Biology* 15 (10): 486. https://doi.org/10.1186/s13059-014-0486-z.

Zeisel, Amit, Hannah Hochgerner, Peter Lönnerberg, Anna Johnsson, Fatima Memic, Job van der Zwan, Martin Häring, et al. 2018. "Molecular Architecture of the Mouse Nervous System." *Cell* 174 (4): 999–1014.e22. https://doi.org/10.1016/j.cell.2018.06.021.

Zeng, Tao, Rongjian Li, Ravi Mukkamala, Jieping Ye, and Shuiwang Ji. 2015. "Deep convolutional neural networks for annotating gene expression patterns in the mouse brain." *BMC Bioinformatics* 16 (1): 147. `https://doi.org/10.1186/s12859-015-0553-9`.

Zhang, Ke, Wanwan Feng, and Peng Wang. 2018. "Identification of spatially variable genes with graph cuts." *bioRxiv*, January, 491472. `https://doi.org/10.1101/491472`.

Zhang, Meng, Stephen W Eichhorn, Brian Zingg, Zizhen Yao, Hongkui Zeng, Hongwei Dong, and Xiaowei Zhuang. 2020. "Molecular, spatial and projection diversity of neurons in primary motor cortex revealed by in situ single-cell transcriptomics." *bioRxiv*, January, 2020.06.04.105700. `https://doi.org/10.1101/2020.06.04.105700`.

Zhang, Wenlu, Daming Feng, Rongjian Li, Andrey Chernikov, Nikos Chrisochoides, Christopher Osgood, Charlotte Konikoff, Stuart Newfeld, Sudhir Kumar, and Shuiwang Ji. 2013. "A mesh generation and machine learning framework for Drosophila gene expression pattern image analysis." *BMC Bioinformatics* 14 (1): 372. `https://doi.org/10.1186/1471-2105-14-372`.

Zhao, Edward, Matthew R Stone, Xing Ren, Thomas Pulliam, Paul Nghiem, Jason H Bielas, and Raphael Gottardo. 2020. "BayesSpace enables the robust characterization of spatial gene expression architecture in tissue sections at increased resolution." *bioRxiv*, January, 2020.09.04.283812. `https://doi.org/10.1101/2020.09.04.283812`.

Zhao, Tuo, Han Liu, Kathryn Roeder, John Lafferty, and Larry Wasserman. 2012. "The Huge Package for High-Dimensional Undirected Graph Estimation in R." *J. Mach. Learn. Res.* 13 (null): 1059–62.

Zheng, Grace X. Y., Jessica M. Terry, Phillip Belgrader, Paul Ryvkin, Zachary W. Bent, Ryan Wilson, Solongo B. Ziraldo, et al. 2017. "Massively parallel digital transcriptional profiling of single cells." *Nature Communications* 8 (1): 14049. `https://doi.org/10.1038/ncomms14049`.

Zhou, Chuan, Ru Huang, Xiaoming Zhou, and Da Xing. 2020. "Sensitive and specific microRNA detection by RNA dependent DNA ligation and rolling circle optical signal amplification." *Talanta* 216: 120954. `https://doi.org/https://doi.org/10.1016/j.talanta.2020.120954`.

Zhou, Jie, and Hanchuan Peng. 2007. "Automatic recognition and annotation of gene expression patterns of fly embryos." *Bioinformatics* 23 (5): 589–96. `https://doi.org/10.1093/bioinformatics/btl680`.

Zhou, Wen, Mary A. Yui, Brian A. Williams, Jina Yun, Barbara J. Wold, Long Cai, and Ellen V. Rothenberg. 2019. "Single-Cell Analysis Reveals Regulatory Gene Expression Dynamics Leading to Lineage Commitment in Early T Cell Development." *Cell Systems* 9 (4): 321–337.e9. `https://doi.org/10.1016/j.cels.2019.09.008`.

Zhu, Junjie, and Chiara Sabatti. 2020. "Integrative Spatial Single-cell Analysis with Graph-based Feature Learning." *bioRxiv*, January, 2020.08.12.248971. https://doi.org/10.1101/2020.08.12.248971.

Zhu, Qian, Sheel Shah, Ruben Dries, Long Cai, and Guo Cheng Yuan. 2018. "Identification of spatially associated subpopulations by combining scRNAseq and sequential fluorescence in situ hybridization data." *Nature Biotechnology* 36 (12): 1183–90. https://doi.org/10.1038/nbt.4260.

Zhu, Y Y, E M Machleder, A Chenchik, R Li, and P D Siebert. 2001. "Reverse Transcriptase Template Switching: A SMART™ Approach for Full-Length cDNA Library Construction." *BioTechniques* 30 (4): 892–97. https://doi.org/10.2144/01304pf02.
